# Genetic affinities between an ancient Greek colony and its metropolis: the case of Amvrakia in western Greece

**DOI:** 10.1101/2025.07.01.662689

**Authors:** Nikolaos Psonis, Eugenia Tabakaki, Despoina Vassou, Stefanos Papadantonakis, Angelos Souleles, Argyro Nafplioti, Georgios Kousis Tsampazis, Angeliki Papadopoulou, Kiriakos Xanthopoulos, Panagiotis Panailidis, Angeliki Georgiadou, Dimitra Papakosta, Sevasti Koursioti, Maria Evangelinou, Varvara Papadopoulou, Paraskevi Evaggeloglou, Elena Korka, Ioannis Christidis, Michael Ioannou, Theodora Kontogianni, Athanasios Arkoumanis, Alexandros Stamatakis, Nikos Poulakakis, Christina Papageorgopoulou, Pavlos Pavlidis

## Abstract

**Background:** During the Ancient Greek colonization, Corinth established a stable network of economic and political ties, by founding colonies connecting southern Greece with the mainland of Epirus and reaching as far as the east Adriatic coast. Amvrakia, one of the main Corinthian colonies founded during the 7th century BCE, was characterized by its strong dependence on its metropolis. Here, we aim to investigate the genetic relationships between the Corinthian metropolis and the Amvrakia colony, the contribution of the local population to the founding genetic pool, as well as the demography of Amvrakia in subsequent periods.

**Results:** During its foundation in the Archaic period, Amvrakia appears to have been shaped by genetic influences from at least two different sources. The first source migrated from the Corinth territory, represented by the Archaic Tenea population and is supported via an Identity By Descent (IBD) analysis. The second source shows a direct ancestry from Late Bronze Age (LBA) / Iron Age Greece, including a local LBA population represented by the Ammotopos site located in close proximity to Amvrakia, as shown by a plethora of independent population genomics analyses. During the subsequent Classical and Hellenistic periods, the population of Amvrakia appears to have slightly differentiated, yet evidence of genetic continuity over time is observed.

**Conclusions:** The migration of Corinthians to Amvrakia contributed to the initial genetic pool of the colony along with the local genetic pool, indicating that the Corinthian colonization included both genetic and cultural transmission between the metropolis and its colony.

## Background

Migrations have been a pervasive element throughout human history. From the out-of-Africa migrations, through the Neolithic demographic transition and the anthropocene, there has been a continuous flow of migratory movements globally. A prominent outcome of such migrations and local interactions is the ancient Greek civilization. Originating in the post-Mycenaean era that is often referred to as Greek Dark Ages, it is widely accepted that the ancient Greek civilization began to solidify during the Archaic period (circa 800-479 BCE) as a consequence of migratory movements and interactions within the Mediterranean and Black Sea regions, a phenomenon known as ancient Greek colonization.

During the ancient Greek colonization, populations from the Aegean Islands, Asia Minor, and continental Greece embarked on numerous expeditions [1–5]. Their search for new homes had numerous reasons: necessity due to internal strife, social conflicts, political strategies, famine or poverty, and the pursuit of opportunities (e.g., new land to farm, more livestock to own, ampler natural resources to exploit). In some cases, the *Emporia (trading posts*) that were established to explore new markets were the predecessor of colonies. From the mid-8th century BCE onwards, the Greek *poleis* (cities), such as Corinth and Eretria, started to expand in a more strategic and sustainable manner. This is particularly hold for Corinth, which evolved into one of the most powerful metropoles during the era of the ancient Greek Colonization through the establishment of *Apoikiai* (colonies) and trading posts across the Mediterranean (Corcyra and Leukas in the Ionian Sea, Syracuse in Sicily), the Adriatic (Epidamnus, Apollonia), the North Aegean (Potidaea in Chalcidice), and mainland Greece (Amvrakia in Epirus, Anaktorion and Sollion in the western part of the geographical unit of Sterea Ellada) (**Figure 1A**). Through its colonies, Corinth developed a trade network encompassing the Aegean, the coasts of Western Greece, the Adriatic, and Sicily [6]. Corinth substantially influenced the lifestyle and the economy of its colonies in the Mediterranean region [7–9].

**Figure 1.**
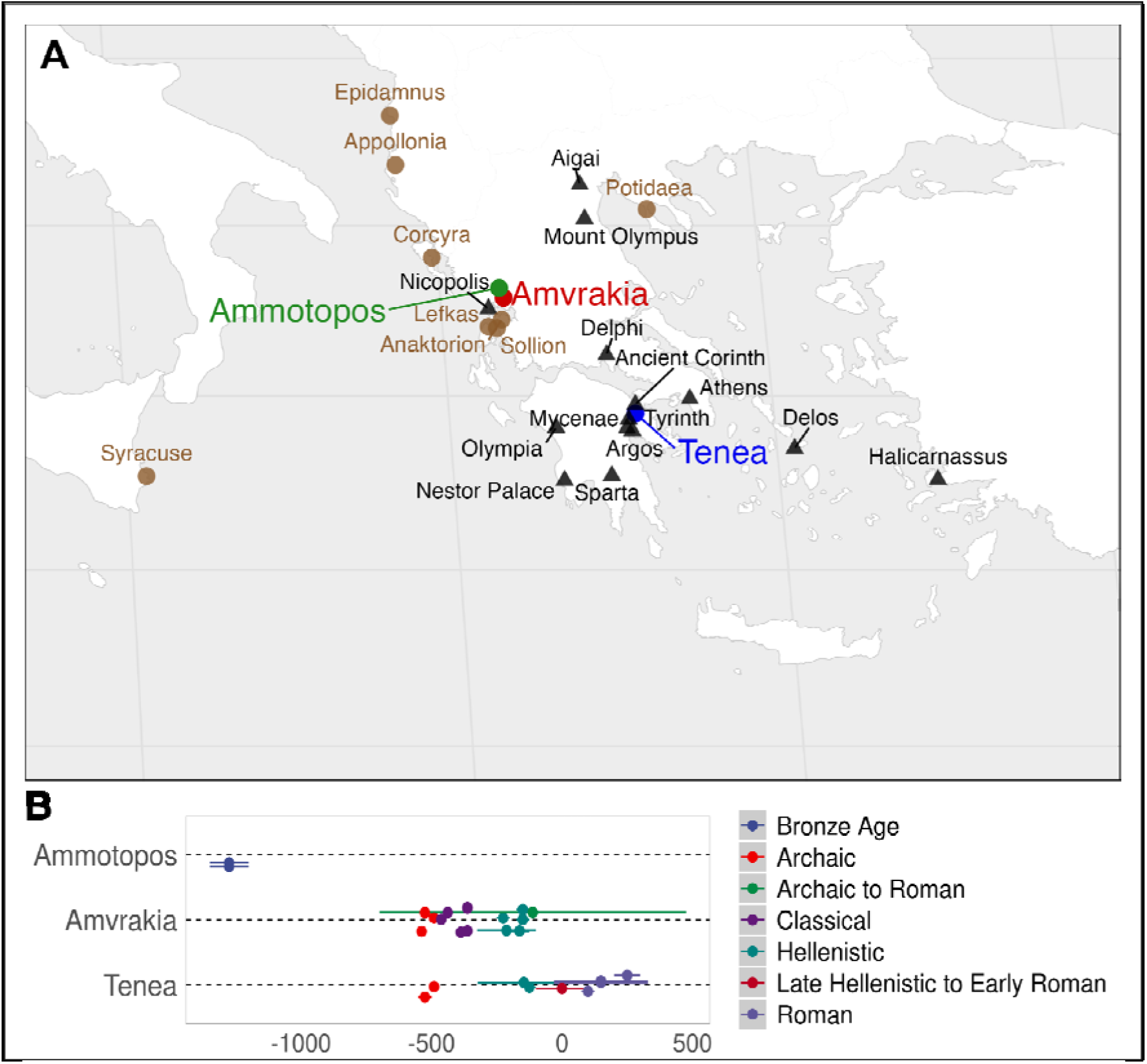
**A**. Location of Ammotopos (green), Amvrakia (red), and Tenea (blue) in the present-day geographical area of Greece. The main Corinthian colonies are displayed in brown color. Other important ancient locations, as well as locations mentioned in the text are also displayed as black triangles. **B.** Temporal coverage of the 26 ancient genomes reported for the first type by our study.

The colony of Amvrakia (also known as Ambracia) was part of this strategic colonization plan. It was founded on the banks of the Arachthos River [10], in the area of present-day northwestern Greece (Epirus regional unit; present-day city of Arta; **Figure 1A**), in the second half of the 7th century BCE. In this region, Corinth had established a trading post, as early as in the 8th century BCE [11], which connected southern Greece with the mainland of Epirus and extended to Apollonia, another Corinthian colony on the eastern Adriatic coast. The foundation of Amvrakia by Corinth is substantiated by the strong presence of Corinthian material culture in the ancient city. Historical literature corroborates the early dependence of Amvrakia from its metropolis [12].

Overall, the relationships between metropoles and their colonies were bidirectional and generally mutually beneficial, with some colonies surpassing their metropolis in cultural and political advancements. Each colony has its unique history, founding myth, potential subsequent establishment of its own colonies, allies, and distinct ties to the metropolis. Despite these individual characteristics, commonalities often exist in the founding myths of most colonies [13]. Based on these elements, archaeologists and historians identify a colony in relation to the metropolis, although this connection is not always straightforward to establish.

While many colonies have been excavated, the debate on the mode, intensity, and tempo of the migratory movements [e.g. 14,15] is still ongoing. Did it occur more like a drift or as an organized colonization? Can we indeed identify biological, linguistic, religious, cultural, or social groups whose origins might as often have been merely invented to establish founding myths as they might have been real? What was the role of the local populations, were they subjects of assimilation or differentiation? The nuclei of settlers attracted numerous others and perhaps local women. What was their role? Many of these open questions are anthropological and population genetic in nature and have been only limited addressed.

Nowadays, ancient colonization can be studied via modern methodological approaches, such as ancient DNA and strontium isotope studies in conjunction with archaeological evidence and literary sources. Up to now, archaeogenomic investigations of ancient Greek colonization are limited to two studies. The first included genome-wide data from inhabitants of the Greek colony of Empúries (Εμπόριον in Greek) in Iberia dated to the the 5th century BCE, who display genetic proximity to both, Late Bronze Age (LBA) Aegean people, and indigenous populations [16]. The second study focused on soldiers who fought for the army of the Greek Sicilian colony of Himera, along with representatives of the civilian population, where both groups included individuals with an LBA Aegean ancestry [17]. Both studies used prehistoric Aegean individuals as a proxy for the ancestral profile of ancient Greeks. This is due to the scarcity of available ancient genomes or genome-wide data from the Eastern Mediterranean covering the period from the Greek Archaic era to Roman Imperial times. The first historical genome-wide data from Greece were generated only recently including four Iron Age (IA) individuals from mainland Greece (Kastrouli near Delphi) [18,19] and the Peloponnese (Palace of Nestor and Tiryns) and one individual from Roman times (Marathon, Attica) [18].

In our study, we focus on the Corinthian colony of Amvrakia to investigate the genetic relationships between a metropolis and its apoikia. For the first time, we assess the contribution of the local population and the colonists to the genetic pool of the founded city. We further examine its demographic evolution from its foundation during the Archaic period until its complete independence in Hellenistic times. Moreover, the assembly of genomes from these yet unsampled periods in the area of present-day Greece also allows for a comparison with preceding prehistoric genetic data from the Eastern Mediterranean, as well as to get insights on their phenotypic characteristics and identify ancient DNA traces from their commensal or pathogenic microbes. Therefore, we sampled ancient bones and teeth from 24 individuals from Archaic, Classical, and Hellenistic Amvrakia (**Figure 1)**, as well as from two individuals from the LBA site of Ammotopos (**Figure 1**) that is located a few kilometers from Amvrakia. The Ammotopos site represents local ancestry before the founding period. Moreover, as a proxy for the genetic pool of Corinth, we sampled 13 individuals from Archaic up to Roman times from the ancient city of Tenea (**Figure 1**), an important settlement located in the eastern periphery of Corinth and in a key strategic position controlling *Kontoporeia*, the shortest path leading from Corinth to Mycenae, and to Argos. Tenea has participated in Corinthian colonization (e.g., to found Syracuse according to ancient written sources [20]).

## Results

We performed pre-screening shotgun NGS analysis on 41 samples from 39 individuals of Amvrakia, Ammotopos, and Tenea burials (**Additional file 1)**. We performed deep whole genome sequencing on 26 out of the 41 samples based on the useful human endogenous content of each sample/library (**Additional file 2)**. All individuals’ genetic data were characterized by an ancient-like DNA signature (i.e., deamination damage and short fragments length; **Additional file 3)**. The final dataset included 15 individuals from Archaic (3), Classical (6), Hellenistic (5), and Archaic-to-Roman (1) Amvrakia, two individuals from LBA Ammotopos, and nine individuals from Archaic (2), Hellenistic (2), Late Hellenistic-to-Early Roman (1), and Roman (4) Tenea (**Figure 1; Additional file 1)**. The mean coverage depth exceeded 0.05× (0.07 - 6.31×) for all 26 individuals (**Additional file 2)**, nine individuals were males and 17 were females (**Additional file 3)**. Contamination evaluation resulted in >98.49% estimated average authenticity levels using ContamMix, a contamination estimate below 4% using schmutzi, and ANGSD contamination estimates below 1.62% based on the X-chromosome in males (**Additional file 3)**. Various haplogroups were detected (**Additional file 3)** regarding both the mitochondrial DNA (J and T major haplogroups in LBA Ammotopos; H, T, and U, in Archaic Amvrakia; H, N, K, and T in Classical Amvrakia; H, J, and W in Hellenistic Amvrakia; T in Archaic Tenea; T and U in Hellenistic Tenea; N and U in Roman Tenea) and the Y-chromosome (G major haplogroup in LBA Ammotopos; J and T in Classical Amvrakia; E in Hellenistic Amvrakia; T in Archaic Tenea; E in Hellenistic Tenea; R and J in Roman Tenea). We further discovered a few cases of close genetic relatedness (see details in **Additional file 7, Section 3.1**). We detected one in Classical Amvrakia [second degree genetic kinships between a female individual (Amv_Epi_Cl_1) and two other individuals that were sisters (Amv_Epi_Cl_5 and Amv_Epi_Cl_6)], one in Hellenistic Amvrakia (mother and daughter: Amv_Epi_Hel_3 and Amv_Epi_Hel_4), and one in Archaic Tenea (mother and son: Ten_Pel_Arch_2 and Ten_Pel_Arch_1). In all downstream kinship-affected analyses (i.e. ADMIXTURE, F-statistics, IBDs), we controlled for the aforementioned relatedness cases by only keeping one single individual with higher coverage for each genetically related group of individuals at first and second degree.

### Comparison between historical and prehistoric genomes from the geographic area of present-day Greece

We grouped the ancient individuals (newly sequenced and “Dataset 1”) primarily according to their main cultural chronology and geography (countries and main regions within) (**Additional file 4**; see also details and rationale about grouping in **Additional file 7, sections 3.5.7-3.5.9**). We computed principal components (PCs) on the Human Origins (HO) SNP set data (“Dataset 3”) [21] using 888 modern genomes from West Eurasia and projected the 26 newly generated as well as 663 published ancient genomes onto the first two PCs (**Figure 2A**). Our new genomes are projected among other ancient eastern Mediterranean genomes. However, the new genomes do not form a tight cluster as they are distributed among several prehistoric (Neolithic to Iron Age) and historical (Archaic to Roman times) individuals from a wide geographical area (Italy, Balkans, Anatolia) with two main clusters emerging (see **Additional file 7, Section 3.5.9** for a detailed presentation of the PCA results). The first cluster includes the majority of the Amvrakia samples (one of the three Archaic, all six Classical, and four of the five Hellenistic samples), the two BA Ammotopos samples, and some of the Tenea samples (both Archaic, two of the three Hellenistic, and the Late Hellenistic-Early Roman). This cluster is primarily close to LBA (1700-1050 BCE) genomes from across present-day Greece, Early Iron Age (1100-500 BCE) genomes from present-day Bulgaria, and the locals of the Ancient Greek colony of Himera in Sicily (780-400 BCE). The second cluster comprises all four Roman Tenea individuals and one of the three Hellenistic Amvrakia individuals. It is placed close to Archaic (750-480 BCE), Roman (27-476 CE), and other Anatolian genomes [including a Hellenistic (510-30 BCE) and an Early Bronze Age (EBA, 3350-2000 BCE) genome], a Roman Imperial (1-530 CE) genome from Italy, and a few Greek genomes from the Roman (250-400 CE), EBA_MBA (2900-1700 BCE), and LBA (1700-1050 BCE) periods. We executed an unsupervised ADMIXTURE analysis on “Dataset 2” [22] for a set of 679 ancient individuals on the 1240K SNP set data [21], including 22 of our newly reported genomes (22 remain after removing close relatives) and 657 ancient genomes, for K=2…10 clusters, where K is the number of putative admixture sources. We display the results for K=3, which is the K with the second lowest cross validation errors (only slightly higher than K=2; see **Additional file 7, Supplementary Figure S34**). The setting of K=3 is at the same time the lowest K that allows to differentiate among genetic clusters associated with three key European ancestral components, namely, Western hunter-gatherers (WHGs), early European farmers (EEFs), and Caucasus hunter-gatherers (CHGs), shown in red, orange, and blue color, respectively, in **Figure 2B**. The majority (19 out of 22) of the newly generated genomes have ancestry proportions that resemble those of previously published LBA and IA genomes from mainland Greece and the Peloponnese, whereas the remaining ones resemble that of Roman times Greece and Italy.

**Figure 2.**
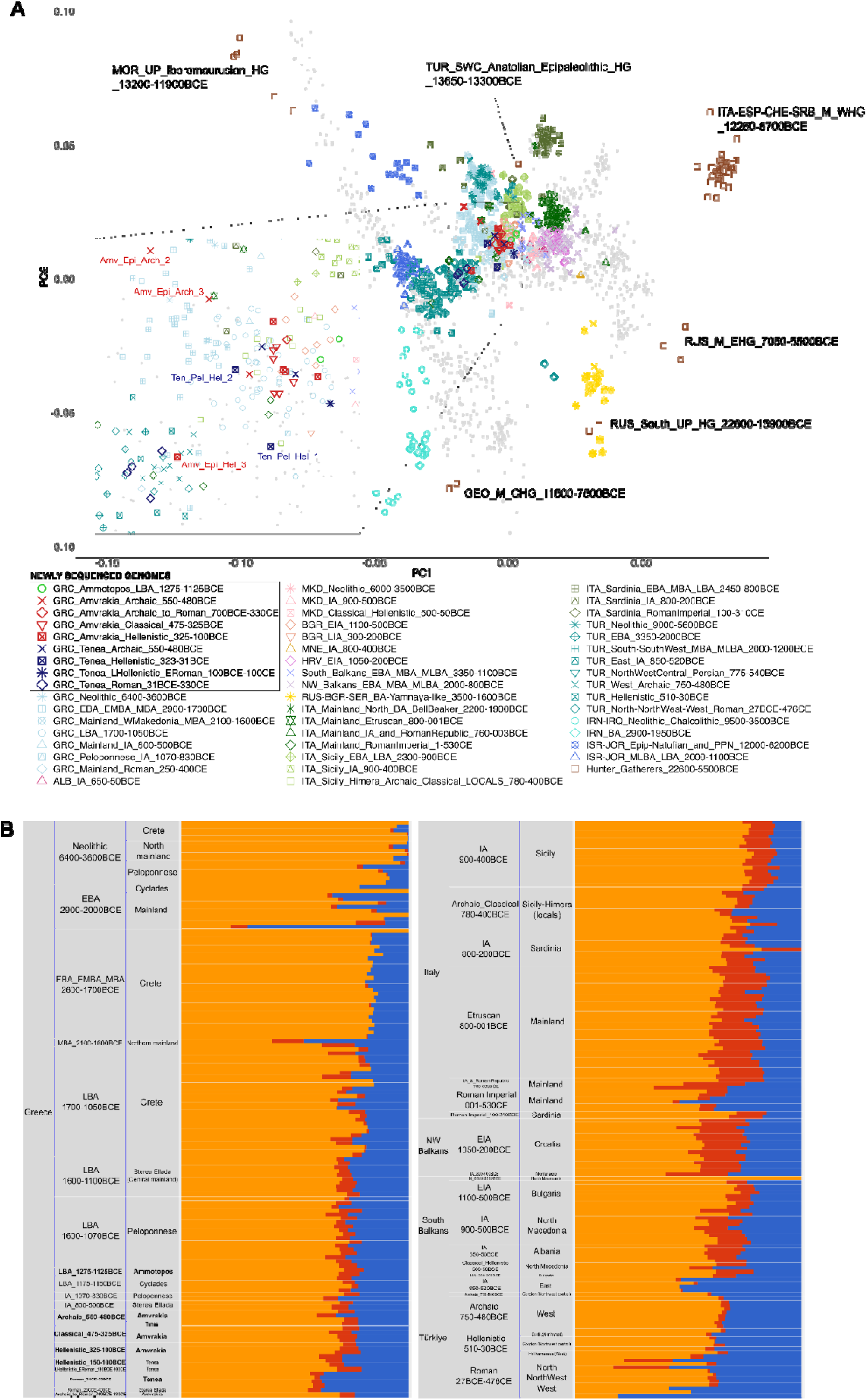
**A.** PCA projection of the 26 newly generated ancient genomes of the present study and 663 other ancient published genomes into the two first PCs generated using 888 genomes from modern West Eurasians (“Dataset 2”). Hunter-Gatherers are indicated, separately, within the plot. South_Balkans includes Albania, North Macedonia, and Bulgaria, whereas NW_Balkans includes Croatia, Montenegro, and Serbia. The analysis was based on 597573 genomic sites of the Human Origins SNP array. GRC=Greece; ALB=Albania; MKD=North Makedonia; BGR=Bulgaria; MNE=Montenegro; HRV=Croatia; RUS=Russia; SER=Serbia; ITA=Italy, TUR=Türkiye, IRN=Iran, IRQ=Iraq, ISR=Israel, JOR=Jordan; BA=Bronze Age; EBA=Early Bronze Age; MBA=Middle Bronze Age; MLBA=Middle-Late Bronze Age; LBA=Late Bronze Age; IA=Iron Age; EIA=Early Iron Age; LIA=Late Iron Age; PPN=Pre-Pottery Neolithic; Epip=Epi-Paleolithic. **B.** Unsupervised ADMIXTURE analysis of 22 newly generated ancient genomes of the present study (four individuals were excluded due to genetic kinship) and 657 other ancient published genomes (“Dataset 1” without six Upper Paleolithic Moroccans) using K=3, LD r^2^ threshold of 0.60, and allele missingness threshold of 40%. K=3 differentiate genetic clusters associated with three key European ancestral components, namely, Western hunter-gatherers (WHGs), early European farmers (EEFs), and Caucasus hunter-gatherers (CHGs), depicted with red, orange, and blue color, respectively. The analysis was based on 433756 genomic sites of the 1240K list. The displayed plot here shows only individuals from the present geographical area of Greece (Neolithic Age to Roman times), as well as historical individuals (Iron Age to Roman times) from adjacent areas (Italy, Balkans, and Türkiye). The full plot is available in **Supplementary Figure S24B** (**Additional file 7, Section 3.5.8**).

We performed ancestry modeling on a per-group and per-individual basis, using ADMIXTOOLS2 [23] and rotating qpAdm using three distinct sets of putative source populations (“Ultimate”, “More_proximate”, and “Most_proximate”; see **Additional file 7, Section 3.5.9** for details), as well as one up to four possible sources. The “Ultimate” set of sources include a, *common to all tested targets*, set of individuals assembled into groups that may be considered a distal ancestor to them, whereas the same applies for the “Most_proximate” set of sources, albeit the groups are dated more recent, and may have a less distal ancestry to the targets. Similarly, the “Most_proximate” set of sources include groups of individuals that are, spatiotemporally, as close as possible to the target, and by definition, each target has *a unique, target-specific* set of individuals and assembled groups. The results indicate that when using the Ultimate set of sources, Neolithic Anatolia appear as the more frequent major source to the ancient Ammotopos, Amvrakia, and Tenea individuals (**Additional file 7, Supplementary Figure S38)**. Similarly, when using the "More_proximate" set of sources, all of our newly sequenced individuals have potential source populations closely related to prehistoric groups from the southern Balkans (**Additional file 7, Supplementary Figure S39)**, particularly those from the area of present-day Greece (see **Additional file 7, Section 3.5.9** for a more detailed presentation). Note, that we also performed a targeted qpAdm analysis using Archaic Amvrakia as target population and LBA Ammotopos and Archaic Tenea as the two potential source populations in order to exclusively test if Archaic Amvrakia can be inferred either as admixed by the other two, or a direct descendant of one of the other two, or none of the above (see **Additional file 7, Section 3.5.9** for details).

In order to determine the genetic relationships of the sampled populations, we utilized the “Most_proximate” set of sources for each target population and performed Outgroup f3 and Admixture f3 analysis. We separately considered the modern African Yoruba and the modern Eastern Asia Hun populations as outgroups to test and avoid potential artifacts associated with outgroup choice. No negative f3 values were detected in any of the pairwise Admixture f3 tests (**Additional file 7, Supplementary Figure S41)**, suggesting that we cannot statistically confirm the presence of admixture between any particular triplet of target and source pairs. However, this does not entirely rule out the possibility of admixture. This finding implies that some populations among the "Most_proximate" sources may still share genetic material even after diverging from their common ancestor.

### Genetic continuity of Archaic Amvrakia from local prehistoric Greek populations

Due to limitations in sample availability, we used the LBA Ammotopos individuals (1275-1125 BCE) as proxy for the local population inhabiting the Archaic Amvrakia area, before the foundation of Amvrakia. In PC space, we find that the two LBA Ammotopos individuals are projected close to other LBA Greek individuals (1700-1050 BCE) and in close proximity to earlier EBA to MLBA individuals from the South Balkans (Albania, North Macedonia, and Bulgaria) (**Figure 2A**). These observations are also supported by ADMIXTURE results (**Figure 2B**), where the two Ammotopos individuals share analogous proportions of the genetic components that are maximized in WHGs, CHGs, and EEFs (red, blue, and orange in **Figure 2B**, respectively) with LBA individuals from mainland Greece and the Peloponnese. Ammotopos inferred to be in rotating qpAdm using its “Most_proximate” set of sources: a two-way admixture between LBA Crete (1700-1250 BCE) and either MBA mainland Greece (2100-1600 BCE) or MBA Albania (1900-1700 BCE); in three-way and four-way admixture models, sources with lower contribution include BA Italy, LBA mainland Greece, and other BA Balkan areas (**Additional file 7, Supplementary Figure S40**Α). The LBA Ammotopos genomes show the closest genetic distance, as estimated by Outgroup f3 tests when using the “Most_proximate” to LBA Ammotopos set of sources, with earlier MBA and LBA individuals from Italy, Crete, and mainland Greece, but neither with earlier LBA individuals from the Peloponnese nor with MBA individuals from Albania (**Additional file 7, Supplementary Figure S40A-B**).

In the PCA, the two LBA Ammotopos individuals are only placed relatively close to one out of the three Archaic Amvrakia individuals (Amv_Epi_Arch_1). The ADMIXTURE results and the genetic components maximized in WHGs, CHGs, and EEFs (red, blue, and orange in **Figure 2B**, respectively), support the genetic similarity between Ammotopos and Archaic Amvrakia (at least for two out of the three Amvrakians). When using LBA Ammotopos as one of the sources, as well as additional sources from the “Most_proximate” to Archaic Amvrakia set of sources in qpAdm analyses, Archaic Amvrakia was either inferred to be a) a non-admixed descendant of LBA Ammotopos, or b) a two-way admixture between LBA Ammotopos, as major source, and either, LBA Cyclades (1175-1150 BCE), or BA Levant (1400-1100 BCE) as minor source. However, if the analyses are not constrained to using LBA Ammotopos as a source, other populations can be inferred to be either non-admixed sources (e.g. LBA Crete, LBA Cyclades, LBA Mainland, IA Peloponnese, and Archaic Tenea) or two- and three-way major admixture sources for the Archaic Amvrakia individuals (**Additional file 7, Supplementary Figure S40**Β).

On the other hand, no shared Identity By Descent (IBD) fragments (**Figure 3**) can be observed between them either. This appears plausible as they are separated by almost 550-750 years. Finally, we do not observe any common mtDNA haplogroups between these two groups.

**Figure 3.**
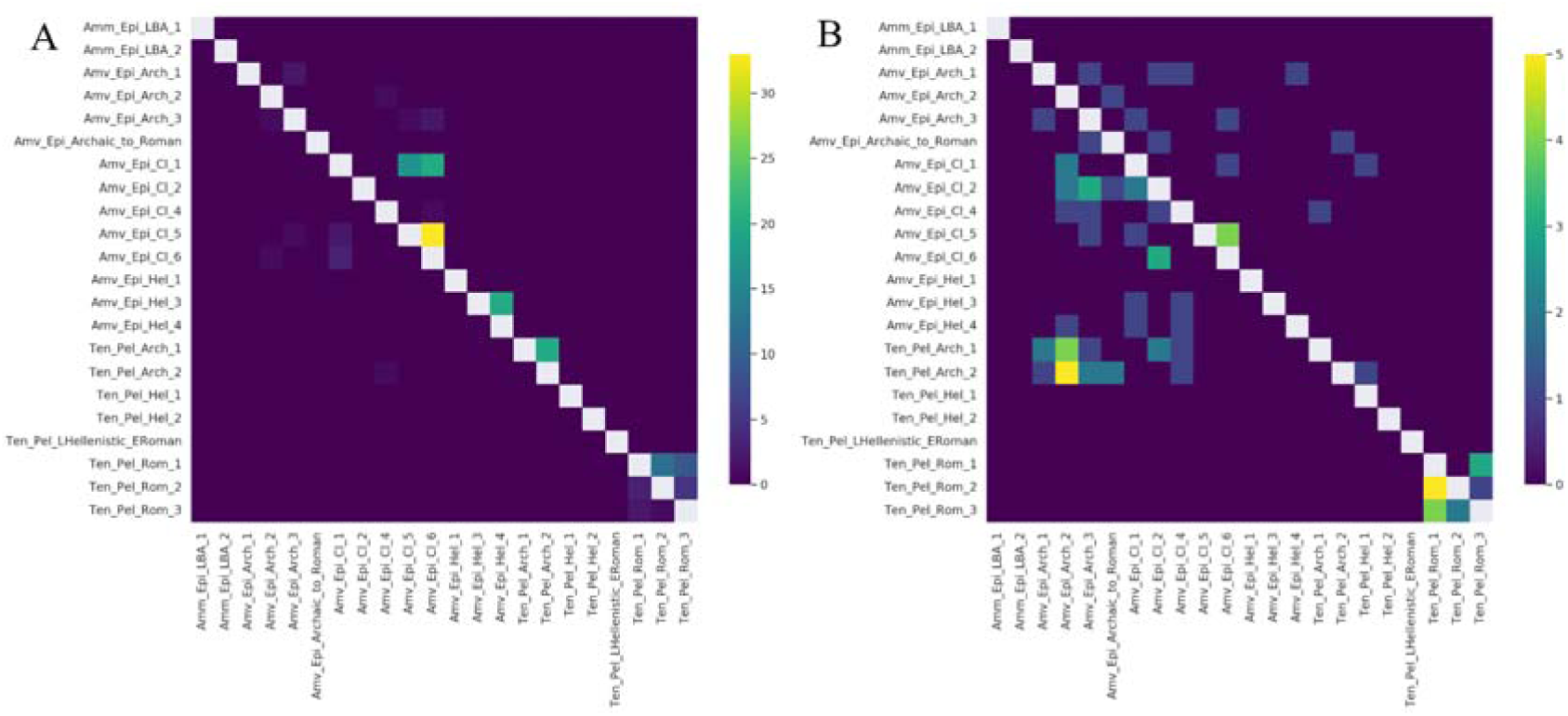
The number of shared IBDs for our newly sequenced WGS data with at least 0.25× coverage (imputed 1240K list of genomic sites) in a pairwise fashion within the >20 cM (**A**; upper right), 16-20 cM (**A**; lower left), 12-16 cM (**B**; upper right), and 8-12 (**B**; lower left) length bins.

Overall, based on the qpAdm and ADMIXTURE results, Archaic Amvrakia appears to have a genetic continuity from an LBA/IA source within the area of present-day Greece, including LBA Ammotopos or a similar genetic pool.

### Genetic affinities between Amvrakia and its metropolis during the Archaic times foundation

Our Archaic Amvrakia and Tenea individuals (550-480 BCE) can be considered as the closest representatives (subset) of the populations involved in the foundation of Amvrakia during the last half of the 7th century BCE. We found that the sampled 6th century BCE individuals from Amvrakia and Tenea, share strong genetic links of recent ancestry, albeit not distant genetic kinship, as demonstrated by the shared IBD segments (yellow, green, petrol, and blue colors in **Figure 3B; bottom-left plot**). These shared IBD segments with a length of 8-12 cM indicate a shared ancestry several generations ago, perhaps when the ancestors of these individuals were part of the same local population in the Corinthian territory. If we assume a human generation length of 27 years [24] and that the probability of an IBD segment of length *l* cM persisting for *t* generations rapidly decreases as exp(-*t* x *l* / 50) according to Ringbauer et al. [25] (50 denotes the distance in cM of two independent loci), then, a 8-12 cM range of segments persists, on average, for approximately 110-170 years. This coincides well with the historically documented Amvrakia foundation time, about 1-1.5 centuries prior to the estimated age of the Archaic individuals. The Archaic Amvrakia and Archaic Tenea populations display analogous proportions of the genetic components that are maximized in WHGs, CHGs, and EEFs based on the ADMIXTURE analysis (red, blue, and orange in **Figure 2B**, respectively). When using Archaic Tenea as one of the sources and additional sources from its “Most_proximate” set of sources in qpAdm analyses, Archaic Amvrakia is either inferred to be a non-admixed descendant population of Archaic Tenea, or a two-way admixture between Archaic Tenea as major source and EIA Croatia (1050-550 BCE) as minor source (**Additional file 7, Supplementary Figure S40**Β). As mentioned before, Archaic Amvrakia has also been inferred to be a non-admixed descendant of IA Peloponnese (the IA Peloponnese source population includes two individuals, one from Nestor Palace near Pylos and one from Tiryns near Tenea). Compared to other populations, Archaic Amvrakia and Archaic Tenea are shown to be highly related based on the genetic distances in Outgroup f3 analyses, when calculated either by using the “Most_proximate” to Archaic Amvrakia set of sources (**Additional file 7, Supplementary Figures S42C**-**D)** or by using the “Most_proximate” to Archaic Tenea set of sources (**Additional file 7, Supplementary Figures S42I**-**J)**. Finally, as an additional evidence of their common genetic origin, the two groups share a common mtDNA haplogroup (T1a4) represented by Amv_Epi_Arch_2 and Tenea_Pel_Arch_1.

On the other hand, in the PC space, the Archaic Amvrakia and Archaic Tenea are placed in distinct positions, mostly due to the position covered by two of the three Amvrakia Archaic individuals (**Figure 2A**). The two Archaic Tenea and one of the three Archaic Amvrakia individuals (Amv_Epi_Arch_1) are placed among the earlier LBA Greek individuals (1700-1050 BCE) and in close proximity to LBA Ammotopos as mentioned above. On the other hand, the remaining Archaic Amvrakia individuals are more differentiated towards the upper left of the PC space and the Levant Neolithic - BA-Yamnaya-like axis. One of them (Amv_Epi_Arch_3), which is the closest to the Greek LBA cluster, overlaps with an Etruscan individual from Italy (800-001 BCE) and is surrounded by Greek genomes mostly from the EBA_MBA (2900-1700 BCE) and the LBA (1700-1050 BCE). The other Archaic Amvrakia individual (Amv_Epi_Arch_2) is placed even further away from the Greek LBA cluster and in close proximity to EBA_MBA (2900-1700 BCE) Greece and a few Greek Neolithic (6400-3600 BCE) and Anatolian Neolithic (9000-5600 BCE) individuals.

According to the targeted qpAdm analysis using Archaic Amvrakia as target population and LBA Ammotopos and Archaic Tenea as the two potential source populations (**Figure 4**) we found that Archaic Amvrakia is inferred either as non-admixed descendant of one of the two sources, or as their admixture product, with parameterization affecting the inference. In the one case that Archaic Amvrakia was inferred as an admixed population, LBA Ammotopos was the major contributor (∼90%) and LBA Ammotopos the minor one (∼10%).

**Figure 4.**
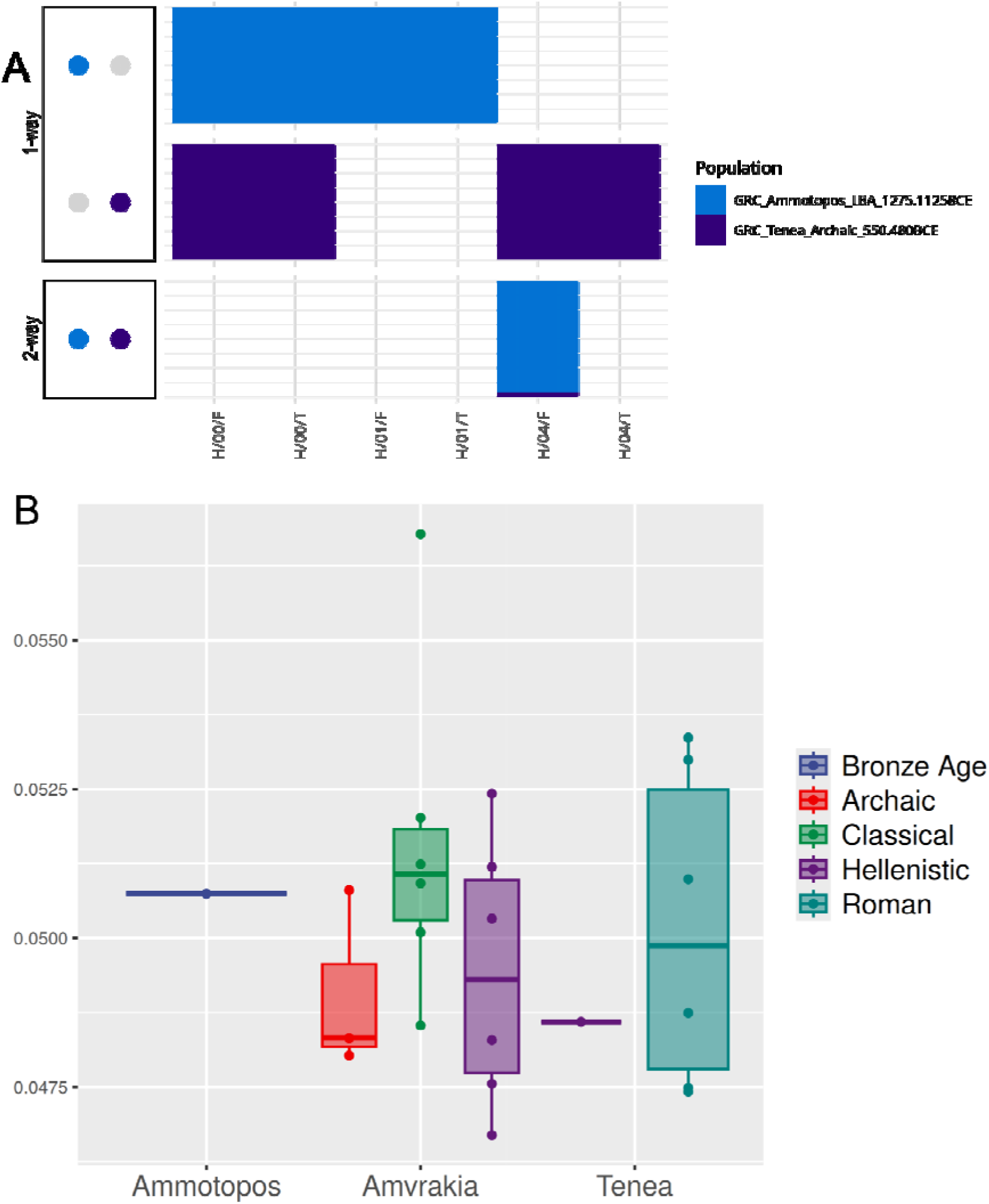
**A.** Scatterplot of targeted qpAdm analysis results at the population level with Archaic Amvrakia as target population and LBA Ammotopos and Archaic Tenea as source populations, using the parameterization schemes “*adjust_pseudohaploid: TRUE - maxmmiss: 0.00/0.10/0.40 - afprod: TRUE/FALSE*”, which correspond to the abbreviations at the horizontal axis. Only the feasible models with a p-value > 0.05 (accepted models) are shown with the points representing the populations involved and the color indicating their admixture proportions. **B.** Boxplot of intrapopulation similarity levels estimated by calculating pairwise Outgroup f3 values within each population and within a given period.

Overall, based on PCA, Outgroup f3, qpAdm, and more importantly, on the IBD analyses, Amvrakia and Tenea appear to have a common past, just prior to the foundation of Amvrakia. Interestingly, only one out of the three Amvrakia individuals appears (by visual inspection of the PCA plot) to have a genetic affinity with the Tenean individuals, despite the fact that two out of the three Archaic Amvrakia individuals share IBDs exceeding 20 cM. This indicates that they had a common ancestor several generations ago. The different localization of the other two individuals from Archaic Amvrakia on the PCA plot, is not due to PCA instability as all three individuals exhibit Pandora Stability support values between 0.80 and 0.86. Could, however, be a methodological artifact (batch effect), as the two aforementioned individuals are the only ones, for whom their DNA was not treated enzymatically to reduce *post-mortem* deamination damage in the sequences. On the other hand, it may suggest the existence of considerable genetic variation in the area during the Archaic times, which is also apparent from the variable Outgroup f3 genetic distances that each of the Archaic Amvrakia individuals have with other populations including the local preceding LBA Ammotopos (**Additional file 7, Supplementary Figures S42C**-**D)**.

### Evolution of the Amvrakia population during the Classical and Hellenistic times

The Classical Amvrakia individuals appear to form a homologous cluster in the PC space, with a comparatively low variability among the individuals, placed in between the LBA ammotopos and one of the Archaic Amvrakia individuals (**Figure 2A**). The high similarity among the Classical Amvrakia individuals is apparent also a) when comparing the proportions of the genetic components maximized in WHGs, CHGs, and EEFs in the ADMIXTURE analysis (red, blue, and orange in **Figure 2B**, respectively), b) by considering the shared IBD segments among the Classical Amvrakia individuals (**Figure 3A)**, and by the within-population, pairwise Outgroup f3 values **(Figure 4B)**. Further, the homozygosity analyses revealed the presence of “long ROH” (sROH_>20_ above 50□cM; **Figure 5**), as termed by [26] and indicating increased inbreeding levels, only during the Classical period in the Amvrakia population, although a degree of consanguinity practices is indicated by the presence of shared sROHs_12-20_ for the earlier Archaic Amvrakia population, too (**Figure 5**). The IBD analyses also yielded distant kin relationships (several generations apart) between Archaic and Classical Amvrakia individuals (Amv_Epi_Arch_3 with Amv_Epi_Cl_-5 and −6; Amv_Epi_Arch_2 with Amv_Epi_Cl_4) (**Figure 3A)**. Classical Amvrakia inferred to be in rotating qpAdm using its “Most_proximate” set of sources: either a non-admixed descendant of a single Greek LBA (Crete, Ammotopos, Sterea Ellada), IA (Peloponnese, Sterea Ellada), or Archaic (Amvrakia, Tenea) source or b) a two-, three-, or four-way admixture among, one of the above as a major source (Archaic Amvrakia being the most frequent) and one (or more) additional minor source(s) from the Eastern Mediterranean and Italy (**Additional file 7, Supplementary Figure S40C)**. Classical Amvrakia shows the closest Outgroup f3 genetic distances, when calculated by using the “Most_proximate” to Classical Amvrakia set of sources, with preceding Greek populations, including Archaic Amvrakia and LBA Ammotopos (**Additional file 7, Supplementary Figures S42E**-**F)**. The closer genetic proximity between Classical Amvrakia and Archaic Amvrakia, compared to Archaic Tenea, is also worth noting when examining Outgroup f3 genetic distances, an observation that indicates local genetic continuity. Overall, a genetic continuity from the Archaic times to the Classical times is suggested for Amvrakia.

**Figure 5.**
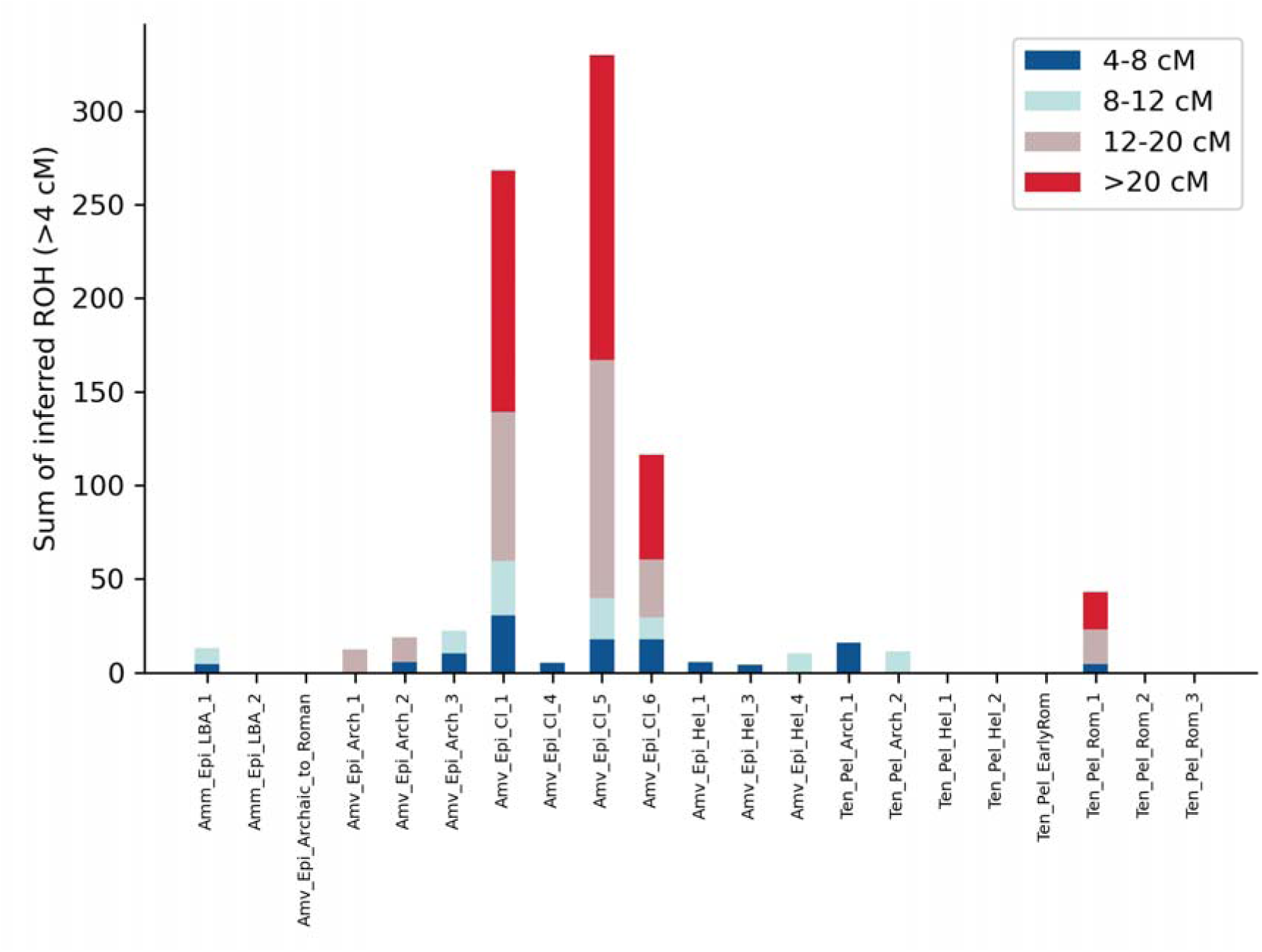
Sums of inferred runs of homozygosity (sROH) per individual in four length bins (4-8 cM, 8-12 cM, 12-20 cM, and above 20 cM). All individuals had more than 300000 genomic sites covered on the 1240K list.

On the other hand, and when considered as a group, the Hellenistic Amvrakia individuals cover a larger area of the PC space than earlier Classical ones (**Figure 2A**). This might be due to the low Pandora Stability support values (0.65 and 0.53, respectively) estimated for two out of the four Hellenistic Amvrakia individuals, albeit the Hellenistic Amrakia individual that is placed far away from the rest (Amv_Epi_Hel_3) has good stability support. More importantly, the Hellenistic Amvrakia individuals are also characterized by slightly different proportions of genetic components maximized in WHGs, CHGs, and EEFs in the unsupervised ADMIXTURE analysis (red, blue, and orange in **Figure 2B**, respectively), both, with respect to each other, as well as with respect to Classical Amvrakia individuals, and compared to Classical AMvrakia a decreased mean, within-population, pairwise Outgroup f3 value **(Figure 4B)**. This is also apparent in the Outgroup f3 genetic distances, when calculated by using the “Most_proximate” to Hellenistic Amvrakia set of sources. Here, each Hellenistic Amvrakia individual shows a notably different amount of similarity (the f3 value, using Yoruba as outgroup population, ranges from 0.04922±0.00111 to 0.05241±0.00184) to genetically proximal LBA and IA populations from Greece (**Additional file 7, Supplementary Figure S42G**-**H)**. Notably, according to the Outgroup f3 genetic distances, Hellenistic Amvrakia individuals retain strong genetic links with the preceding Classical Amvrakia, as also manifested by the shared IBDs between two pairs of individuals from these periods (**Figure 3B**). Note that IBD sharing is *not* observed between Hellenistic Amvrakia and Hellenistic Tenea. We also observe a high Outgroup f3 measured genetic affinity with the Classical population living in the geographical area of present-day North Macedonia, albeit their ADMIXTURE proportions are slightly different (**Figure 2B)**. Given the unavailability of an Ancient Macedonian genome from Hellenistic times, the latter observation may be considered as testimony of genetic affinity between Ancient Macedonian territory and Amvrakia, corresponding to the creation of connections during the occupation of Amvrakia by the Macedonians or its subsequent rise as a capital of the Epirus Kingdom [27]. Hellenistic Amvrakia inferred to be in rotating qpAdm using its “Most_proximate” set of sources: a non-admixed descendant of a single source (i.e., Archaic Amvrakia, Classical Amvrakia, LBA Ammotopos, Hellenistic Tenea, Archaic Tenea) or as a two-way admixture between the aforementioned populations; in three-way and four-way admixture models, several other sources with lower contribution appear, such as IA Italy, IA Balkans, IA Greece, Classical North Macedonia, etc. (**Additional file 7, Supplementary Figure S40D**).

Overall, during the subsequent Classical and Hellenistic periods the population of Amvrakia appears to have slightly differentiated in comparison to the Archaic period, yet evidence of genetic continuity over time is observed.

### Evolution of the Tenean population from Archaic to Roman times

As mentioned above, Archaic Tenea (or at least a closely related population) contributed genetically to the foundation of Amvrakia, and has a close genetic proximity to Archaic Amvrakia, as well as to other preceding LBA and IA Mainland Greek and Peloponnese populations (**Additional file 7, Supplementary Figures S42I**-**J)**. Archaic Tenea inferred to be in rotating qpAdm using its “Most_proximate” set of sources: a non-admixed descendant of multiple BA (Cyclades, Crete, Sterea Ellada, Ammotopos, Albania) and IA (Türkiye, Peloponnese, Sardinia, Bulgaria) single sources or a two-, three-, or four-way admixture among, one of the above as a major source (LBA Cyclades, LBA Crete, and IA Peloponnese being the most frequent major sources) and one (or more) additional minor source(s) from Eastern Mediterranean and Italy (**Additional file 7, Supplementary Figure S40E**).

Compared to Archaic Tenea, the subsequent Hellenistic Tenea has also its closest Outgroup f3 genetic distances, when calculated by using the “Most_proximate” sources to Hellensistic Tenea, with preceding Greek populations, including Archaic Tenea, indicating a degree of local continuity (**Additional file 7, Supplementary Figures S42M**-**N)**. In contrast to the Archaic Tenea individuals, Hellenistic Tenea individuals are not placed very close to each other in the PC space (**Figure 2A**), although in the unsupervised ADMIXTURE analysis, the Hellenistic Tenea individuals are characterized by homogeneous proportions of the genetic components maximized in WHGs, CHGs, and EEFs (red, blue, and orange in **Figure 2B**, respectively). In the same plot (**Figure 2B**), when comparing the Archaic and the Hellenistic Tenea, the latter shows a lower EEFs proportion, but a higher CHGs proportion. Finally, Hellenistic Tenea is inferred to be in rotating qpAdm using its “Most_proximate” set of sources: a direct descendant of either Archaic Amvrakia, or Classical Amvrakia, or LBA Ammotopos, or Hellenistic Amvrakia, or Archaic Tenea or as a two-way admixture between the aforementioned populations; in three-way and four-way admixture models, several other sources with lower contribution appear, such as IA Italy, IA Balkans, IA Greece, Classical North Macedonia, etc (**Additional file 7, Supplementary Figure S40F**).

One of Tenea individuals is dated in the transitional phase of the Late Hellenistic period to the Early Roman times, hence it was not grouped within either the Hellenistic or the Roman Tenea. In the PCA, this individual is placed close to Classical and Hellenistic Amvrakia individuals, as well as close to Archaic and Hellenistic Tenea individuals (**Figure 2A**). In the unsupervised ADMIXTURE analysis, this individual has similar proportions of the genetic components maximized in WHGs, CHGs, and EEFs (red, blue, and orange in **Figure 2B**, respectively) with the earlier dated Archaic, Classical, and Hellenistic individuals from Amvrakia and Tenea. It also shows the closest Outgroup f3 genetic distances, when calculated by using its “Most_proximate” set of sources, with preceding and/or contemporary to it Greek and south Balkan populations rather than NW Balkan, Anatolian, Italian, and Levant populations (**Additional file 7, Supplementary Figures S42K**-**L)**. It does not share large IBDs with any other newly sequenced individual, though, indicating that it is not sharing a relatively recent common ancestor with any of them.

We found that the Roman Tenea individuals form a cluster on the PCA plot that is distinct from most of the preceding Archaic, Classical, and Hellenistic Amvrakia and Tenea individuals. More specifically, they are projected among a) contemporary to them Roman individuals from Türkiye, Italy, and Greece, b) earlier prehistorical (EBA, MBA, and LBA) Greek and Anatolian individuals, and c) historical (Archaic and Hellenistic) Anatolian genomes (**Figure 2A**). Roman Tenea individuals are homogeneous regarding the proportions of the unsupervised ADMIXTURE genetic components, which are maximized in WHGs, EEFs, and CHGs (red, blue, and orange in **Figure 2B**, respectively). As shown in **Figure 2B**, these components appear in analogous proportions in other published Roman genomes from Greece and mainland Italy (Imperial Roman times). According to the Outgroup f3 analyses using the “Most_proximate” sources to Roman Tenea, earlier LBA and

IA Greek populations are among the most closely related to the Roman Tenea ones, together with the Late Hellenistic - Early Roman Tenea individual, in contrast to earlier Hellenistic populations from Greece and Türkiye or to preceding Tenea time periods (**Additional file 7, Supplementary Figures S42O**-**P)**. Roman Tenea is inferred to be in rotating qpAdm using its “Most_proximate” to Roman Tenea set of sources: a direct descendant of either Archaic Tenea, or Hellenistic Tenea, or as a two-way admixture between a major source (Archaic Tenea, Hellenistic Tenea, Classical Amvrakia, and Hellenistic Amvrakia) and a minor source with lower contribution (BA Levant, LBA and IA Greece, IA Balkans, Hellenistic Türkiye); in three-way and four-way admixture models, several other sources with lower contribution appear, such as IA Italy, Classical North Macedonia, etc (**Additional file 7, Supplementary Figure S40G**).

According to the shared IBDs analyses, that we conducted on three out of the four Roman Tenea individuals (Ten_Pel_Rom_-1, −2, and −3; **Figure 3**) due to coverage issues, they exhibit an intermediate genetic kinship relationship that is a few generations apart, albeit exceeding the 3rd degree as they were not inferred as being closely related (**Additional file 7, Section 3.5.3)**. In addition, these three individuals have the same matrilineality, as they belong to the same mtDNA haplogroup (N1a1a+152 or N1a1a2 depending on the software used) in contrast to the fourth individual, whereas two of these aforementioned three male individuals have distinct Y-chromosome haplogroups (**Additional file 3)**. Given that these three individuals were children with intermediate genetic relatedness, same matrilineality, and different patrilineality, we can assume that they were members of an extensive family that also displayed a degree of matrilocality.

### Phenotypic and microbial metagenomic insights on ancient Greek historical populations

We estimated eye color, skin color, and hair shade for all individuals. Hair color was estimated for 19 out of 26 individuals due to coverage (**Additional file 5**). All newly sequenced individuals from a) LBA Ammotopos, b) Archaic, Amvrakia, and Hellenistic Amvrakia, and c) Archaic, Hellenistic and Roman Tenea showed the highest probability for brown eyes (average p-value: 0.95, 95% CI: 0.92-0.98). Most individuals likely had an intermediate skin tone (average p-value: 0.64, 95% CI: 0.58-0.70), whereas three of them had a darker skin color (two from Classical Amvrakia and one from Roman Tenea). The individuals from Classical Amvrakia showed the highest probability for dark skin (∼0.93) and dark to black (note that our newly sequenced individuals are not of African genetic origin as shown above) skin color (∼0.50) categories, respectively. The individual from Roman Tenea had a notable probability for dark skin (∼0.58). Similarly, most individuals likely had brown hair (average p-value: 0.62, 95% CI: 0.59-0.65) with a dark shade (average p-value: 0.80, 95% CI: 0.74-0.86). Notably, the Late Hellenistic - Early Roman individual from Tenea still exhibited a high probability for red hair (∼0.64). Among the ancient individuals analyzed so far, only four have shown a high probability of having red hair. Three of these cases were located in central and northern Europe, specifically in present-day Hungary and Latvia, dating between 1500 and 4000 years BP [28]. The sole case in southern Europe, dating to approximately 2500 years BP, was discovered in the Bezdanjača cave in present-day Croatia [28]. Overall, these results are in line with previously published archaeogenomics results, stating that in antiquity, darker skin tones (most often associated with the “Intermediate” skin category in the HIrisPlex-S model) were typically found in southern Europe, whereas lighter skin tones were more prevalent in northern Europe [28].

Regarding lactase persistence, none of the individuals (for whom the specific SNP was covered, 14/26; **Additional file 5**) were found to be lactose tolerant, even the more recent ones, suggesting a different evolutionary history for this phenotype in the Greek region compared to Central Europe. In previous studies on Greece, lactose tolerance-associated alleles were found to be virtually absent in the Aegean’s first Neolithic farmers [29–31] and remained so until the Bronze Age [30]. Therefore, our results further indicate the absence or at least the low frequency of the lactase persistence phenotype in ancient Greece. In Central Europe however, lactose tolerance was detected at low, but noticeable frequencies (∼7%) as early as ∼5000 BCE, with strong selection pressures continuing over the last 3000 years from around 1500 BCE onwards [32].

Regarding sensitivity to fats (15/26 individuals covered the specific SNP), as well as the muscle contraction type (15/26 individuals covered the specific SNP) and muscle performance (11/26 individuals covered the specific SNP), seven individuals showed a moderately increased sensitivity to fats, eight individuals had an ACTN3 genotype associated with improved muscle performance typically seen in sprinters, whereas seven individuals likely had impaired muscle performance. Lastly, five individuals had higher muscle strength associated with the ACVR1B gene (**Additional file 5**). The ACTN3 gene has been previously studied due to its potential influence by the cultural transition from hunter-gatherer to farming societies [33]. Although some results suggest positive selection acting on this gene (particularly in Europe), the evidence remains inconclusive [34].

Regarding beta thalassemia (and malaria resistance), the most common genetic disorder in modern Greece [35,36], none of the newly sequenced Greek individuals were found to carry associated alleles on the 29 SNPs examined (**Additional file 5**; only one individual had zero SNPs coverage). Despite its high modern day frequency, previous aDNA studies investigating beta thalassemia variants in Greece have not identified any alleles associated with the disorder either [29,37].

No microbial DNA belonging to ancient systemic pathogens was detected in any of the samples examined (**Additional file 6**). Given that most of our bone samples were petrous bones, this is not surprising as this bone material is not well-suited for this purpose [e.g. 38], despite its utility in human archaeogenomics analyses due to its high human endogenous DNA content. In dental samples, however, we did observe DNA traces belonging to ancient human oral bacteria, including those associated with severe forms of periodontal disease, such as *Porphyromonas gingivalis* and *Tannerella forsythia*.

## Discussion

### Insights in the genetic profile of historical Greece

Our study is among the first to produce whole-genome sequencing (WGS) archaeogenomic data -and one of the earliest when considering genome-wide SNP-targeted approaches-from individuals dating to the historical periods of modern-day Greece, specifically from the Archaic to Roman eras. By assembling genomes from these previously unsampled periods, we can compare them with earlier prehistoric genetic data from the Eastern Mediterranean, gain insights into their phenotypic traits, and identify traces of ancient DNA from associated commensal or pathogenic microbes.

The consensus of our findings (PCA, ADMIXTURE, qpAdm using the “Most_Proximate” set of sources) shows that the Amvrakia and Tenea individuals can be considered as descendants of the LBA and IA populations of the southern Balkans, especially the area of present-day Greece. In addition, the Roman Tenea individuals appear to have an additional minor contribution from the east, represented by BA Levant and Hellenistic Türkiye (Northwest and West, including Halikarnassos) in the “Most_proximate” qpAdm analysis. Overall, a local genetic continuity is suggested from the LBA/IA Greece to the Archaic, Classical, and Hellenistic Greece, as well as in a lower degree to Roman Greece, albeit for the latter the spatial sampling is inadequate to justify such a generalization.

Regarding their external phenotypes, our newly sequenced individuals most likely had brown eyes, an intermediate or a darker skin tone, and brown hair with a dark shade, similar to the preceding LBA and IA individuals from Greece [18,19,39,40]. Regarding the other phenotypes examined, they display an intermediate frequency in fats sensitivity and muscle-related traits, as well as no occurrence of lactase persistence or Beta thalassemia. Finally, no systematic pathogen DNA was found in any of the individuals, albeit we detected taxa associated with severe forms of periodontal disease.

### Manifestation of the genetic links between Amvrakia and its metropolis and the contribution of the locals

The ancient Greeks employed the term “apoikia” (Αποικία), best translated as “home away from home”, to highlight the separation and/or connection between the metropolis and the new “oikos” (see **Additional file 7, Section 1.1** for definitions of oikos, apoikia, metropolis, polis, kome) [41]. According to the founding myth people from the metropolis were participating in the foundation of an apoikia; a process that could have had occurred in the form of either a single or multiple migration (translocation) events, at various points in time and with a plethora of socioeconomic (e.g. population increase, lack of farmland, better life opportunities due to competition) and political (e.g. political exiles) drivers and characteristics (e.g. organized military expedition, male-oriented) [42,43]. For instance, in the case of Amvrakia, the Corinthians had already established an “Emporion” (trading post) prior to founding Amvrakia [11]. Emporia, however, were not characterized by the permanent translocation of a group(s) of people in contrast to an apoikia that implies permanent settlement to the new city [44,45]. Admittedly, in archaeology there is a clear cultural connection between the apoikia of Amvrakia and the metropolis of Corinth [11]. In Greek literature, there are references to the relocation of groups of people, such as the oracle tablets from Dodona asking the god for guidance on whether it was a wise decision to move to Ambrakia [46]. However, there is no direct biological evidence, though not disputed either, that people migrated *en masse* and to what extent this had an impact on the local population (to the point it had developed up to that point).

In the present study, our primary research goal was to identify genetic affinities and study the consequences of Corinthian colonization in the demography of the population of Amvrakia. Overall, Amvrakia and Tenea seem to share a common history just before the establishment of Amvrakia, based on analyses such as PCA, Outgroup f3, qpAdm, and especially IBD (see main figures). Two of the three Archaic Amvrakia individuals share IBD segments with the two Archaic Tenea individuals exceeding 20 cM, indicating a common ancestor several generations back. These observations constitute direct genetic evidence of the affinity that the Amvrakia has with its metropolis during its foundation period of Amvrakia and its early years, supporting the scenario that Corinth contributed, both culturally and genetically, during the foundation of Amvrakia.

One may consider that Archaic Tenea constitute a suboptimal population to be used as representative of the Corinthian genetic pool, given the lack of genomes from the polis of Corinth *per se*. Tenea was a “kome” (politically may be a subdivision of a larger polis) of Corinth, belongs to the region of Corinthia, and has participated in the Corinthian colonization efforts (records survive only for Syracuse). Therefore, while there is no ancient text or other source or inscriptions indicating that the people of Tenea were actively involved in the colonization process of Amvrakia, we can not exclude the presence of individuals from Tenea in the foundation of Amvrakia either. Indeed, the elevated shared IBDs (**Figure 3**) between Amvrakia and Tenea constitute evidence that closely related individuals to the Tenea ones, may have been members of the group(s) that translocated to Amvrakia.

One of the main challenges in our study is to determine the genetic influence and contribution of the local and neighboring people of the pre-colonial area of Amvrakia, in the initial formation of Amvrakia and its population composition. The close genetic proximity of Archaic Tenea and LBA Ammotopos individuals could indicate complex genetic relationships between these regions before the Archaic period. Whereas Greek colonies were the meeting grounds of culturally and genetically diverse people [41], as for example has been found for Sicilian Himera [17], in the case of Amvrakia the indigenous people appear to have similar ancestry with the newcomers, as indicated by the genetic profiles (see main figures with ADMIXTURE, Outgroup f3, and qpAdm results) of the LBA Ammotopos population and Archaic Tenea, respectively. The potential recent common ancestry, coupled with the increased gene-flow that someone may expect in such a small spatial scale, and the common cultural (e.g. worship) and linguistic elements, hinder the identification of their differentiation. Even so, we observed signals of admixture between the two aforementioned populations in targeted qpAdm analyses (**Figure 4A**), with the local component being the major contributor and the newcomers the minor one.

### The evolution of Amvrakia and Tenea in the post-Archaic times

As corroborated by the aforementioned genetic analysis, at least two different genetic pools contributed to the founding process in Amvrakia. A local one, that has direct ancestry from LBA/IA Greece, including the LBA Ammotopos population, and another one that migrated from the Corinthian territory and is represented by the Archaic Tenea population. Consequently, Archaic Amvrakia exhibits a fair amount of genetic variation as suggested by our genetic analyses results (see main figures of ADMIXTURE, Outgroup f3, and perhaps PCA too). One important question is how much different genetically, if any, was Amvrakia during the subsequent Classical and Hellenistic periods, and if the genetic links between the metropolis and the colony remained the same or not. We found that during the Classical and Hellenistic periods, the population of Amvrakia seems to be only slightly differentiated genetically (during Hellenistic times) when compared to the Archaic period (according to qpAdm, IBD, Outgroup f3, ADMIXTURE), providing evidence of genetic continuity over time. Importantly, no shared IBDs are observed during Hellenistic times between the two areas, in contrast to the Archaic and Classical periods.

Evidently, in the Greek literature there are records of the dependency and close relationship between Amvrakia and Corinth during Classical times, at least in the political sense [43]. For example, Amvrakia, together with nearby sister-colonies, became an ally of Sparta during the Peloponnesian War (431-404 BCE) according to Thucydides [47,48]. Amvrakia also joined Corinth in the anti-Spartan alliance that was formed after the Peloponnesian War and before the Corinthian War in 394 BCE [49]. On the other hand, during the Hellenistic times, there is archeological evidence that Corinth lost its influence into Amvrakia, following a general trend for the Corinthian colonies in Western Greece [50]. Importantly, at the end of the Classical times, Amvrakia came under the rule of Macedonians under Philip II (338 BCE), although it maintained a semi-autonomous status [27]. During Hellenistic times, the Macedonians ceded it to the Kingdom of Epirus. King Pyrrhus, the king of Mollosians, established it as the capital of the Epirus Kingdom in 294 BCE and Amvrakia experienced notable prosperity during the century to follow [51].

Hence, we may conclude that Amvrakia retain its genetic ancestry from the Archaic times during the subsequent Classical times, a period characterized, however, by lower diversity and population size in Amvrakia (as manifested also by the increased consanguinity practices; see ROH results; **Figure 5**). Nevertheless, during Hellenistic times, although genetic continuity from Classical times is observed, a higher diversity characterizes Amvrakia, which during that time has become totally independent politically from Corinth, a fact that is also translated into the genetic level. We should note, though, that the lack of shared IBDs between Amvrakia and Hellenistic Tenea could additionally or alternatively mean a genetic change in the “pool” of Tenea. Indeed, Tenea shows also a differentiation in ADMIXTURE between Archaic and Hellenistic times, which is moderated if we include the Late Hellenistic to Early Roman times individual in the comparison.

In the case of Tenea, the most striking differentiation (see main figures with IBD, ADMIXTURE, and PCA results) happens during Roman times, during which, despite that genetic continuity is observed from the Hellenistic times, a gene flow is also inferred to had been occurred from an east source (see qpAdm and Outgroup f3 results).

## Conclusions

Local genetic continuity is suggested from the LBA/IA Greece to the subsequent historical periods of Greece. Amvrakia and Tenea display strong genetic affinities indicating a recent genetic past, even before the foundation of Amvrakia by the Corinthians. The latter procedure is characterized by the major genetic contribution of the metropolis and the minor influence of the local population in the genetic pool of the newly founded apoikia of Amvrakia.

## Methods

### Ancient DNA Laboratory Work

All analytical procedures, that is sample processing, DNA extraction, and genomic library preparation were performed in the cleanroom facilities of the Ancient DNA Lab at the Institute of Molecular Biology and Biotechnology at the Foundation for Research and Technology - Hellas (IMBB-FORTH). Laboratory work details are provided in Supplementary Information (**Additional file 7, Section 3.1**). In brief, DNA extraction was performed using optimized protocols for ancient DNA [52–54]. Double-stranded, blunt-end libraries without pretreatment were constructed according to published protocols [55,56] with some minor modifications (see supplement for details), and pre-screening shotgun whole genome sequencing was performed on an Illumina NextSeq500 platform. Deeper sequencing on the 26 selected samples/libraries was performed on freshly-produced, double-indexed libraries, that were partially pre-treated [57] -except for two samples- with the USER^TM^ enzyme (New England BioLabs Inc., USA), in an Illumina Novaseq6000 platform (2×150 bp).

### Sequence data processing and initial analyses

Data processing was performed using the ancient DNA analysis pipeline mapache v.0.3.0 [58] at three levels: (a) at the FastQ level corresponding to the sequence reads in each FastQ file, (b) at the library level, corresponding to multiple BAM files from the same (PCR amplified) library, and (c) at the level of individuals, corresponding to multiple BAM files from the same individual. Data processing details are provided in Supplementary Information (**Additional file 7, Sections 3.2**). Analyses at the FastQ level included quality control, residual adapter trimming and filtering, pair-end read merging, mapping to a human reference genome (hs37d5), read group incorporation, sorting and quality filtering of mapped reads, and statistical reports. Analyses at the library level included BAM file merging (of the same indexed library) and de-duplication, post-mortem DNA damage profiling, soft-clipping of damaged bases, and statistical reports. Finally, analyses at the individual level included BAM file merging (of the same individual), local indel realignment, final BAM file generation, statistics reporting, and genetic sex inference.

Contamination was assessed by using three distinct contamination estimation approaches (see details in **Additional file 7, Section 3.3)**; contamMix v.1.0-10 [59], schmutzi v.1.5.6 [60], and the *contamination* function of ANGSD v.0.941-6-g67b6b3b [61]. The classification to mitochondrial haplogroups was performed with HaploGrep3 v.3.3.2.1 [62] and HaploCart v.1.0 [63] and to Y-chromosome haplogroups with Yleaf v.3.1 [64] and Yhaplo v.1.1.2 [65] (see details in **Additional file 7, Section 3.4)**. The genetic relatedness analyses included Relationship Estimation from Ancient DNA (READ) [66] and KIN v.3.1.3 [67] (see details in **Additional file 7, Section 3.5.3)**.

### Population genomics analyses

The 26 newly whole-genome-sequenced ancient individuals were studied at the population level, both by assembling them into a distinct stand-alone dataset, but also in the context of previously published modern and ancient data as different analyses required the assembly of appropriate, distinct datasets (**Additional file 7, Section 3.5.6; Additional file 4**).

For the analyses as a distinct stand-alone dataset (see details in **Additional file 7, Sections 3.5.4-5**), we first performed Runs Of Homozygosity (ROHs) analysis on 1240K pseudo-haploidised data using the hapROH v.64 [26] software and the 1000 Genomes Project as a reference panel (see details in **Additional file 7, Section 3.5.4**). Then, we imputed the 26 newly produced ancient genomes of this study by using the 1000 Genomes phase 3 [68] dataset as a reference and GLIMPSE v.1.1.1 [69], as well as the ATLAS pipeline v.0.9 [70] and HapMap phase II NCBI b37 genetic map [71] (see details in **Additional file 7, Section 3.5.5**). Finally, we performed Identity-by-Descent (IBD) segments screening using the ancIBD v0.5 tool [25] (see details in **Additional file 7, Section 3.5.5**).

For the analyses in the context of previously published modern and ancient data (see details in **Additional file 7, Sections 3.5.7-10**), three datasets were assembled and merged with our data: “Dataset 1” that includes 663 ancient individuals with collected data over a list of ∼1.24 million genomic sites, known as 1240K [21], “Dataset 2” that included all the individuals of “Dataset 1” except of six Upper Paleolithic Iberomaurusian hunter-gatherers from Tafolrat, Morocco, as two-thirds of their ancestry originates from sub-Saharan Africa [72], and “Dataset 3” that includes 888 modern West Eurasian individuals genotyped on the Human Origins SNP (HO; 597573 sites) array [21] and the data from the ancient individuals of “Dataset 1”, albeit restricted to the HO sites.

PCA projection of ancient data onto modern ones (newly generated genomes and “Dataset 3”) was performed with the smartpca function of EIGENSOFT v.7.2.1 [73], using default parameters and the lsqproject: YES option (see details in **Additional file 7, Section 3.5.7**). The stability of the PCA was assessed using Pandora v.2.0.0 [74]. Clustering of ancient genomes was performed using unsupervised ADMIXTURE v.1.3.0 [22] with K=2, 3,…,10 after filtering the dataset (newly generated genomes and “Dataset 2”) for linkage disequilibrium (LD) and allele missingness (see details in **Additional file 7, Section 3.5.8**).

For the F-statistics analyses, we grouped the ancient individuals (newly sequenced here and “Dataset 1”) according to geographical, chronological, and cultural contexts, and performed the analysis on a per-group (closely related individuals were omitted) and per-individual basis (see details in **Additional file 7, Section 3.5.9**). Ancestry modeling was performed with qpAdm in a rotating fashion (when a given population is not included in the test as one of the putative source populations, then it used as a reference population) as recommended by [75], using ADMIXTOOLS2 v.2.0.0 [23] and three different sets of potential sources: a) distant genetic sources (“Ultimate”) that characterize the general ancestry of ancient Western Europeans following Lazaridis et al. [18], b) spatiotemporally, more proximate sources (“More-proximate”), and c) spatiotemporally, as close as possible sources, by limiting the temporal range only to individuals dated earlier than each target, and the spatial range only to the Eastern Mediterranean and adjacent areas (“Most-proximate”). Additionally, we also performed a targeted qpAdm analysis using Archaic Amvrakia as target population and LBA Ammotopos and Archaic Tenea as the two potential source populations. Admixture f3 and Outgroup f3 (using, separately, the modern African Yoruba and the modern Eastern Asia Han populations as outgroups) analyses were performed with ADMIXTOOLS2. Within-population genetic similarity levels were estimated by calculating the pair-wise Outgroup f3 values within each population and within a given period.

### Additional genetic analyses: phenotype estimation and microbial metagenomics

Phenotypic estimation was performed for hair, eye, and skin color prediction using HirisPlex-S [76], as well as for specific metabolic traits (lactase persistence and sensitivity to fats), human muscle characteristics (muscle contraction type and muscle performance), and the genetic disorder of beta thalassemia (resulting to malaria resistance). Details can be found in **Additional file 7, Section 3.6**.

Ancient microbial DNA screening analyses were performed with the v.1.0.0 aMeta [77] metagenomic pipeline (see details in **Additional file 7, Section 3.7**), by using the aMeta-provided, pre-built, full NCBI non-redundant *nucleotide* (NT) database containing records of microbial, vertebrate, non-vertebrate, and plant organisms.

### Strontium oxygen isotopic analyses

Strontium isotope ratios were measured from tooth enamel samples from 14 human burials of Amvrakia from Archaic (n=8) and Classical (n=6) chronological contexts, whereas three of them were also analyzed for corresponding ^87^Sr/^86^Sr signatures in tooth dentine samples (**Additional file 7, Additional Table A2)**. The analytical protocol for ^87^Sr/^86^Sr analysis has been detailed in earlier publications [e.g. 78] and it is also provided in the Supplementary Information (**Additional file 7, Section 4.3.2**).

## Abbreviations

BCE: Before Common Era
BA: Bronze Age
BP: Before Present
CE: Common Era
CHGs: Caucasian Hunter Gatherers
EEFs: Early European Farmers
IA: Iron Age
IBD: Identity By Descent
LBA: Late Bronze Age
LD: Linkage Disequilibrium
NGS: Next Generation Sequencing
PCA: Principal Component Analysis
PC(s): Principal Component(s)
PCR: Polymerase Chain Reaction
ROH: Runs of Homozygosity
SNP: Single Nucleotide Polymorphism
WGS: Whole Genome Sequencing
WHGs: Western Hunter Gatherers

## Declarations

### Ethics approval and consent to participate

Permissions for sampling and analysis of the biological excavation findings of the present study have been acquired by the Directorate of Conservation of Ancient and Modern Monuments, Ministry of Culture, Greece, (protocol numbers ΥΠΠΟΑ/ΓΔΑΠΚ/ΔΣΑΝΜ/ΤΕΕ/Φ77/281858/183008/2818/254, ΥΠΠΟΑ/ΓΔΑΠΚ/ΔΣΑΝΜ/ΤΕΕ/Φ77/250940/175997/2927/179, ΥΠΠΟΑ/Φ77/225862, ΥΠΠΟΑ/Φ77/342765, ΥΠΠΟΑ/Φ77/225402, ΥΠΠΟΑ/80102/22-3-2022). This study has been approved by the Research Ethics committee of the Foundation for Research and Technology Hellas (FORTH).

### Consent for publication

Not applicable.

### Availability of data and materials

The mapache configuration (for runs 1 and 2) and samplelist files that were used in the present study, the SNPs lists (1240K, 5M), the assembled datasets in EIGENSTRAT format, as well as various software outputs are accessible in https://doi.org/10.5281/zenodo.10848927. Scripts used are available in https://github.com/Himsself1/APOIKIA_Analysis. Additionally, the genetic data in raw FastQ formatted files are accessible in NCBI Short Read Archive (SRA) under the BioProject accession PRJNA1143893.

### Funding

This study was supported by the project “APOIKIA” implemented within the framework of the special actions "AQUACULTURE" - "INDUSTRIAL MATERIALS" - "OPEN INNOVATION IN CULTURE" and co-financed by the European Regional Development Fund (ERDF) of the European Union and Greek national funds through the Operational Program "Competitiveness, Entrepreneurship & Innovation (EPAnEK) of the NSRF 2014-2020 under the supervision of the General Secretariat for Research and Innovation (GSRI) of Greece (Project code: Τ6ΥΒΠ-00191).

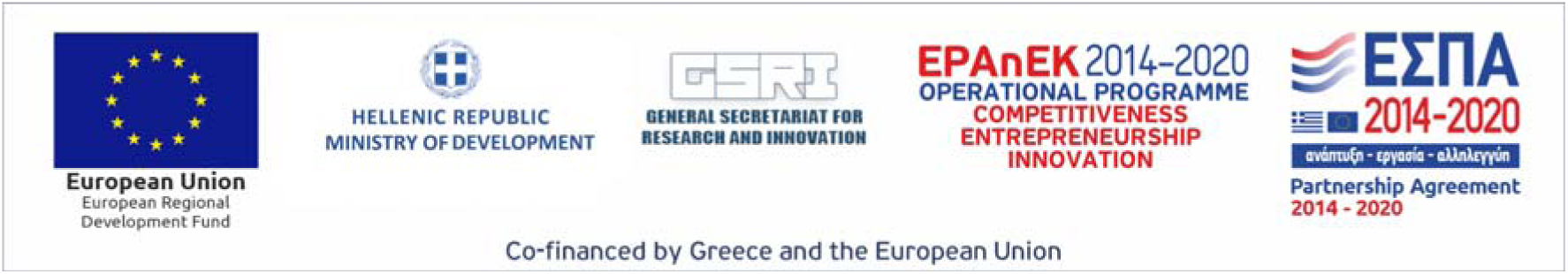

Alexis Stamatakis was funded by the European Union (EU) under Grant Agreement No 101087081 (CompBiodiv-GR).

### Author contribution

**Nikolaos Psonis**: Conceptualization, Methodology, Formal analysis, Investigation, Data Curation, Writing - Original Draft, Writing - Review & Editing, Visualization, Project administration **Eugenia Tabakaki**: Conceptualization, Writing - Original Draft, Writing - Review & Editing, Project administration **Despoina Vassou**: Investigation, Validation, Writing - Original Draft, Writing - Review & Editing, Project administration **Stefanos Papadadonakis**: Formal analysis, Writing - Review & Editing, Software, Visualization **Angelos Souleles**: Formal analysis, Writing - Original Draft, Writing - Review & Editing, Visualization **Argyro Nafplioti**: Formal analysis, Investigation, Writing - Original Draft, Writing - Review & Editing **Georgios Kousis Tsampazis**: Writing - Review & Editing, Visualization **Angeliki Papadopoulou**: Writing - Review & Editing, Visualization **Kiriakos Xanthopoulos**: Writing - Review & Editing **Panagiotis Panailidis**: Writing - Review & Editing **Angeliki Georgiadou**: Writing - Review & Editing **Dimitra Papakosta**: Writing - Review & Editing **Sevasti Koursioti**: Writing - Review & Editing, Investigation **Maria Evangelinou**: Software **Varvara Papadopoulou**: Resources, Writing - Review & Editing **Paraskevi Evageloglou**: Resources, Writing - Review & Editing **Elena Korka**: Resources, Writing - Review & Editing **Ioannis Christidis**: Writing - Review & Editing **Michael Ioannou**: Writing - Review & Editing **Theodora Kontogianni**: Writing - Review & Editing **Athanasios Arkoumanis**: Writing - Review & Editing **Alexandros Stamatakis**: Resources, Writing - Review & Editing **Nikos Poulakakis**: Writing - Review & Editing, Supervision **Christina Papageorgopoulou**: Writing - Original Draft, Writing - Review & Editing, Supervision **Pavlos Pavlidis**: Formal analysis, Writing - Review & Editing, Software, Supervision.

## Acknowledgements

We thank the Genomics Facility of IMBB-FORTH for the pre-screening library sequencing.

## Supplementary Information Captions

**Additional file 1 (.xlsx). Supplementary Table S1.** Information about the individuals and the samples used in this study, including wetlab details.

**Additional file 2 (.xlsx). Supplementary Table S2.** Sequencing and mapping statistics at the fastQ, library, and individual levels.

**Additional file 3 (.xlsx). Supplementary Table S3.** Post mortem DNA damage profiling (library level), contamination estimates (individual level), genetic sex estimation, and haplogroup assignments (mtDNA and Y-chromosome)

**Additional file 4 (.xlsx). Supplementary Table S4.** Sample assignment per analysis and associated population labels.

**Additional file 5 (.xlsx). Supplementary Table S5.** Phenotype reconstruction results.

**Additional file 6 (.xlsx). Supplementary Table S6.** Metagenomics analysis (aMeta) heatmap scores.

**Additional file 7 (.pdf) Supplementary Information.** Detailed information on archaeological background, samples, methodology, and results including supplementary figures.

## References

1. Graham AJ. Colony and Mother City in Ancient Greece. Manchester University Press; 1999. Available from: https://books.google.com/books/about/Colony_and_Mother_City_in_Ancient_Greece.html?hl=&id=z6XnAAAAIAAJ

2. Ridgway D. The First Western Greeks. CUP Archive; 1992. Available from: https://books.google.com/books/about/The_First_Western_Greeks.html?hl=&id=9F44AAAAIAAJ

3. Grammenos DV, Petropoulos EK. Ancient Greek Colonies in the Black Sea. Thessaloniki: Archaeological Institute of Northern Greece; 2003. Available from: https://books.google.com/books/about/Ancient_Greek_Colonies_in_the_Black_Sea.html?hl=&id=UtdoAAAAMAAJ

4. Tsetskhladze GR. Greek Colonisation: An Account of Greek Colonies and Other Settlements Overseas. Leiden, Boston, and Köln: Brill; 2008. Available from: https://books.google.com/books/about/Greek_Colonisation.html?hl=&id=PIgTAQAAIAAJ

5. Petropoulos EK. Problems in the history and archaeology of the Greek colonization of the Black Sea. In: Grammenos DV, Petropoulos EK, editors. Ancient Greek Colonies in the Black Sea. 2003. p. 17–92. Available from: https://www.academia.edu/32112108/ANCIENT_GREEK_COLONIES_IN_THE_BLACK_SEA_2_Grammenos_D_V_and_E_K_Petropoulos_eds_British_Archaeological_Reports_International_Series_1679_Oxford_2007

6. Hammond N. The classical age of Greece. Harper & Row Publishers, Inc. USA; 1975. Available from: https://cir.nii.ac.jp/crid/1130000797935601664

7. Reitsema LJ, Kyle B, Vassallo S. Food traditions and colonial interactions in the ancient Mediterranean: Stable isotope evidence from the Greek Sicilian colony Himera. Journal of Anthropological Archaeology. 2020;57:101144. Available from: https://www.sciencedirect.com/science/article/pii/S0278416519301734

8. Kaponis A. (In Greek) The Corinthian colonies around the Amvrakikoσ gulf from their foundation to the time of Philip II. 2020; Available from: https://pergamos.lib.uoa.gr/uoa/dl/object/2897438

9. Keenleyside A. Dental pathology and diet at Apollonia, a Greek colony on the Black Sea. International Journal of Osteoarchaeology. 2008;18:262–79. Available from: https://onlinelibrary.wiley.com/doi/abs/10.1002/oa.934

10. Papadopoulou V. RES GESTAE. The work of the Ephorate of Antiquities of Arta during the years 2014 – 2020, Arta 2020 / RES GESTAE. 2020; Available from: https://www.academia.edu/51173824/RES_GESTAE_The_work_of_the_Ephorate_of_Antiquities_of_Arta_during_the_years_2014_2020_Arta_2020_RES_GESTAE_%CE%A4%CE%B1_%CE%A0%CE%B5%CF%80%CF%81%CE%B1%CE%B3%CE%BC%CE%AD%CE%BD%CE%B1_%CF%84%CE%B7%CF%82_%CE%95%CF%86%CE%BF%CF%81%CE%B5%CE%AF%CE%B1%CF%82_%CE%91%CF%81%CF%87%CE%B1%CE%B9%CE%BF%CF%84%CE%AE%CF%84%CF%89%CE%BD_%CE%86%CF%81%CF%84%CE%B1%CF%82_%CE%BA%CE%B1%CF%84%CE%AC_%CF%84%CE%B1_%CE%AD%CF%84%CE%B7_2014_2020_%CE%86%CF%81%CF%84%CE%B1_2020

11. Vokotopoulou I. (n Greek) Epirus in the 8th and 7th centuries BCE. Annuario della Scuola Acheologica di Atene e delle Missioni Italiane in Oriente. 1982;LX. Available from: https://www.scuoladiatene.it/dal-1981-al-1990/1982.html

12. Malkin I. A Small Greek World: Networks in the Ancient Mediterranean. OUP USA; 2011. Available from: https://books.google.com/books/about/A_Small_Greek_World.html?hl=&id=CKQXm8sNgNkC

13. Malkin I. Foundations. A Companion to Archaic Greece. John Wiley & Sons, Ltd; 2009 p.373–94. Available from: https://onlinelibrary.wiley.com/doi/abs/10.1002/9781444308761.ch19

14. Hornblower S. Thucydides and “Chalkidic” Torone (IV.110.1). Oxford Journal of Archaeology. 1997;16:177–86. Available from: https://onlinelibrary.wiley.com/doi/abs/10.1111/1468-0092.00033

15. Papadopoulos JK. Archaeology, Myth-History and the Tyranny of the Text: Chaldike, Torone and Thucydides. Oxford Journal of Archaeology. 1999;18:377–94. Available from: https://onlinelibrary.wiley.com/doi/abs/10.1111/1468-0092.00091

16. Olalde I, Mallick S, Patterson N, Rohland N, Villalba-Mouco V, Silva M, et al. The genomic history of the Iberian Peninsula over the past 8000 years. Science. 2019;363:1230–4. Available from: 10.1126/science.aav4040

17. Reitsema LJ, Mittnik A, Kyle B, Catalano G, Fabbri PF, Kazmi ACS, et al. The diverse genetic origins of a Classical period Greek army. Proc Natl Acad Sci U S A. 2022;119:e2205272119. Available from: 10.1073/pnas.2205272119

18. Lazaridis I, Alpaslan-Roodenberg S, Acar A, Açıkkol A, Agelarakis A, Aghikyan L, et al. The genetic history of the Southern Arc: A bridge between West Asia and Europe. Science. 2022;377:eabm4247. Available from: 10.1126/science.abm4247

19. Skourtanioti E, Ringbauer H, Gnecchi Ruscone GA, Bianco RA, Burri M, Freund C, et al. Ancient DNA reveals admixture history and endogamy in the prehistoric Aegean. Nat Ecol Evol. 2023;7:290–303. Available from: 10.1038/s41559-022-01952-3

20. Boardman J. The Greeks Overseas: Their Early Colonies and Trade. Thames and Hudson; 1999. Available from: https://play.google.com/store/books/details?id=EqHAQgAACAAJ

21. Mallick S, Micco A, Mah M, Ringbauer H, Lazaridis I, Olalde I, et al. The Allen Ancient DNA Resource (AADR): A curated compendium of ancient human genomes. bioRxiv. 2023; Available from: 10.1101/2023.04.06.535797

22. Alexander DH, Novembre J, Lange K. Fast model-based estimation of ancestry in unrelated individuals. Genome Res. 2009;19:1655–64. Available from: 10.1101/gr.094052.109

23. Maier R, Flegontov P, Flegontova O, Işıldak U, Changmai P, Reich D. On the limits of fitting complex models of population history to f-statistics. Elife. 2023;12. Available from: 10.7554/eLife.85492

24. Wang RJ, Al-Saffar SI, Rogers J, Hahn MW. Human generation times across the past 250,000 years. Science advances. 2023;9. Available from: https://pubmed.ncbi.nlm.nih.gov/36608127/

25. Ringbauer H, Huang Y, Akbari A, Mallick S, Olalde I, Patterson N, et al. Accurate detection of identity-by-descent segments in human ancient DNA. Nat Genet. 2024;56:143–51. Available from: 10.1038/s41588-023-01582-w

26. Ringbauer H, Novembre J, Steinrücken M. Parental relatedness through time revealed by runs of homozygosity in ancient DNA. Nat Commun. 2021;12:5425. Available from: 10.1038/s41467-021-25289-w

27. Angeli A. (In Greek) The cemeteries of Amvrakia during the Archaic and Classical times. National Documentation Centre (EKT); 2021. Available from: https://www.didaktorika.gr/eadd/handle/10442/49590?locale=en

28. Lazaridis I, Alpaslan-Roodenberg S, Acar A, Açıkkol A, Agelarakis A, Aghikyan L, et al. A genetic probe into the ancient and medieval history of Southern Europe and West Asia. Science. 2022; Available from: https://www.science.org/doi/10.1126/science.abq0755

29. Hofmanová Z, Kreutzer S, Hellenthal G, Sell C, Diekmann Y, Díez-del-Molino D, et al. Early farmers from across Europe directly descended from Neolithic Aegeans. Proceedings of the National Academy of Sciences. 2016;113:6886–91. Available from: https://www.pnas.org/doi/abs/10.1073/pnas.1523951113

30. Marchi N, Winkelbach L, Schulz I, Brami M, Hofmanová Z, Blöcher J, et al. The genomic origins of the world’s first farmers. Cell. 2022;185:1842–59.e18. Available from: 10.1016/j.cell.2022.04.008

31. Mathieson I, Lazaridis I, Rohland N, Mallick S, Patterson N, Roodenberg SA, et al. Genome-wide patterns of selection in 230 ancient Eurasians. Nature. 2015;528:499–503. Available from: https://www.nature.com/articles/nature16152

32. Burger J, Link V, Blöcher J, Schulz A, Sell C, Pochon Z, et al. Low Prevalence of Lactase Persistence in Bronze Age Europe Indicates Ongoing Strong Selection over the Last 3,000 Years. Curr Biol. 2020;30:4307–15.e13. Available from: 10.1016/j.cub.2020.08.033

33. Brooke JL, Larsen CS. The Nurture of Nature: Genetics, Epigenetics, and Environment in Human Biohistory. Am Hist Rev. 2014;119:1500–13. Available from: https://academic.oup.com/ahr/article/119/5/1500/44610

34. Amorim CEG, Acuña-Alonzo V, Salzano FM, Bortolini MC, Hünemeier T. Differing evolutionary histories of the ACTN3*R577X polymorphism among the major human geographic groups. PLoS One. 2015;10:e0115449. Available from: 10.1371/journal.pone.0115449

35. Boussiou M, Karababa P, Sinopoulou K, Tsaftaridis P, Plata E, Loutradi-Anagnostou A. The molecular heterogeneity of β-thalassemia in Greece. Blood Cells Mol Dis. 2008;40:317–9. Available from: https://www.sciencedirect.com/science/article/pii/S1079979607002549

36. Georgiou I, Makis A, Chaidos A, Bouba I, Hatzi E, Kranas V, et al. Distribution and frequency of beta-thalassemia mutations in northwestern and central Greece. Eur J Haematol. 2003;70:75–8. Available from: 10.1034/j.1600-0609.2003.00017.x

37. Hughey JR, Du M, Li Q, Michalodimitrakis M, Stamatoyannopoulos G. A search for β thalassemia mutations in 4000year old ancient DNAs of Minoan Cretans. Blood Cells Mol Dis. 2012;48:7–10. Available from: https://www.sciencedirect.com/science/article/pii/S1079979611001744

38. Margaryan A, Hansen HB, Rasmussen S, Sikora M, Moiseyev V, Khoklov A, et al. Ancient pathogen DNA in human teeth and petrous bones. Ecol Evol. 2018;8:3534–42. Available from: https://onlinelibrary.wiley.com/doi/abs/10.1002/ece3.3924

39. The genomic history of the Aegean palatial civilizations. Cell. 202;184:2565–86.e21. Available from: 10.1016/j.cell.2021.03.039

40. Lazaridis I, Mittnik A, Patterson N, Mallick S, Rohland N, Pfrengle S, et al. Genetic origins of the Minoans and Mycenaeans. Nature. 2017;548:214–8. Available from: 10.1038/nature23310

41. van Dommelen P. Colonialism and Migration in the Ancient Mediterranean. Annu Rev Anthropol. 2012;41:393–409. Available from: https://www.annualreviews.org/content/journals/10.1146/annurev-anthro-081309-145758

42. Morakis A. THUCYDIDES AND THE CHARACTER OF GREEK COLONISATION IN SICILY. Class Q. 2011;61:460–92. Available from: https://www.cambridge.org/core/product/identifier/S0009838811000188/type/journal_article

43. Kaponis AS. Intertemporal memories of a shifting unity. C&M. 2024;221–57. Available from: https://tidsskrift.dk/classicaetmediaevalia/article/view/145251

44. Bresson A. Quatre emporia antiques : Abul, La Picola, Elizavetovskoie, Naucratis. Rev études anc. 2002;104:475–505. Available from: https://www.persee.fr/doc/rea_0035-2004_2002_num_104_3_4880

45. Demetriou D. What is an emporion? A reassessment. Historia. 2011; Available from: http://www.jstor.org/stable/41342851

46. Pease AS. Notes on the Delphic Oracle and Greek Colonization. Class Philol. 1917;12:1–20. Available from: http://www.jstor.org/stable/262478

47. Holladay AJ. Sparta’s role in the First Peloponnesian War. J Hell Stud. 1977;97:54–63. Available from: https://www.cambridge.org/core/journals/journal-of-hellenic-studies/article/abs/spartas-role-in-the-first-peloponnesian-war/1638F06647CE44F04FD58AC359ECA059

48. Katsivardelos P. The Military and Political Role of the Allies of Sparta in the Peloponnesian War [MLitt(R)]. [Ann Arbor :]: University of Glasgow; 1992. Available from: https://theses.gla.ac.uk/75255/

49. Perlman S. The causes and the outbreak of the Corinthian war. Class Q. 1964;14:64–81. Available from: http://www.jstor.org/stable/637630

50. Kaponis A. The image of Corinthian colonies founded around the Ambracian Gulf through the ancient written tradition. Mare Ponticum. 2022;10. Available from: https://www.academia.edu/100444454/%CE%91%CE%BD%CF%84%CF%8E%CE%BD%CE%B9%CE%BF%CF%82_%CE%A3_%CE%9A%CE%B1%CF%80%CF%8E%CE%BD%CE%B7%CF%82_%CE%97_%CE%B5%CE%B9%CE%BA%CF%8C%CE%BD%CE%B1_%CF%84%CF%89%CE%BD_%CE%BA%CE%BF%CF%81%CE%B9%CE%BD%CE%B8%CE%B9%CE%B1%CE%BA%CF%8E%CE%BD_%CE%B1%CF%80%CE%BF%CE%B9%CE%BA%CE%B9%CF%8E%CE%BD_%CF%80%CE%B5%CF%81%CE%AF_%CF%84%CE%BF%CE%BD_%CE%91%CE%BC%CE%B2%CF%81%CE%B1%CE%BA%CE%B9%CE%BA%CF%8C_%CE%BA%CF%8C%CE%BB%CF%80%CE%BF_%CE%BC%CE%AD%CF%83%CE%B1_%CE%B1%CF%80%CF%8C_%CF%84%CE%B7_%CE%B3%CF%81%CE%B1%CF%80%CF%84%CE%AE_%CF%80%CE%B1%CF%81%CE%AC%CE%B4%CE%BF%CF%83%CE%B7_%CF%84%CE%B7%CF%82_%CE%B1%CF%81%CF%87%CE%B1%CE%B9%CF%8C%CF%84%CE%B7%CF%84%CE%B1%CF%82_A_S_Kaponis_The_image_of_Corinthian_colonies_founded_around_the_Ambracian_Gulf_through_the_ancient_written_tradition

51. Domínguez AJ. Not Only “invincible in arms, a glorious warrior” (Plut. Pyrrh. 11.8). Pyrrhus and the Administration of the Epirote Kingdom. Klio. 2022 [cited 2024 Sep 4];104:550–86. Available from: https://www.degruyter.com/document/doi/10.1515/klio-2021-0033/html

52. Harney É, Cheronet O, Fernandes DM, Sirak K, Mah M, Bernardos R, et al. A minimally destructive protocol for DNA extraction from ancient teeth. Genome Res. 2021;31:472–83. Available from: 10.1101/gr.267534.120

53. Rohland N, Glocke I, Aximu-Petri A, Meyer M. Extraction of highly degraded DNA from ancient bones, teeth and sediments for high-throughput sequencing. Nat Protoc. 2018;13:2447–61. Available from: 10.1038/s41596-018-0050-5

54. Orfanou E, Himmel M, Aron F, Haak W. Minimally-invasive sampling of pars petrosa (os temporale) for ancient DNA extraction v2. protocols.io. ZappyLab, Inc.; 2020. Available from: https://www.protocols.io/view/minimally-invasive-sampling-of-pars-petrosa-os-tem-bqd8ms9w

55. Allentoft ME, Sikora M, Sjögren K-G, Rasmussen S, Rasmussen M, Stenderup J, et al. Population genomics of Bronze Age Eurasia. Nature. 2015;522:167–72. Available from: 10.1038/nature14507

56. Meyer M, Kircher M. Illumina sequencing library preparation for highly multiplexed target capture and sequencing. Cold Spring Harb Protoc. 2010;2010:db.prot5448. Available from: 10.1101/pdb.prot5448

57. Rohland N, Harney E, Mallick S, Nordenfelt S, Reich D. Partial uracil–DNA–glycosylase treatment for screening of ancient DNA. Philos Trans R Soc Lond B Biol Sci. 2015;370:20130624. Available from: 10.1098/rstb.2013.0624

58. Neuenschwander S, Cruz Dávalos DI, Anchieri L, Sousa da Mota B, Bozzi D, Rubinacci S, et al. Mapache: a flexible pipeline to map ancient DNA. Bioinformatics. 2023;39. Available from: 10.1093/bioinformatics/btad028

59. Fu Q, Mittnik A, Johnson PLF, Bos K, Lari M, Bollongino R, et al. A revised timescale for human evolution based on ancient mitochondrial genomes. Curr Biol. 2013;23:553–9. Available from: 10.1016/j.cub.2013.02.044

60. Renaud G, Slon V, Duggan AT, Kelso J. Schmutzi: estimation of contamination and endogenous mitochondrial consensus calling for ancient DNA. Genome Biol. 2015;16:224. Available from: 10.1186/s13059-015-0776-0

61. Korneliussen TS, Albrechtsen A, Nielsen R. ANGSD: Analysis of Next Generation Sequencing Data. BMC Bioinformatics. 2014;15:356. Available from: 10.1186/s12859-014-0356-4

62. Schönherr S, Weissensteiner H, Kronenberg F, Forer L. Haplogrep 3 - an interactive haplogroup classification and analysis platform. Nucleic Acids Res. 2023;51:W263–8. Available from: 10.1093/nar/gkad284

63. Rubin JD, Vogel NA, Gopalakrishnan S, Sackett PW, Renaud G. HaploCart: Human mtDNA haplogroup classification using a pangenomic reference graph. PLoS Comput Biol. 2023;19:e1011148. Available from: 10.1371/journal.pcbi.1011148

64. Ralf A, Montiel González D, Zhong K, Kayser M. Yleaf: Software for Human Y-Chromosomal Haplogroup Inference from Next-Generation Sequencing Data. Mol Biol Evol. 2018;35:1291–4. Available from: https://academic.oup.com/mbe/article-pdf/35/5/1291/24704771/msy032.pdf

65. David Poznik G. Identifying Y-chromosome haplogroups in arbitrarily large samples of sequenced or genotyped men. bioRxiv. 2016. p. 088716. Available from: https://www.biorxiv.org/content/10.1101/088716v1.abstract

66. Kuhn JMM, Jakobsson M, Günther T. Estimating genetic kin relationships in prehistoric populations. PLoS One. 2018;13:e0195491. Available from: https://journals.plos.org/plosone/article/file?id=10.1371/journal.pone.0195491&type=printable

67. Popli D, Peyrégne S, Peter BM. KIN: a method to infer relatedness from low-coverage ancient DNA. Genome Biol. 2023;24:1–22. Available from: https://genomebiology.biomedcentral.com/articles/10.1186/s13059-023-02847-7

68. A global reference for human genetic variation. Nature. 2015;526:68–74. Available from: https://www.nature.com/articles/nature15393

69. Rubinacci S, Ribeiro DM, Hofmeister RJ, Delaneau O. Efficient phasing and imputation of low-coverage sequencing data using large reference panels. Nat Genet. 2021;53:120–6. Available from: 10.1038/s41588-020-00756-0

70. Link V, Kousathanas A, Veeramah K, Sell C, Scheu A, Wegmann D. ATLAS: Analysis Tools for Low-depth and Ancient Samples. bioRxiv. 2017. p. 105346. Available from: https://www.biorxiv.org/content/10.1101/105346v2.abstract

71. International HapMap Consortium, Frazer KA, Ballinger DG, Cox DR, Hinds DA, Stuve LL, et al. A second generation human haplotype map of over 3.1 million SNPs. Nature. 2007;449:851–61. Available from: 10.1038/nature06258

72. van de Loosdrecht M, Bouzouggar A, Humphrey L, Posth C, Barton N, Aximu-Petri A, et al. Pleistocene North African genomes link Near Eastern and sub-Saharan African human populations. Science. 2018;360:548–52. Available from: 10.1126/science.aar8380

73. Patterson N, Price AL, Reich D. Population Structure and Eigenanalysis. PLoS Genet. 2006;2:e190. Available from: https://journals.plos.org/plosgenetics/article/file?id=10.1371/journal.pgen.0020190&type=printable

74. Haag J, Jordan AI, Stamatakis A. Pandora: A Tool to Estimate Dimensionality Reduction Stability of Genotype Data. bioRxiv. 2024. p. 2024.03.14.584962. Available from: https://www.biorxiv.org/content/10.1101/2024.03.14.584962v1

75. Harney É, Patterson N, Reich D, Wakeley J. Assessing the performance of qpAdm: a statistical tool for studying population admixture. Genetics. 2021;217. Available from: 10.1093/genetics/iyaa045

76. Chaitanya L, Breslin K, Zuñiga S, Wirken L, Pośpiech E, Kukla-Bartoszek M, et al. The HIrisPlex-S system for eye, hair and skin colour prediction from DNA: Introduction and forensic developmental validation. Forensic Sci Int Genet. 2018;35:123–35. Available from: 10.1016/j.fsigen.2018.04.004

77. Pochon Z, Bergfeldt N, Kırdök E, Vicente M, Naidoo T, van der Valk T, et al. aMeta: an accurate and memory-efficient ancient metagenomic profiling workflow. Genome Biol. 2023;24:242. Available from: 10.1186/s13059-023-03083-9

78. Nafplioti A. Tracing population mobility in the Aegean using isotope geochemistry: a first map of local biologically available 87Sr/86Sr signatures. J Archaeol Sci. 2011;38:1560–70. Available from: https://www.sciencedirect.com/science/article/pii/S0305440311000604

