## Additional file 7. Supplementary Info and Figures for "Genetic affinities between an ancient Greek colony and its metropolis: the case of Amvrakia in western Greece"

#### Supplementary Information

#### Table of Contents

1. Archaeological Background 3
1.1. Ancient Greek Colonization 3
1.2. Corinthian colonization 5
1.3 The colony of Amvrakia 5
1.4 Tenea as a proxy for Ancient Corinthian ancestry 7
2. Archaeological and anthropological context of sampled individuals 10
2.1 Amvrakia cemeteries 10
2.1.1 Graves associated with the deep-sequenced individuals of the present study 12
2.2. Ammotopos - “Kokkino Lithari” site 25
2.3. Cemeteries of Tenea 27
3. Ancient DNA Analysis 40
3.1 Sample Preparation 40
3.2 Read Processing, Damage estimation, Genetic sex determination 42
3.2.1 Analyses at the FastQ level 43
3.2.2 Analyses at the library level 44
3.2.3 Analyses at the individual level 45
3.3 Contamination estimation 47
3.4 Uniparental haplogroup estimation 48
3.4.1 mtDNA 48
3.4.2 Y-chromosome 49
3.5 Population Genomics analysis 50
3.5.1 Lists of genomic sites 50
3.5.2 Genotype calling and pseudohaploidization 50
3.5.3 Genetic relatedness analysis 51

3.5.4 Runs of homozygosity 55
3.5.5 Imputation and Identity-by-Descent segments screening 55
3.5.6 Merging with public data 57
3.5.7 Principal Component Analysis 58
3.5.8 Population clustering analysis (ADMIXTURE) 61
3.5.9 f3 Statistics and Ancestry Proportion Analysis (qpAdm) 72
3.6 Phenotypes 105
3.6.1. Pigmentation 105
3.6.2. Monogenic Phenotypes 105
3.7 Microbial Metagenomics 107
3.8 Visualization 107
4. Provenance, mobility and diet analysis using stable isotopes 109
4.1 Strontium isotope ratio analysis of bioarchaeological skeletal remains: principles 109
4.2 Geological context of the study area 110
4.3 Materials and Methods 112
4.3.1 Samples 112
4.3.2 Sample preparation and analysis 112
4.4 Results and Discussion 112
5. Supplementary Information References 114

### 1. Archaeological Background

***Eugenia Tabakaki, Christina Papageorgopoulou, Aggeliki Georgiadou, Elena Korka, Paraskevi Evaggeloglou, Ioannis Christidis, Michael Ioannou, Panagiotis Panailidis***

#### 1.1. Ancient Greek Colonization

Migrations are ubiquitous in human history. From the great out-of-Africa journey, over the Neolithic demographic transition, and up to present times, there has been a constant flow of migratory waves across the planet.

An important outcome of migratory expeditions and local interactions is the ancient Greek civilization, a period of political, philosophical, artistic, and scientific achievements that formed a legacy with substantial influence on Western civilization. Having its roots in the post-Mycenaean era usually described as the Greek Dark Ages, it is almost universally recognised that it started to take shape during the Archaic period (c. 800-479 BCE), as an outcome of migratory movements and interactions in the Mediterranean and the Black Sea, known as ancient Greek colonization.

The Ancient Greek Colonization started during the 8th century BCE. During this period, *poleis*[1] (cities) in continental Greece, the Aegean, and along the coast of Asia Minor initiated colonization campaigns, both towards the West, including western Greece, southern Italy, Sicily, south France, and Spain, and towards the East, expanding from the northern Aegean coast and the islands, all the way to the Black Sea [3–7]. In particular, the northern Aegean and north-western Greece were heavily colonized by major Euboean centers, Corinth, the Cycladic islands, and Ionian cities. At present more than 100 colonies from Amvrakia and Corcyra to Chalcidice, the Thermaic Gulf, and further East to the Thracian coast and the islands of Thasos and Samothrace are known. The Colonisation of the Black Sea was initially undertaken by the Greek cities of Asia Minor under the leading role of Miletus and until the end of the 6th century BCE when the Athenian expansion policy began to overshadow all other attempts in the area with a record breaking 200 colonies [7,8].

Ancient Greeks used the word *apoikia* (*Αποικία*), a term that may best be translated as "home away from home", predominantly emphasizing the separation, but also the

[1] The central focus of civilization for the Greeks, after the oikos or family unit, was the polis (plural: poleis). Polis is usually translated as ‘city-state’, as polis was generally an independent state, with its own laws, customs, political system, military force, currency and sometimes calendar. According to Aristotle those who did not live in a polis were ‘tribeless, lawless, heartless’, and to the Greeks the fact that they lived in a city-state was proof that they were a civilized people. But the polis should also in Aristotle’s opinion be limited in size and self-sufficient [1]. The concept of the polis mattered to the Greeks. They did not just live in poleis, they found it important to live in poleis rather than in some other form of political community. Every Greek colony was founded as a polis or became a polis not long after its foundation. Nevertheless, no one has ever investigated how many poleis there were and which settlements were actually poleis. For the colonies there is no comprehensive study at all. The polis and the concept of polis have been investigated either in general or in relation to one individual polis. The general studies are mostly based on sources relating to Athens, and most of the individual studies deal with the Athenian democratic polis of the Classical period or with Archaic and Classical Sparta [2].

connection between the *metropolis*[2] ("mother city") and the new *oikos*[3] [9]. The colonies were usually sovereign states and not dominions. Therefore scholars emphasize the misleading connotation of the term "colonization" compared to the modern colonial era [10,11]. The relationships between the metropoles and colonies were bidirectional and typically beneficial for both entities. In some cases, the colonies outperformed the metropoles with respect to cultural and political developments. Each colony had its individual history and founding myth, colonies of its own, allies, and special ties to the metropolis. Despite these distinct profiles, common features are observed among the foundation myths of most colonies, such as the substantial role of the Delphic oracle in indicating new lands; the figure of the *oikistes*, the divine or mortal hero whose name was often given to the colony and may have led the group of settlers; the foundation of the first sanctuaries and the determination of the social order following the *nomima* (the laws and traditions of the metropolis) [12]. Based on these data, archaeologists and historians identify a colony *vis-à-vis* the metropolis, although this connection is not always straightforward.

Their search for new homes was instigated for a plethora of reasons: internal strife, social conflicts, political strategies, famine or poverty, a quest for better opportunities. *Emporia* (trading posts) were in some cases the predecessors of colonies. Yet, starting in the mid-8th century BCE the Greek *poleis* (city-states), as well as regional or ethnic groups started to expand with more targeted, longer-term intentions. The founding of hundreds of new cities, within a wide and geographically diverse range, the interaction of the settlers with heterogeneous indigenous populations, and the relatively short time in which the colonies evolved into major cultural, commercial, and political centers indicates that Greek Colonization constitutes one of the most influential phenomena in European ancient history and archaeology.

Although numerous colonies have been excavated, the debate surrounding the mode, intensity, and pace of these migratory movements [e.g. 13,14] remains unresolved. Were these movements more akin to a gradual drift, or did they represent an organized colonization effort? Can we reliably identify biological, linguistic, religious, cultural, or social groups whose origins might have been as much a product of invented founding myths as of reality? What role did the local populations play—were they assimilated or differentiated? The settlement

[2] The Greek colonization is temporally and causally connected with the very creation of the Greek city as an organized city - urban community of citizens. We can not actually date the birth of the city state. The oldest legal text of Greek antiquity, an inscription of the Cretan Dreros (c. 630 BCE), presupposes the existence of the city in the sense of the city-state. The term "metropolis" itself implies the existence of a city. In Greek "metropolis" meant the "mother city" and refers to the relationship between cities and colonies. The term metropolis has a highly complex definition. The 'mother-city' of a Greek colony (apoikia) usually nominated the founder (oikistēs), conducted rituals of divination and departure, organized a body of settlers, and formulated the charter of their individual status [4].

[3] The term "Oikos" in ancient Greece did not mean only "house", as it does today, nor did it denote only a family (i.e. the sum of the members of a group of people who are connected by family ties). The oikos, as Aristotle says in his Politics, was the smallest unit, the smallest component of society; of course, especially in archaic societies, the oikos was also linked to the possession of land (this was no longer necessarily true in societies such as 5th century Athens, with its developed commercial and craft economy). The characteristic of the house, however, at the ideological level, was that it had a continuity through time, it had a past (the ancestors) and a future (the descendants). Therefore, it was the duty of every adult male to respect the past of his house and to ensure its continuity into the future. Therefore, the social attitude and ethics of each man had consequences not only individually, but also in terms of the maintenance and preservation of the prestige of his house.

nuclei likely attracted many others, possibly including local women. But what was their role? Many of these pressing questions are anthropological and population genetic in nature, and they have been only partially addressed to date.

#### 1.2. Corinthian colonization

Colonization was not a flight into the unknown, but the creation of a stable network of economic and political ties [15,16]. This is particularly evident in the case of Corinth, which grew into a powerful metropolitan city through the foundation of colonies and trading posts across the Mediterranean (Corcyra, Leukas, Syracuse), the Adriatic (Epidamnus, Apollonia), the Aegean (Potidaea in Chalcidice), and mainland Greece (Amvrakia). Historical texts and archaeological evidence (artifacts, burials, grave goods, coins) reveal the substantial influence of Corinth on the colonies' lifestyle and economy [17–19]. Based on its colonies, Corinth developed a trade route, from the Saronic Sea, through the Corinthian Gulf, along the western coasts of Sterea Ellada (geographical unit of modern-day Greece), Epirus, and the Adriatic to Sicily [20]. The main trading stations (and colonies at the same time) were Anactorion, Ithaca, Lefkas, Corcyra [21], Amvrakia [18,22], Apollonia [22,23], and Epidamnus [22].

Through this network, Corinth established a dominance in the West, as observed by Thucydides. Within a generation after the foundation of Syracuse, by about 700 BCE, Corinth monopolized the western trade. This is substantiated by the archaeological findings of Corinthian origin covering the end of the 8th century BCE and the beginning of the 7th century BCE in the West that exceed the findings from all other Greek poleis combined [24]. Corinth exported pottery, textiles, metalwork, ivory-carving, sculpture, and imported grain, as well as the raw materials it lacked. During this period, the Corinthian architectural style experienced a new phase of rapid commercial development.

Overall, the city of Corinth was an active and competitive metropolis in Archaic times and had intense relations with the cities it founded. The colonies remained under Corinthian control, that is, having their coins minted in Corinth as late as the fifth century BCE. However, there also existed conflicts between the metropolis and its colonies. The dispute between the Corinthians and Corcyraeans in the mid of the 5th century BCE over the control of Epidamnus is a representative example [25].

Corinth contributed to the history of the ancient Mediterranean world through the foundation of its colonies, some of which evolved into major cities of the ancient and modern world.

#### 1.3 The colony of Amvrakia

Amvrakia (also known as Ambracia) was founded by Corinth during the last half of the 7th century BCE on the banks of the river Arachthos [26] and at a short distance from the northern coast of the Amvrakian Gulf [22]. Already in the 8th century BCE, before the formal establishment of the colony, Corinth had established a trading post to support the wide trade network it had developed along the Ionian and Adriatic coasts. Individual burials found in the northern part of the city confirm this [27].

Amvrakia was founded on an important crossroad that connected southern Greece with the mainland of Epirus and reached as far as Apollonia, another Corinthian colony on the eastern Adriatic coast. Amvrakia had access to maritime trade routes through the Arachthos river. In the area of the Amvrakian Gulf, Corinth had also founded the colonies of Lefkada,

Anaktorion, and Sollion known as the “sister-cities” of Amvrakia. They all formed part of the Corinthian trade network.

According to the foundation myth, the Corinthians, led by Gorgos (oikist), son of Cypselos, the tyrant of Corinth, founded a colony on the banks of the Arachthos river [28]. The foundation of Amvrakia, is evidenced by the strong presence of the metropolis in the material culture of the ancient city. The archaeological, epigraphic, and numismatic finds indicate that there was a strong relationship between the colony and its metropolis. Amvrakia strengthened Corinth's economic influence and commercial control in northwestern Greece. This colony, situated at the verge of the Hellenic world, was the connecting link that united the Greek city-states, especially in the Epirus-Illyrian region.

Based on archaeological evidence, the latter archaic city of Amvrakia had strong fortifications protecting the city, and was centrally planned and organized. The city was built according to a geometric urban masterplan. Streets of N-S direction intersected with avenues creating residential blocks. Each urban block measured 150 × 30 m and contained about twenty houses. This urban organization remained unchanged throughout the centuries. Each house, following the principles of isometry and isonomy had the same dimensions (approximately 15 × 15 m). The urban plan of Amvrakia dedicated the north-western part of the town to the public life of the town [29]. The administrative and religious center with monumental public buildings and temples was located in the north-western part of the city. Private residences of the classical and Hellenistic periods had almost identical shapes with those of the archaic period. Successive phases of the private houses of the Archaic, Classical, and Hellenistic periods are a common place in Amvrakia. This means that life continued uninterrupted in the city.

During the 6th century BCE, the city developed demographically and politically and transitioned from tyranny to a form of representative democracy. In 582 BCE, it acquired a democratic constitution by rebelling against the tyrants as stated by Aristotle [30]. In the 5th century BCE, the city minted its own coins following the Corinthian model.

The Amvrakians took part in the Persian Wars and the Peloponnesian War supporting the metropolis of Corinth. The prime of Amvrakia was during the Hellenistic period under the reign of Pyrrhus, the King of Epirus (c. 319-272 BCE). In 295 BCE, Pyrrhus established Amvrakia as the capital of his kingdom. Representative public and private buildings and spaces constituted a model city that provided a good quality of life to its citizens until the conquest by the Romans under Aemilius Paulus in 167 BCE. Amvrakia declined in 31 BCE, when the residents were forced to populate the neighboring Nicopolis, which was a city founded by Octavian Augustus [31].

### 1.4 Tenea as a proxy for Ancient Corinthian ancestry

Tenea was an important settlement located in the eastern periphery of Corinth at a key strategic position controlling Kontoporeia, the shortest path leading from Corinth to Mycenae and Argos (**Supplementary Figure S1**). For many centuries, Tenea was the largest and most important "*Kome*[4]" (town) in the eastern region of Corinth. The status of Tenea as Kome was different to that of the independent polis. It was a community or settlement, in the Dorian Peloponnese, which joined forces [2].

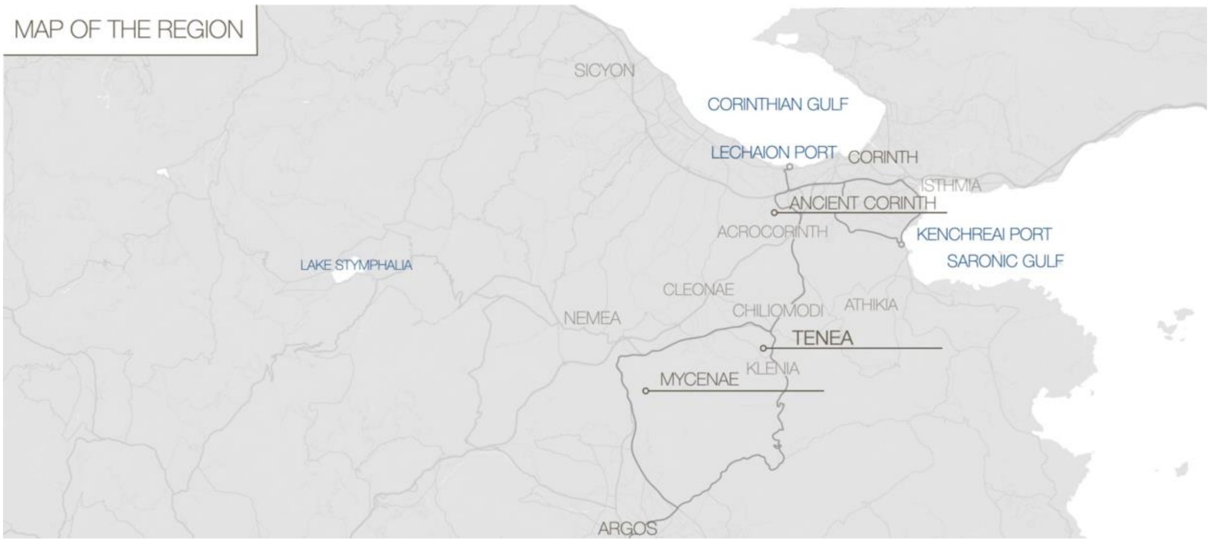


**Supplementary Figure S1.** Map of the Corinthia region, with the respective location of Tenea and Ancient Corinth.

The city is mentioned in Strabo [1] and Pausanias [2]. Strabo provides more information on Tenea than almost all the smaller towns of the Peloponnese, which emphasizes its importance [32]. According to written sources, Tenea was the place where Oedipus was brought up in the summer palace of Polybus, king of Corinth. In the Archaic and Classical periods, the Greek city of Tenea in Corinthia was part of the city-state of Corinth. Wiseman [33] mentions "Citizens of all the towns of the Korinthia evidently considered themselves, throughout most of the antiquity citizens of Corinth". Tenea developed substantially, and, in the 8th century BCE, a large part of its population actively participated in the Greek colonization under Corinth. Together with Corinth, the two cities provide an example of a common colonization. For instance, in 734 BCE, residents of Tenea participated in the colonization of Syracuse led by Archias from Corinth [16]. During this period, although being under the direct influence of Corinth, Tenea had already evolved into a populous and

[4] The status of kome is different to that of the independent polis. It is a form of community, "municipality" or "village", but not as developed and valuable as the polis, which is the perfect form of human society. We hear about komai in the Dorian Peloponnese, in some parts of central and western Greece, in Makedonia and Thrace and along the west coast of Asia Minor. It is believed that the term kome was Dorian, whereas the term demos was used in the non-Dorian parts of Hellas. Kome is more a notion of how the Greeks designated and classified settlements that were not Poleis. The overlap between the two terms seems to occur principally when kome is used in a political sense about a subdivision of a larger polis [2].

prosperous city with an extensive region, *Teneatis*[5], which is confirmed by the latest archaeological evidence. The participation of Tenea in the colonization of Syracuse shows that it was a well-developed city and that the colonization contributed decisively to the prosperity of its society, due to the commercial contacts that were developed. Tenea had always had a tendency for independence. During Hellenistic times, Tenea attempted unsuccessfully to gain autonomy from the Corinthians by minting its own currency [34]. Only during the late Hellenistic period, Tenea became a free city in the sense that it had its own government and was not under Corinthian authority any more. Its development, however, attained its peak during the Roman period, since it was the only city of Corinth that was not destroyed by the Romans [35]. This political status appears to have been maintained by the Romans after they had conquered Greece. According to Pausanias: "The inhabitants (of Tenea) say that they are Trojans who were taken prisoners in Tenedos by the Greeks, and were permitted by Agamemnon to dwell in their present home". The common Trojan mythical origin of the residents of Tenea and the Romans, from the lineage of Tennes, king of Tenedos, and the trojan hero Aeneas [22], seems to have created a strong link between them. The city was favored by the Romans, not only because of their supposed common ancestry from Troy but also because of its support for them during the war against the Achaean League. The myth of common ancestry came to help and justified the Roman decision. The earliest reference to Tenea being independent from Corinth, sometime before 146 BCE, is by Strabo who claims that the city had already gained independence when it joined the Romans against Corinth in that year: "Tenea prospered more than the other settlements (in Corinthia), and finally even had a government of its own, and, revolting from the Corinthians, joined the Romans, and endured after the destruction of Corinth". Tenea remained an important center during the Roman era, until the 6th century CE, when it was abandoned due to the raids of the Avaro-Slavs.

Today, systematic archaeological research[6] conducted in the area gradually reveals aspects of the ancient city. In general, the first results of the research show rich and remarkable activity in Tenea from the Archaic to the late Roman times. However, the oldest settlement evidence in the area dates to the Early Bronze Age. During excavations, remains of the Early Helladic period were revealed, such as a built ritual deposit that is about four meters deep, as well as part of the Early Helladic settlement, which came to light for the first time. The discovery of evidence for Early Helladic habitation in the area confirms the strategic importance of the site over the course of centuries. New findings from prehistoric Corinth and other prehistoric settlements around Tenea, help us gain new insights regarding the area between the modern towns of Chiliomodi and Klenia before the Greek colonization of Sicily and Southern Italy. It is worth noting the strong presence of imported pottery from Aegina, Attica, Argolis, Corinth, and the Cyclades, which indicates the contacts Tenea had developed with distant regions.

All the above evidence that has been uncovered in the context of the project prove that Tenea was indeed a developed area, with significant activity from the prehistoric to the late Roman period. The participation of Tenea at the colonization of Syracuse supports the

[5] Teneatis included the present-day villages of Stefani, Agios Vasileios, Athikia, Mapsos, Koutalas, Agionori, and Spathovouni.

[6] The archaeological program, known as "Tenea Project", is the first systematic archaeological research in the region of ancient Tenea and is being conducted since 2013 under the direction of Dr. Elena Korka, implemented by the Directorate of Prehistoric and Classical Antiquities of the Hellenic Ministry of Culture. It is also supported by an interdisciplinary team in the context of various scientific collaborations with Greek academic institutions.

observation that the city was in a state of economic, demographic, and cultural prosperity [36]. Regarding the colonization of Amvrakia, there is no ancient text or other source or inscriptions indicating that the people of Tenea were actively involved in the colonization process. However, as Tenea was a *kome* of Corinth, belongs to the region of Corinthia and has participated in its colonization efforts, therefore we can also not exclude their presence in the foundation of Amvrakia either.

### 2. Archaeological and anthropological context of sampled individuals

***Eugenia Tabakaki, Aggeliki Georgiadou, Kiriakos Xanthopoulos, Panagiotis Panailidis, Dimitra Papakosta, Varvara Papadopoulou, Elena Korka, Paraskevi Evaggeloglou, Ioannis Christidis, Michael Ioannou, Theodora Kontogianni, Arkoumanis Athanasios***

#### 2.1 Amvrakia cemeteries

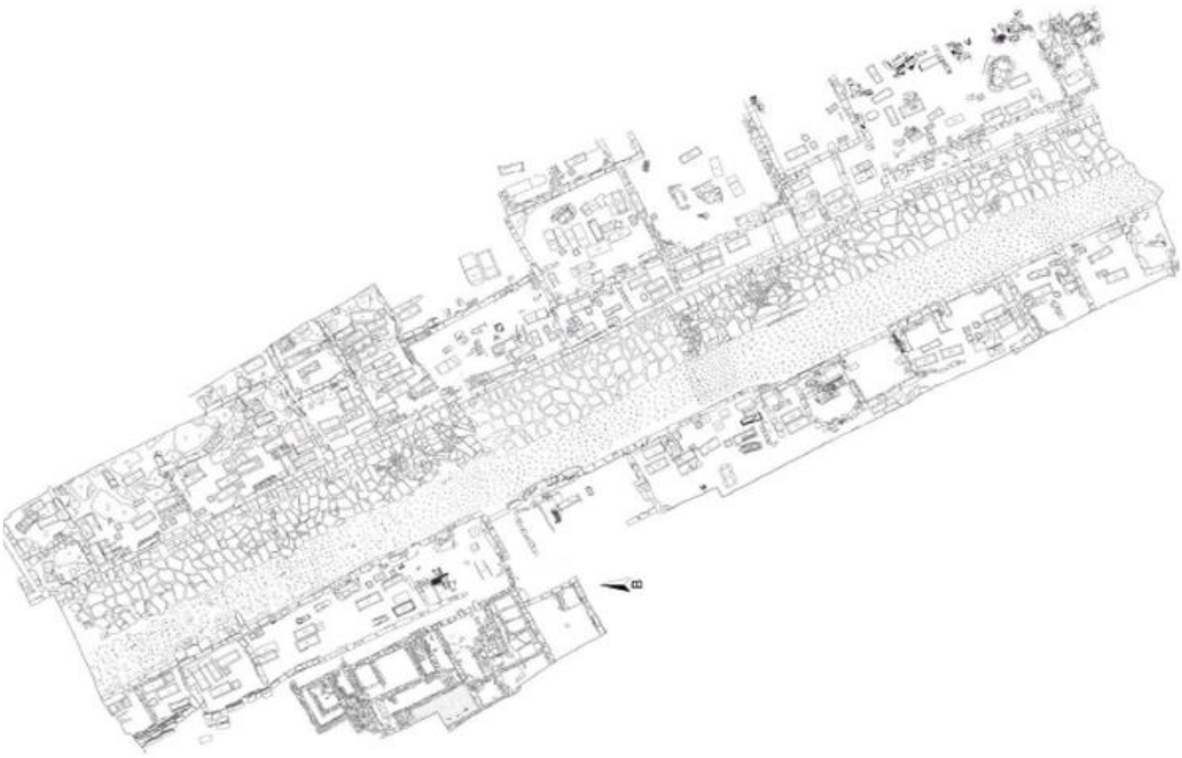

**Supplementary Figure S2.** The western cemetery of Ancient Amvrakia.

The earliest burial evidence dates back to the 8th-7th century BCE, before the official foundation of the city as a Corinthian colony. Individual burials in jars found in the northern part of the city date back to this period. From the 6th century BCE onwards, the two cemeteries of the city were organized in areas outside the city walls, namely on the eastern and south-western fringes of the Peranthis hill.

The southwestern cemetery (**Supplementary Figure S2)** was larger and better organized [37]. A monumental road, which started from the southern gate of the city wall and led to the port in the Amvrakian Gulf, traversed it. This 10-12 m wide road was paved in its eastern part with an elevated pavement, while in its western part it was made of simpler materials. On either side of the avenue, there were retaining limestone walls with elaborate carving, which defined the fronts of the square enclosures (periboloi). The most monumental enclosure was the “Polyandrion” (a cenotaph), erected by the city of Amvrakia in honor of its dead warriors.

The enclosures contained a large number of burials, dating from the 6th century BCE to the early Roman period. The arrangement of the tombs inside the “periboloi” is very dense and testifies the intensive use of the necropolis until the late Hellenistic period. The Π-shaped burial enclosures, bearing facades, contained various cist, tile-covered, and pit graves, sarcophagi, as well as burial cases with copper or clay vessels holding the remains of the cremations. In many cases, the Hellenistic tombs are found on the cist graves and burial cist cases of the Classical era [27].

The cemetery took its final form during the late Classical and especially the Hellenistic period. Graves were made for one individual with some exceptions that contained two burials. During the Hellenistic period reusing the graves became common practice. These graves were probably used by extended families. Numerous burial offerings were placed in the tombs. A total of 700 graves have been investigated in the Necropolis [26].

In Amvrakia, inhumation was the most common practice and cremation was very limited. It appears that all the dead were buried in an extended posture, with the hands parallel to their body. The orientation rule is N-S and the head of the deceased is placed toward the south. In general, the predominant tomb type is the cist grave, in different variations. More than half of the graves are of the cist type. The limestone cist graves were also used as family tombs. In many cases they contained more than one burial, that is, the bones of the older burials were set aside for the new burial. In some cases, the graves also contained cremation urns or vessels. Porous sarcophagi have also been excavated, but they do not represent the common burial type.

During the last period of the Archaic era, especially the years 500-480 BCE, the number of burials in both cemeteries of the city increased substantially. The majority of adult burials are accompanied by at least one drinking vessel as grave goods.

The cemeteries of Amvrakia are different in comparison with the Epirus hinterland, (tombs in the valley of Gormos in Pogoni, Vitsa in Zagori, Liatovouni in Konitsa, and Dourouti Ioannina). In some cases in Epirus, tumuli had been erected (covering a period: LBA - 3rd century BCE), but never became the main burial monument type. So far, they are only found in two areas, Ephyra and Pogoni [38]. The construction of tumuli, is linked with symbolic acts that promote and maintain the collective identity and continuity of preceding communities. [39].

In other cases, graves were often arranged in clusters, while in some cases the graves were arranged around a particular burial. The Amvrakia cemeteries do not appear to follow the conservatism of the burial customs of Epirus, which exhibits a lack of innovations. The simple grave types in the Epirus hinterland and the absence of grave markers are in contrast to the cemeteries of Amvrakia. It is therefore obvious that Amvrakia adopted the burial customs of southern Greece, such as those of Corinth and Athens. The Amvrakia burial customs of the 6th century BCE follow those of the metropolis. Yet, during the early colonial stages of Amvrakia it appears that burial customs were still different from those of the metropolis [40].

The eastern cemetery of the city was smaller in size than the western one. The plots that have been investigated in the eastern cemetery are scattered. Therefore we do not have the comprehensive picture we do have for the western cemetery. The eastern cemetery did not have the monumentality that the western cemetery had acquired, at least during the Late Classical period. It had a 6m wide burial street with burial enclosures on both sides and a paved pavement along the eastern side [41]. The archaeological research from the beginning of the 20th century to the present day has revealed a large section of the Amvrakia eastern cemetery, with burials dating from the Late Archaic, Classical, and Hellenistic periods. This proves the continuous use and functionality of the burial site. Most graves are cist-shaped, followed by the pit graves, and the small cist-cases. The absence of graves dating to the late Hellenistic period indicates the progressive abandonment of the eastern cemetery [42].

Particularly important evidence in modern research are the funerary stelae of Amvrakia, one of the largest sets of inscribed monuments we have to date in northwestern Greece. Most of the stelae come from the western cemetery; numerically fewer are those from the eastern cemetery [43].

##### 2.1.1 Graves associated with the deep-sequenced individuals of the present study

**Archaic period**

**-Grave CVII**: The limestone cist-grave CVII, oriented north-south, was located at southwestern cemetery of Amvrakia (Theodorou plot) and it was excavated on 19/08/1997 in trench D2. The grave is of small dimensions and measures approximately: 2.00 × 0.65 × 0.64 m. The floor of the cist tomb is paved with gravel and mud. The opening was covered by a large limestone slab. The grave was reused. It contained a primary (orientation N-S; Anthro ID 5) and a secondary burial (Anthro ID 781), each of one individual. The primary burial was an adult individual (18-50 years old) with undetermined sex (after macroscopic estimation). The secondary burial (earlier remains) was pushed to the sides of the cist when the grave was reopened. No grave goods were found, but a fragmented iron nail was discovered. There is no direct dating, as the chronology is mostly based upon ceramics. The burials were dated to the late Archaic period, second half of the 6th century BCE (ca. 550-500 BCE).
Deep-sequenced individual: Individual A (primary burial), **Amv_Epi_Arch_1**.

**-Grave LXIX**: Pithos (storage vessel) subadult (infant) burial (**Supplementary Figure S3**). The pithos was located in the burial enclosure γ, southwestern cemetery of Amvrakia (Kommenos plot) and it was excavated on 04/10/2011. The pottery vessel is undecorated and was found in a vertical position at a depth of 12.62 m from the surface. The pithos contained a single inhumation of an infant (Anthro ID 267). The pithos was fragmented. Four ceramic pots were found in situ. Three of them were Corinthian imports and the kyathos was probably made by an Amvrakian workshop [40]. The burial was dated to the late Archaic period, third quarter of the 6th century BCE (ca. 550-525 BCE). It is worth mentioning that the burial of adults in pithoi did not occur in the burial customs of Corinth. This seems to be a practice of the local population before colonization. Lefkada and Corfu, also Corinthian colonies, buried their dead in pithoi as a common practice in northwestern Greece in pre-colonial times.
Deep-sequenced individual: **Amv_Epi_Arch_1**.

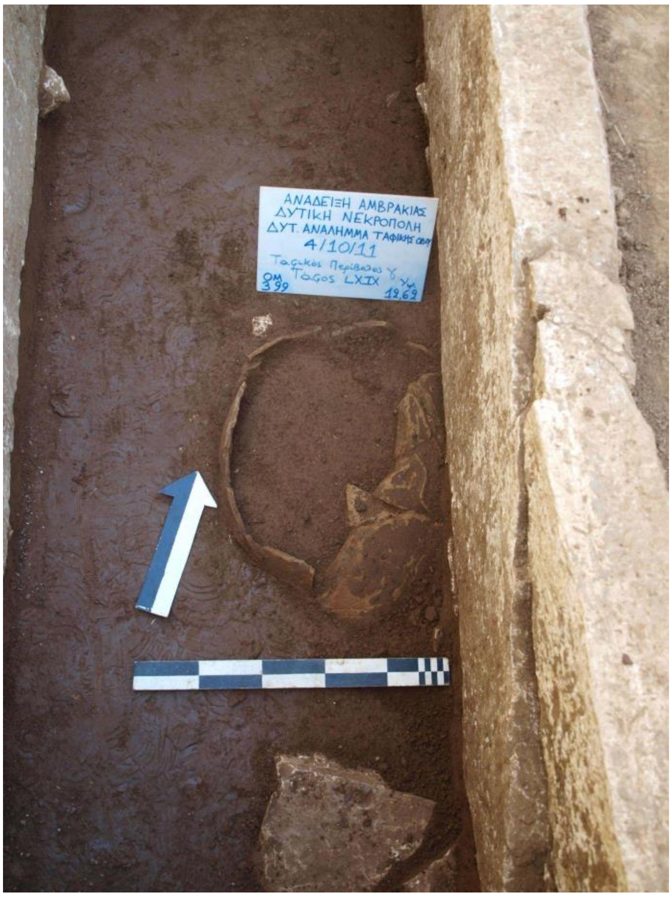


**Supplementary Figure S3.** Grave LXIX, Western Necropolis of Amvrakia.

-**Grave CXXVI**: The limestone cist-grave CXXVI (**Supplementary Figure S4**), oriented S-E and N-W, was located in burial enclosure E, at the southwestern cemetery of Amvrakia (Kommenos plot) and it was excavated on 25/10/2012 at a depth of 11.84 m from the surface. The grave contained two burials (I: primary - II: secondary). A burial of a 35 year old male (Anthro ID 1) and a burial of a 35-45 years old female (Anthro ID 51), were identified by the excavation team [40]. The floor of the cist tomb was paved with gravel and mud. One individual (I) was accompanied by a black-figured flask located above the right shoulder. Additionally, two bronze 'scrapers' (stleggides) for cleaning the dust and the remaining oil from their skin after training and a fragmented bronze end of a musical instrument (all offerings for the man) were found. There is no direct dating, as the chronology is based mostly upon ceramics. The burials were dated to the late Archaic period ca. 500-480 BCE.
Deep-sequenced individual: Individual I (burial I), **Amv_Epi_Arch_3**.

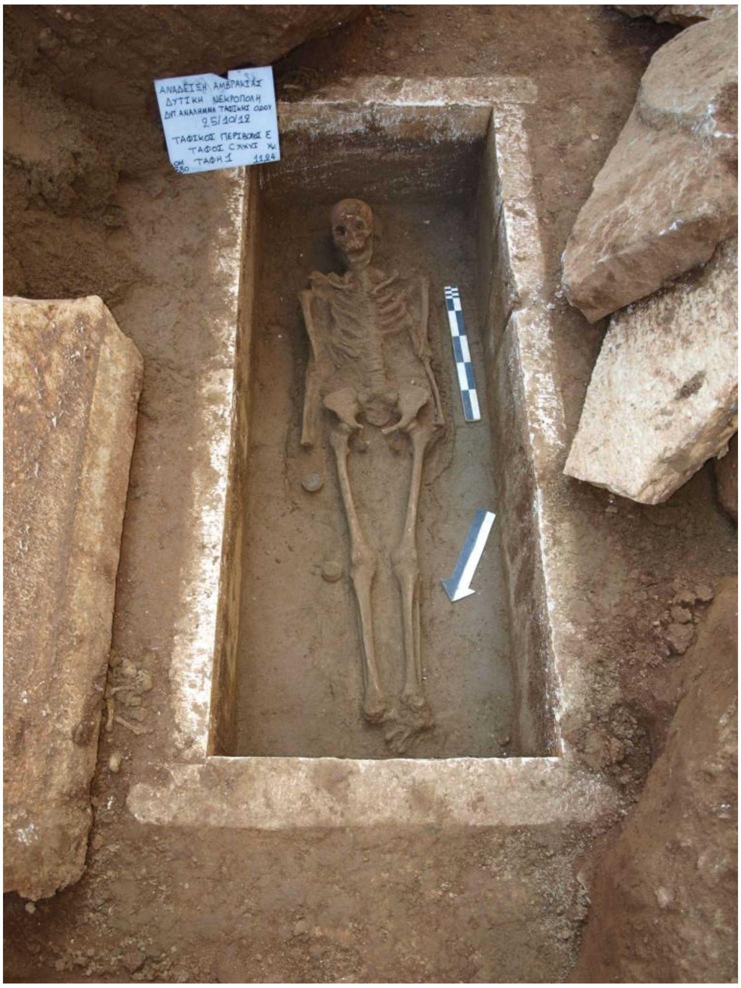


**Supplementary Figure S4.** Grave CXXVI (primary), Western Necropolis of Amvrakia.

#### Classical period

-**Grave CXIX**: The limestone cist-grave CXIX was located in burial enclosure E, at the southwestern cemetery of Amvrakia (Kommenos plot) and it was excavated on 25/09/2012, at a depth of 12.49 m from the surface. A fragmented limestone slab was found *in situ*, probably covering the grave opening. The grave was reused. It contained a primary (orientation S-N; Anthro ID 7) and a secondary burial (**Supplementary Figure S5;** Anthro ID 8), each of one individual. The earlier remains were pushed to the north side of the cist when the grave was reopened. One individual was accompanied by a kyathos (drinking-cup for wine) and two small lekythoi and a bronze ring. According to A. Aggeli, a burial of a 45-49 years old woman was found in situ. A secondary burial of a female individual over 60 years of age, with no grave goods, was also recognised [40]. There is no direct dating, as the chronology is based mostly upon ceramics. The burials were dated to the late Classical period, second quarter of the 4th century BCE (ca. 375-350 BCE).
Deep-sequenced individual: Retrieval individual (=secondary burial), **Amv_Epi_Cl_1**.

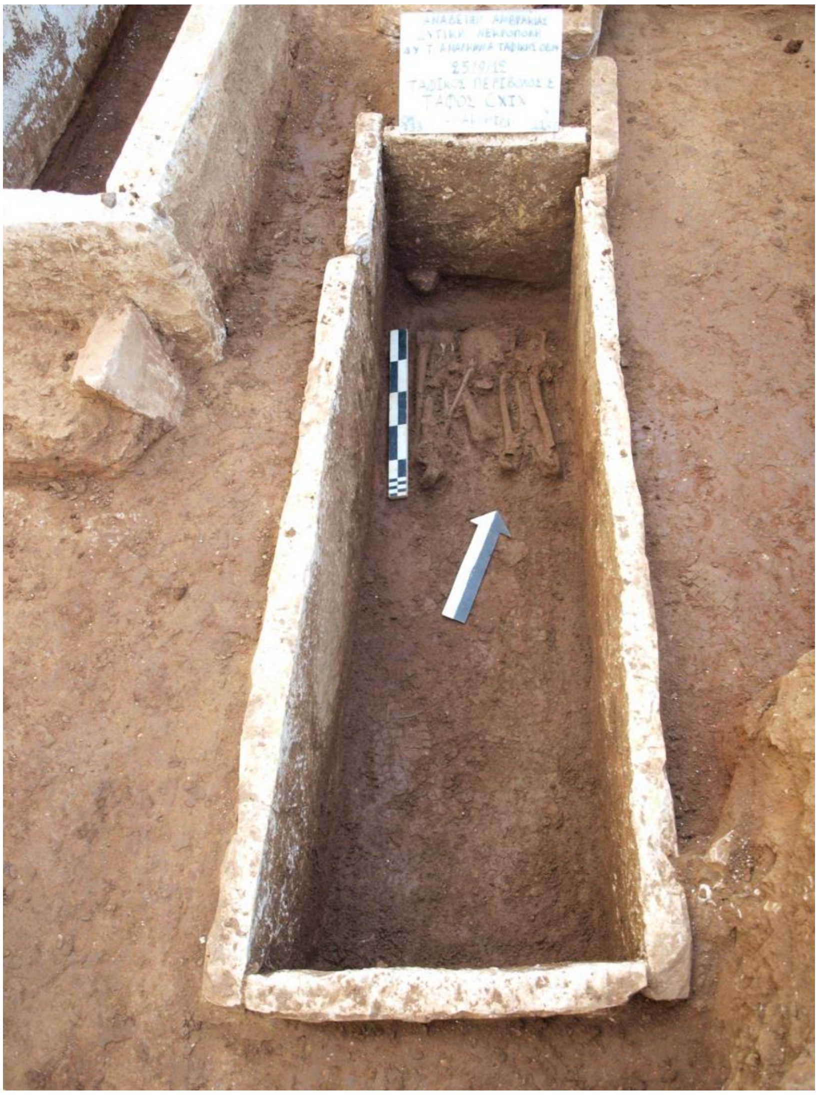


**Supplementary Figure S5.** Grave CXIX (retrieval), Western Necropolis of Amvrakia.

-**Grave CCCXXXIX**: The limestone cist-grave CCCXXXIX (**Supplementary Figure S6**), oriented north-south, was located at the southwestern cemetery of Amvrakia (Kommenos plot) and it was excavated on 30/07/2014 in trench E18 at a depth of 14.76 m from the surface. The opening was covered by a large limestone slab. Grave CCCXXXIX, which had dimensions 1,95 m × 0,67 m, height: 0,53 m is located north of the grave CCCXXXVII. It contained two burials, each of one individual: burial I (Anthro ID 16) and II (Anthro ID 11), both with a S-N orientation. According to A. Aggeli, the main burial belonged to a female, with a poor state of skeletal preservation. The head was oriented to the south, probable age of 50 years (after macroscopic estimation). A canastron was placed at both sides of the body at the level of the neck. Underneath, of the female burial, a male burial of a young man 18-19 years old (after macroscopic estimation), in extended posture, and heading to the south, was discovered. One ceramic vessel, a black figure skyphos (drinking-cup) and two bronze 'scrapers' (stleggides) for cleaning the dust and the remaining oil from their skin after training are associated with the male burial [40]. There is no direct dating, as the chronology is based mostly upon ceramics. The burials were dated to the Classical period, third quarter of the 5th century BCE (ca. 450-425 BCE).

Deep-sequenced individual: Individual II (burial II), **Amv_Epi_Cl_2**.

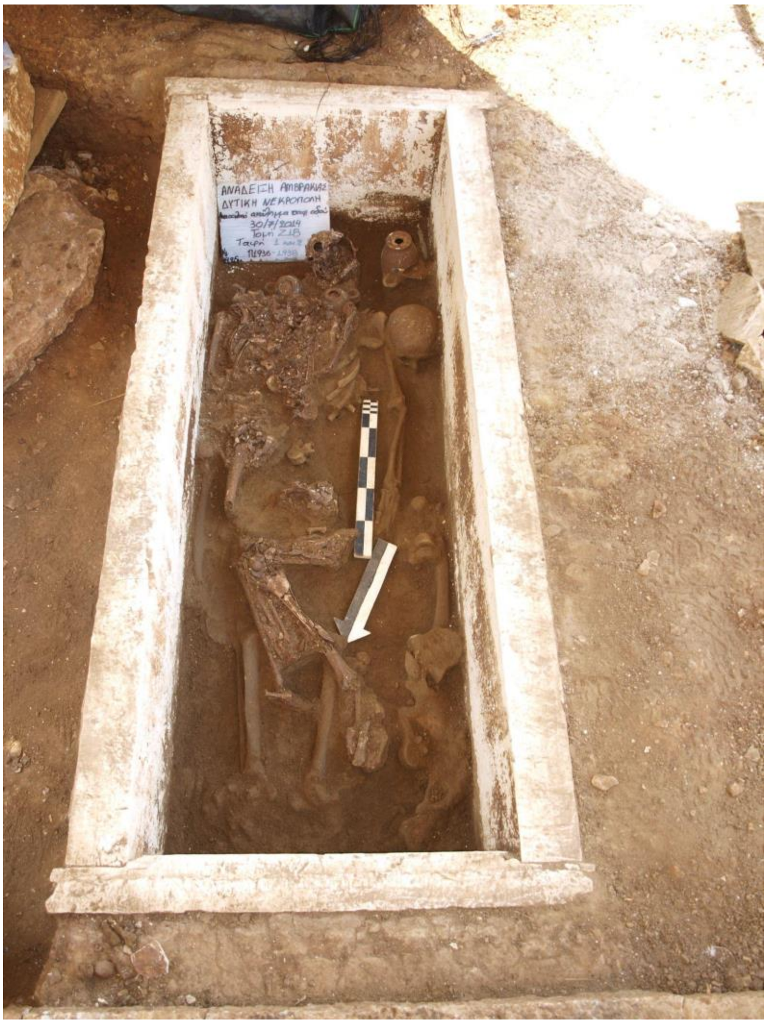


**Supplementary Figure S6.** GraveCCCXXXIX, Western Necropolis of Amvrakia

-**Grave CXLVII**: The limestone cist-grave CXLVII (**Supplementary Figure S7**) was located in the southwestern cemetery of Amvrakia (Kommenos plot) and it was excavated on 17/06/2013 in trench A12 at a depth of 13.25 m from the surface. A limestone slab was found *in situ*. The grave was located and partially constructed below the cist XL. The grave contained one individual (orientation SE-NW; Anthro ID 12). The individual (probably female, after macroscopic estimation) was accompanied by a red figured lekythos (oil container) and a silver coin (not well preserved)[40]. There is no direct dating, as the chronology is based mostly upon ceramics. The burials were dated to the Classical period, first quarter of the 4th century BCE (ca. 400-370 BCE).
Deep-sequenced individual: **Amv_Epi_Cl_3**.

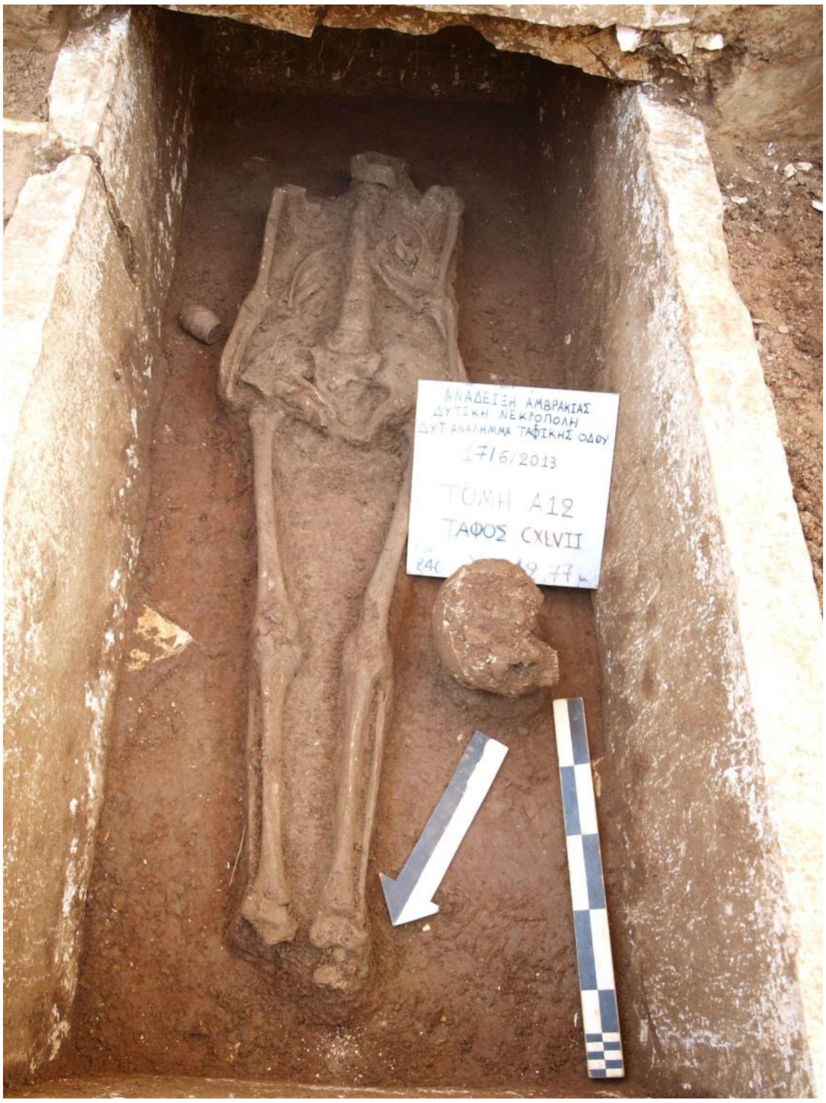


**Supplementary Figure S7.** Grave CXLVII, Western Necropolis of Amvrakia.

-**Grave CCCLXXXVIII**: The pit-grave CCCLXXXVIII (**Supplementary Figure S8**) was located at the southwestern cemetery of Amvrakia (Kommenos plot) and it was excavated on 28/07/2015 in trench E21 at a depth of 15.28 m from the surface. The grave was located and partially constructed below the burial CCCLXXXVI. The grave contained two individual burials. Both primary (Anthro ID 14) and secondary (Anthro ID 13) burials had orientation N-S. The individuals were accompanied by a small lekythos (oil container) [40]. There is no direct dating, as the chronology is based mostly upon ceramics. The burials were dated to the Classical period, second quarter of the 5th century BCE (ca. 475-450 BCE).
Deep-sequenced individual: Individual II (burial II), **Amv_Epi_Cl_4**.

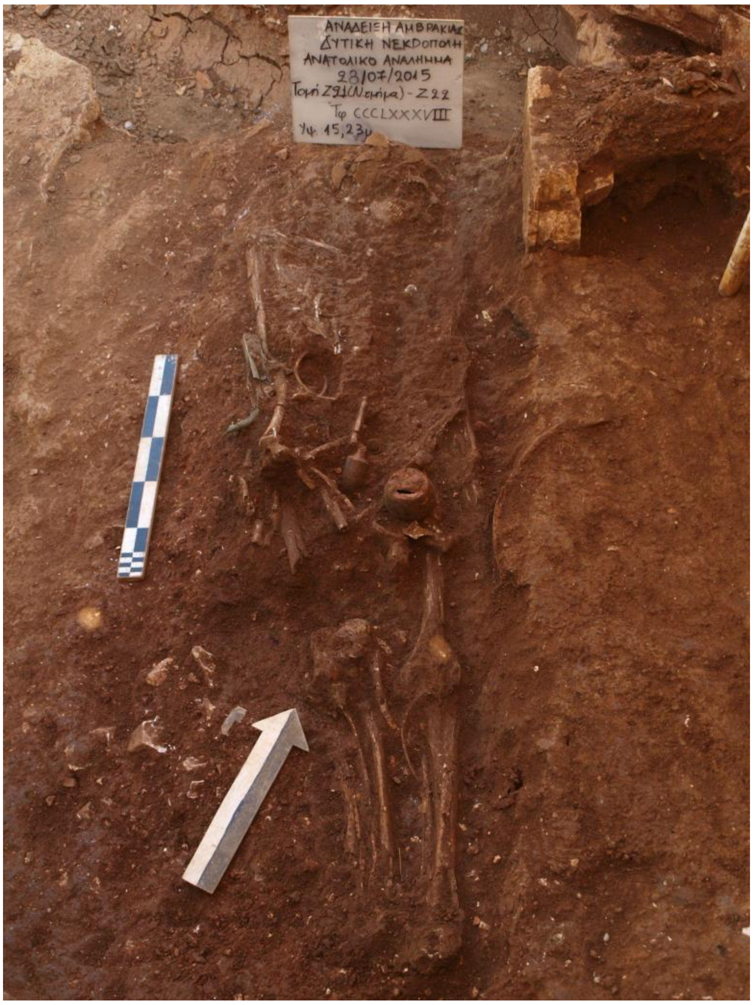


**Supplementary Figure S8.** Grave CCCLXXXVIII, Western Necropolis of Amvrakia.

-**Grave CV**: The pit-grave CV (**Supplementary Figure S9**) was located in burial enclosure ε, at the southwestern cemetery of Amvrakia (Kommenos plot) and it was excavated on 30/03/2012 at a depth of 13.10 m from the surface. A limestone slab was found *in situ*. The grave contained distracted secondary burials of two subadults represented by two temporal bones (bonedata_30032012_cv1 and bonedata_30032021_cv2). The estimated age of the subadults was 4 and 4-18 years old (after macroscopic estimation), respectively. One ceramic vessel, a black-figured skyphos (drinking-cup), is associated with the burials. A clay rattle and a clay doll figurine were also found in situ [40]. The chronology is based upon ceramics. The burials were dated to the late Classical period, second quarter of the 4th century BCE (ca. 375-350 BCE).

Deep-sequenced individuals: Retrieval individual I (=secondary burial I), **Amv_Epi_Cl_5** and retrieval individual II (=secondary burial II), **Amv_Epi_Cl_6**.

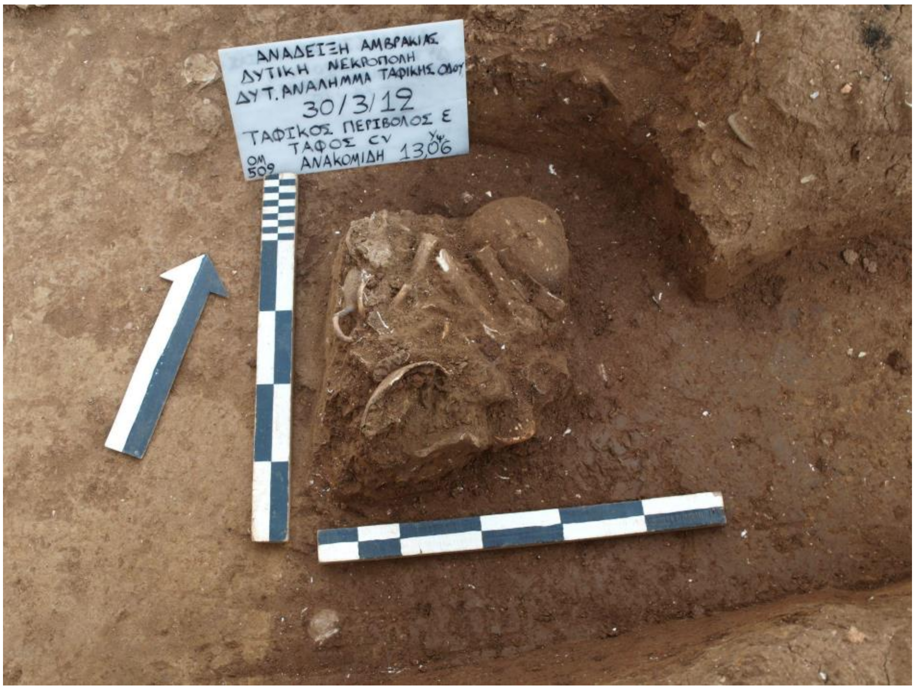


**Supplementary Figure S9.** Grave CV, Western Necropolis of Amvrakia.

**Hellenistic period**

-**Grave CCVIII**: The build limestone cist-grave CCVIII (**Supplementary Figure S10**), oriented E-W, was located at the southwestern cemetery of Amvrakia (Kommenou plot) and it was excavated on 19/07/2012 in trench E14. It was excavated at a depth of 15.90 m from the surface. A pile of stones was found *in situ*. The grave was reused. According to the excavation log book the grave contained two primary (orientation E-W; Anthro ID 211-212) and three secondary burials (Anthro ID 214-216). The earlier remains were pushed to the side of the cist when the grave was reopened. One ceramic vessel, a skyphos (drinking-cup), is associated with one of the burials. There is no direct dating, as the chronology is based mostly upon ceramics. The burials were dated to the Hellenistic period, 1st-3rd quarter of 2nd century BCE (ca. 200-125 BCE).
Deep-sequenced individual: primary burial individual Anthro ID 212, **Amv_Epi_Hel_1**.

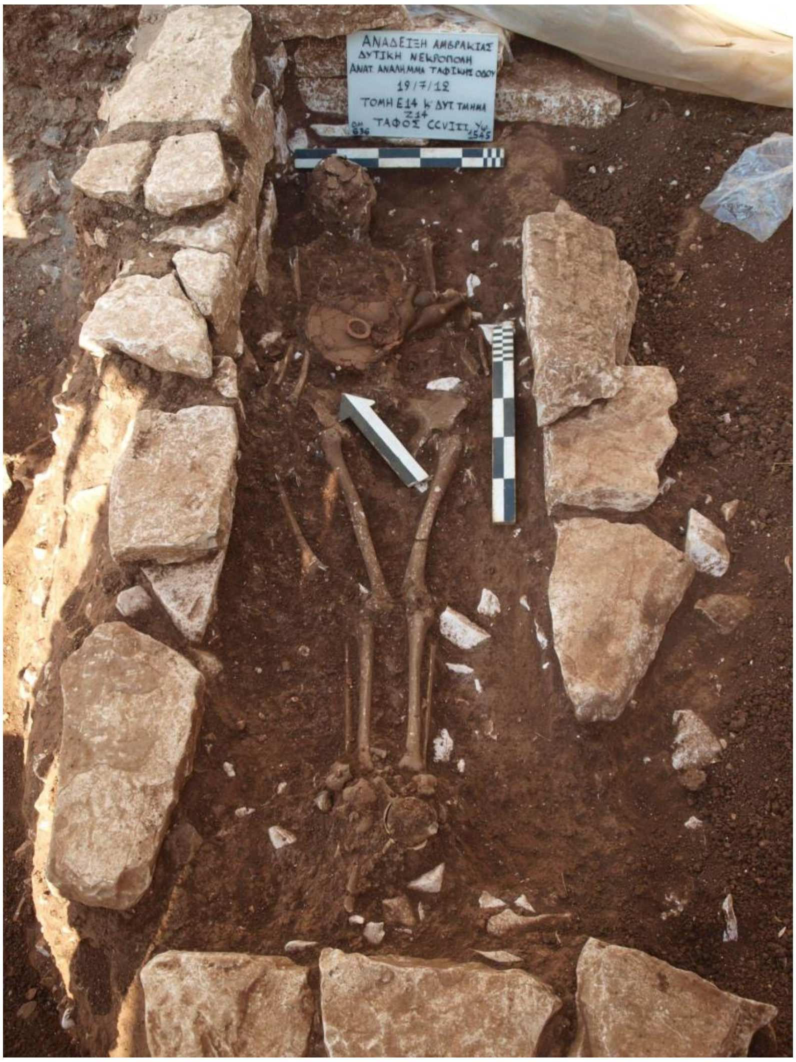


**Supplementary Figure S10.** Grave CCVIII, Western Necropolis of Amvrakia.

-**Grave CXCIV**: The pit-grave CXCIV (**Supplementary Figure S11**), oriented E-W, was located at the southwestern cemetery of Amvrakia (Kommenou plot) and it was excavated on 26/06/2012 in trench Section E12 and Benchmark E11 - E12, and Section E13 (North of Wall 103), at a depth of 14.90 m from the surface. The grave contained one primary burial (orientation N-S; Anthro ID 656). An unidentified Hellenistic bronze coin found *in situ*, is associated with the burial. There is no direct dating, as the chronology is based mostly upon ceramics. The burial was dated to the Hellenistic period, 2nd half of 3rd century BCE (ca. 250-200 BCE).
Deep-sequenced individual: **Amv_Epi_Hel_2**.

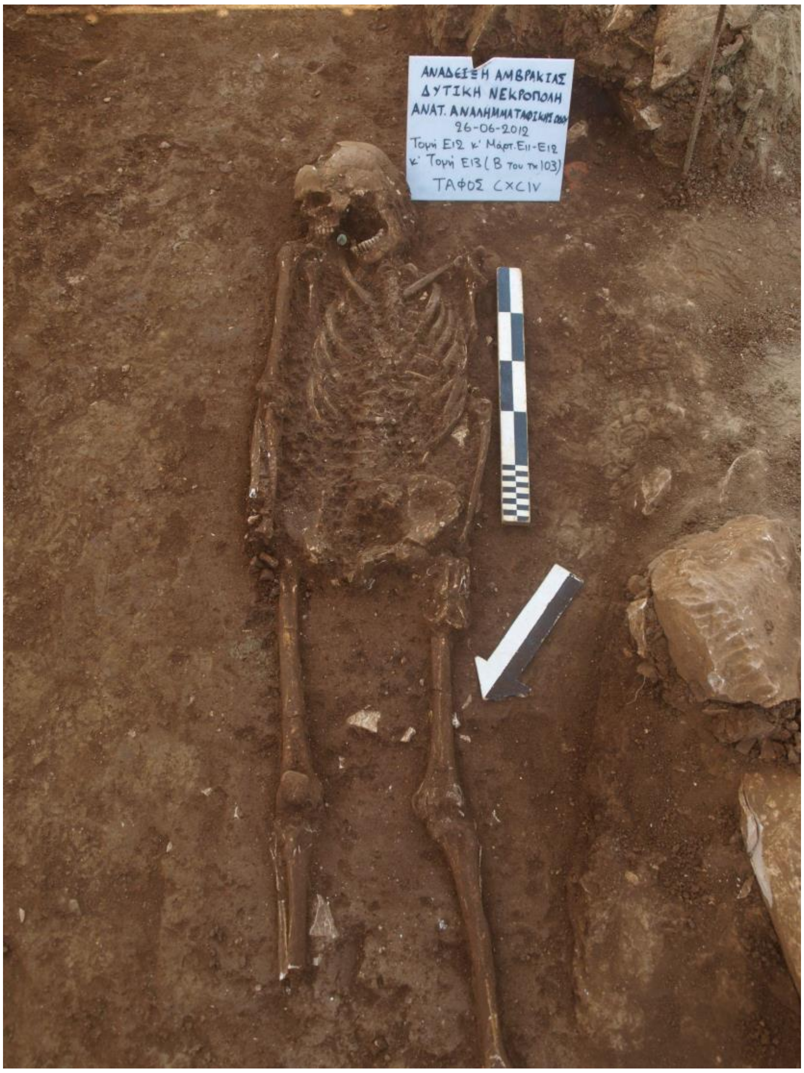


**Supplementary Figure S11.** Grave CXCIV, Western Necropolis of Amvrakia.

-**Grave CCXLV**: The pit-grave CCXLV (**Supplementary Figure S12**), oriented S-N, was located at the southwestern cemetery of Amvrakia (Kommenou plot). It was excavated on 05/10/2012 at a depth of 14.52 m from the surface. The grave contained one primary burial (orientation S-N; Anthro ID 389) and one secondary burial (Anthro ID 390). A golden “danake” (coin), served as the so-called Charon's obol, was placed in the individual's mouth. It was found *in situ* among the individual’s teeth. There is no direct dating, as the chronology is based mostly upon ceramics. The burials were dated to the Hellenistic period, 2nd-3rd quarter of 2nd century BCE (ca. 175-125 BCE).

Deep-sequenced individuals: Retrieval individual (=secondary burial), **Amv_Epi_Hel_3** and primary burial individual, **Amv_Epi_Hel_4**.

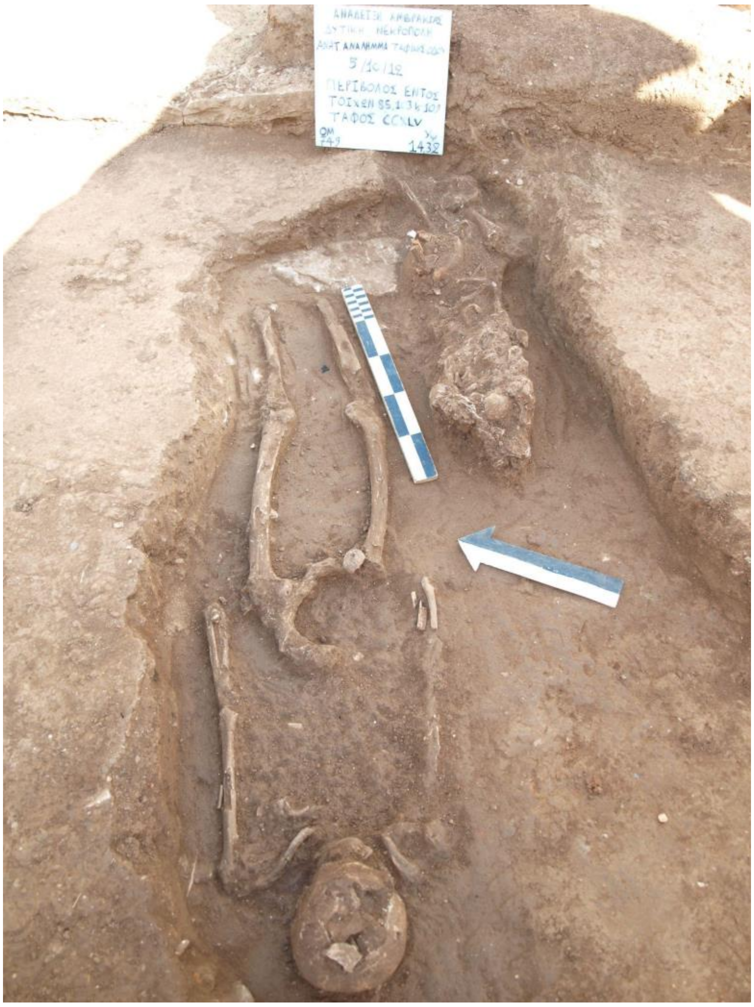


**Supplementary Figure S12.** Grave CCXLV, Western Necropolis of Amvrakia.

-**Grave CCXXIII**: The limestone cist-grave CCXXIII (**Supplementary Figure S13**), oriented N-S, was located at the southwestern cemetery of Amvrakia (Kommenos plot) and it was excavated on 30/08/2012, in the precinct within the walls 105, 107, and 110, at a depth of 14.62 m from the surface. Broken limestone slabs were found *in situ*. The grave contained a total of five burials; three primary burials (Anthro ID 438 - 440), two secondary burials (Anthro ID 441-442), and one cremation burial (bonedata_30082012_CCXXIII) found in an amphora. The cremation amphora was found in the west side of the tomb. It held a lid made of lead (M89/OM697). The burials were accompanied by a terracotta spindle-shaped unguentarium (perfume bottle). There is no direct dating, as the chronology is based mostly upon ceramics. The burials were dated to the Hellenistic period. Due to absence of associated grave finds with the secondary burials, the dating is based on the usage period of the grave; late 4th to the 2nd century BCE (ca. 325-100 BCE).
Deep-sequenced individual: Retrieval individual (=secondary burial) Anthro ID 441, **Amv_Epi_Hel_5**.

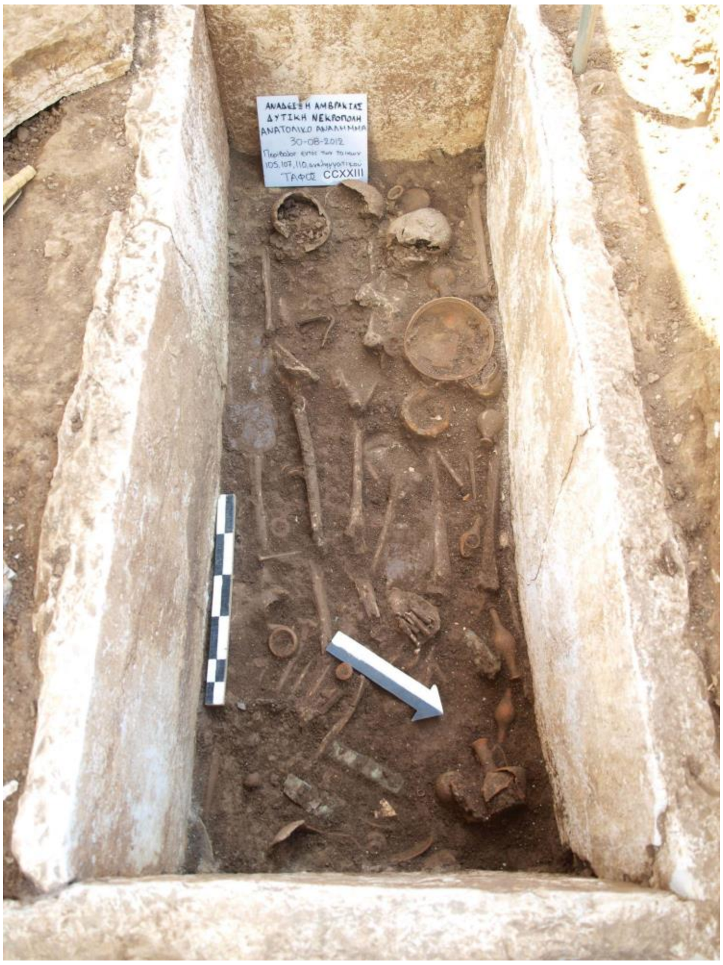


**Supplementary Figure S13.** Grave CCXXIII, Western Necropolis of Amvrakia.

**Undated burial**

-**Grave CCXLIII**: The limestone cist-grave CCXLIII (**Supplementary Figure S14**), oriented E-W, was located at the southwestern cemetery of Amvrakia (Kommenos plot) and it was excavated on 04/10/2012, in the perivolos within walls 85, 103, and 109, at a depth of 14.54 m from the surface. A limestone slab was found *in situ*. The grave contained two burials: one primary burial (orientation N-S; Anthro ID 661) and one secondary burial, as well as several isolated teeth (bonedata_04102012_CCXLIII). A fragmented iron stleggis (body scraper) was found *in situ*. Neither direct, nor indirect (e.g. ceramics-based) dating is available. Due to absence of -chronology-indicative- associated grave finds with these burials, their dating is based on the usage period of the necropolis; Archaic to Roman period (ca. 700 BCE - 476 CE).

Deep-sequenced individual: bonedata_04102012_CCXLIII, **Amv_Epi_Archaic_to_Roman**.

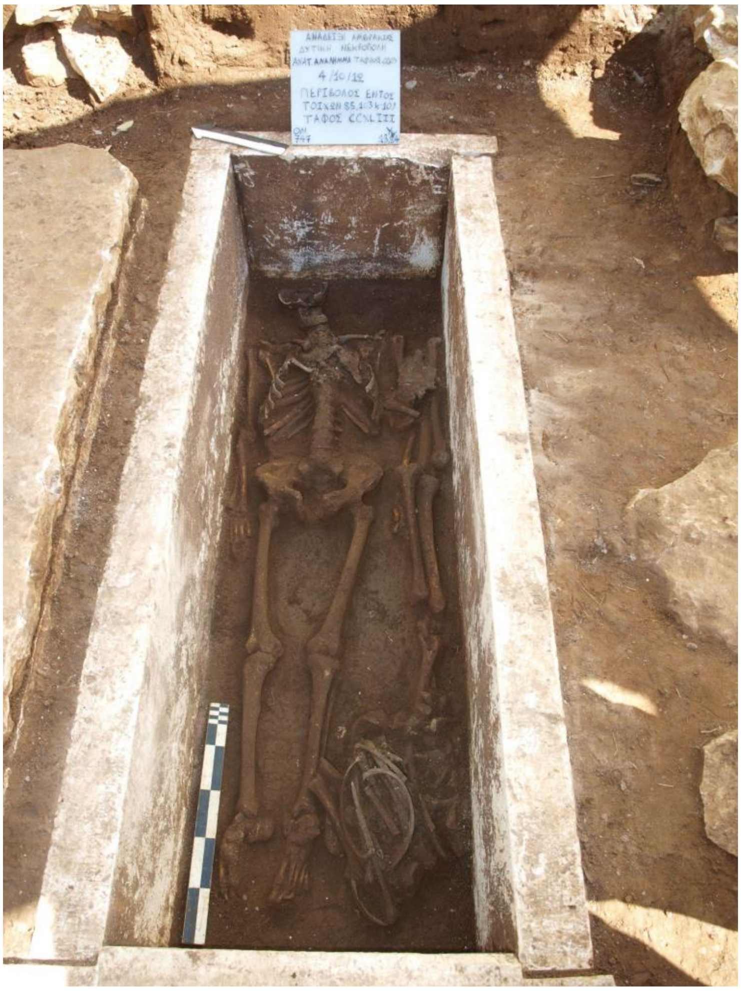

**Supplementary Figure S14.** Grave CCXLIII, Western Necropolis of Amvrakia.

#### 2.2. Ammotopos - “Kokkino Lithari” site

A prehistoric settlement of great importance was discovered (2015 - 2016) during construction work for the Ionian highway connecting western Greece to Athens. The site is situated at a crossroad of people and goods for the topography of the area, connecting Athamania, Molossida, and the Amvrakian plain. Remains of dry masonry constructions with finds of pottery, (big storage vessels), numerous stone tools etc. are dated to the Middle and Late Bronze Age (c. 2000-1000 BCE). The new archaeological site, named “Kokkino Lithari", is located on the steep side of the hill directly opposite of House A of Orraos. The hill of "Kokkino Lithari" is a site where Neolithic structures and stone tools have been found. One of the most interesting finds was a grave located on the southern part of the hill, with walls made of limestone slabs. The tomb contained four burials (**Supplementary Figure S15**), an unglazed drinking vessel (kantharos type of local production), and a bronze ring. Another grave located closeby, at the Kastri hill, was uncovered. The Kastri tomb was dated to the Late Helladic (LH IIIA2- IIIB) period (ca. 1350-1200 BCE). This tomb contained five burials, a bronze knife, a necklace of stone beads, fragmented pottery (which were later restored and reconstructed to a Mycenaean jar and a Mycenaean global alabastron), and four Mycenaean kylikes from unstratified layers. The pottery clearly indicates the influence of the Mycenaean workshops in the area [26,44–46].

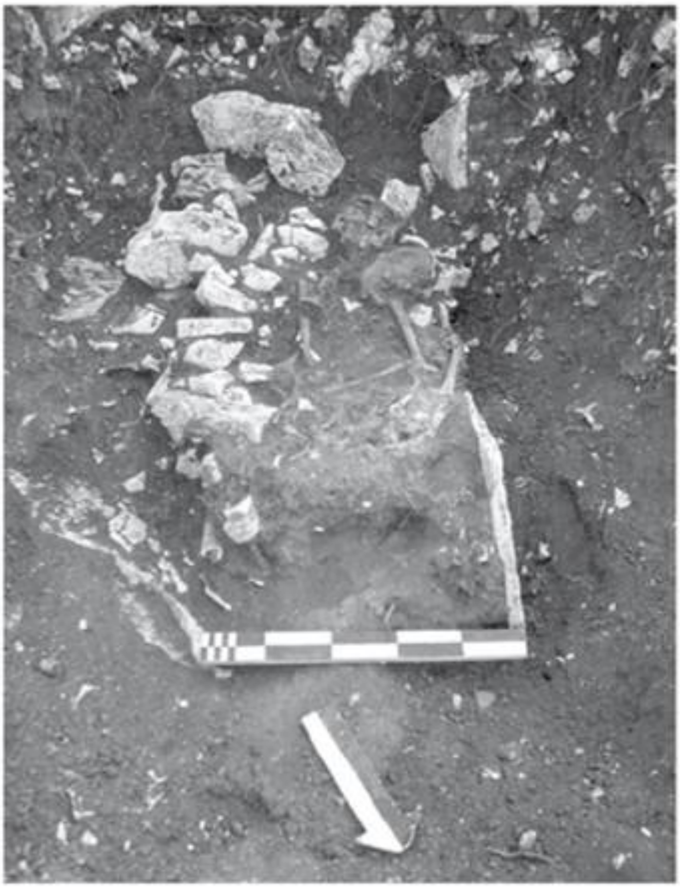

**Supplementary Figure S15.** Grave I, Kokkino Lithari site in Ammotopos.

**Sampled Individuals**

-**Grave I**: The grave was built with limestone slabs. It was located in the southern part of the "Kokkino Lithari" hill (IONIA Highway, Χ.Θ. 153+975). It was excavated on 26 & 27-05-2015, and contained four skeletons preserved from the femur and above. Two of them were oriented S-N (Anthro ID 17, Anthro ID 18), a third one in a N-S orientation, whereas the skeletal remains of a fourth burial were pushed toward the margins of the grave. No particular care was taken during the burial to ensure the position and orientation of the deceased. The grave goods were a bronze ring and a ceramic kantharos type vessel (drinking vessel) of probable local origin [46]. The finds date the burials to the Late Bronze Age (c.1350 - c.1125 BCE), and $C^{14}$ dating of individuals I and II narrows it down to 1275-1125 calBCE [3225-3075 calBP (95.4%)].
Deep-sequenced individuals: Individual I (**Amm_Epi_LBA_1**) and individual II (**Amm_Epi_LBA_2**).

#### 2.3. Cemeteries of Tenea

Since 2013 until today, ancient Tenea has been systematically excavated in the context of the "Tenea Project" that is supported by an interdisciplinary team. Important remains of the ancient city have been uncovered, which relate to its public and private life. The excavation of a large part of its cemeteries yielded important information concerning the living conditions of its inhabitants, as well as the broader image of the city in terms of its organization and layout, customs and practices.

Initially, in 1984, the archaeological service brought to light the archaic porous sarcophagus of Chiliomodi, which is now exhibited in the Archaeological Museum of Ancient Corinth [47]. Inside the sarcophagus, a female burial aged 18–28 years was preserved in an outstretched position, oriented SW/NE. The burial was enriched with, among other things, a metal and wooden pin, a bronze mirror, a pyxis - kalathos, and a handless decorated pyxis with curved walls, which are representative examples of Corinthian art during the early 6th century BCE [48]. The uniqueness of the find, however, lies in the covering slab of the sarcophagus, which had a painted representation on its inner side in ad secco technique. Two lions heraldically placed on an off-white background are depicted with an antefix in between in the form of a palmate. The image is framed by a red band and is definitely related to the monumental painting of the Archaic period [47,48].

Around the sarcophagus of 1984, a cluster of Archaic graves was discovered in the years 2013-2015 (**Supplementary Figure S16**), which yielded a substantial number of grave goods, many of which introduce new typological shapes in the relevant bibliography [49]. More specifically, four burials in porous monolithic sarcophagoi, which date from the beginning of the 6th to the beginning of the 5th century BC, have come to light. These are burials of a child (Grave 02), a woman about 50 years old (Grave 03), and two men (Grave 04 and 05), aged about 40-50 years and over 50 years old. The burials were enriched with ceramic and metal objects that were located, either inside, or around the graves. The graves were not uniformly oriented. The child's burial (Grave 02) stands out, around and within which 58 ceramic, metal and lead objects that accompanied the burial were found. Typologically, the findings consist of oenochoes, aryballoi, pyxides, as well as a lekanis, a hydriske, a lekythos and two bronze lekanides that date to the Corinthian art of the early 6th century BCE. Remarkable finds are a double askos with a rope handle and trefoil strained spout, as well as two small-sized handless vases, for which no parallel finds have been identified so far [36]. The other graves had undisturbed burials in an extended position-oriented SW/NE. Graves 03 and 04 contained only one burial each, and date to the mid and late 6th century BCE. Grave 05 dates to the beginning of the 5th century BCE and contained six vases, two kylikes, two lekythoi, a skyphos. and an oenochoe. In the proximity of the grave a modern deposit was found, in which two lekanides with lids, two kylikes, two skyphoi, and an oenochoe were found [36].

At a short distance from the archaic cluster, a Hellenistic period pit grave was found, which housed the remains of two adults. The burials were adorned with a krateriskos, a miniature oinochoe, a miniature cup, and a lamp, which can be dated through the four coins minted in the reign of Ptolemy III that were also placed as grave goods.

Hellenistic burials were also found at the southwestern border of the cemetery zone, at the "Palaio Scholeio" area (**Supplementary Figure S19**). The burials were organized in the area north and east of a semi-underground cistern, violating in some cases earlier structures for their encapsulation. The excavation revealed a total of ten graves of the Hellenistic period, the majority of which were reused in Roman times. The number and type of grave goods found inside the undisturbed graves is impressive, with vases representative of the Hellenistic period, such as miniature vases and unguentaria, but also more refined shapes, such as the lagynos and the funerary calpe found in Grave 17. Also noteworthy are the metal objects that came to light, such as bronze mirrors, iron strigils, and a bronze oinochoe. A gold-plated bronze wreath of myrtle leaves and fruits, a gold ring, and other metal jewellery are included, as well as gold danakes.

The graves belong to the types of pit graves, monolithic sarcophagoi, but also stone cofins with porous covering slabs. The reuse of the graves during the Roman period resulted in the identification of sidelined anthropological remains on the outside of the graves and the coexistence of Hellenistic and Roman finds. The macroscopic study of the anthropological remains found inside and outside the graves played a key role in the separation of the main and secondary burials.

The Roman burials identified as having reused Hellenistic graves and others in upper excavation layers in the same area, belong to a well-organized Roman cemetery around the perimeter of an above-ground funerary monument discovered in 2016 (**Supplementary Figure S22**). The monument is a two-room above-ground and temple-shaped funerary monument of the 1st - 2nd century CE. with dimensions 10.53 × 5.82 m, orientation E/W, and entrance to the West. In the burial chamber, five built cist graves are formed with dimensions of 2.00 × 0.68 m each in a circumferential layout that has the shape of the Greek letter “Π”. Monuments of similar typology, roofed with vaults, can be found in Patras, Argos, Nicopolis, Ostia, and Asia Minor [35][36]. The funerary monument of Tenea is one of the few above-ground burial monuments found so far in the Corinthia regional unit of present-day Greece, yet without there being another of similar typology.

Around the monument, an organized cemetery of Roman times was revealed, dating from the 3rd century CE, up to the 5th century CE. The graves are distinguished according to their typology into pit graves, kalyvites pit graves, and jar burials. The burials were found richly adorned with lamps, glass, gold, silver, lead, bronze and iron jewellery, vases of everyday use, glass vessels, metal tools, bone jewellery and tools, organic remains, coins, and shoe nails [36,49].

As all the above evidence suggests, the "Palaio Scholeio" area and specifically the location of the funerary monument, was a place with long burial use and ritual value in antiquity, near the ancient city and the residential web.

##### 2.3.1 Graves associated with the deep-sequenced individuals of the present study

**Archaic period (Faneromeni-Kamareta site)**

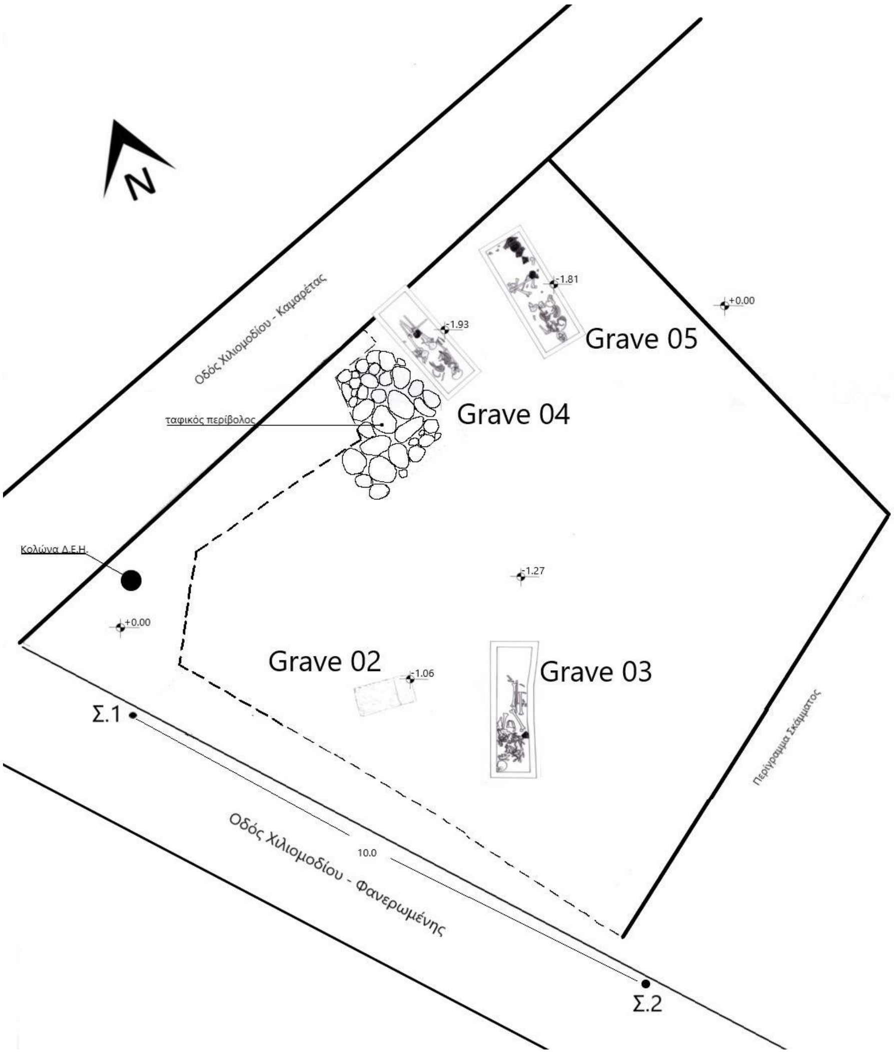


**Supplementary Figure S16.** The location of graves 3 and 5 in the Faneromeni-Kamareta site of Ancient Tenea.

-**Grave 03**: The porous sarcophagus (**Supplementary Figure S17**), oriented N-S, was excavated on 08/09/2014, in trench E1-I, in the cemetery of Tenea (Faneromeni-Kamareta, Tsirtsis plot). The sarcophagus carried a porous lid. The dimensions of the sarcophagus are 2.20 × 0.09 × 0.45 m. A layer of mortar covers much of the sarcophagus. The mortar thickness is 1-3.5 cm. The interior of the sarcophagus contained one burial of an adult. It contained one skyphos (drinking-cup) and an iron nail, all found *in situ*. The chronology is based mostly on the ceramics and dates to the late Archaic period (ca. 550-500 BCE).
Deep-sequenced individual: **Ten_Pel_Arch_2**.

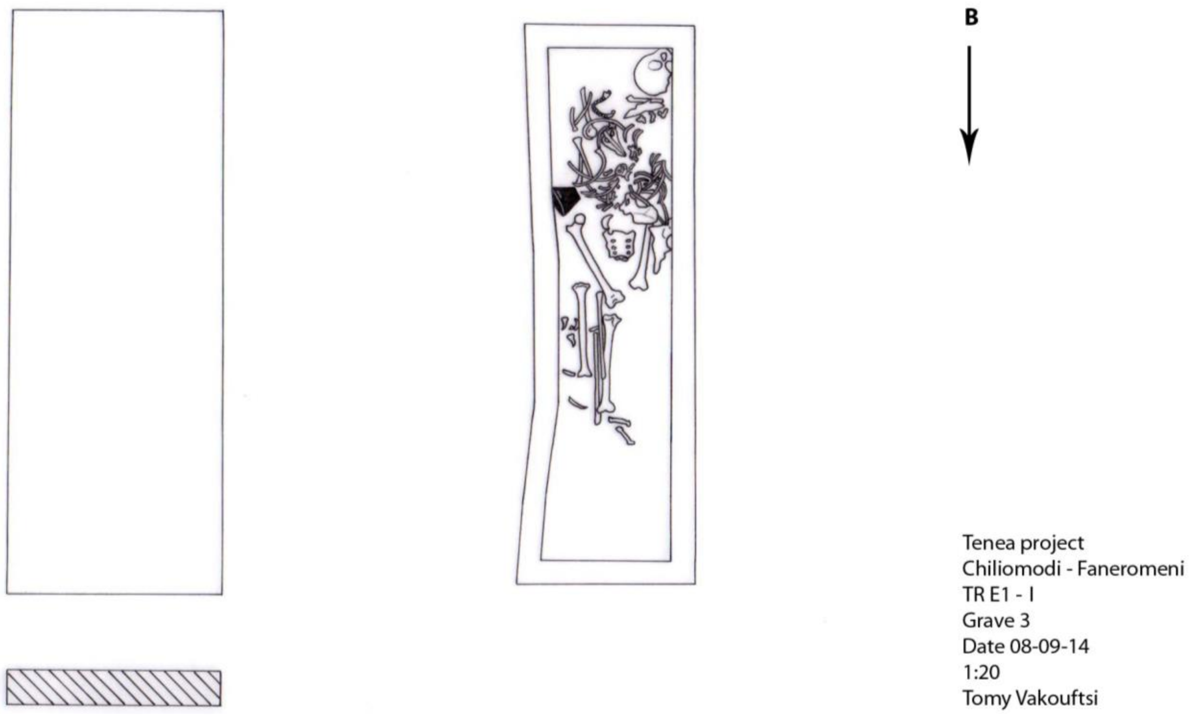


**Supplementary Figure S17.** The sarcophagus of grave 5 in the Faneromeni-Kamareta site of Ancient Tenea.

-**Grave 5**: The porous sarcophagus (**Supplementary Figure S18)**, oriented NE-SW, was excavated on 08/10/2014, in trench E1-I, in the cemetery of Tenea (Faneromeni-Kamareta, Tsirtsis plot). The sarcophagus carried a porous lid. The dimension of the sarcophagus are 0.80 × 0.82 × 2.10 m. A modern layer covers much of the sarcophagus. Its thickness is 1 cm. The interior of the sarcophagus contained one burial of an adult. It contained six vases (among them two kylikes, two lekythoi, one skyphos, and one oinochoe) and one iron ring. The chronology is based on the ceramics and dates to the late Archaic period (ca. 500-480 BCE). Deep-sequenced individual codes: **Ten_Pel_Arch_1**.

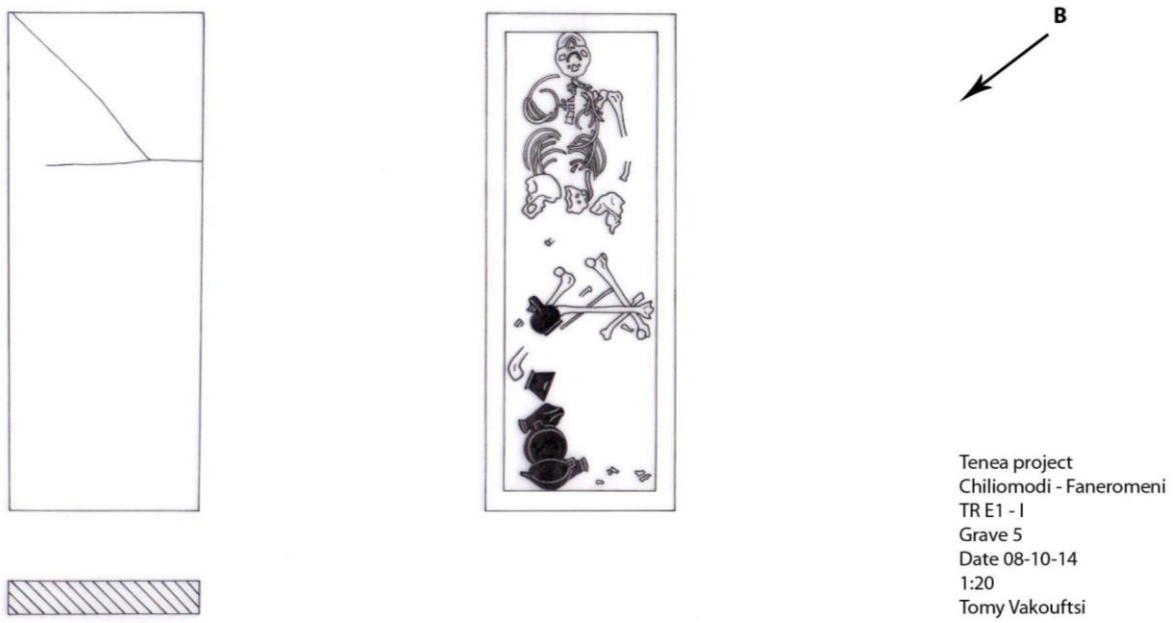


**Supplementary Figure S18.** The sarcophagus of grave 5 in the Faneromeni-Kamareta site of Ancient Tenea.

**Hellenistic and Roman periods (Cemetery of Tenea)**

**Hellenistic period**

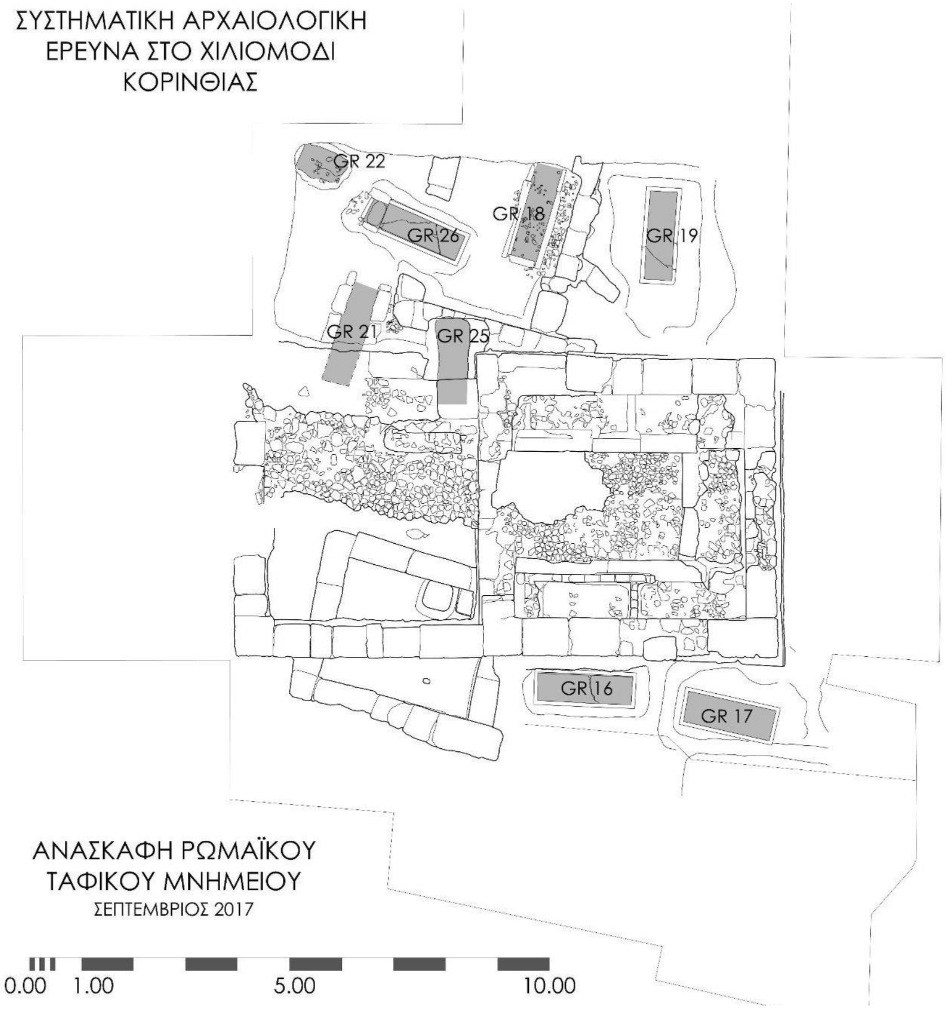


**Supplementary Figure S19.** The Hellenistic graves in the cemetery of Tenea (Palaio Scholeio, Hasikidis plot).

-**Grave 18**: The porous sarcophagus, oriented N-S, was excavated on 24/09/2017 in trench 2017/3, in the cemetery of Tenea (Palaio Scholeio, Hasikidis plot). The grave was located north of a cistern used for ritual purposes, upon which the Roman grave monument (Grave Monument I) was built. Grave 18 contained the burial of one adult. The individual was accompanied by seven fusiform unguentaria, a terracotta lamp, a skyphos, two miniature vases, two pytharia, a gold *Danake*, a ring with semi-precious stone, a folded mirror, a bronze wreath with gilded leaves and myrtle fruits, red-colored pigments, metal fragments, a bone hinge, etc. The chronology is based on the ceramics and dates to the Hellenistic period (ca. 150-100 BCE).
Deep-sequenced individual: **Ten_Pel_Hel_1**.

-**Grave 26 (re-used Hellenistic grave)**: The porous sarcophagus (**Supplementary Figure S20**), oriented E-W, was excavated on 09/10/2017 in trench 2017/2 in the cemetery of Tenea (Palaio Scholeio, Hasikidis plot). The sarcophagus carried a porous lid. The dimensions of the grave are 1.91 × 0.53 m. The interior of the sarcophagus contained three burials of three adults. The coffin itself, as well as one of the burials (Individual 2), dates to the Hellenistic times (323–31 BCE), while the remaining burials were located inside the sarcophagus during the Roman period. The sarcophagus contained various offerings, including a glass unguentarium, six ceramic unguentaria, two ceramic oinochoe, a bronze oinochoe, a skyphos, a glass vessel, a ceramic pedestal, a silver coin, etc. The chronology is based mostly on the ceramics.

Deep-sequenced individual codes: Individual 2, **Ten_Pel_Hel_2**.

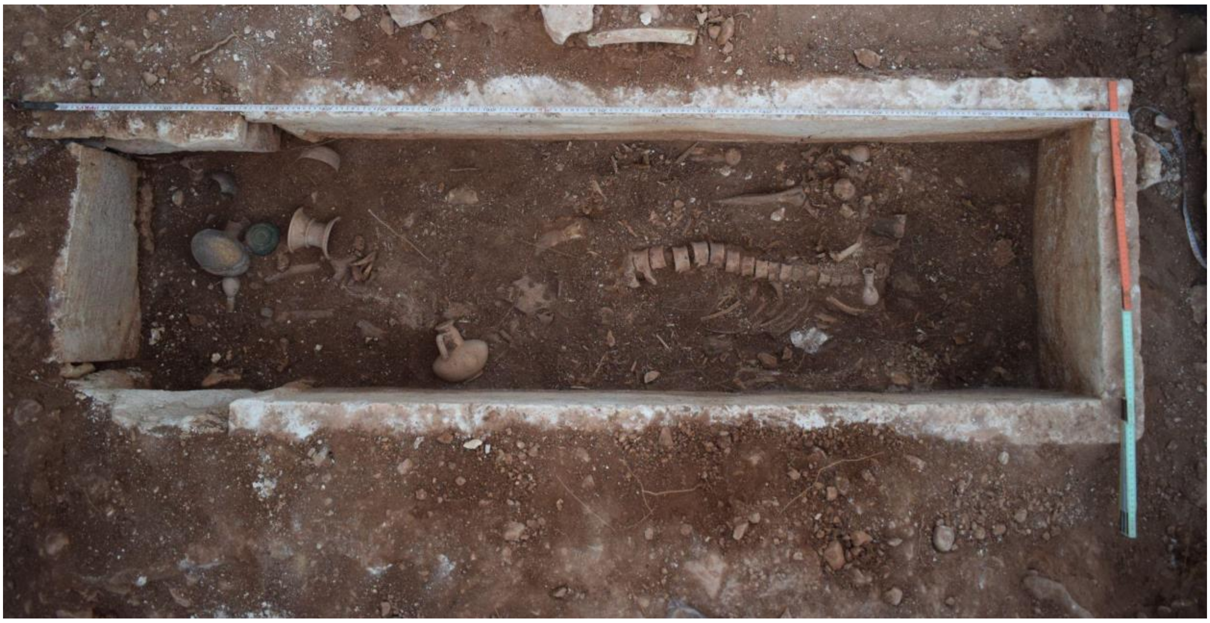

**Supplementary Figure S20.** Grave 26, cemetery of Tenea (Palaio Scholeio, Hasikidis plot).

-**Grave 33 (re-used Hellenistic grave)**: The pit-grave 33 (**Supplementary Figure S21**), oriented W-E, was excavated on 20/09/2018, in trench 2018/3, in the cemetery of Tenea (Palaio Scholeio, Hasikidis plot). The grave was covered with a porous lid, upon which a roman vase was found. The dimensions of the grave are 1.31 × 0.42 m. The grave contained the burial of one adult. The individual was accompanied by a fragmented Hellenistic unguentarium (perfume bottle). From the evidence of the burial assemblage, it is not possible to determine whether the skeletal remains belong to a burial contemporary with the Hellenistic vase found inside the pit or contemporary with the Roman vase found on top of the covering lid. In either case it appears that the grave was disturbed during the Roman period either for reuse or because of the construction (foundation) of the Roman wall found parallel to the grave. Putative date: late Hellenistic - early Roman period, 1st century BCE - 1st century CE (ca. 100 BCE - 100 CE).
Deep-sequenced individual: **Ten_Pel_LHellenisticERoman**.

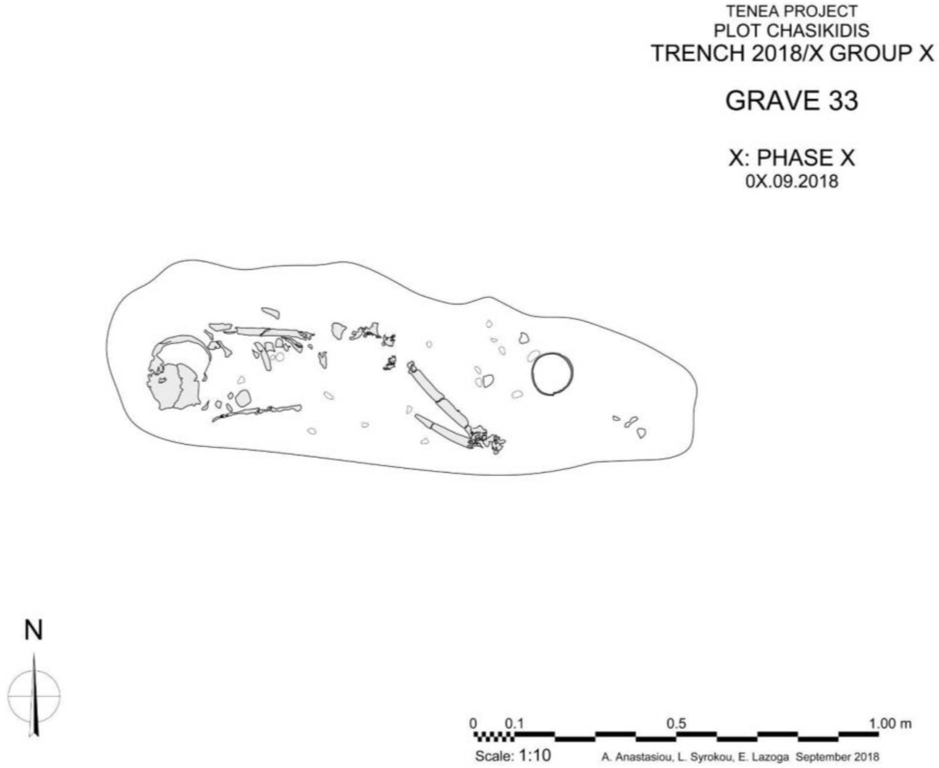


**Supplementary Figure S21.** Grave 33, cemetery of Tenea (Palaio Scholeio, Hasikidis plot).

**Roman period**

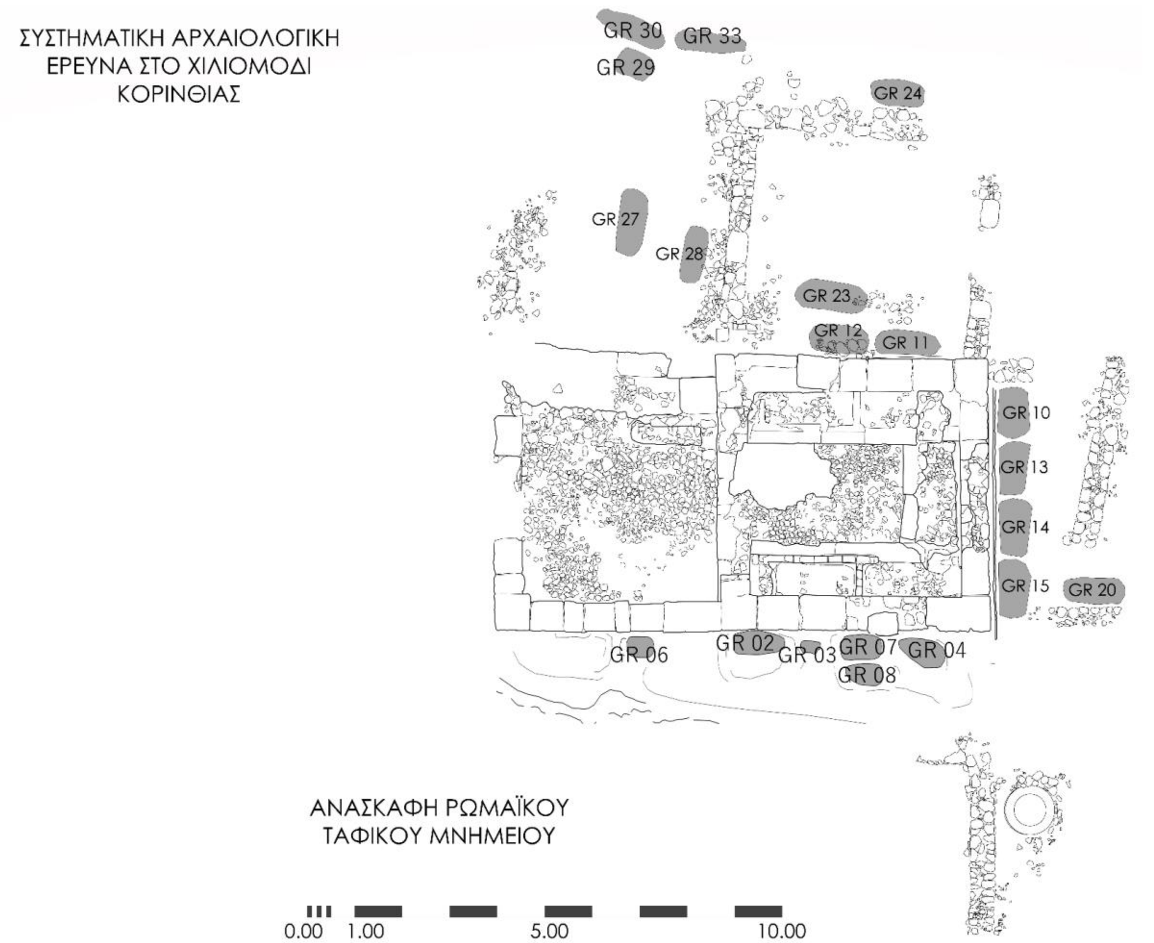


**Supplementary Figure S22**. The Roman graves in the cemetery of Tenea (Palaio Scholeio, Hasikidis plot).

-**Grave 07**: The pit - kalyvites grave (**Supplementary Figure S23**), oriented E-W, was excavated on 06/10/2016, in the cemetery of Tenea (Palaio Scholeio, Hasikidis plot). The grave was found intact, with dimensions 1.50 × 0.60 m. The grave contained the burials of three children. The main child, aged about 5 years old (Individual 1) was placed in an extended position, while the skulls and some long bones of two other children were found toward the west of the main burial. The individuals were accompanied by a glass unguentarium (perfume bottle), a needle and several nails. The chronology is based mostly on the grave offerings and dates to the Roman Period (3rd century CE).

Deep-sequenced individual: Individual 1, **Ten_Pel_Rom_3**.

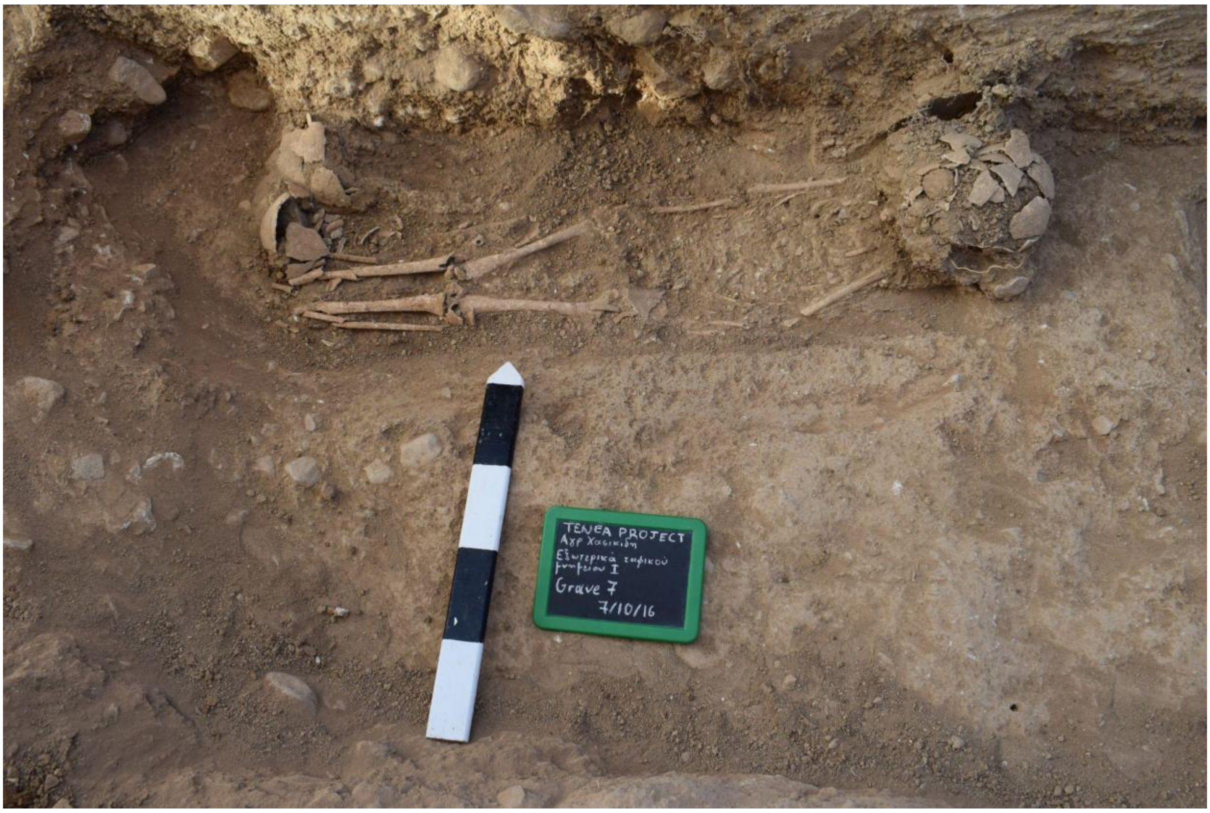


**Supplementary Figure S23.** Grave 07, cemetery of Tenea (Palaio Scholeio, Hasikidis plot).

-**Grave 15**: The pit - kalyvites grave (**Supplementary Figure S24**), oriented N-S, was excavated on 15/09/2017 in trench 2017/4, in the cemetery of Tenea (Palaio Scholeio, Hasikidis plot). The grave was found intact, with dimensions 1.17 × 0.52 m. The grave contained the burial of one child, aged about 7 years. The individual was accompanied by a lamp bearing an image of Aphrodite, a ceramic plate and a coin. The chronology is based mostly on the grave offerings and dates to the Roman period, (2nd century CE).

Deep-sequenced individual: **Ten_Pel_Rom_2**.

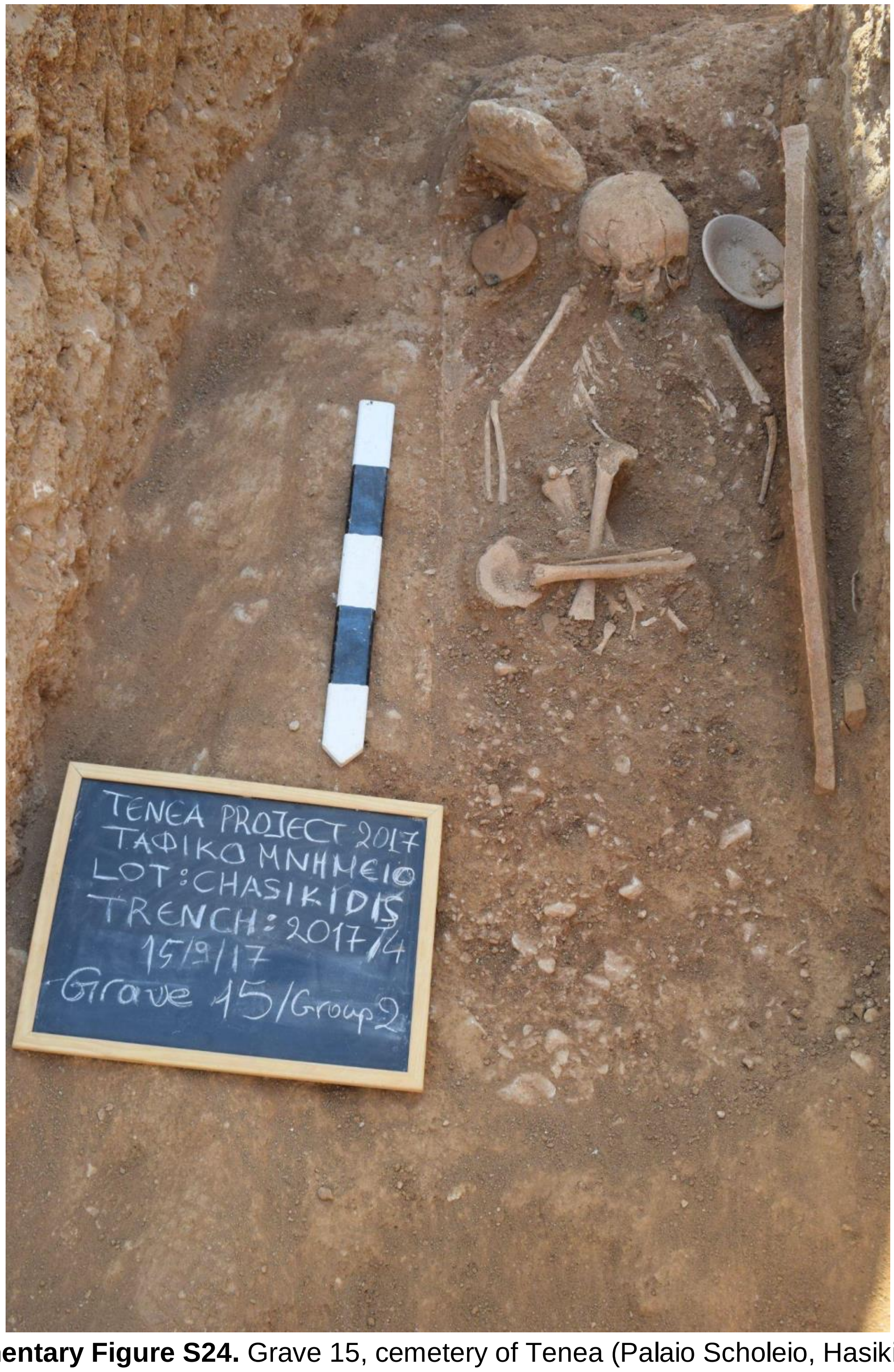


**Supplementary Figure S24.** Grave 15, cemetery of Tenea (Palaio Scholeio, Hasikidis plot).

-**Grave 22**: The pit-grave (**Supplementary Figure S25**), oriented NW-SE, was excavated on 06/10/2017 in trench 2017/2, in the cemetery of Tenea (Palaio Scholeio, Hasikidis plot). The pit grave carried a porous lid. The dimensions of the grave are 1.00 × 0.60 m. The grave contained the burial of a woman aged around 18 years old and a fetus around 20–22 weeks old, found in the pelvic area. The individuals were accompanied by a gold foil, while remains, probably of a wooden carrier, were collected along with iron nails. From the evidence of the burial assemblage, it is not possible to determine the chronology of the grave, although it most likely belongs to the roman period (31 BCE - 330 CE).
Deep-sequenced individual: Individual 1 (adult), **Ten_Pel_Rom_4**.

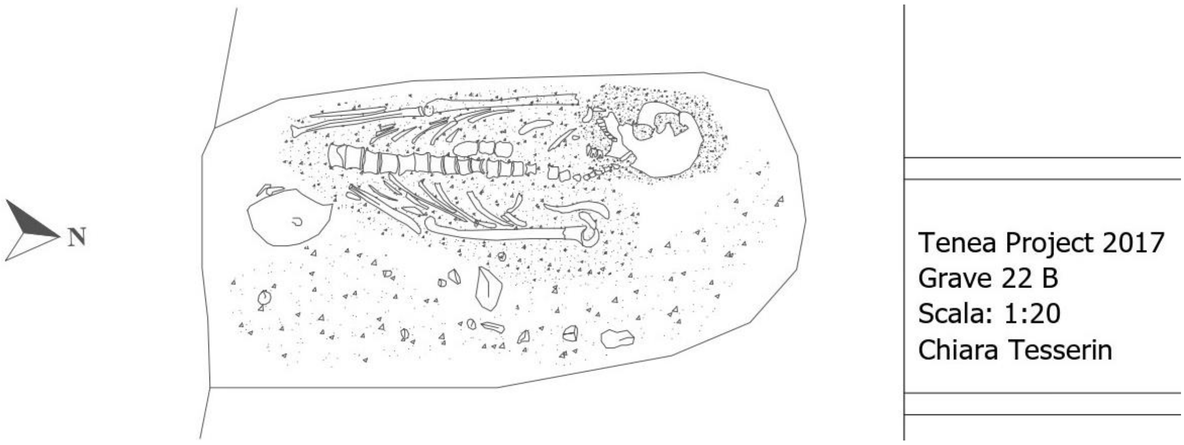


**Supplementary Figure S25.** Grave 22, cemetery of Tenea (Palaio Scholeio, Hasikidis plot).

-**Grave 30**: The pit - kalyvites grave (**Supplementary Figure S26**), oriented SE-NW, was excavated on 07/09/2018 in the cemetery of Tenea (Palaio Scholeio, Hasikidis plot). The grave contained three burials of two adults (Individual 1 and Individual 3) and one child (Individual 2). The individuals were accompanied by a gold earring, handless vase, lekanis, bone pins, and a Hellenistic coin placed as *Charon's obol*. The chronology is based on the grave offerings and dates to the Roman period (3rd - 4th century CE).
Deep-sequenced individual codes: Individual 2 (child), **Ten_Pel_Rom_1**.

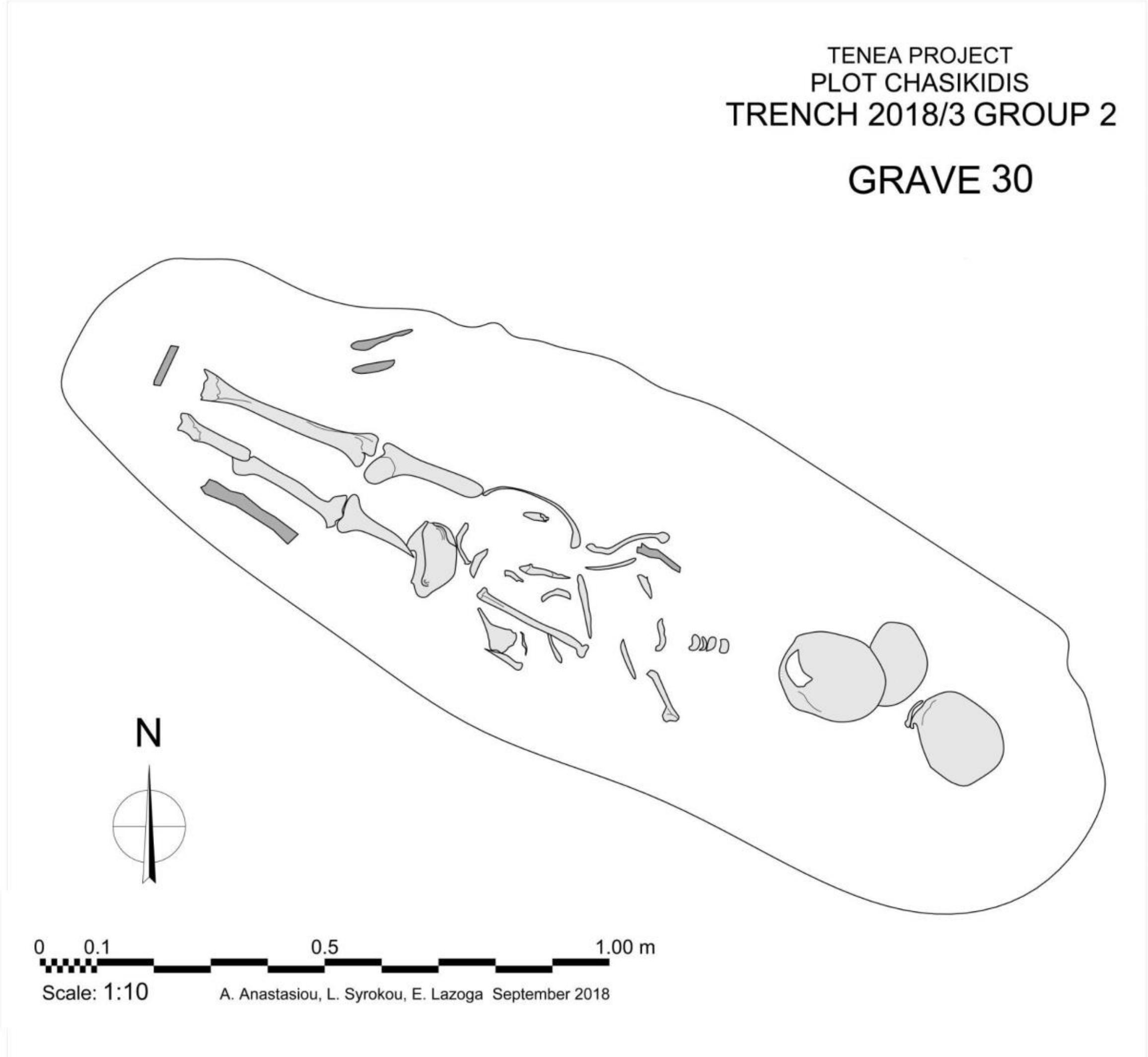


**Supplementary Figure S26.** Grave 30, cemetery of Tenea (Palaio Scholeio, Hasikidis plot).

### 3. Ancient DNA Analysis

#### 3.1 Sample Preparation

***Despoina Vassou, Sevasti Koursioti, and Nikolaos Psonis***

All analyses, that is, sample processing, DNA extraction, and genomic library preparation were performed in the cleanroom facilities of the Ancient DNA Lab at IMBB-FORTH. Negative controls (DNA-free) were included in all steps of the experimental procedure (DNA extraction, library preparation, and PCR amplification) to control for exogenous DNA contamination. Details on the analyses conducted for each sample are provided in **Additional file 1**.

*Dental samples processing*

For all dental samples, we used the minimally destructive dental root cementum decalcification method of Harney et al. [50]. Briefly, the outer surface of the tooth was decontaminated by a series of gentle washes using a cotton swab, starting with water, then 0.5% sodium hypochlorite solution (bleach), water again to remove all bleach traces, and finally absolute ethanol to remove all water traces. The samples were left to dry completely and were subsequently UV irradiated (6 J/cm2 at 254 nm) in a UVP CL-1000 UV Crosslinker for 10 mins on each side. Each dental root was submerged in a 1 ml extraction buffer (0.45M EDTA, 0.05% Tween 20, Proteinase K 0.25 mg/ml) at 42 ℃ under mild agitation. The buffer was exchanged after 24 hours in some cases, if cementum was still visible. DNA extraction and purification was performed using the magnetic beads-based protocol of Rohland *et al*. [51] with buffer D.

*Pars petrosal samples processing*

For the majority of the temporal bone samples, the outer surface of the petrous area was cleaned using a diamond disk at low speed with a multitool drill (ProLab Basic Laboratory Control, Bien Air, Switzerland). *Pars petrosal* was isolated and powdered using pliers. For one sample (Amv_Epi_Arch_3), we used the minimally destructive petrous bone powder sampling method of Orfanou *et al*. [52] that involves targeted drilling toward the petrous area without cutting the bone. Before drilling, the outer surface of the petrous bone was decontaminated in the same way as described above for the teeth. For all *pars petrosal* samples, approximately 50-150 mg of powder were used for demineralization, DNA extraction and purification using either the magnetic beads-based or the large-volume column-based protocol of Rohland *et al*. [51] with buffer D.

*Genomic libraries and sequencing*

Double stranded, blunt-end libraries were constructed following published protocols [53,54], albeit by substituting buffer BL01 with buffer D [55] to enrich for smaller fragments of DNA. No library pre-treatment was used during pre-screening. The number of amplification/indexing PCR cycles for each library was determined with qPCR. Single or double indexing was performed for pre-screening purposes. Purification of amplified libraries was performed twice with AMPure XP Beads (BeckMan Coulter, Inc., USA) at a 1:1.8 ratio (DNA:beads). Quality control and quantification of the libraries was performed with Qubit

(Thermo Fisher Scientific, Inc., USA) and Bioanalyzer (Agilent Technologies, Inc., USA). Endogenous DNA content and post-mortem deamination (PMD) damage was initially estimated via shallow sequencing on an Illumina NextSeq500 platform using either, single-end (1×75), or paired-end (2×75) chemistry (Genomics Facility, IMBB-FORTH, Greece).

For deeper sequencing on selected samples and libraries (useful mapped content >1.0%), we prepared fresh libraries (from previous or new DNA extractions) following the same procedure as above, albeit with two modifications: a) pre-treatment of the libraries with the USER™ enzyme (New England BioLabs Inc., USA) for 30 min [partial UDG-treatment method of Rohland *et al.* [56]] in order to reduce deamination misincorporation in the DNA sequence data and b) double indexing with unique 8bp barcodes in order to reduce potential cross-sample contamination issues during indexing. USER treatment was not performed for two samples (Amv_Epi_Arch_2 and Amv_Epi_Arch_3). Deeper sequencing was performed in an Illumina Novaseq6000 platform (Macrogen, Inc., South Korea), using paired-end (2×100 and 2×150) chemistry.

#### 3.2 Read Processing, Damage estimation, Genetic sex determination

***Nikolaos Psonis***

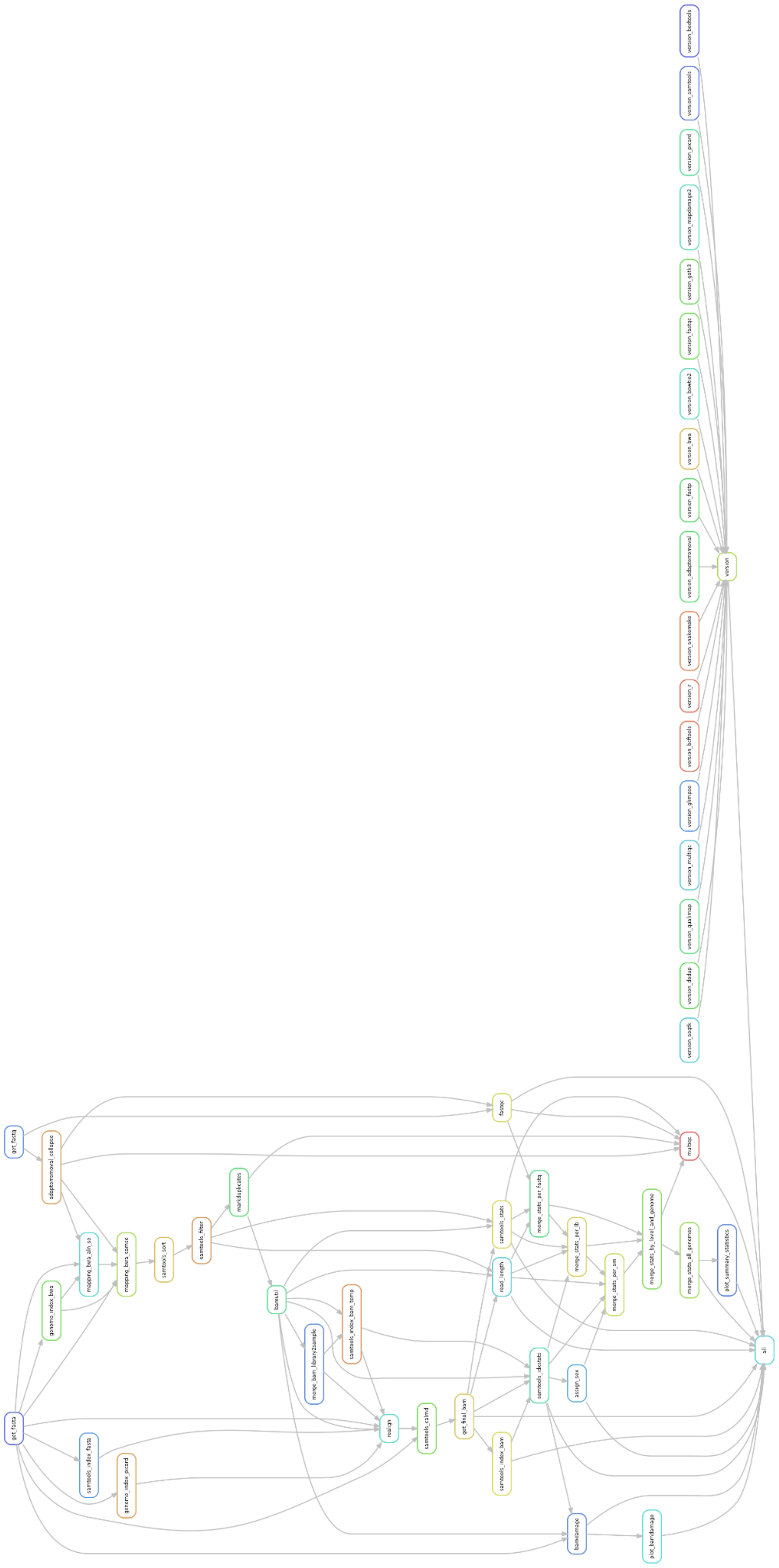

**Supplementary Figure S27.** Rulegraph of the mapache pipeline used in this study, as produced by snakemake. The graph depicts the second mapache run that includes bamUtil.

Initial analyses using the raw reads were performed at three levels: (a) at the FastQ level corresponding to the sequencing reads in each FastQ file, (b) at the library level, corresponding to multiple BAM files from the same (PCR amplified) library, and (c) at the level of individuals, corresponding to multiple BAM files from the same individual. All analyses were performed as implemented in the mapache v.0.3.0 commit f1316e1 [57] pipeline (**Supplementary Figure S27**) by using the snakemake v.7.18.2 workflow manager [58] with parameter `--notemp` to retain intermediate files that are required for downstream analyses (e.g., the BAM files at the FastQ or the library level). In order to ensure reproducibility of the results, our mapache configuration (config.yaml) and samplelist files are available at https://doi.org/10.5281/zenodo.10848927.

Note that pre-screening data were exclusively used to identify the most promising libraries for deep sequencing and are not included in the final dataset (i.e., not merged with the deep sequencing data).

All computational analyses were conducted on an AMD EPYC 7452 system with 64 physical cores and 1 TB of RAM running Ubuntu 20.04.6 LTS.

##### 3.2.1 Analyses at the FastQ level

We received already de-multiplexed sequences from the sequencing facility. De-multiplexing relied on the two 6- to 8-bp barcodes of each double-indexed library. Initial quality control was performed for each FastQ file using FastQC v.0.11.9 (http://www.bioinformatics.babraham.ac.uk/projects/fastqc/). Using AdapterRemoval v.2.3.2 [59] the raw sequences were filtered using a base quality of 2 [default value; values > 2 yield a larger amount of short sequences that will not be propagated to the mapping step; stricter base quality filtering (Illumina proposed values of >20-30 that equal to 0.01-0.001 error probability) is applied *after* mapping (see downstream analyses)] to remove low quality bases from the read ends, trimmed for Illumina adapters and stretches of ambiguous bases (Ns). Paired-end reads were merged by requiring at least 11 bp (default value; a corresponding parameter exploration with 9 up to 11bp showed no substantial difference in results) overlap between the pairs, whereas merged reads shorter than 30 bp were discarded to avoid mere random matches [see e.g. 60] in the next step (parameters used: `--trimqualities --gzip --trimns --collapse --minalignmnetlength 11 --minlength 30`). Only the output with fully (non-truncated) collapsed reads was used for downstream analyses. FastQC was then used again to verify that (i) trimming was successful and (ii) to assess the post-adapter-removal quality of the reads.

The merged reads were mapped to the 1000 Genomes project version of the human reference genome *hs37d5* [61] using BWA v.0.7.17 [62] and the *aln/samse* algorithm using settings optimized for aDNA reads, including disabling the seed length (`-l 1024`) and using `-n 0.01` (the fraction of missing alignments given a 2% uniform base error rate) and `-o 2` (maximum number of gap openings) to allow for higher sensitivity [63] and to minimize reference bias in downstream population genomics analyses [64]. Informative read groups (RG) were also added during the above step (using `-r`) to keep track of PCR-parallels. SAMtools v.1.14 [65] was used to sort the uniquely mapped reads by chromosome order (*sort* function) and to filter them using the *view* function for a mapping quality of 30 (`-q30`) such as to only keep alignments with but a few mismatches, remove reads with flag 4 (denoting an unmapped read) in their header (`-F4`), and index (*index* function) the final BAM file. Note that indexing was also performed in all BAM outputs generated by the intermediate steps outlined below.

Trimming and mapping metrics reported by mapache at the FastQ level include (a) the absolute number and proportion of fully collapsed reads from the total number of raw reads (reported by mapache as trim_prop and reads_trim, respectively), as well as their mean length (length_reads_trimmed) and (b) the absolute number and the proportion of mapped reads - including duplicates - from the total number of raw reads, known as *mapped content* (reported by mapache as mapped_raw and endogenous_raw, respectively), as well as their mean length (length_mapped_raw). All of the above metrics are provided in **Additional file 2**.

##### 3.2.2 Analyses at the library level

Multiple BAM files from the same library were merged with SAMtools *merge*. Duplicated sequences (PCR clones and single amplification clusters incorrectly detected as being multiple clusters by the sequencer's optical sensor; also known as optical duplicates) at the library level were removed using the *MarkDuplicates* function of the Picard software tool.

Mapping metrics calculated by mapache at the library level included the same metrics as at the FastQ level, but also (a) the absolute number and proportion of duplicate reads in the overall number of mapped reads (reported by mapache as duplicates and duplicates_prop, respectively) and (b) the absolute number and proportion of uniquely mapped (non-duplicate) reads in the overall number of reads, known as *useful mapped content* (reported by mapache as mapped_unique and endogenous_unique, respectively), as well as their length (length_mapped_unique). In addition, we manually calculated (c) the *endogenous DNA content* (ratio of number of mapped reads, including duplicates, over the number of fully (non-truncated) collapsed reads), (d) *the efficient endogenous DNA content* (number of mapped reads, excluding duplicates, to the number of fully collapsed reads). All mapping metrics mentioned above are provided in **Additional file 2**.

Based on a visual inspection of the damage plots (see below) and to avoid incorporating incorrect sequence information caused by *post-mortem* deamination at either end of the reads, we soft-clipped (a) six bases at either ends of each read in libraries obtained by shotgun sequencing *without* UDG (USER$^{TM}$) treatment and (b) two bases at either ends of each read from libraries treated *with* UDG. Soft clipping was performed using the *trimBam* function of bamUtil v.1.0.15 [66].

To investigate the level of *post-mortem* DNA degradation, such as DNA fragmentation and deamination (C-to-T for both ds- and ss-libraries and G-to-A transitions for ds-libraries only), the uniquely mapped deduplicated sequences were analyzed with a modified version[7] (see https://github.com/sneuensc/mapache/wiki/3.-Config-file-(parameters)) of bamdamage [67] as implemented in the mapache pipeline using 10,000 reads per BAM file. By default, mapping quality and base quality thresholds were set to 30 and 20, respectively, in order to keep well-aligned and high-quality reads only. Deamination damage values are provided in **Additional file 3**.

[7] the following changes were made:
- speed up: there is now a subsampling (every nth alignment) if desired. Can be specified in the config file.
- Output is not just the pdf, but also the underlying data. The figure is improved and the y-axis of R1 and R2 have the same scale.

Note that in the mapache pipeline, the bamUtil program is executed before mapdamage. Thus, in order to properly estimate deamination damage, the mapache pipeline was run twice as described in the next section.

##### 3.2.3 Analyses at the individual level

Multiple BAM files from the same individual were merged via the SAMtools *merge* function. We did not remove identical sequences at the individual level as these represent original DNA fragments from different cells of the same individual, rather than duplication artifacts from our analysis procedure. Local indel re-alignment was performed at the individual level using the RealignerTargetCreator and IndelRealigner tools of GATK v.3.8 [68] after recomputing the MD tag using the SAMtools *calmd* function. The mean genome depth (as well as that of the X and Y chromosomes and of the human mitogenome) coverage was determined at the individual level using Qualimap v.2.2.2d [69]. Mapping metrics calculated by mapache at the individual level were the same as those mentioned for the library level, as well as the mean depths of coverage. All mapping metrics mentioned above are provided in **Additional file 2**. Overall, the total raw sequences per individual ranged from 77232103 to 2379802838, the proportion of fully collapsed reads was between 65.45% and 84.21%, the mapped content varied between 1.27% and 48.42% with a proportion of duplicated reads ranging from 16.57% to 27.80%. The mean coverage depth exceeded ~0.05× (0.07 - 6.31×) for all 26 individuals.

Genetic sex inference was performed using two approaches. First, we used the Rx method [70] as implemented in mapache. This method relies on the ratio of the normalized X-chromosome coverage depth to the normalized autosomal depth. Secondly, we manually applied (not in the mapache pipeline) the Ry method [71] using the python script provided by the authors (the input was piped via SAMtools *view*). This method is based on the ratio of the reads mapped exclusively to Y, and to both, X, and Y chromosomes, respectively. The results are provided in **Additional file 3**. Overall, the genetic sex inference analyses confirmed that nine individuals were males and 17 were females. Both methods used mostly agreed with each other (albeit the Ry method failed to assign sex in three cases).

As mentioned above, in the mapache pipeline the program bamUtil is executed at the library level *before* mapdamage. Thus, the mapache pipeline was run twice: (a) First, we disabled the bamUtil program in the mapache configuration file. The BAM files of the run containing *post-mortem* damage were not used for downstream analyses, with the exception of a mtDNA-based contamination estimation analysis (see section 4.3). (b) For the second run, we renamed the main output directories of mapache (*0_2_library*, *0_3_sample*), enabled bamUtil, and generated the BAM files at the library and individual level using data without *post-mortem* damage. We used these BAM files for downstream population genomics analyses.

All individuals' genetic data were characterized by an ancient-like DNA signature. More specifically, in the USER-treated libraries, the C-to-T and G-to-A deamination damage at the two first bases of the read ends showed a "spike" pattern, with the C-to-T damage at the first base of the 5' end of the reads (**Additional file 3**) ranging (among the different libraries) from 7.74% to 36.97%. The two non-USER-treated libraries (256_1_B_lys2_ex1_lib2 and 299_B_lys1_ex1_lib1) displayed the classic "smiley" pattern and their 1st-base-5'-end C-to-T damage was 54.09% and 42.30%, respectively. In conjunction with the deamination damage, the mean fragment length of the deduplicated mapped reads of each library (**Additional file 2**) correspond to the characteristics of degraded DNA, ranging from ~39 bp to ~80 bp (collapsed-reads; 150 sequencing chemistry was used). As expected, the damage plots at the library level did not show the deamination effect for the second run, in which bamUtil was used.

#### 3.3 Contamination estimation

***Nikolaos Psonis***

Genome authenticity was further verified by using three distinct contamination estimation approaches, all performed at the individual level. One approach is X-chromosome-based and two are mtDNA-based. In the analyses below, indexing of BAM and FastA files was performed with the SAMtools *index* and bwa *index* functions, respectively.

First, we performed an mtDNA-based contamination estimation by using contamMix v.1.0-10 [72] with a minimum base quality filter of 30 (default value) to only retain bases of good quality and a set of 311 modern mitochondrial genomes from around the globe [73] serving as sources of potential contamination. For this method, it is necessary to construct the consensus mtDNA sequence from the BAM files. To this end, the majority-rule consensus sequence was computed with ANGSD v.0.941-6-g67b6b3b [74] using the following parameters: `-doCounts 1 -minMapQ 30 -minQ 30 -doFasta 2`, in order to exclusively use well-aligned reads. Hence, as input files in contamMix we used: (a) `--samFn` (BAM format): mitochondrial mapped reads (MT-reads) from the full alignment file with clipped deaminated bases (second mapache run; see above), extracted using SAMtools *view* and (b) `--malnFn` (FastA format): a multiple sequence alignment (MSA) containing the consensus mitogenome above and the aforementioned 311 worldwide modern mitochondrial genomes. The MSA was computed with the automatic mode of mafft v.7.505 [75].

Then, we performed a second, independent mtDNA-based contamination estimation with schmutzi v.1.5.6 [76]. MT-reads from the full alignment file that contained deaminated bases (first mapache run; see above), were extracted using SAMtools *view* and realigned against rCRS only (using bwa and SAMtools; same parameters as in **Section 3.2**) as proposed by the software developers. Recomputing the MD tag of the mtDNA BAM files that provides information about the reference base and is used for SNP/indel calling without taking into account the reference was performed with SAMtools *calmd*. According to the workflow recommended by the developers for ancient samples, we first applied the *contDeam* method to estimate initial contamination and endogenous deamination rates. Depending on the library type, we set the `--library` option to single or double and we also considered only two bases at both read ends as being deaminated (`--lengthDeam 2`). Following this, we called the actual schmutzi method (*mtcont*), that is, the iterative procedure without the prediction of the contaminant (`--notusepred`), but also with contaminant prediction. As potential contamination sources we used a database of 197 mitochondrial allele frequencies that are provided with the schmutzi software.

Finally, we used an X-chromosome-based contamination estimation on XY samples by using the contamination function of ANGSD based on haploid X-chromosomal regions (`X:5000000-154900000`). First, we ran ANGSD with the options `-minMapQ 30`, `-minQ 30`, `-doCounts 1`, and `-iCounts 1`, in order to only use well-aligned reads. Then, we executed contamination providing the publicly available HapMap file HapMapChrx.gz (https://github.com/ANGSD/angsd/tree/master/RES).

All contamination estimation analyses mostly supported each other (**Additional file 3**) resulting in >98.49% authenticity levels in ContamMix, <4% contamination estimates in schmutzi (if analysis applicable; both with and without `--notusepred`), and <1.62% contamination estimates based on X-chromosome in XY samples.

#### 3.4 Uniparental haplogroup estimation

***Nikolaos Psonis***

The classification to mitochondrial and Y-chromosomal haplogroups was performed using data at the individual level from reads with clipped deaminated bases (second mapache run; see above).

##### 3.4.1 mtDNA

We generated the mtDNA consensus sequence for each individual by using the MT-reads from the BAM file after mapping against *hs37d5* (extracted with SAMtools *view*). The majority-rule consensus sequence in both cases was called with ANGSD using the following parameters: `-doCounts 1 -minMapQ 30 -minQ 30 -setMinDepth 2 -doFasta 2`, in order to only use well-aligned reads and avoid misincorporation of sequencing errors. The classification to haplogroups was performed with the webtools HaploGrep3 v.3.3.2.1 [77] and HaploCart v.1.0 [78] for the sake of comparison and cross-validation. The results are given in **Additional file 3**. The mitochondrial haplogroup assignment by the two methods used was almost identical, albeit in a few cases (10/26), the two methods produced different assignments to the most external leaf of the mitochondrial tree.

The LBA Ammotopos samples were assigned to the J1c(or 2) and T2b3 haplogroups. Amvrakia included H15a1b, T1a4, and U5a1g1 during the Archaic period, H, K1a2, N1a1a1(or b), and T2b(6 or 3c) during the Classical period, and H46, J2b1(or a) and W(+194 or 9) during the Hellenistic period. An additional individual from the late Classical - Hellenistic times was assigned to the H5a(3 or 3a) haplogroup. Finally, Tenea included T1a4 during the Archaic period, T2n and U3a3 during the Hellenistic period, and N1a1a(+152 or 2), U1a1c(or 1), and U3a3 during the Roman period.

According to the Allen Ancient DNA Resource (Version 8; aadr_v.54.1.p1_1240K_public; [79]), the aforementioned haplogroups have been observed: a) J1c in multiple individuals in Neolithic to Medieval Europe and Middle East, including IA Greece, b) J1c2 only in Neolithic to Medieval Europe, but not in Greece, c) T2b3 in prehistoric Europe, but not in Greece, d) H15a1b, in only one individual in LBA-EIA Armenia, e) T1a4 only in a couple of Neolithic to Medieval Europe, but not in Greece, f) U5a1g1 in prehistoric Eurasia, but not in Greece, g) H in individuals in Neolithic to Medieval Europe and Middle East, including Neolithic, BA, but also Roman Greece, h) K1a2 in individuals in Neolithic to Medieval Europe and Middle East, including Neolithic and BA Greece, i) N1a1a1 predominantly from Neolithic Europe, but not in Greece, j) N1a1a1b has not been observed, k) T2b6 has not been observed, l) T2b3c only in Neolithic Ireland, m) H46 in prehistoric Europe, but not in Greece, n) J2b1 in prehistoric and historical Europe and Middle East, but not in Greece, o) J2b1a in Neolithic to Medieval Europe, but not in Greece, p) W194 in prehistoric Europe, but not in Greece, q) W9 only in one individual from Ottoman Anatolia, r) H5a3 in BA Germany, only, s) H5a3a in a couple ancient European individuals, but not in Greece, t) T2n has not been observed, u) U3a3 only in BA Jordan, v) N1a1a+152 in prehistoric Europe and Middle East, but not in Greece, w) N1a1a2 has not been observed, , x) U1a1c in prehistoric and historical West Asia, and y) U1a1c1 in BA Iran and in Medieval Russia.

##### 3.4.2 Y-chromosome

Classification into Y-chromosomal haplogroups was performed using Yleaf v.3.1 [80] with a minimum read quality of 30 (`-q 30`), a minimum percentage of a base result for acceptance of 90 (`-b 90`), in order to only use well-aligned reads and avoid misincorporation of sequencing errors and the *hg19* reference genome (`-rg hg19`) and its accompanying predict_haplogroup.py python script. The minimum prediction score was 0.95 (set by default). This version of Yleaf uses YFull (v.10.01) for the underlying haplogroup tree structure.

For the sake of comparison and verification, we also used Yhaplo v.1.1.2 [81]. First, by piping the BCFtools v.1.15 [82] *mpileup* and *call* modules, we called and filtered SNPs found at the Y chromosome (`-r Y`), kept bases with base quality>=30 (`-q 30`) and reads with mapping quality>=30 (`-Q 30`) to ensure that only well-aligned reads were used, and we downgraded mapping quality for reads containing excessive mismatches (`--adjust-MQ 50`; value recommended by the developers) to avoid keeping (small, ancient) reads that originate from another region or a different species. We also set ploidy to 1 (`--ploidy 1`; Y-chromosome is haploid) and selected the multiallelic caller (`-m`) designed for rare-variant calling. Then, using the BCFtools *norm* module, we performed normalization (left-alignment and normalization of indels; check if REF alleles match the reference; split of multiallelic sites into multiple rows; recovery of multiallelics from multiple rows). Finally, we ran Yhaplo with the -aao parameter in order to generate all auxiliary outputs. Yhaplo uses the ISOGG Y-DNA Haplogroup Tree (2016.01.04; https://isogg.org/tree/). The results are given in **Additional file 3**. The Y-chromosome haplogroup assignment to the major haplogroups by the two methods used in the present study was identical (9/9). However, due to partial coverage of the Y-chromosome, the estimated haplogroups may not represent the assignment of the samples to the most external node of the Y-chromosomal tree.

The male LBA Ammotopos sample was assigned to G2a2b2a1a1c1a, the male Amvrakia samples were assigned to J2 and T1a2 (Classical period) and E1b1b1 (Hellenistic period), and the male Tenea samples were assigned to T1a2 (Archaic period), E1b1b1a1b1 (Hellenistic period), and R1b1a2a2a and J2a1b1 (Roman period). The Allen Ancient DNA Resource (Version 8; aadr_v.54.1.p1_1240K_public; [79]) does not include entries from prehistoric or historical Greece for any of the aforementioned haplogroups.

#### 3.5 Population Genomics analysis

***Stefanos Papadantonakis, Angelos Souleles, Pavlos Pavlidis, Angeliki Papadopoulou, and Nikolaos Psonis***

##### 3.5.1 Lists of genomic sites

We used two different lists of genomic sites, one containing ~1.24 million sites, known as 1240K [83] that is extensively being used in human archaeogenomics research, and another one containing ~5 million sites (5M_auto; 5M hereinafter) that was recently generated [84]. The lists are provided at https://doi.org/10.5281/zenodo.10848927.

##### 3.5.2 Genotype calling and pseudohaploidization

For each of the aforementioned two lists of genomic sites (see section 4.5.1), SNP calling (in pileup format) per individual was performed with the SAMtools *mpileup* function by providing the position of sites in the reference genome, disabling the per-Base Alignment Quality, known as BAQ (`-B`) to reduce reference bias, ignoring read groups (RG) tags (one BAM = one sample), and skipping alignments and bases with mapping and base quality, respectively, smaller than 30, to only retain the well-aligned reads. Random pseudohaploidization was performed with the *pileupCaller* module of SequenceTools v.1.5.2 (https://github.com/stschiff/sequenceTools) by providing the pileup file and an EIGENSTRAT snp file with the positions of each list of genomic sites, using the `--randomHaploid` parameter, and selecting EIGENSTRAT as output format. The sex field in the output individual (ind) file was annotated using the result of the genetic sex inference above (see section 4.2). Calculation of coverage depth for the distinct lists of genomic sites was performed with the eigenstrat_snp_coverage.py v.1.1.0 python script (https://github.com/TCLamnidis/EigenStratDatabaseTools).

We merged individual EIGENSTRAT files to assemble different datasets (see below) with the EIGENSOFT v.7.2.1 [85] *mergeit* function and converted them to PACKEDPED (BED) files with the *convertf* function of the same software package.

##### 3.5.3 Genetic relatedness analysis

To assess genetic relatedness, we used two different approaches developed for aDNA data, Relationship Estimation from Ancient DNA (READ) [86] and KIN v.3.1.3 [87]. READ can estimate up to 2nd-degree genetic relationships, whereas KIN identifies up to 3rd-degree genetic relatives provided at least 0.05× sequence coverage. KIN can also disentangle siblings from parent-child pairs. As lists of genomic sites we used both, the 1240K, and 5M lists for the sake of comparison [following the procedure outlined in 88]. For the READ analysis, we first extracted the 22 autosomal chromosomes (only for the 1240K panel, 5M includes only autosomal sites) using PLINK v.1.90b6.21 [89] and selected tped as output format (`--recode transpose`). Then, we executed READ using the median normalization method to decrease the influence of outliers. For the KIN analyses, we used the *KINgaroo* module without contamination correction (`–cnt 0`) to generate the input files from bamfiles for the *KIN* module that followed. Although we report all resulting genetic relationships (https://doi.org/10.5281/zenodo.10848927), we considered only those as being valid, which yielded |Z|>1 for READ and Δ*LL*>1 for KIN, as proposed by the respective tool developers. We discovered three cases of genetic relatedness, two in Ancient Amvrakia, and one in ancient Tenea:

<u>Case 1 - Classical Amvrakia</u>

Grave CV (**Supplementary Figure S28)** contained two individuals (Amv_Epi_Cl_5 and Amv_Epi_Cl_6) dated to the Classical period (375-350 BCE). Both individuals were anthropologically estimated to be children (2-12 years old), whereas their genetic sex was inferred as being female for both (XX; **Supplementary Table 3; Additional file 3**). They also shared the same mt-DNA haplogroup (K1a2; **Supplementary Table 3**; **Additional file 3**). These two Individuals were inferred, by both READ and KIN, as 1st-degree genetic relatives, with KIN identifying them as siblings. Hence, these two individuals can confidently be recognised as sisters.

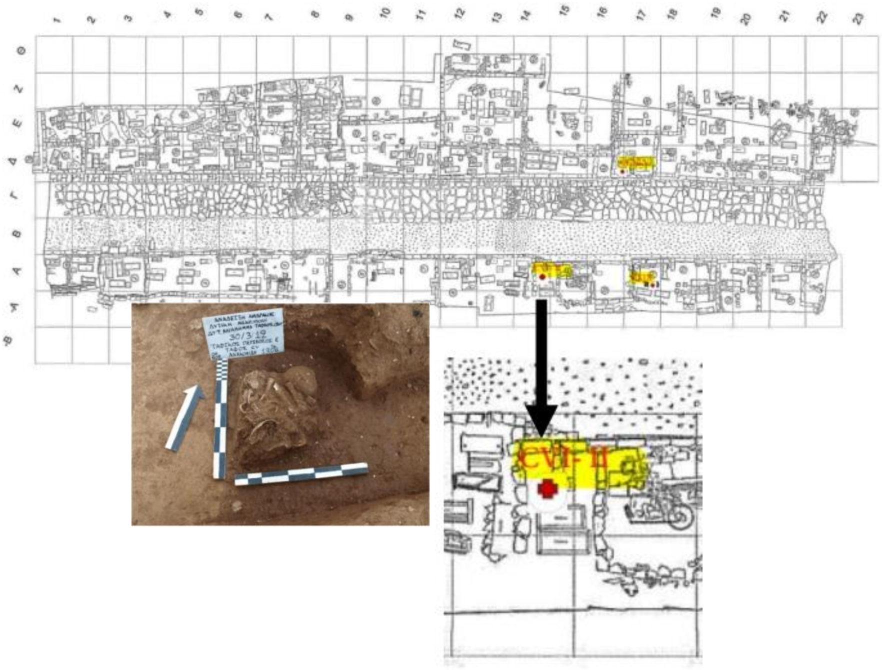

**Supplementary Figure S28.** The location of grave CV in the western necropolis of Ancient Amvrakia and the CV1 burial (retrieval).

Additionally, these two sisters were inferred (by both READ and KIN) to also have 2nd-degree genetic relationships with a third individual (Amv_Epi_Cl_1), found in a nearby grave (CXIX A; retrieval; **Supplementary Figure S29**) dated to around the same time (375-350 BCE). Anthropologically, this individual was estimated to be a young (20-34 years old) female (confirmed by genetic sex analysis; **Supplementary Table 3; Additional file 3**) and it belongs to a different mtDNA haplogroup (N1a1a1; **Supplementary Table 3; Additional file 3**). Hence, this individual could either be an aunt or the grandmother of the sisters (from their father's side), or their stepsister from a different mother. Based on the dating of these three burials and the age-at-death of Amv_Epi_Cl_1, the second scenario (grandmother-granddaughters) seems the least possible (although feasible).

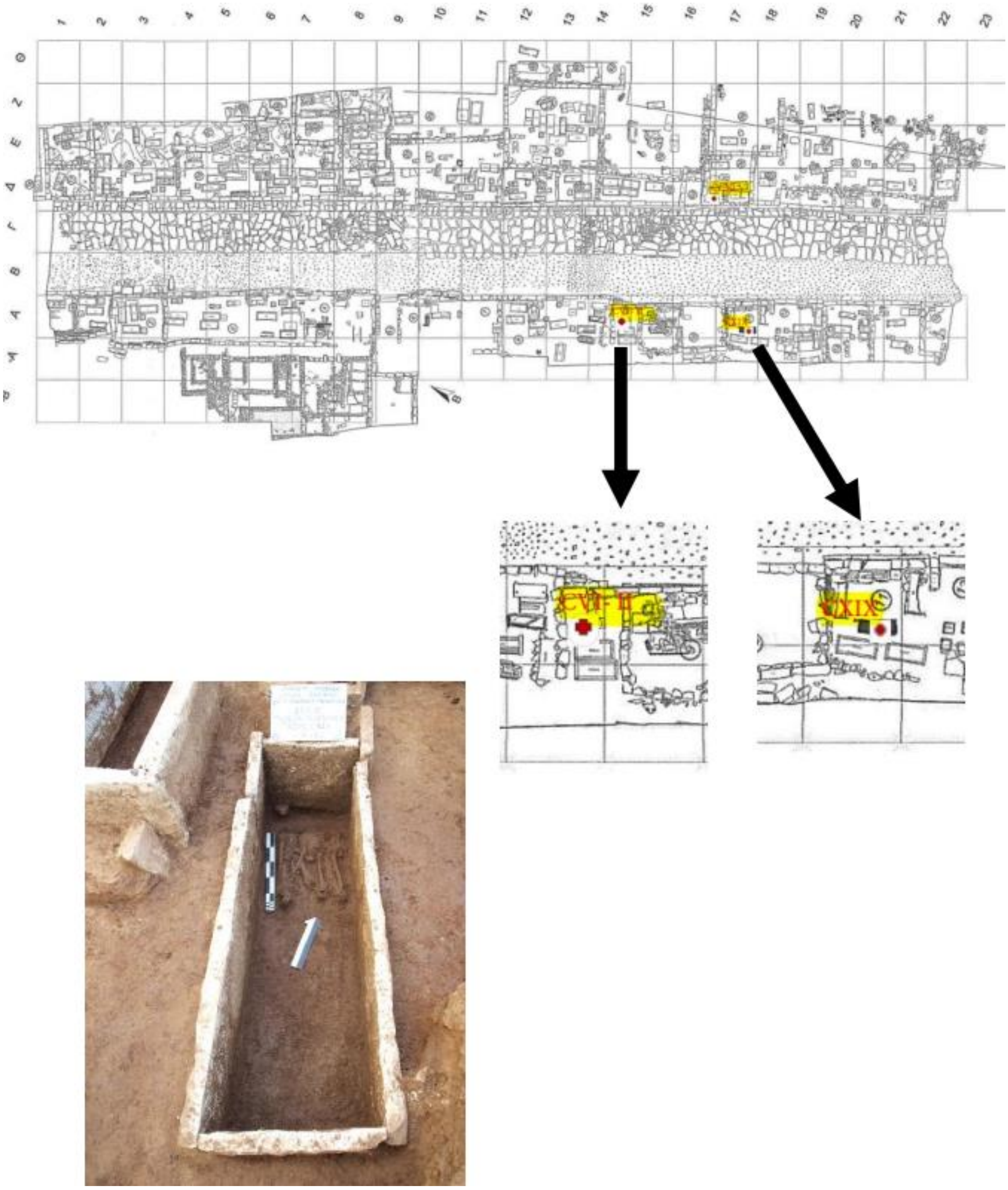


**Supplementary Figure S29.** The location of grave CXIX in the western necropolis of Ancient Amvrakia and the CXIX A burial (retrieval).

Case 2 - Hellenistic Amvrakia

Grave CCXLV (**Supplementary Figure S30**) contained two individuals (Amv_Epi_Hel_3 and Amv_Epi_Hel_4) dated to the Hellenistic period. They were buried with a small chronological difference (175-125 BCE), as the burial of Amv_Epi_Hel_3 is a retrieval, whereas the one of Amv_Epi_Hel_4 is a primary burial. Anthropologically, the first individual was estimated to be an old (>50 years old) male (possibly), whereas the second individual was also old (>50 years old), albeit the sex could not be determined. According to our genetic analyses, both individuals were females and share the same mtDNA haplogroup (W+194) (**Supplementary Table 3; Additional file 3**). READ and KIN inferred them to have a 1st-degree genetic relationship, with KIN indicating a parent-child relationship. Based on the fact that the burial of Amv_Epi_Hel_3 is a retrieval, we consider it most likely that this individual was the mother and Amv_Epi_Hel_4 the daughter, although the grave dating and the age-at-death cannot exclude the opposite.

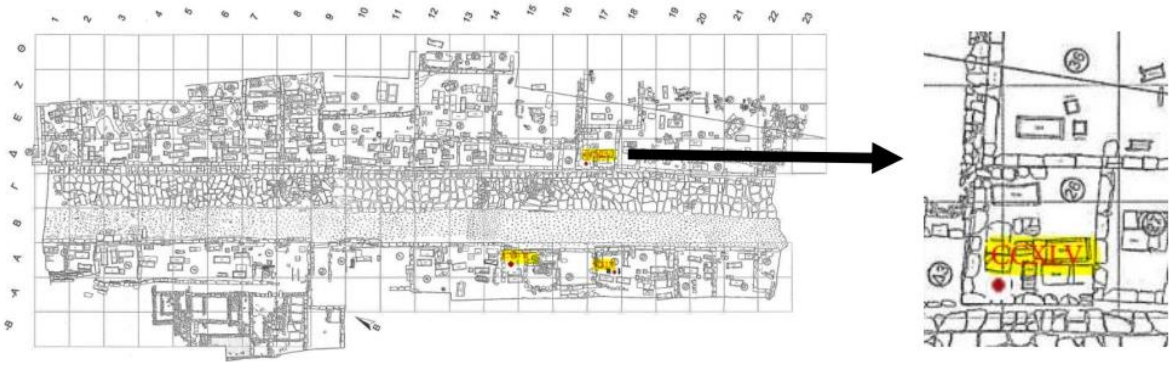


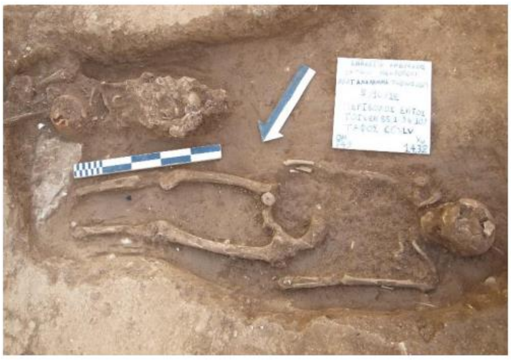

**Supplementary Figure S30.** The location of grave CCXLV in the western necropolis of Ancient Amvrakia and the two burials (retrieval and primary).

Case 3 - Archaic Tenea

At the Faneromeni-Kamareta site, two related individuals were found in two neighboring graves (sarcophagi) dated to the Archaic period (**Supplementary Figure S31**). Individual Ten_Pel_Arch_1 was found in grave 5 dated to 500 BCE, whereas individual Ten_Pel_Arch_2 was found in grave 3 dated to 550-500 BCE. The first individual was anthropologically determined to be an old (>50 years old) male, whereas the second a middle-aged (50 years old) female. The genetic analyses confirmed their genetic sex and indicated that they belong to the same mtDNA haplogroup (T1a4) (**Supplementary Table 3**; **Additional file 3**). READ and KIN inferred their genetic relationship as being of 1st degree, with KIN yielding a parent-child relationship. Given that they share the same mtDNA haplogroup, the most likely scenario is that they are mother (Ten_Pel_Arch_2) and son (Ten_Pel_Arch_1), rather than father and daughter.

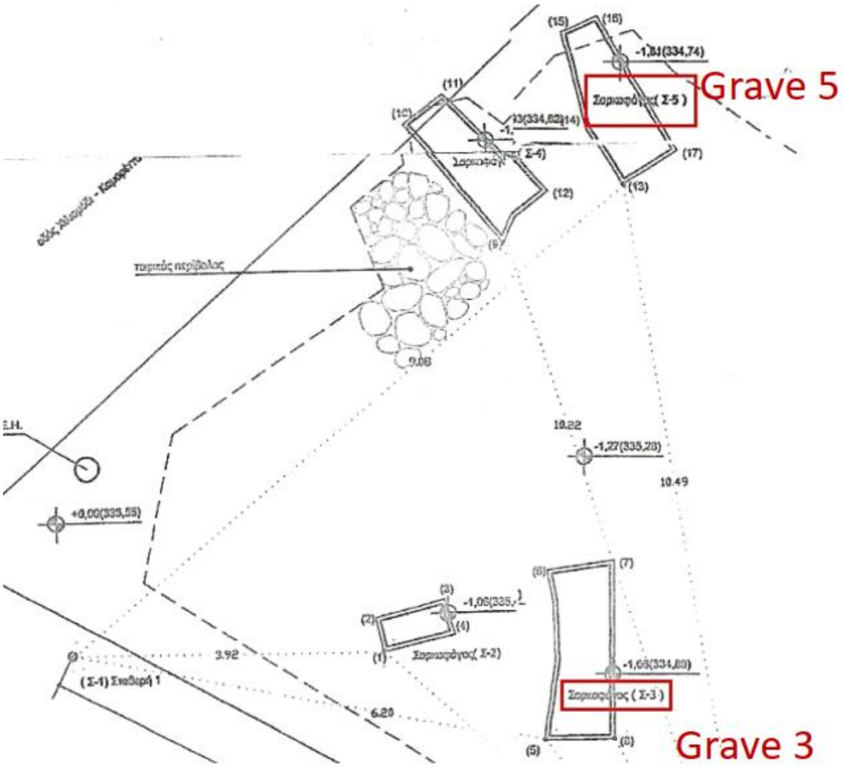


**Supplementary Figure S31.** The location of graves 3 and 5 in the Faneromeni-Kamareta site of Ancient Tenea.

##### 3.5.4 Runs of homozygosity

Analyses of per individual Runs Of Homozygosity (ROH) levels were performed using the hapROH v.64 [90] software on the 1240K pseudo-haploid data in EIGENSTRAT format. We applied this method to individuals covering more than 300000 [91,92] sites on the 1240K panel (21/26). We followed the proposed pipeline presented in the Jupyter Notebook file called callROH_vignette.ipynb (https://www.dropbox.com/sh/eq4drs62tu6wuob/AABM41qAErmI2S3iypAV-j2da?dl=0), using the 1000 Genomes Project as a reference panel (in hdf5 format). The output of HapROH, as well as the script are available at https://doi.org/10.5281/zenodo.10848927.

**Figure 5** displays the total length of ROH segments exceeding 4 cM that were detected in all the individuals analyzed here. The segments are categorized into four bins based on their length: 4-8 cM, 8-12 cM, 12-20 cM, and >20 cM. Shorter ROH segments in the 4-8 cM bin indicate a small population size, while longer segments in the >20 cM category suggest an isolated population and/or consanguinity practices. Individuals with a total of more than 50 cM of ROHs in the >20 cM category are considered possible offsprings of close kin [90]. **Supplementary Figure S32** presents a linear regression scatterplot showing the relationship between the total sum of ROHs and the genomic coverage of each individual, along with Pearson's r correlation coefficient. The p-value > 0.01 indicates no statistically significant correlation between detected ROHs and genomic coverage, indicating that the detected ROHs are not affected by sequencing depth.

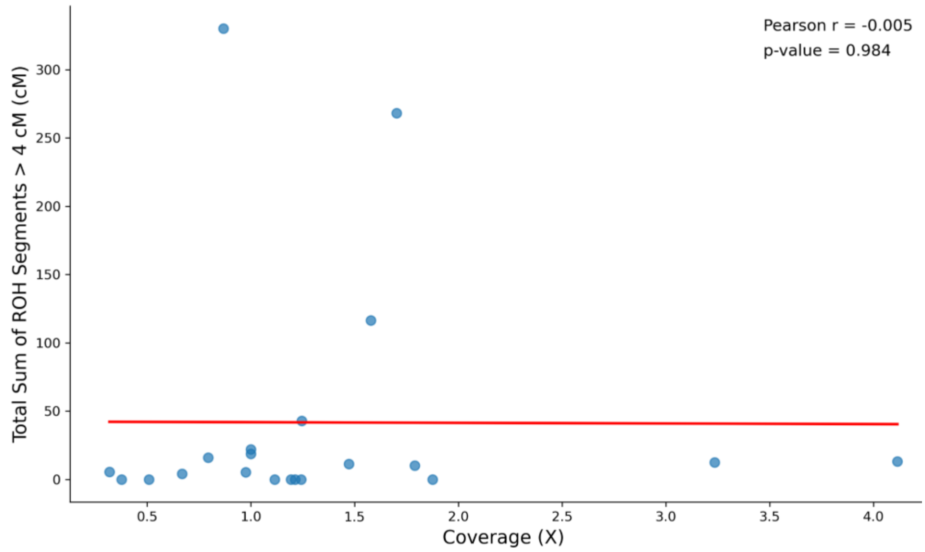


**Supplementary Figure S32.** Linear regression scatterplot showing the relationship between the total sum of ROHs and the genomic coverage of each newly sequenced individual. Estimated Pearson's r correlation coefficient, as well as p-value shown in top right corner.

##### 3.5.5 Imputation and Identity-by-Descent segments screening

We imputed the newly produced ancient genomes of this study by using the 1000 Genomes phase 3 [61] dataset as a reference and GLIMPSE v.1.1.1 [93]. We first reduced the 1kGP sites list by removing singletons (uninformative for imputation), and keeping only biallelic SNPs with the BCFtools v.1.14 view (`-m 2 -M 2 -c 2`) module as per [94]. This resulted in a total of 43285119 sites. Then, to generate genotype likelihoods, we used the

ATLAS pipeline v.0.9 [95], by estimating *post-mortem* damage (`task=PMD`) and calculating genotype probabilities (`task=call method=MLE`). The *GLIMPSE_chunk* function was used to create smaller genomic chunks [window size: 2000000 and buffer-size: 200000 [94]] and *GLIMPSE_phase* to perform imputation for each chunk. For this step, chromosomal VCF files and a genetic map are required. We used the VCF files produced by ATLAS containing the ~43 million sites and the HapMap phase II NCBI *b37* genetic map [96]. Finally, the chunks were ligated and phased with *GLIMPSE_ligate* and *GLIMPSE_sample*, respectively**.**

To infer shared Identity By Descent (IBD) segments between pairs of individuals in our dataset, we used the ancIBD v.0.5 tool [97], following the recommended by the developers processing procedure for our newly sequenced WGS data with at least 0.25× coverage (22 out of 26; Amv_Epi_Cl_3, Amv_Epi_Hel_2, Amv_Epi_Hel_5, and Ten_Pel_Rom_4 were excluded). Firstly, as ancIBD parameters are optimized for the 1240K list of genomic sites, the phased/imputed data were reduced only to those sites. In addition the VCF files were transformed into hdf5 format. Both of these processes were conducted using the *ancIBD.IO.prepare_h5.vcf_to_1240K_hdf* function. Then, the IBD segments were called using the *hapBLOCK_chroms* function with standard parameters used (`l_model='h5'`, `e_model='haploid_gl2'`, `h_model='FiveStateScaled'`, `t_model='standard'`, `p_col='variants/AF'`, `ibd_in=1`, `ibd_out=10`, `ibd_jump=400`, `min_cm=6`, `cutoff_post=0.99`, `max_gap=0.0075`), as proposed by the software developers. Finally, the create_ind_ibd_df function was used to create summary data for the pairwise shared IBD segments and to perform a quality control filtering, by removing IBD segments of low SNP density (IBD segments with less than 220 SNPs per centiMorgan). The inferred IBD segments are categorized by length into four groups: 8-12 cM, 12-16 cM, 16-20 cM, and greater than 20 cM. Segments longer than 20 cM indicate closer relatedness, as only a few meiotic events are required to break up such long segments, whereas the shorter segments represent genealogical connections that are a few hundred years deep [97]. The output of ancIBD, as well as the script used are provided in https://doi.org/10.5281/zenodo.10848927. The number of shared IBDs within each of the four length bins, in a pairwise fashion, is presented in **Figure 3**.

Shared IBDs segments for the >20 cM bin indicate a) the individuals determined by READ and KIN (see **Section 3.5.3** above) to have a 1st- (Amv_Epi_Cl_-5 and -6; Amv_Epi_Hel_-3 and -4; Ten_Pel_Arch_-1 and -2) and 2nd-degree (Amv_Epi_Cl_-5 and -6 with Amv_Epi_Cl_1) genetic relationship (yellow and green colors in **Figure 3A; upper-right plot**) as being closely related, b) a relationship of intermediate genetic kinship (a few generations apart) among three Roman Tenea individuals (Ten_Pel_Rom_-1, -2, and -3; petrol colors in **Figure 3A; upper-right plot**), and c) distant kin relationships (several generations apart) between Archaic and Classical Amvrakia individuals (Amv_Epi_Arch_3 with Amv_Epi_Cl_-5 and -6; Amv_Epi_Arch_2 with Amv_Epi_Cl_4), between Archaic Amvrakia individuals (Amv_Epi_Arch_-1 and -3), and between Classical Amvrakia individuals (Amv_Epi_Cl_-4 and -6). Shared IBD segments for the 16-20 cM bin (**Figure 3A; bottom-left plot)**, in addition to the above relationships, also indicate some degree of distant genetic relationship between other pairs of Classical Amvrakia individuals (Amv_Epi_Cl_1 with Amv_Epi_Cl_-6 and -5, respectively). Shared IBD segments in the 12-16 cM and 8-12 cM bins, also indicate the presence of strong ancestral genetic links (yellow, green, petrol, and blue colors in **Figure 3B**) a) between Archaic Amvrakia and Archaic Ternea (Amv_Epi_Arch_-1,-2, and -3 with Tenea_Pel_Arch_-1 and -2 combination of pairs), b) between Classical Amvrakia and Archaic Tenea (Amv_Epi_Cl_-2 and -4 with Tenea_Pel_Arch_-1 and -2 combination of pairs), c) between Classical Amvrakia and Hellenistic Tenea (Amv_Epi_Cl_1 with Tenea_Pel_Hel_1), d) between Archaic Tenea and Hellenistic Tenea (Tenea_Pel_Arch_2 with Tenea_Pel_Hel_1), e) between Classical Amvrakia and Hellenistic Amvrakia (Amv_Epi_Cl_-1 and -4 with Amv_Epi_Hel_-3 and -4, respectively), and f) between Hellenistic Amvrakia and Archaic Amvrakia (Amv_Epi_Cl_4 and Amv_Epi_Arch_1). Notably, the absence of shared IBD segments is observed a) between LBA Ammotopos and any other sample and b) between Roman Tenea and any other non-Roman sample from either Tenea, or Amvrakia.

##### 3.5.6 Merging with public data

The 26 ancient individuals were studied at the population level in the context of previously published data for modern Western Eurasians individuals and prehistoric as well as historic (Iron age to Roman times) ancient individuals. The rationale for this sample selection is a) the inclusion of the spatiotemporally most closely related populations to our newly sequenced individuals and b) the inclusion of more distantly related populations that are required for specific, additional analyses (see below for details). Note that, different analysis types and methods required assembling distinct datasets from this data pool. Details on which sample was used in each analysis (and under which group label) can be found in **Supplementary Table 4** (**Additional file 4**). The details of each dataset assembly are provided in the following:

<u>**“Dataset 1”**: 663 ancient individuals (published)</u>

This dataset included a) 542 ancient individuals [98–125] whose data derive from the ‘1240K’ (1233013 sites) SNP capture essay [83] downloaded from the Allen Ancient DNA Resource (Version 8; aadr_v.54.1.p1_1240K_public; [79]) in PACKEDANCESTRYMAP format, b) 1240K data from 111 ancient individuals [126,127] that at the time of manuscript preparation had not yet been integrated into the AADR database (https://reich.hms.harvard.edu/sites/reich.hms.harvard.edu/files/inline-files/Reitsema2022PNAS_Ancient_1240K.zip; https://figshare.com/projects/Genotype_data_for_103_individuals_from_study_Ancient_DNA_reveals_admixture_history_and_endogamy_in_the_prehistoric_Aegean_/156152; PACKEDANCESTRYMAP format), and c) WGS data from 10 individuals generated by Koptekin et al. [84] (not in the AADR database, either), which we processed, starting from raw FastQ data (obtained after personal communication with the authors), using the same pipeline as the one used for the 26 samples of the present study (the mapache samplelist file for these 10 individuals is available at https://doi.org/10.5281/zenodo.10848927). All individuals fulfill the following selection criteria: a) they were genetically unrelated (above >2nd genetic kinship degree), b) they covered at least 100000 positions of the 1240K list, and c) their contamination assessment in AADR was not tagged as “QUESTIONABLE”. We merged data from these distinct sources into a single PACKEDANCESTRYMAP file using the EIGENSOFT *mergeit* function. This dataset was used in F-statistics analyses (see below).

<u>**“Dataset 2”**: 657 ancient individuals</u>

This is the same as “Dataset 1” albeit excluding six Upper Paleolithic Iberomaurusian hunter-gatherers from Tafolrat, Morocco. This dataset was used in the ADMIXTURE analysis (see below). The samples from Tafolrat were excluded from this analysis as two-thirds of their ancestry originates from sub-Saharan Africa [121].

**“Dataset 3”**: 663 ancient and 888 modern individuals

This dataset included a) 888 modern West Eurasian individuals genotyped on the Human Origins SNP (HO; 597573 sites) array [83]; most of the HO data were downloaded from the Allen Ancient DNA Resource (Version 8; aadr_v.54.1.p1_HO_public; [79]) and the rest are available in [127], both in the PACKEDANCESTRYMAP format, and b) the data from the ancient individuals of “Dataset 1”, albeit restricted to the HO sites. Individuals that were either tagged as outliers (_o suffix), as being genetically related, as *to-be-ignored*, and as having been obtained via whole genome amplification (_*wga* suffix) were omitted. This dataset was used in the PCA analysis (see below).

#### 3.5.7 Principal Component Analysis

Principal Component Analysis (PCA) was used to summarize the relationships among our 26 ancient samples in the context of the previously published ancient and modern genomes of “Dataset 3”. PCA was performed using the EIGENSOFT *smartpca* function, default parameters, and the `lsqproject: YES` and `numoutlieriter: 0` options, in order to project the data of the ancient samples onto the PCs calculated for the modern samples, and also, to disable outlier checking and removal, respectively. The stability of the PCA was assessed using Pandora v.2.0.0 [128] using `n_replicates: 100` as suggested by the developers and `kmeans_k: 4` and supplying the smartpca_optional_settings field in the configuration file with the aforementioned projection-related options above. Note that the kmeans_k setting is of minor importance as we mainly focus on the stability of individuals in the PCAs conducted on the 100 bootstrap replicates. The Pandora Stability estimate was 0.95 (convergence achieved after 25 pseudo-replicates), with an average ± standard deviation support value of 0.89 ± 0.07, and a median of 0.90. Regarding the newly sequenced individuals, their support values were ranging between 0.70 and 0.86, with the exception of the following four individuals: Amv_Epi_Cl_3 (0.67), Amv_Epi_Hel_1 (0.65), Amv_Epi_Hel_2 (0.53), and Ten_Pel_Rom_4 (0.67). Their instability in PCA placement is shown in **Supplementary Figure 33**. Their lack of stability is due to coverage as three out of these four genomes are among the four least covered ones (<0.15×), whereas all of them are among the six genomes with the lowest mean depth coverage (<0.32×).

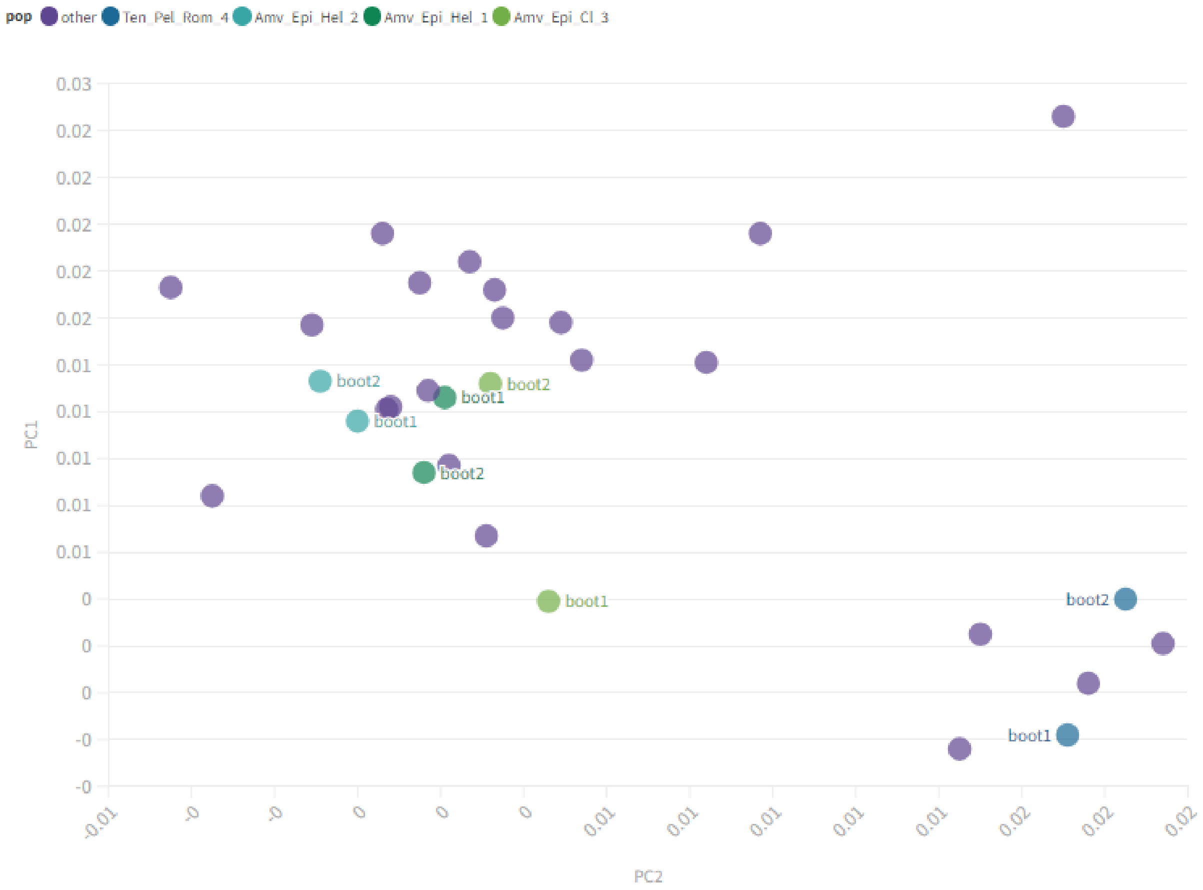


**Supplementary Figure S33.** Plot of two bootstrapped PCA replicates created using Pandora showing the variability in placement of the four newly sequenced individuals with a low Pandora Stability estimate (<0.70). For clarity we plot only the individuals of the present study and both of the replicates only for the four focal individuals.

The grouping of samples, which is given in **Supplementary Table 4** (**Additional file 4**), reflects the spatiotemporal and cultural origin of the samples, meaning that each group is a combination of a) present-day countries name (3-digit codes) and b) relative archaeological period (e.g. Neolithic, Late Bronze Age, Archaic, Classical etc). In all cases we have also added the actual temporal range covered. In order to provide clarity to the plot and as we wanted to focus more on the temporal space of the newly sequenced individuals (LBA to Roman times) we merged some earlier (prehistoric) samples into single, genetically homogeneous, groups. As a result of the above, the grouping for the PCA analysis includes groups, such as GRC_Neolithic_6400-3600BCE, GRC_EBA_EMBA_MBA_2900-1700, and GRC_LBA_1700-1050BCE, which all three include, both, mainland and insular populations. However, we kept GRC_Mainland_WMakedonia_MBA_2100-1600BCE as a separate group as its genetic ancestry is quite different from the rest of the Greek MBA individuals [100]. In the cases of other Balkan countries excluding Greece, we merged the prehistoric samples into South_Balkans_EBA_MBA_MLBA_3350-1100BCE (Albania, Bulgaria, North Makedonia) and NW_Balkans_EBA_MBA_MLBA_2000-800BCE (Croatia, Serbia, Montenegro). Historical times were not merged together (see e.g. MKD_IA_900-500BCE and MKD_Classical_Hellenistic_500-50BCE), whatsoever. Other merged groups are Hunter_Gatherers_22600-5500BCE (all Paleolithic and Mesolithic HGs), IRN-IRQ_Neolithic_Chalcolithic_9500-3500BCE (Iran/Iraq Neolithics and Chalcolithics), ISR-

JOR_Epip-Natufian_and_PPN_12000-6200BCE (Natufians and Pre-pottery Neolithics from the Levant), and RUS-BGR-SER_BA-Yamnaya-like_3500-1600BCE (individuals with significant Yamnaya-like ancestry). In the case of Italy we divided the groups geographically in Mainland, Sicily, and Sardinia, whereas we also separated the Etruscans (ITA_Mainland_Etruscan_800-001BCE) from other mainland contemporary to them individuals (ITA_Mainland_IA_and_RomanRepublic_760-003BCE). In the case of Türkiye, we divided the samples temporarily, but in some cases that the samples of a given period do not cover the entire area of Türkiye we also mention the regions (see e.g. TUR_South-SouthWest_MBA_MLBA_2000-1200BCE, TUR_West_Archaic_750-480BCE). Note that this grouping does not affect the results and is used only for better visualization.

The result of the PCA projection for the first two PCs is presented in **Figure 2A.** According to the results, the newly generated genomes are, in general, clustered together and overlap with other ancient eastern Mediterranean genomes, albeit they do display considerable variation. Two main clusters are formed, with three additional samples standing out.

The first cluster includes the majority of the Amvrakia samples (one of the three Archaic, all six Classical, and four out of five Hellenistic), the two BA Ammotopos samples and some of the Tenea samples (both of the two Archaic and two out of three Hellenistic). This cluster overlaps mostly with LBA (1700-1050 BCE) genomes from the entire present-day area of Greece, EIA (1100-500 BCE) genomes from present-day area of Bulgaria, as well as with the local population of the Ancient Greek colony of Himera in Sicily (780-400 BCE). Moreover, some additional samples are also placed close to this cluster including a Greek EBA_MBA (2800-1700 BCE) individual and a Greek IA individual from the Peloponnese (1070-800 BCE). Close, but not overlapping, several other genomes are observed, including most of the Greek EBA_MBA (2800-1700 BCE) and the rest of the Greek IA (1070-800 BCE and 800-500 BCE, respectively), a few Roman Imperial individuals from the Italian mainland (1-530 CE), a few from the South Balkans EBA_MBA_MLBA (3350-1100 BCE), and two more from the IA (900-500 BCE) and Classical-Hellenistic (500-50 BCE) present-day area of North Makedonia.

The second cluster includes all four Roman Tenea individuals, as well as one of the three Hellenistic Amvrakia individuals. This cluster, in comparison to the first one, is placed slightly more towards the bottom left of the PC space and the Iran-Neolithic/CHG - WHG axis. It mostly overlaps with Archaic (750-480 BCE), Roman (27-476 CE), and other Anatolian genomes [including a Hellenistic (510-30 BCE) and an EBA (3350-2000 BCE)], a Roman Imperial (1-530 CE) genome from Italy, as well as a few Greek genomes from the Roman (250-400 CE), the EBA_MBA (2900-1700 BCE), and the LBA (1700-1050 BCE) times. Close, but not overlapping, are a few additional Greek EBA_MBA (2900-1700 BCE) and Italian Roman Imperial (1-530 CE) genomes, as well as some EBA (3350-2000 BCE) and Hellenistic (510-30 BCE) genomes from the present-day area of Türkiye.

Between these two clusters, the remaining of the two Hellenistic Tenea individuals is placed, with close affinities to a local individual of the Ancient Greek colony of Himera, Sicily (780-400 BCE) and an IA individual from the Peloponnese (1070-800 BCE).

Finally, the two remaining Archaic Amvrakia individuals are closer to the first cluster, but more differentiated towards the upper left of the PC space and the Levant Neolithic - BA-Yamnaya-like axis. One of them, the closest to the first cluster, entirely overlaps with an Etruscan individual from Italy (800-001 BCE) and is surrounded by Greek genomes mostly from the EBA_MBA (2900-1700 BCE), and the LBA (1700-1050 BCE). The other Archaic Amvrakia individual is placed relatively far away from the first cluster, in close proximity to

EBA_MBA (2900-1700 BCE) Greece and a few Greek Neolithic (6400-3600 BCE) and Anatolian Neolithic (9000-5600 BCE) individuals.

##### 3.5.8 Population clustering analysis (ADMIXTURE)

We performed unsupervised ADMIXTURE v.1.3.0 [129] analysis using “Dataset 2” in order to examine the broader patterns of ancestry of our newly generated genomes in comparison to other ancient individuals. In preparation for this analysis, we used Plink v.1.9 [130] for dataset manipulations and pruned the dataset based on three different linkage disequilibrium $r^2$ thresholds (0.40, 0.60, and 0.80) on a sliding window of 200 kbp with a step of 25 variant counts (`--indep-pairwise 200 25 0.4/0.6/0.8`), as well as three allele missingness (`--geno`) thresholds (40%, 60%, and 80%), resulting in distinct final SNP counts per dataset (**Additional Table A1**). Before LD pruning and allele missingness filtering, we removed the newly sequenced samples *Amv_Epi_Cl_5*, *Amv_Epi_Cl_4*, *Amv_Epi_Hel_3*, and *Ten_Pel_Arch_1* as they have a 1st-/2nd-degree genetic kinship with individuals that exhibit higher coverage in the dataset. We performed the analysis for K=2-10, using the `--haploid=”*”` flag given that our data are pseudohaploidized and we determined the best K value and LD and allele missingness thresholds by using ADMIXTURE’s internal block jackknife routine to estimate cross validation errors (**Supplementary Figure S34**). The script is provided in https://doi.org/10.5281/zenodo.10848927 and includes the following R packages: argparse v.2.2.3 [131], doMC v.1.3.8 [132], foreach v.1.5.2 [133], ggh4x v.0.2.8 [134], ggthemes v.5.0.0 [135], gridExtra v.2.3 [136], reshape [137], and stringr v.1.5.1 [138].

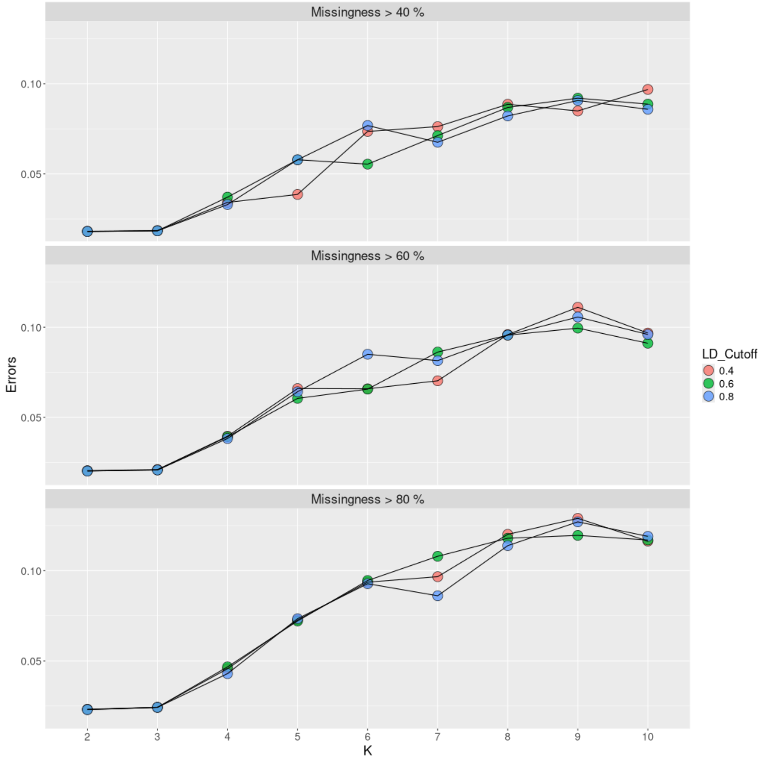


**Supplementary Figure S34.** ADMIXTURE block jackknife cross validation errors for K=2-10 and a linkage disequilibrium $r^2$ threshold (LD_Cutoff) of 0.40, 0.60, and 0.80, respectively. SNPs whose presence was below the respective missingness threshold, as well as SNPs exceeding the LD threshold on a sliding window of 200 kbp with a step of 25 variant counts, were removed.

**Additional Table A1.** Resulting number of SNPs after filtering the dataset used for ADMIXTURE analysis using distinct linkage disequilibrium (LD) and missingness thresholds.

| LD threshold ($r^2$) | Allele missingness threshold (%) | # SNPs |
|---|---|---|
| 0.80 | 80 | 919167 |
| 0.80 | 60 | 719225 |
| 0.80 | 40 | 434297 |
| 0.60 | 80 | 913376 |
| 0.60 | 60 | 717163 |
| 0.60 | 40 | 433756 |
| 0.40 | 80 | 858751 |
| 0.40 | 60 | 694490 |
| 0.40 | 40 | 426668 |

We observed similar results among the pruning/filtering options for K=2-4, albeit the allele missingness threshold of 80% was slightly worse than the 60% and 40%. For higher values, an allele missingness filter of 40% seems to marginally outperform the other two configurations regardless of the selected LD cutoff. The fewest errors were observed for K=2 and K=3. All resulting ADMIXTURE plots are provided in https://doi.org/10.5281/zenodo.10848927. The resulting plots of ADMIXTURE analysis using an LD threshold of 0.60 and an allele missingness threshold of 40% are presented in **Figure 2B** (K=3; only historical individuals plotted) and in **Supplementary Figure S35 (A-I)** (K=2-10; all individuals plotted). The grouping of samples under the same population label is given in **Supplementary Table 4** (**Additional file 4**)**.** Here, we applied an analogous grouping procedure as for the PCA analyses, with the following modifications: a) we further subdivided LBA to Roman times Greece geographically, in order to separate Crete, the Cyclades, and the Peloponnese from the mainland, b) we subdivided the Hunter Gatherers (into WHG, CHG, etc.), and c) we divided the Natufians and the pre-pottery Neolithics of the Levant. Note that this grouping does not affect the analytical results and is merely used for better visualization.

When focusing on K=3, which is the lowest K that allows to differentiate genetic clusters associated with three key European ancestral components, namely, Western hunter-gatherers (WHGs), early European farmers (EEFs), and Caucasus hunter-gatherers (CHGs), depicted with red, orange, and blue color, respectively in **Figure 2B** and **Supplementary Figure S35B**, our samples appear to comprise all three of the above ancestral components. Most of our samples (excluding the Roman Tenea ones), have ancestry proportions that resemble those of previously published LBA and IA genomes from mainland Greece: a high EEFs proportion, followed by a lower CHGs proportion, and a small WHGs proportion. The Roman Tenea genomes display a higher CHGs proportion than the rest of our samples, a reduced EEFs proportion, and an analogous WHGs proportion, thus resembling other published Roman genomes from Greece and Italy.

## A. K=2

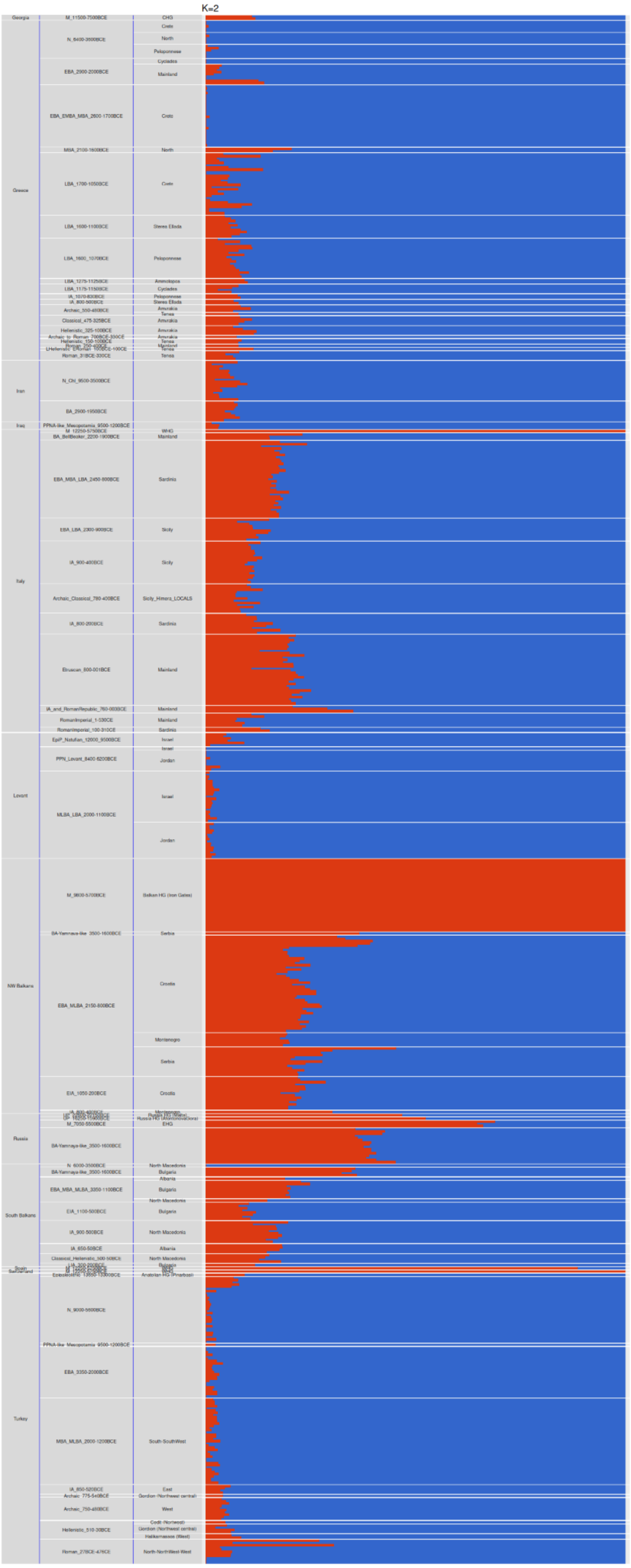

K=2
Georgia
Greece
Iran
Iraq
Italy
Levant
NW Balkans
Russia
South Balkans
Turkey

**B. K=3**

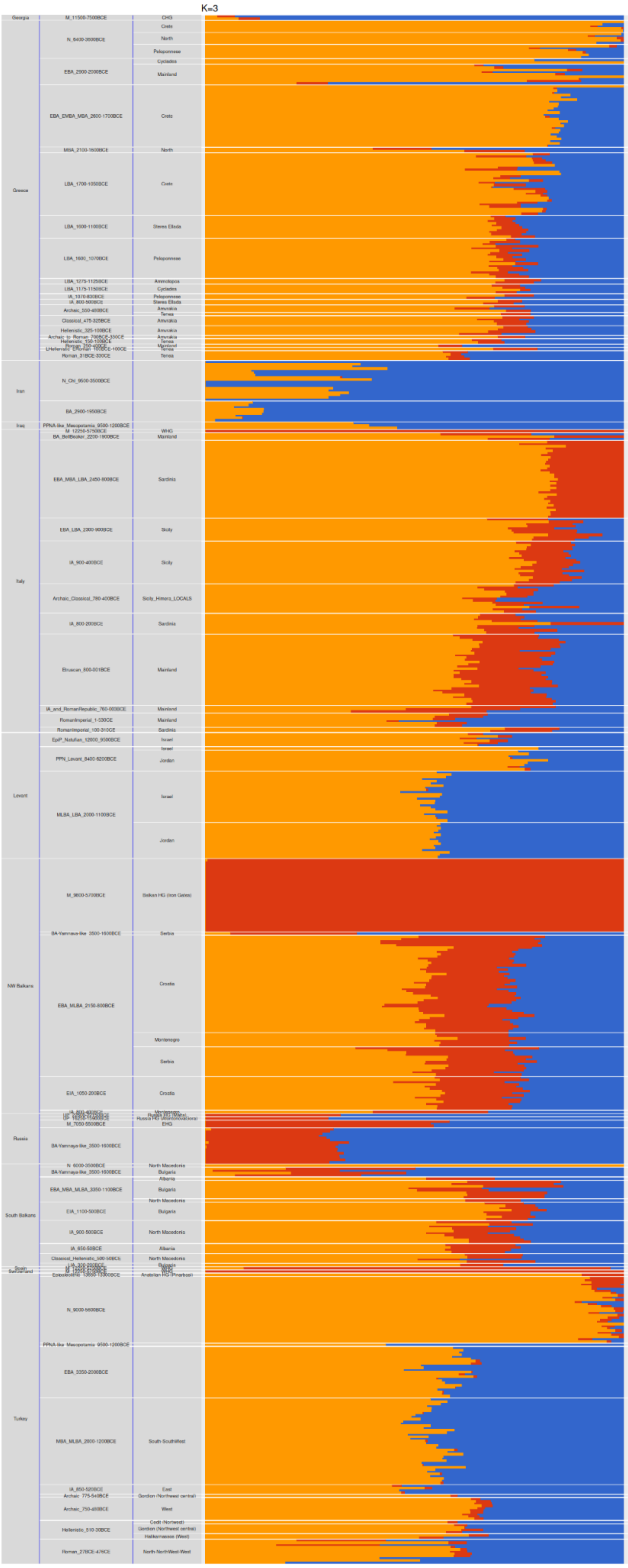

**C. K=4**

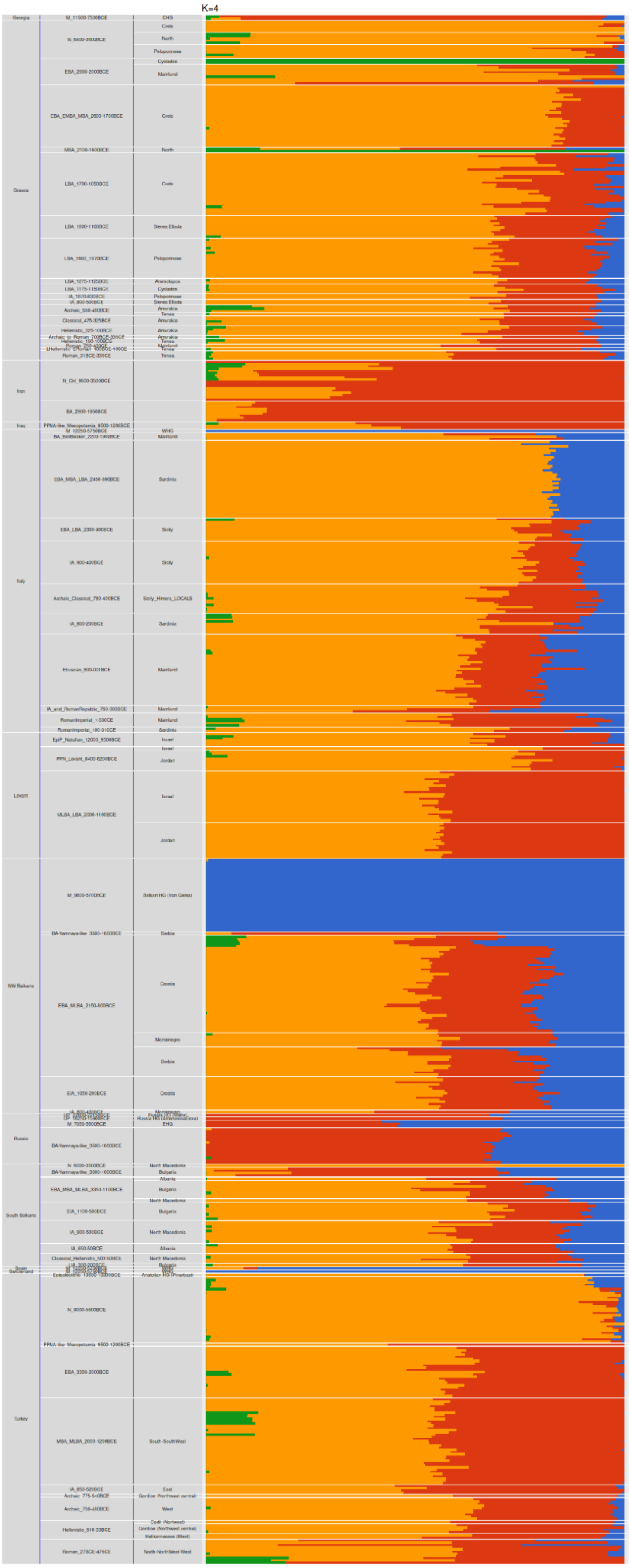
K=4
Georgia
Greece
Iran
Iraq
Italy
Levant
NW Balkans
Russia
South Balkans
Turkey

**D. K=5**

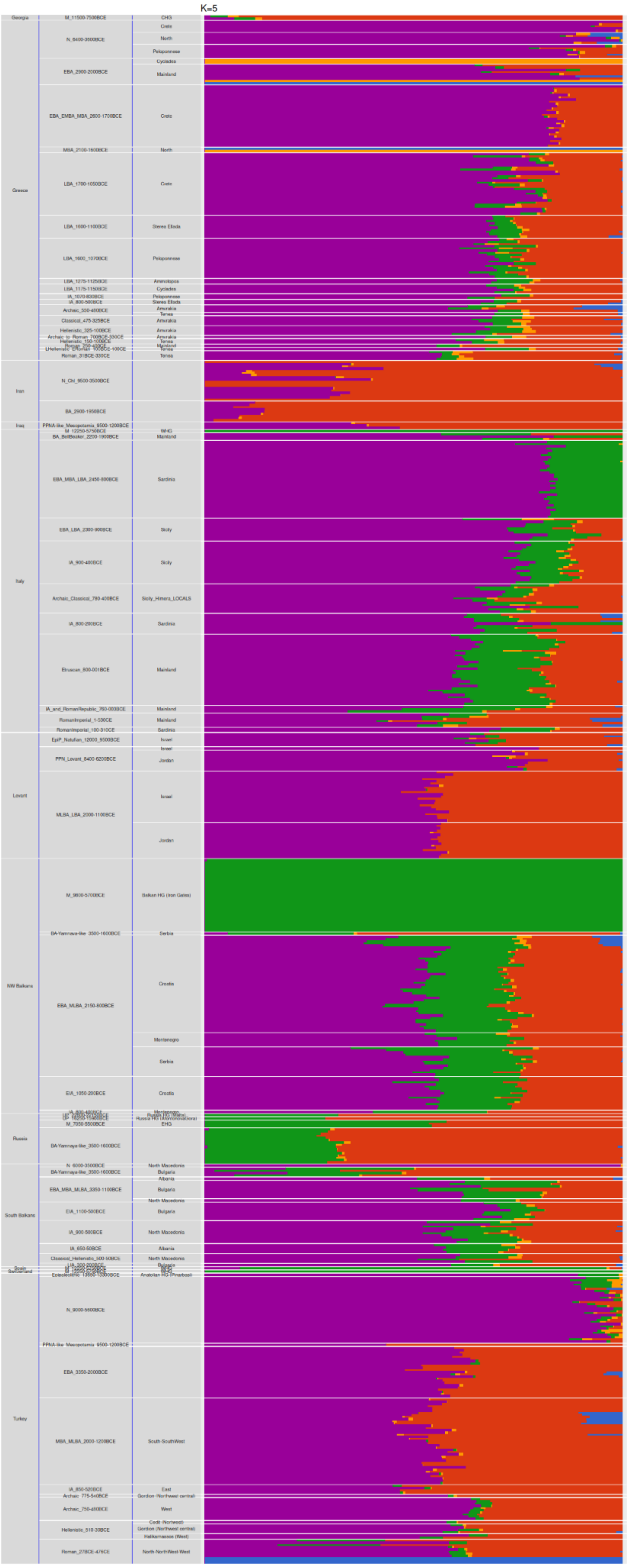

K=5
Georgia
Greece
Iran
Iraq
Italy
Levant
NW Balkans
Russia
South Balkans
Turkey

## E. K=6

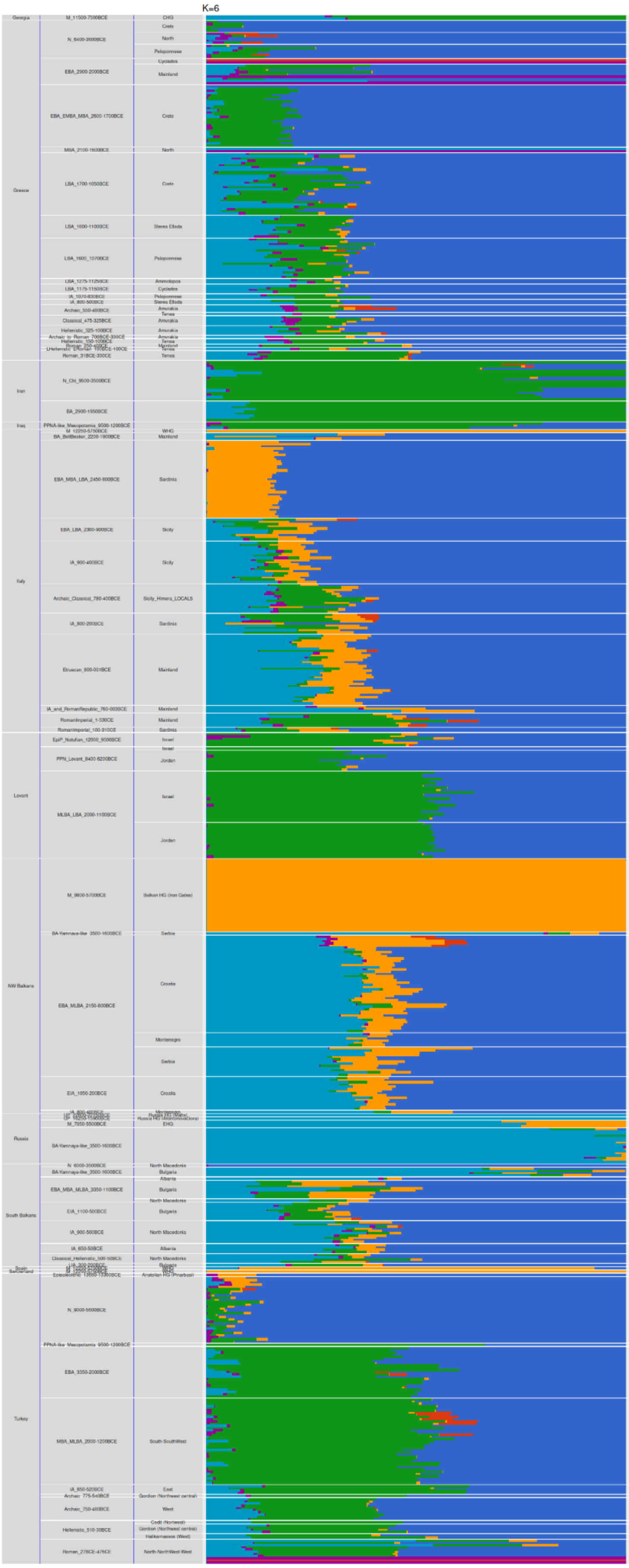
K=6
Georgia
M_11500-7500BCE
CHG
Greece
N_6400-3500BCE
Crete
North
Peloponnese
EBA_2900-2000BCE
Cyclades
Mainland
EBA_EMBA_MBA_2600-1700BCE
Crete
MBA_2100-1600BCE
North
LBA_1700-1050BCE
Crete
LBA_1600-1100BCE
Sterea Ellada
LBA_1600_1070BCE
Peloponnese
LBA_1275-1125BCE
Ammotopos
LBA_1175-1150BCE
Cyclades
IA_1070-830BCE
Peloponnese
IA_800-500BCE
Sterea Ellada
Archaic_550-480BCE
Amvrakia
Tenea
Classical_475-325BCE
Amvrakia
Hellenistic_325-100BCE
Amvrakia
Archaic_to_Roman_700BCE-330CE
Amvrakia
Hellenistic_150-100BCE
Tenea
Roman_31BCE-330CE
Tenea
Iran
N_Chl_9500-3500BCE
BA_2900-1950BCE
Iraq
PPNA-like_Mesopotamia_9500-1200BCE
WHG
BA_BellBeaker_2200-1900BCE
Mainland
Italy
EBA_MBA_LBA_2450-800BCE
Sardinia
EBA_LBA_2300-900BCE
Sicily
IA_900-400BCE
Sicily
Archaic_Classical_780-400BCE
Sicily_Himera_LOCALS
IA_800-200BCE
Sardinia
Etruscan_800-001BCE
Mainland
IA_and_RomanRepublic_760-000BCE
Mainland
RomanImperial_1-530CE
Mainland
RomanImperial_100-310CE
Sardinia
Levant
EpiP_Natufian_12000_9500BCE
Israel
PPN_Levant_8400-6200BCE
Israel
Jordan
MLBA_LBA_2000-1100BCE
Israel
Jordan
NW Balkans
M_9800-5700BCE
Balkan HG (Iron Gates)
BA-Yamnaya-like_3500-1600BCE
Serbia
EBA_MLBA_2150-800BCE
Croatia
Montenegro
Serbia
EIA_1050-200BCE
Croatia
Russia
M_7050-5500BCE
EHG
BA-Yamnaya-like_3500-1600BCE
South Balkans
N_6000-3500BCE
North Macedonia
BA-Yamnaya-like_3500-1600BCE
Bulgaria
Albania
EBA_MBA_MLBA_3350-1100BCE
Bulgaria
North Macedonia
EIA_1100-500BCE
Bulgaria
IA_900-500BCE
North Macedonia
IA_650-50BCE
Albania
Classical_Hellenistic_500-50BCE
North Macedonia
Turkey
N_9000-5600BCE
PPNA-like_Mesopotamia_9500-1200BCE
EBA_3350-2000BCE
MBA_MLBA_2000-1200BCE
South-SouthWest
IA_850-520BCE
East
Archaic_775-540BCE
Gordion (Northwest central)
Archaic_750-480BCE
West
Hellenistic_510-30BCE
Cedit (Northwest)
Gordion (Northwest central)
Halikarnassos (West)
Roman_27BCE-476CE
North-NorthWest-West

## F. K=7

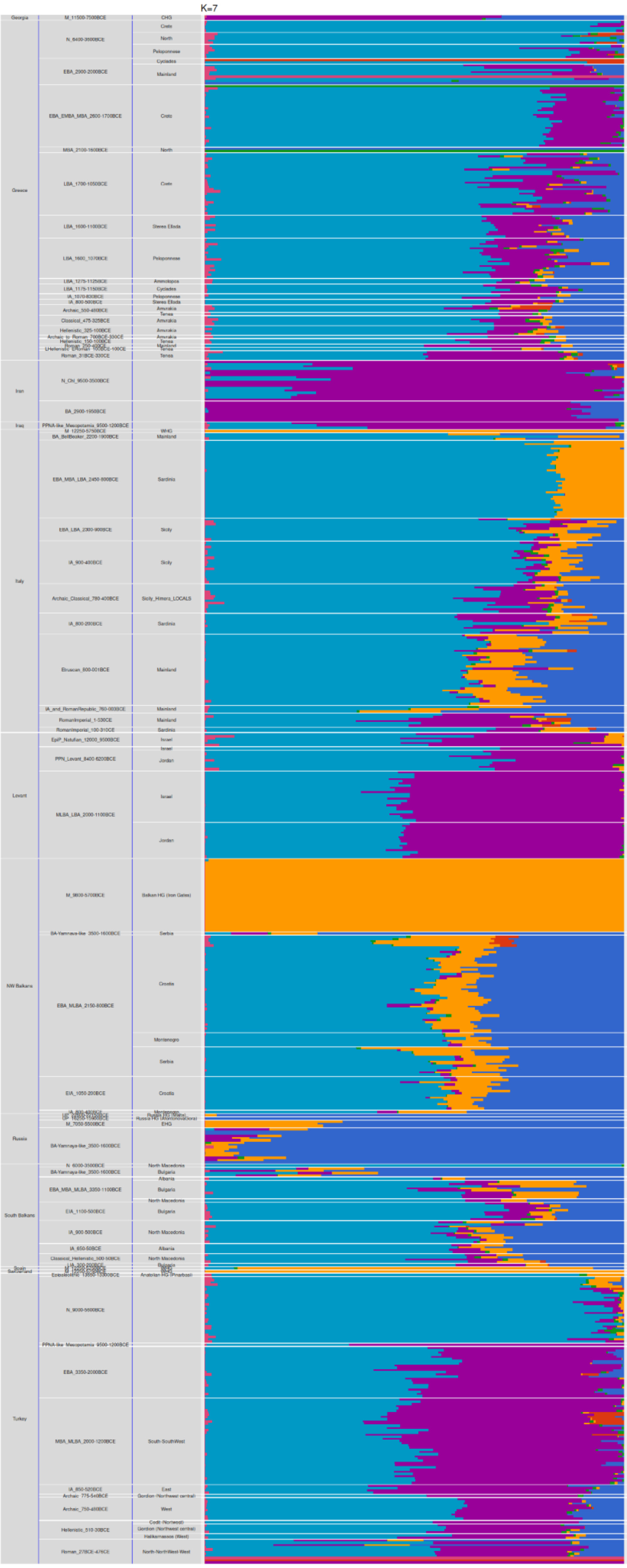
K=7
Georgia
Greece
Iran
Iraq
Italy
Levant
NW Balkans
Russia
South Balkans
Turkey

## G. K=8

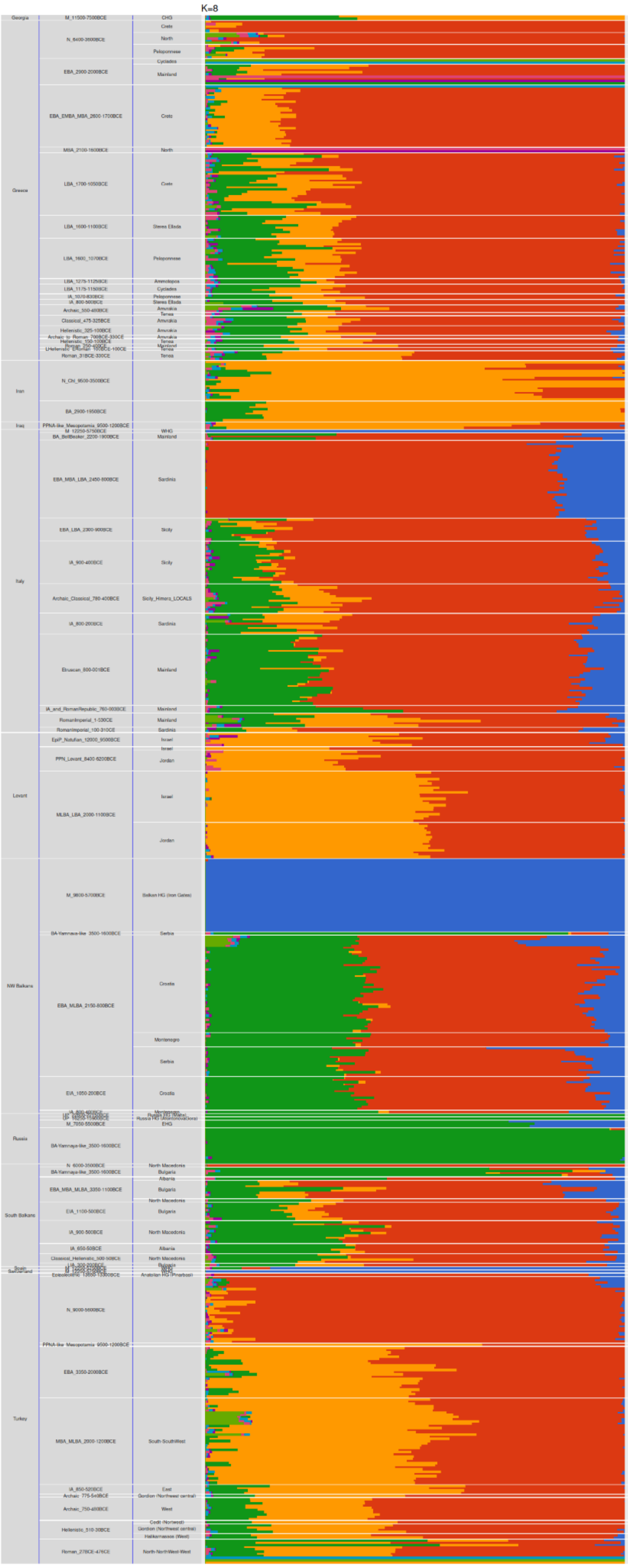

## H. K=9

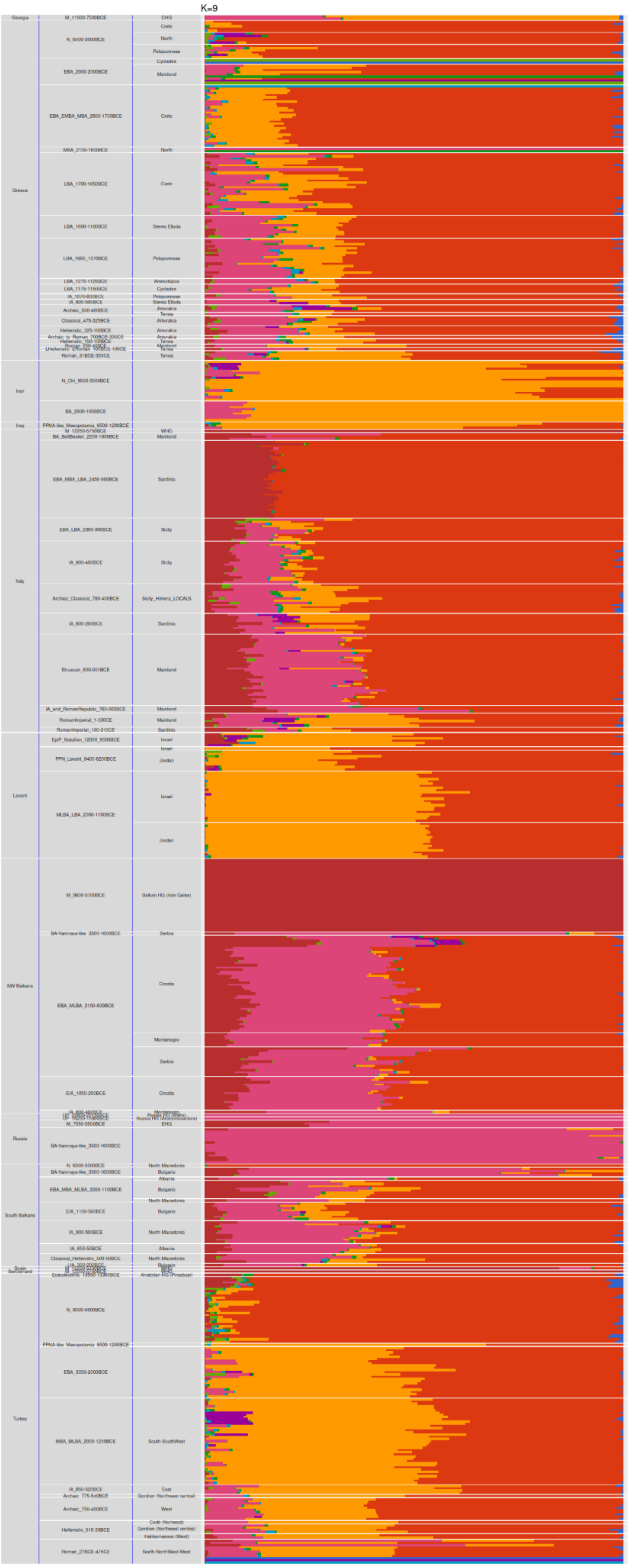

**I. K=10**

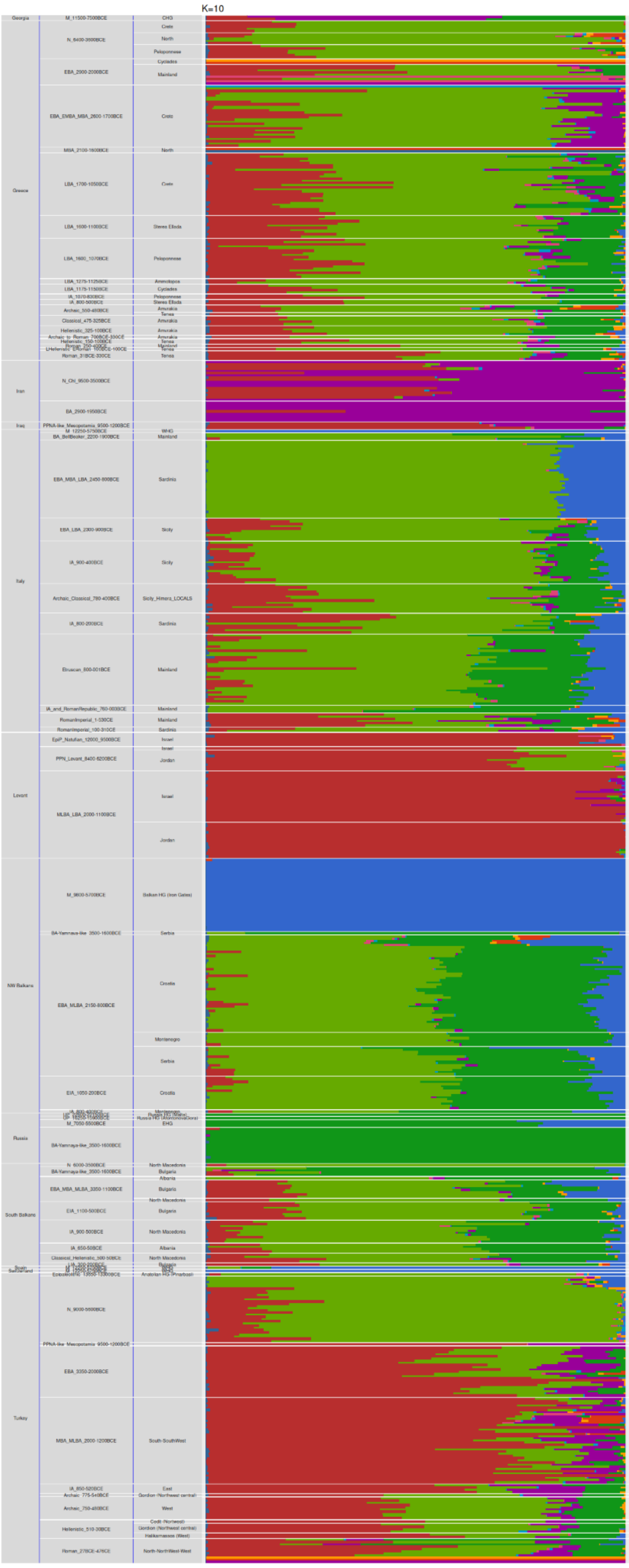


**Supplementary Figure S35.** ADMIXTURE analysis of 22 newly generated ancient genomes of the present study (four individuals were excluded due to genetic kinship) and 657 other ancient published genomes (“Dataset 1”) using K=2-10 (**A-I plots**), an LD $r^2$ threshold of 0.60 and an allele missingness threshold of 40%. The analysis was based on 433756 genomic sites of the 1240K list.

##### 3.5.9 f3 Statistics and Ancestry Proportion Analysis (qpAdm)

**Benchmarking and simulation analysis**

Admixture *f3 simulation analysis*

Interestingly, when we calculated Admixture f3 values for all plausible population triplets, the resulting values were all positive (see details and results of the analysis below). Such a behavior is interesting because historically our sampled populations should have exchanged migrants extensively. Using simulations, we examined whether Admixture f3 values might have been affected by processes, such as the gene flow (known as migration rate in population genetics; not to be confused with migration in the archaeological sense) between the source populations and/or population size changes in the target (admixed) population. We employed a scenario with eight populations as depicted in **Supplementary Figure S36**.

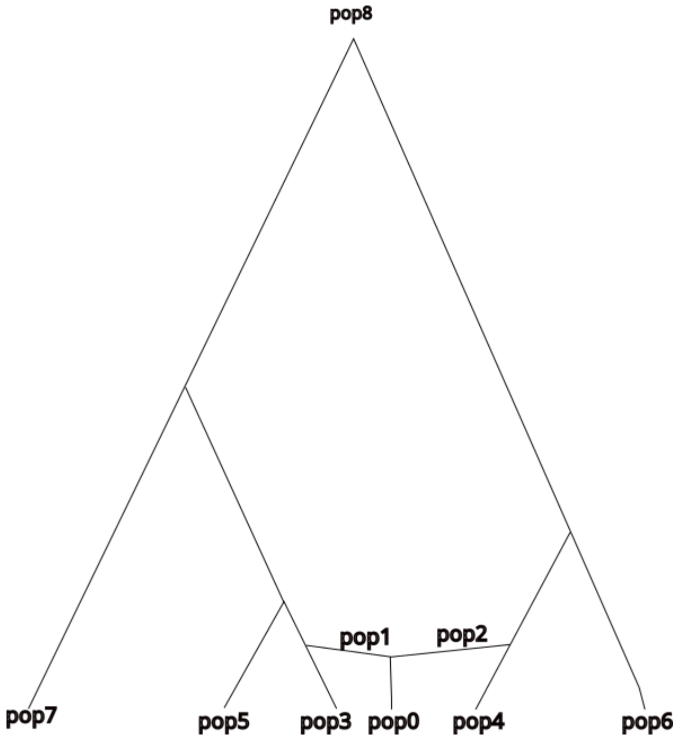


**Supplementary Figure S36:** Population pop0 results as an admixture of pop1 and pop2. We have no sampled individuals from pop1 and pop2. Ten individuals have been sampled from each of the pop3 to pop7. Thus, pop3 and pop4 are considered the right sources of pop0 since no samples have been obtained from pop1 and pop2. To test how f3 behaves, we applied an instantaneous bottleneck just before sampling pop0 (bottleneck strength: 0, 10, 50, 100, 150 - 0 indicates no bottleneck). In addition, we applied two migration scenarios. In the first, only pop3 and pop4 exchange migrants, whereas in the second all populations pop3-pop7 experience various degrees of migration rates.

We then applied an instantaneous bottleneck to pop0 of various strengths ($b = 0, 10, 50, 100, 150$). $b$ resembles the coalescent time that corresponds to the bottleneck, since a bottleneck accelerates the coalescent rate. Thus, 0 means no bottleneck and 150 is the most severe bottleneck. In addition, we allowed for gene flow between pop3 and pop4 (scenario 1) or between all sampled populations (scenario 2). The migration rates we tried are the following: *m=0, 0.001, 0.05, 0.1. m* indicates the proportion of the destination populations (forward in time) composed of migrants. Results are presented in **Supplementary Figure S37**. We observe that even in this very simple scenario (without all complications of missing data, small sample sizes etc), there are combinations that give no negative values of the (Admixture) f3 statistic.

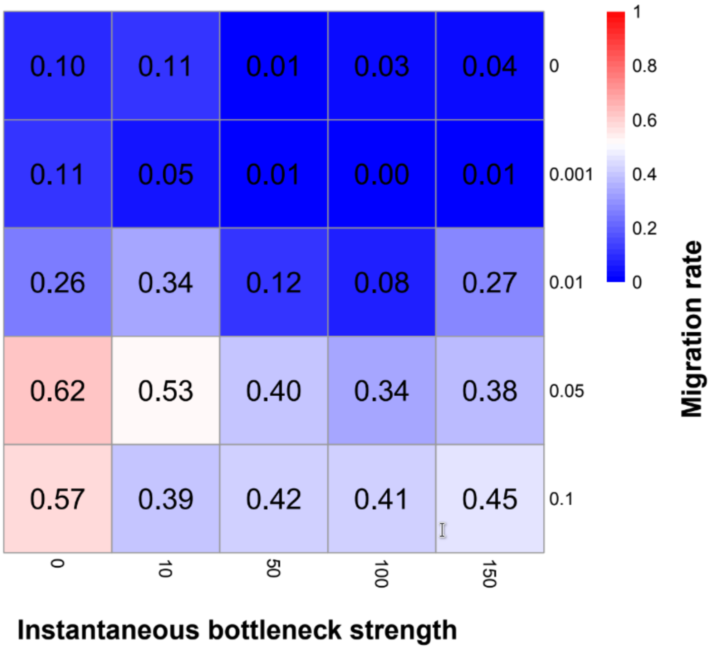


**Supplementary Figure S37**: The proportion of negative values of the f3 statistic (average value for 20 replications) for all plausible triplets of the 8 populations depicted in **Supplementary Figure S36**. For bottleneck strength 100 and migration rate 0.001, none triplet exhibited a negative value for the Admixture f3 statistic. Thus, even if there is admixture and also gene flow, Admixture f3 did not support the scenario of admixture.

**F-statistics analyses**

For the F-statistics analyses we grouped the ancient individuals (public and newly sequenced here; “**Dataset 1**”) according to their general genetic ancestry and their chronological and archaeological contexts, and performed the analysis on a per-group and per-individual basis. Here, we applied the same grouping appg8roach as for the ADMIXTURE analyses, albeit different analyses required generating different subdatasets (**Supplementary Table 4; Additional file 4**; see also details below)**.** Samples *Amv_Epi_Cl_5*, *Amv_Epi_Cl_4*, *Amv_Epi_Hel_3*, and *Ten_Pel_Arch_1* were removed from per-group analyses as they have a 1st-/2nd-degree genetic kinship relationship with individuals that exhibit higher coverage in the dataset. They were included, nonetheless, in the per-individual analyses.

**Ancestry modeling was performed with qpAdm using the ‘rotating population’ strategy** [139]. This approach begins by identifying a set of "candidate" populations from which we iteratively select a defined number of "sources" of ancestry for our "target" population (1 to 4 populations in our analyses). Once the sources are selected, they are removed from the candidate list, and all remaining populations are classified as the "right or outgroup" populations. Next, we iteratively fit qpAdm models for each combination of sources, and right populations, keeping the target fixed. We extract p-values and z-scores. A model is called "feasible" and is further examined if p-value > 0.05 and all z-scores are positives. The rotating population analysis was performed using the R interface of ADMIXTOOLS2 v.2.0.0 [140]. We performed three sets of rotating qpAdm runs by always keeping the same targets, but changing the potential sources. As separate targets we used the focal populations of Ammotopos, Amvrakia, Tenea during different time periods (LBA, Archaic, Classical, Hellenistic, Roman, etc). For the first run we used distant genetic sources (henceforth denoted as "Ultimate" sources) that characterize the general ancestry of ancient Western Europeans following Lazaridis et al. [141]. The same sources were used for all targets in order to compare their general genetic ancestry (see **Supplementary Table 4; Additional file 4** for details). For the second run, we performed the analysis under the same setting, but replaced the "Ultimate" sources with spatiotemporally more proximate ones (henceforth denoted as "More_proximate" sources; see **Supplementary Table 4; Additional file 4** for details) in order to again compare the ancestry modeling of the different targets using alternative sources that may better represent their common ancestry. For the third run, the goal was to identify the immediate sources of each target. Therefore, in this run, each target had a different set of potential sources that was, spatiotemporally, as close to the target as possible: the temporal range was limited to those individuals that dated earlier than the target, whereas the spatial range was restrained to the Eastern Mediterranean and adjacent areas, including Italy, the Balkans, and the Middle East ("Most_proximate" sources, hereinafter; see **Supplementary Table 4**; **Additional file 4** for details). Additionaly, we performed a targeted qpAdm analysis using Archaic Amvrakia as target population and LBA Ammotopos and Archaic Tenea as the two potential source populations in order to exclusively test if Archaic Amvrakia can be inferred either as admixed by the other two, or a direct descendant of one of the other two, or none of the above. In this run as reference (right) populations we used, predominantly, representatives from IA Eastern Mediterranean (see **Supplementary Table 4**; **Additional file 4)**. For comparison reasons we perform the analyses using various parameters including a) *adjust_pseudohaploid*: *TRUE/FALSE*, b) *maxmiss* values of 0.00, 0.10, and 0.40, and c) *afprod: TRUE/FALSE*. The two latter parameters regulate the influence of missing data in the qpAdm inferences. The script is provided at https://doi.org/10.5281/zenodo.10848927.

Feasible models (accepted models) with a p-value > 0.05 and positive z-scores (or equivalently all weights being positive) were visualized using upset-stacked barplots. The points in this scatterplot represent the populations involved, and the color gradient indicates their admixture proportions. The models were sorted based on the number of involved sources (1-4). All the per-group and per-individual qpAdm scatterplots are provided in https://doi.org/10.5281/zenodo.10848927, along with all the rotating qpAdm analysis outputs (tables; "model1" in the file names correspond to populations-based results, whereas model2 to individual-based results).

For all the "Ultimate"-based runs, many feasible models were produced when using *maxmiss: 0.00*, with an Anatolian Neolithic source being the most frequent major contributor. When applying a *maxmiss: 0.40*, no feasible models were produced in most cases. An intermediate value of *maxmiss: 0.01* resulted in several inferences, again with an Anatolian Neolithic source being the most frequent major contributor. The parameter *afprod: TRUE* produces similar results, in most cases, albeit it seems to should necessarily be enabled when using *maxmiss: 0.40*, as when disabled most results are non feasible. Similar results are observed for the "More_proximate"-based runs, with a prehistoric Southern Balkan source (especially Greece) being the most frequently inferred major contributor. The same pattern is observed for the "Most_proximate"-based runs, too. Given the above observations, as the results are too many to present all of them, we present the ones for the parameterization scheme "*adjust_pseudohaploid*: *TRUE - maxmmiss: 0.10 - afprod: TRUE*". The results for our newly sequenced genomes using the "Most_proximate" to each target set of sources (**Supplementary Figures S40**), as well as the targeted to Archaic Amvrakia run (**Figure 4**), are presented in the main text. For the "Ultimate" and the "More_proximate" set of sources the results (**Supplementary Figures S38-S39**) are presented in the following:

- Ammotopos inferred to be in rotating qpAdm using:
    - a) the "Ultimate" set of sources: a four-way admixture among the Anatolian Neolithic Barcin (6500-5900BCE) as a major source (>60%) and other minor sources, including Neolithic Iran, Chalcolithic Iran, and Russian EBA Yamnaya-like ancestry (**Supplementary Figure S38A)**,
    - b) the "More_proximate" set of sources: a two-way admixture between a southern Balkan MBA source [MBA mainland Greece (2100-1600 BCE) or MBA Albania (1900-1700 BCE)] and either Neolithic Greece (6400-3600 BCE) or EBA Greece (2900-2000 BCE). When excluding Neolithic Greece (6400-3600 BCE) that is the oldest of them, then the remaining models include EBA Greece (2900-2000 BCE) as the source with the greatest contribution (~0.6%) and one of the two MBA sources as the second greatest (~0.4%); in three-way and four-way admixture models, sources with lower contribution include BA Italy and other BA Balkan areas (**Supplementary Figure S39A).**
- Archaic Amvrakia inferred to be in rotating qpAdm using:
    - a) the "Ultimate" set of sources: either a three-way or a four-way admixture among the Anatolian Neolithic Barcin (6500-5900BCE) as a major source (>60%) and other minor sources, including Neolithic Iran, Chalcolithic Iran, and Russian EBA Yamnaya-like ancestry (**Supplementary Figure S38B**).
    - b) the "More_proximate" set of sources: a three-way admixture among two major sources (of ~45%, each), Neolithic Greece (6400-3600 BCE) and MBA Mainland Greece (2100-1600 BCE), and LBA Crete (1700-1250 BCE) as a third minor source (**Supplementary Figure S39B**).
- Classical Amvrakia inferred to be in rotating qpAdm using:
    - a) the "Ultimate" set of sources: either a three-way or a four-way admixture among the Anatolian Neolithic Barcin (6500-5900BCE) as a major source (>60%) and other minor sources, including Neolithic Iran, Chalcolithic Iran, and Russian EBA Yamnaya-like ancestry (**Supplementary Figure S38C**),
    - b) the "More_proximate" set of sources: a two-way admixture between EBA Greece (2900-2000 BCE) as the main source (~0.6%) and MBA mainland Greece (2100-1600 BCE) as the second source (~0.4%): in the three-way admixture model, sources with lower contribution include LBA Crete (1700-1250 BCE), as well as BA Türkiye and LBA Peloponnese although with quite low contribution in both cases (**Supplementary Figure S39C**).
- Hellenistic Amvrakia inferred to be in rotating qpAdm using:
    - a) the "Ultimate" set of sources: a four-way admixture among an Anatolian Neolithic major source (>25-60%), either Barcin (6500-5900BCE) or Boncuklu Hoyuk (8300-7600 BCE), and other minor sources, including Neolithic Iran, Chalcolithic Iran, and Russian EBA Yamnaya-like ancestry (**Supplementary Figure S38D**),
    - b) the "More_proximate" set of sources: a two-way admixture between EBA Greece (2900-2000 BCE) as the main source (~0.6%) and MBA Albania (2100-

    1600 BCE) as the second source (~0.4%); in three-way and four-way admixture models, several other sources with lower contribution appear, such as MBA and LBA Greece, BA Italy, BA Balkans, etc. (**Supplementary Figure S39D**).

- Archaic Tenea inferred to be in rotating qpAdm using:
    - a) the “Ultimate” set of sources: either a two-, three-, or four-way admixture among an Anatolian Neolithic major source (>25-60%), either Barcin (6500-5900BCE) or Boncuklu Hoyuk (8300-7600 BCE), and other minor sources, including Neolithic Iran, Chalcolithic Iran, and Russian EBA Yamnaya-like ancestry (**Supplementary Figure S38E**),
    - b) the “More_proximate” set of sources: a two-way admixture between EBA Greece (2900-2000 BCE) as main source (~0.6%) and a southern Balkan MBA source [e.g. MBA mainland Greece (2100-1600 BCE) or MBA Albania (1900-1700 BCE)] as a second source (~0.4%), with Neolithic Greece (6400-3600 BCE) and BA Crete (2900-1700) being interchangeable with EBA Greece; in three-way and four-way admixture models, sources with lower contribution include BA Italy and other BA Balkan populations (**Supplementary Figure S39E**).
- Hellenistic Tenea is inferred to be in rotating qpAdm using:
    - a) the “Ultimate” set of sources: a four-way admixture among the Anatolian Neolithic Barcin (6500-5900BCE) as a major source (>60%) and other minor sources, including Neolithic Iran, Chalcolithic Iran, and Russian EBA Yamnaya-like ancestry (**Supplementary Figure S38F**),
    - b) the “More_proximate” set of sources: a two-way admixture between either EBA Greece (2900-2000 BCE) as main source (~0.7%) and Yamnaya-like BA Balkans (2900-2000 BCE) as the second source (~0.3%) or MBA Albania (2100-1600 BCE) (~0.6%) and EBA Greece (2900-2000 BCE) (0.4%) or MBA mainland Greece (2100-1600 BCE) ~(~0.6%) and LBA mainland Greece (Sterea Ellada; 1600-1300 BCE) (0.4%); in three-way and four-way admixture models, several other sources with lower contribution appear, such as LBA Crete, LBA Peloponnese, BA Levant, BA Türkiye etc (**Supplementary Figure S39F**).
- Roman Tenea is inferred to be in rotating qpAdm using:
    - a) the “Ultimate” set of sources: either a three-way or a four-way admixture among two major sources (~40-45% each), Anatolian Neolithic Barcin (6500-5900BCE) and CHG (11500-7500 BCE), and other minor sources, including Natufian Levant (12000-9500 BCE), Neolithic Iran, and PPN Levant (**Supplementary Figure S38G**),
    - b) the “More_proximate” set of sources: a three-way admixture among MBA_MLBA South-Southwest Türkiye (2000-1300 BCE) as the source with the greatest contribution (~0.4%), EBA Greece (2900-2000 BCE) as the second greatest (~0.35%), and MBA mainland Greece (2100-1600 BCE) as the third greatest (~0.25%); alternatively, in four-way admixture model it can be modeled as admixture among the three aforementioned sources and LBA Mainland Greece (Sterea Ellada; 1600-1300 BCE) (**Supplementary Figure S39G**).

**A.**

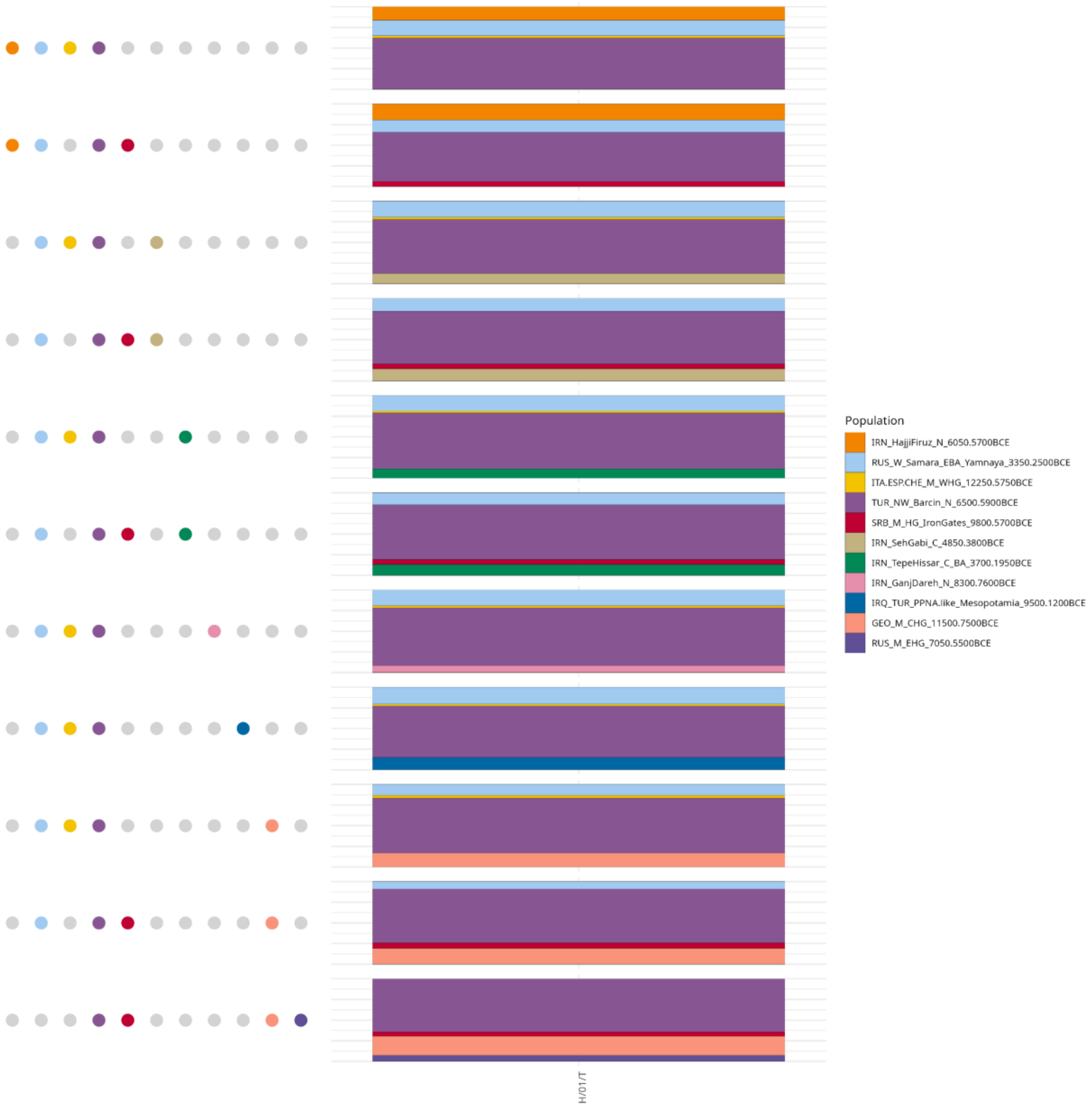
Population
IRN_HajjiFiruz_N_6050.5700BCE
RUS_W_Samara_EBA_Yamnaya_3350.2500BCE
ITA.ESP.CHE_M_WHG_12250.5750BCE
TUR_NW_Barcin_N_6500.5900BCE
SRB_M_HG_IronGates_9800.5700BCE
IRN_SehGabi_C_4850.3800BCE
IRN_TepeHissar_C_BA_3700.1950BCE
IRN_GanjDareh_N_8300.7600BCE
IRQ_TUR_PPNA.like_Mesopotamia_9500.1200BCE
GEO_M_CHG_11500.7500BCE
RUS_M_EHG_7050.5500BCE
T/10/H

**B.**

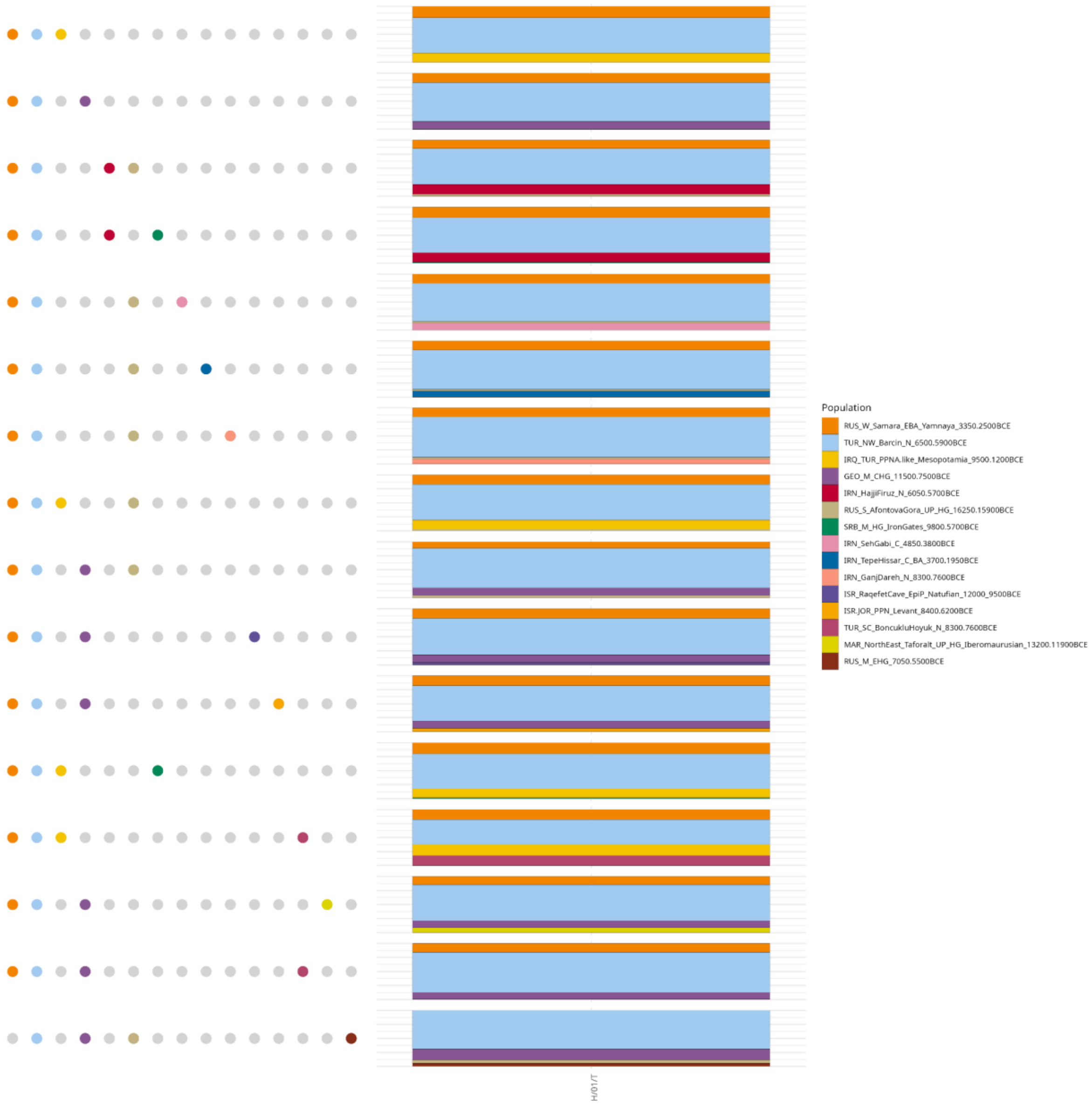
Population
RUS_W_Samara_EBA_Yamnaya_3350.2500BCE
TUR_NW_Barcin_N_6500.5900BCE
IRQ_TUR_PPNA.like_Mesopotamia_9500.1200BCE
GEO_M_CHG_11500.7500BCE
IRN_HajjiFiruz_N_6050.5700BCE
RUS_S_AfontovaGora_UP_HG_16250.15900BCE
SRB_M_HG_IronGates_9800.5700BCE
IRN_SehGabi_C_4850.3800BCE
IRN_TepeHissar_C_BA_3700.1950BCE
IRN_GanjDareh_N_8300.7600BCE
ISR_RaqefetCave_EpiP_Natufian_12000_9500BCE
ISR.JOR_PPN_Levant_8400.6200BCE
TUR_SC_BoncukluHoyuk_N_8300.7600BCE
MAR_NorthEast_Taforalt_UP_HG_Iberomaurusian_13200.11900BCE
RUS_M_EHG_7050.5500BCE

**C.**

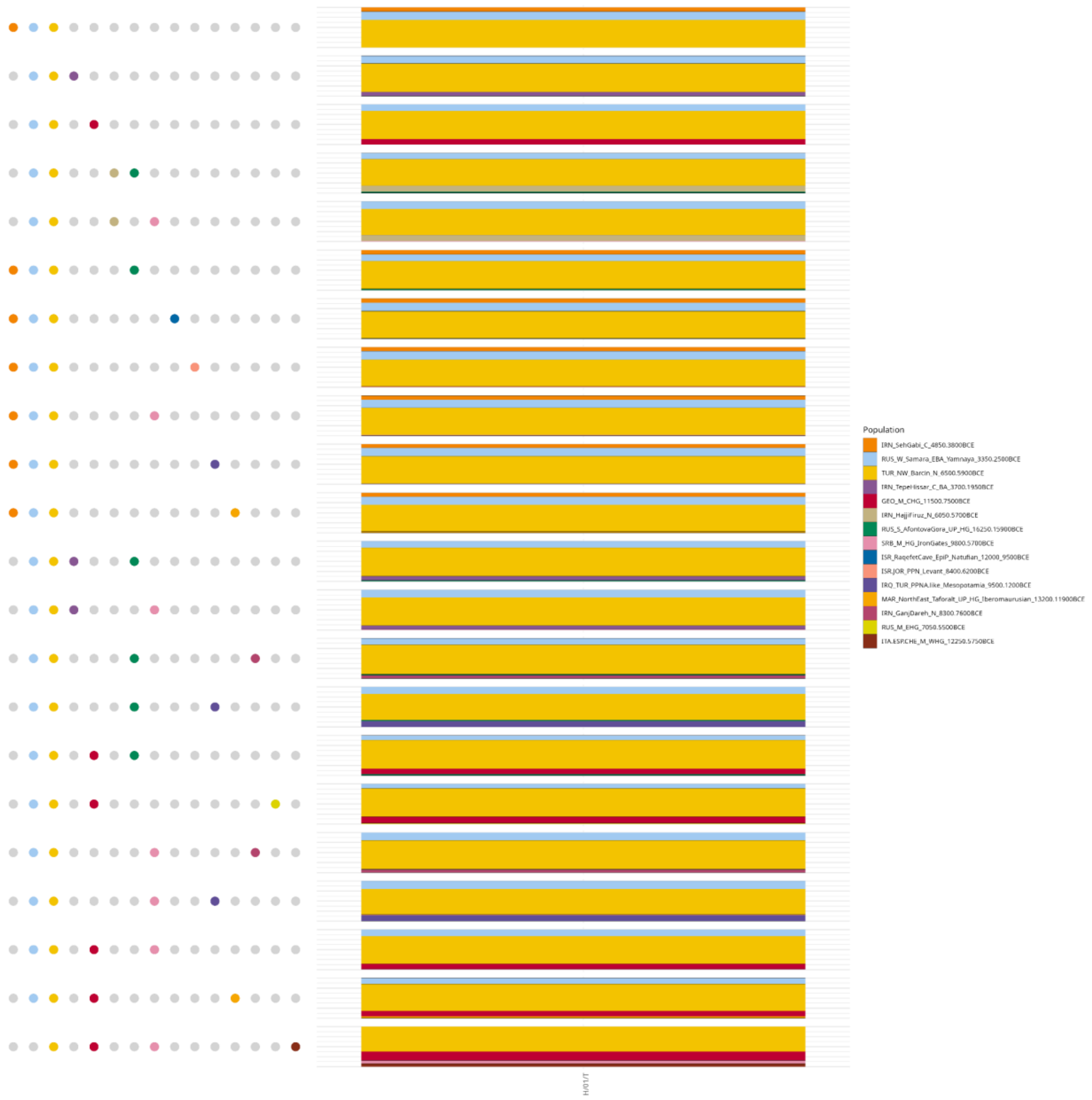

Population
IRN_SehGabi_C_4850.3800BCE
RUS_W_Samara_EBA_Yamnaya_3350.2500BCE
TUR_NW_Barcin_N_6500.5900BCE
IRN_TepeHissar_C_BA_3700.1950BCE
GEO_M_CHG_11500.7500BCE
IRN_HajjiFiruz_N_6050.5700BCE
RUS_S_AfontovaGora_UP_HG_16250.15900BCE
SRB_M_HG_IronGates_9800.5700BCE
ISR_RaqefetCave_EpiP_Natufian_12000_9500BCE
ISR.JOR_PPN_Levant_8400.6200BCE
IRQ_TUR_PPNA.like_Mesopotamia_9500.1200BCE
MAR_NorthEast_Taforalt_UP_HG_Iberomaurusian_13200.11900BCE
IRN_GanjDareh_N_8300.7600BCE
RUS_M_EHG_7050.5500BCE
ITA.ESP.CHE_M_WHG_12250.5750BCE

**D.**

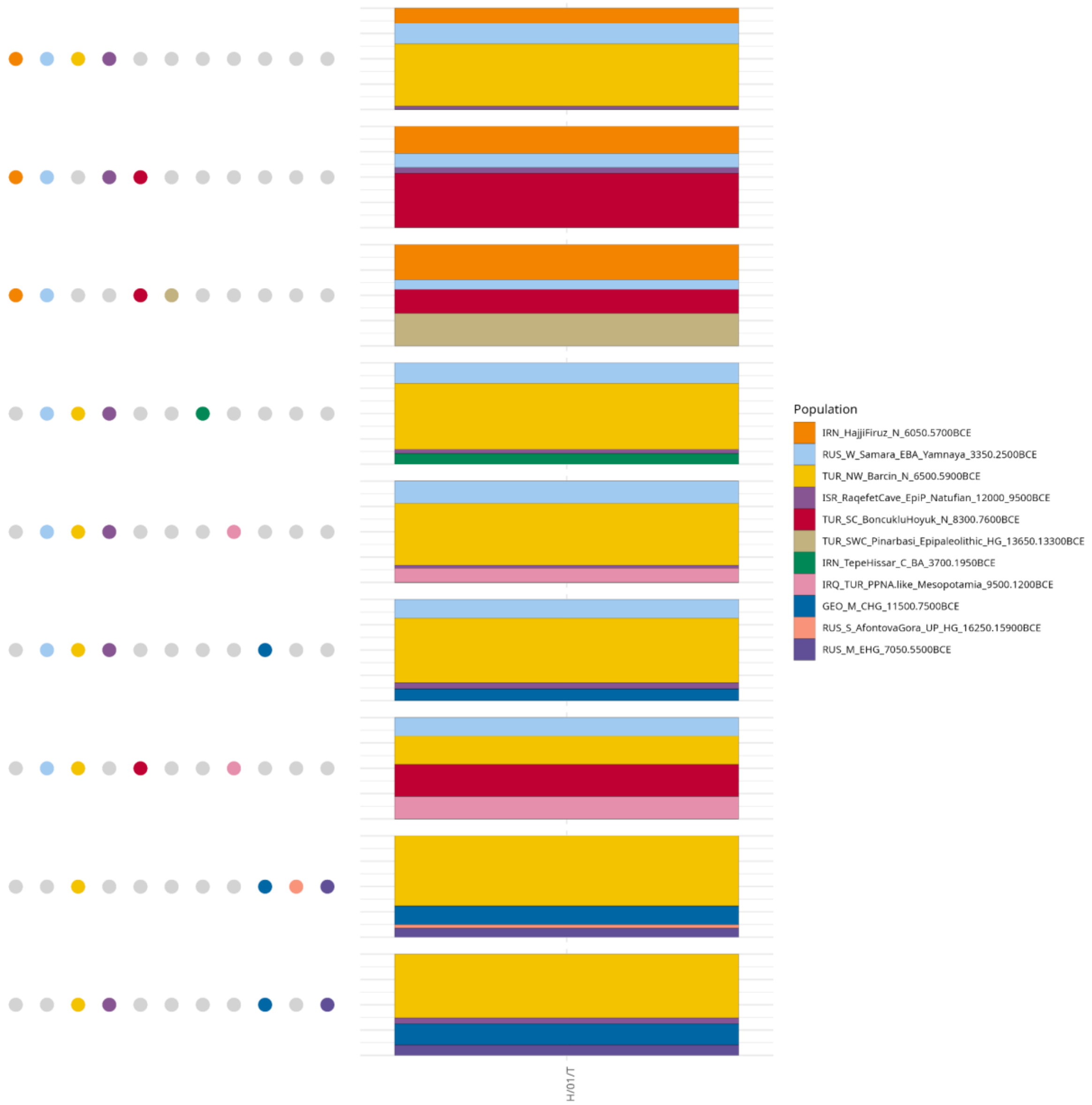
H/01/T
Population
IRN_HajjiFiruz_N_6050.5700BCE
RUS_W_Samara_EBA_Yamnaya_3350.2500BCE
TUR_NW_Barcin_N_6500.5900BCE
ISR_RaqefetCave_EpiP_Natufian_12000_9500BCE
TUR_SC_BoncukluHoyuk_N_8300.7600BCE
TUR_SWC_Pinarbasi_Epipaleolithic_HG_13650.13300BCE
IRN_TepeHissar_C_BA_3700.1950BCE
IRQ_TUR_PPNA.like_Mesopotamia_9500.1200BCE
GEO_M_CHG_11500.7500BCE
RUS_S_AfontovaGora_UP_HG_16250.15900BCE
RUS_M_EHG_7050.5500BCE

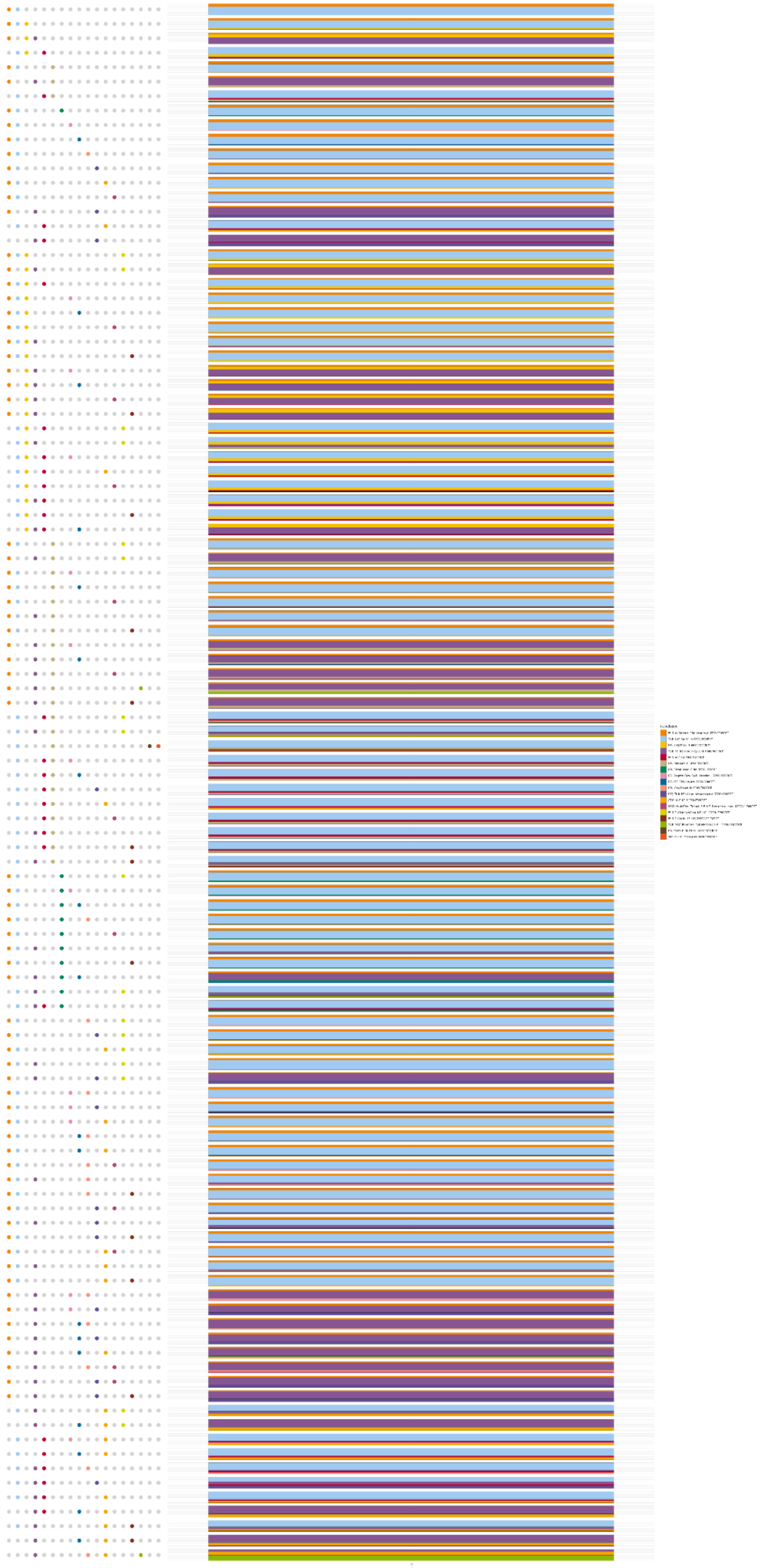

1732 **E.**

1733 **F.**

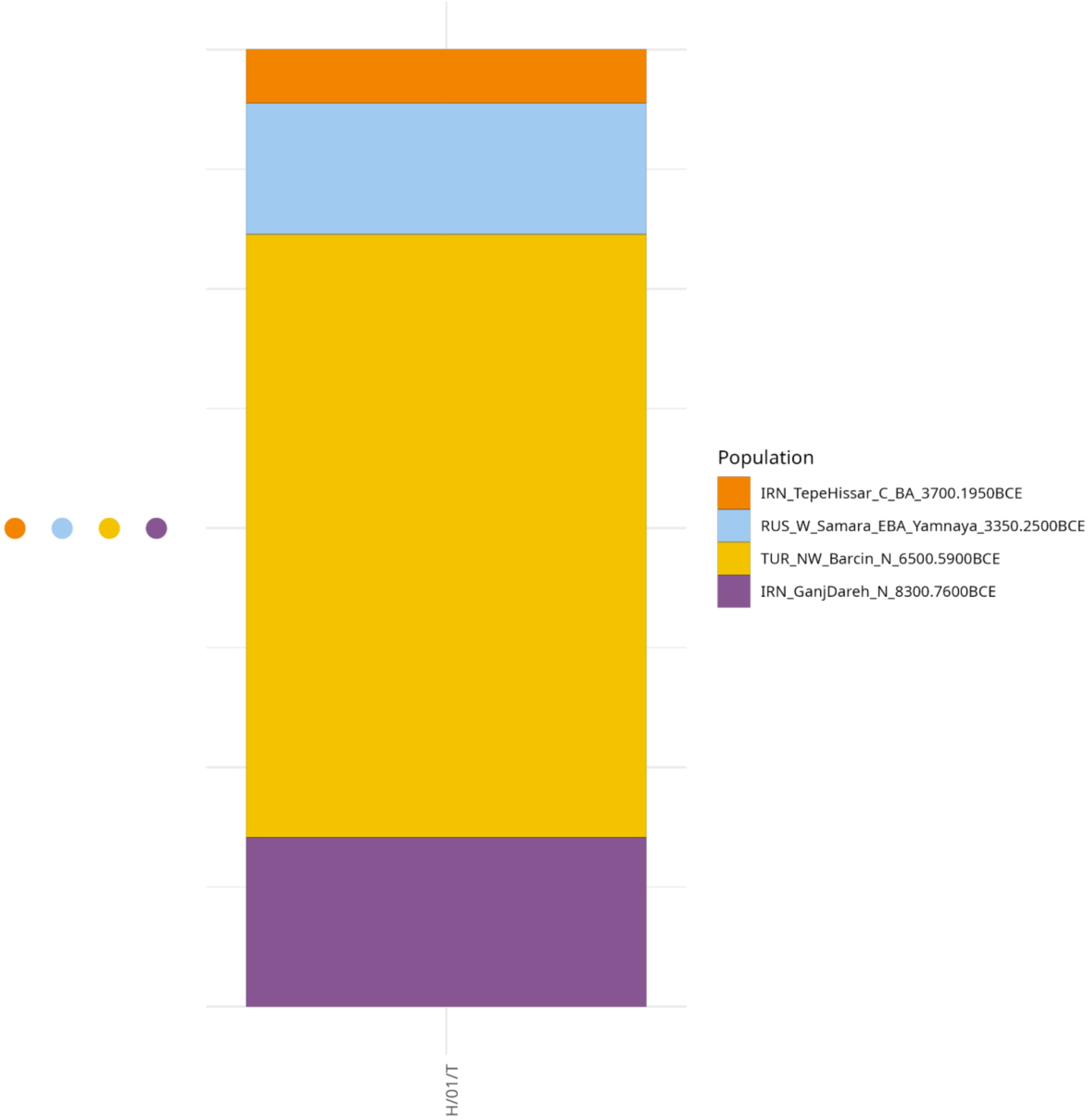
Population
IRN_TepeHissar_C_BA_3700.1950BCE
RUS_W_Samara_EBA_Yamnaya_3350.2500BCE
TUR_NW_Barcin_N_6500.5900BCE
IRN_GanjDareh_N_8300.7600BCE
H/01/T

1734

**G.**

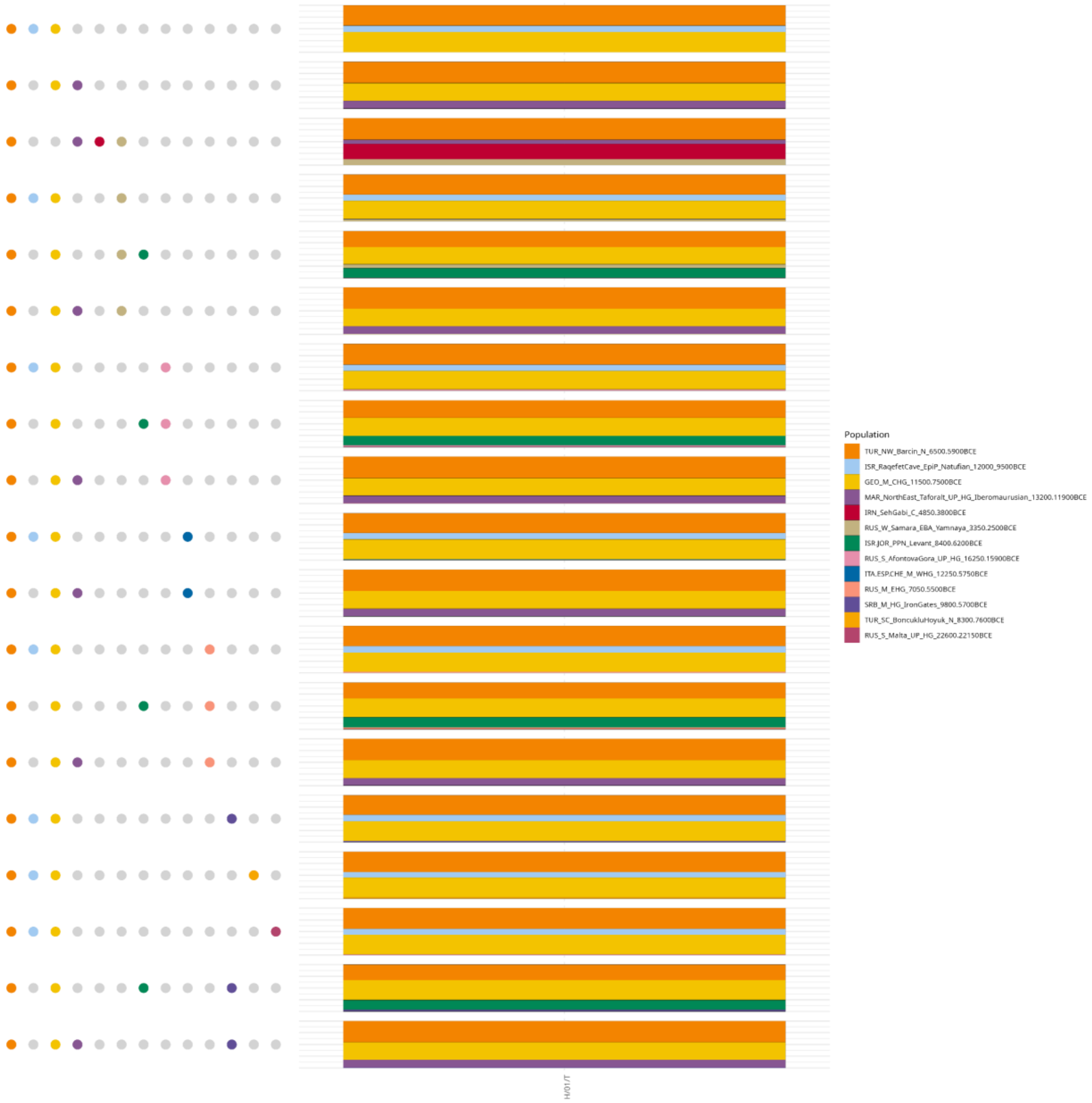


**Supplementary Figure S38.** Scatterplot of qpAdm analyses results at the population level using the "Ultimate" sample-set as putative sources, using the parameterization scheme "*adjust_pseudohaploid: TRUE - maxmmiss: 0.10 - afprod: TRUE*". Only the feasible models with a p-value > 0.05 (accepted models) are shown with the points representing the populations involved and the colors indicating their admixture proportions. The models were sorted based on the number of involved sources (1-4). The tested target population is **A.** LBA Ammotopos **B.** Archaic Amvrakia, **C.** Classical Amvrakia **D.** Hellenistic Amvrakia **E.** Archaic Tenea **F.** Hellenistic Tenea **G.** Roman Tenea.

**A.**

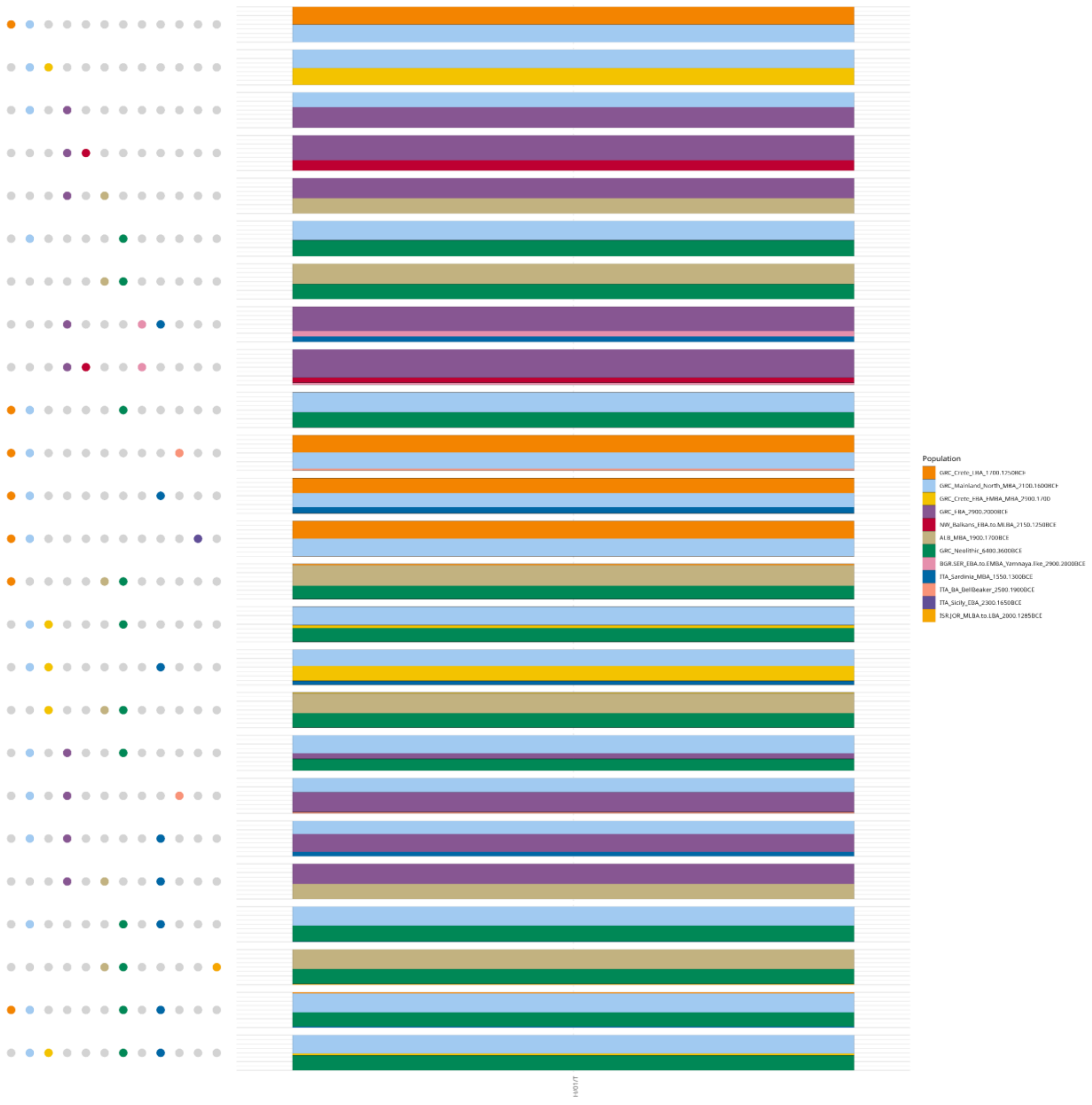

Population
GRC_Crete_LBA_1700.1250BCE
GRC_Mainland_North_MBA_2100.1600BCE
GRC_Crete_EBA_EMBA_MBA_2900.1700
GRC_EBA_2900.2000BCE
NW_Balkans_EBA.to.MLBA_2150.1250BCE
ALB_MBA_1900.1700BCE
GRC_Neolithic_6400.3600BCE
BGR.SER_EBA.to.EMBA_Yamnaya.like_2900.2000BCE
ITA_Sardinia_MBA_1550.1300BCE
ITA_BA_BellBeaker_2500.1900BCE
ITA_Sicily_EBA_2300.1650BCE
ISR.JOR_MLBA.to.LBA_2000.1285BCE

**B.**

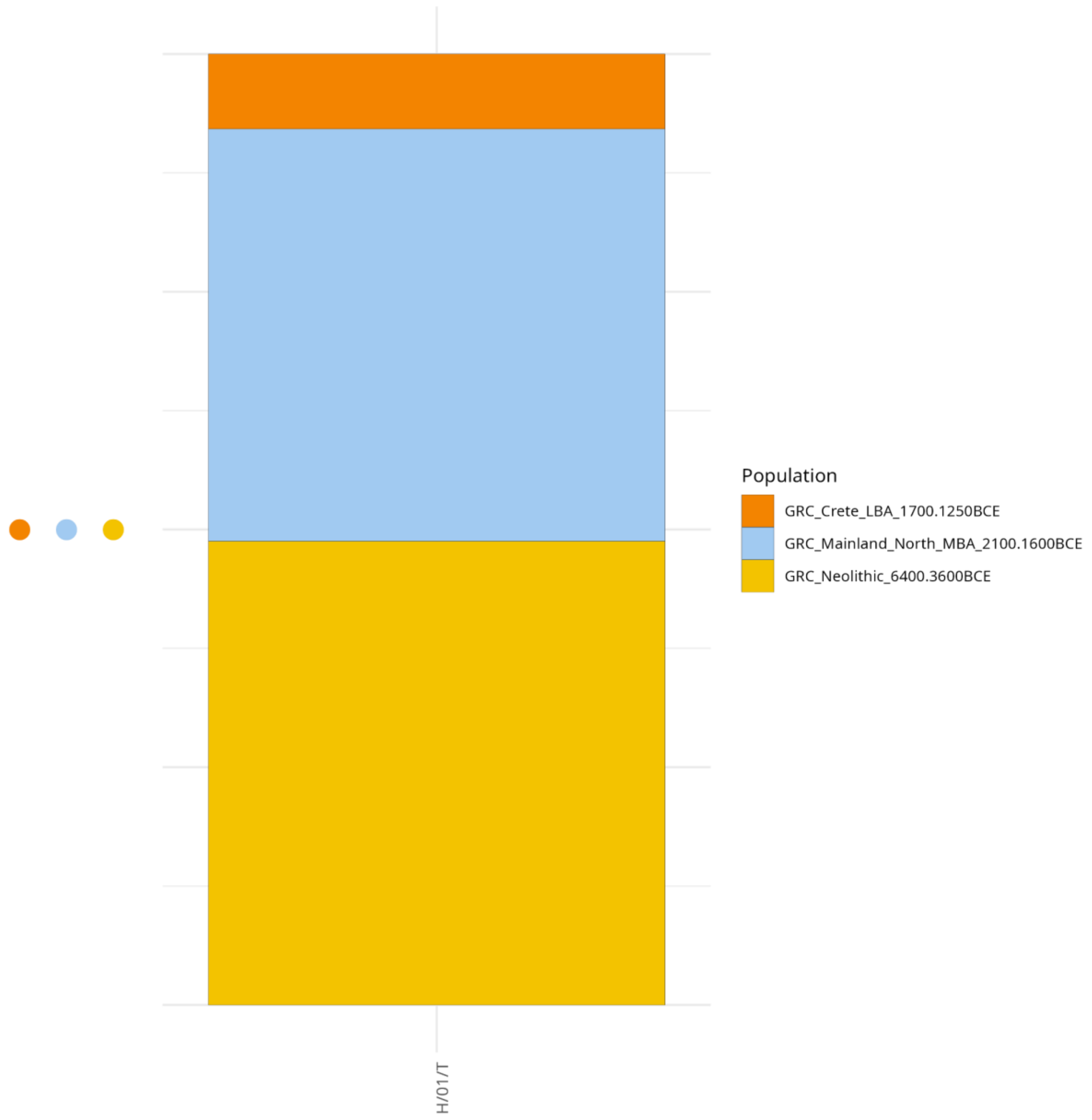
Population
GRC_Crete_LBA_1700.1250BCE
GRC_Mainland_North_MBA_2100.1600BCE
GRC_Neolithic_6400.3600BCE
H/01/T

**C.**

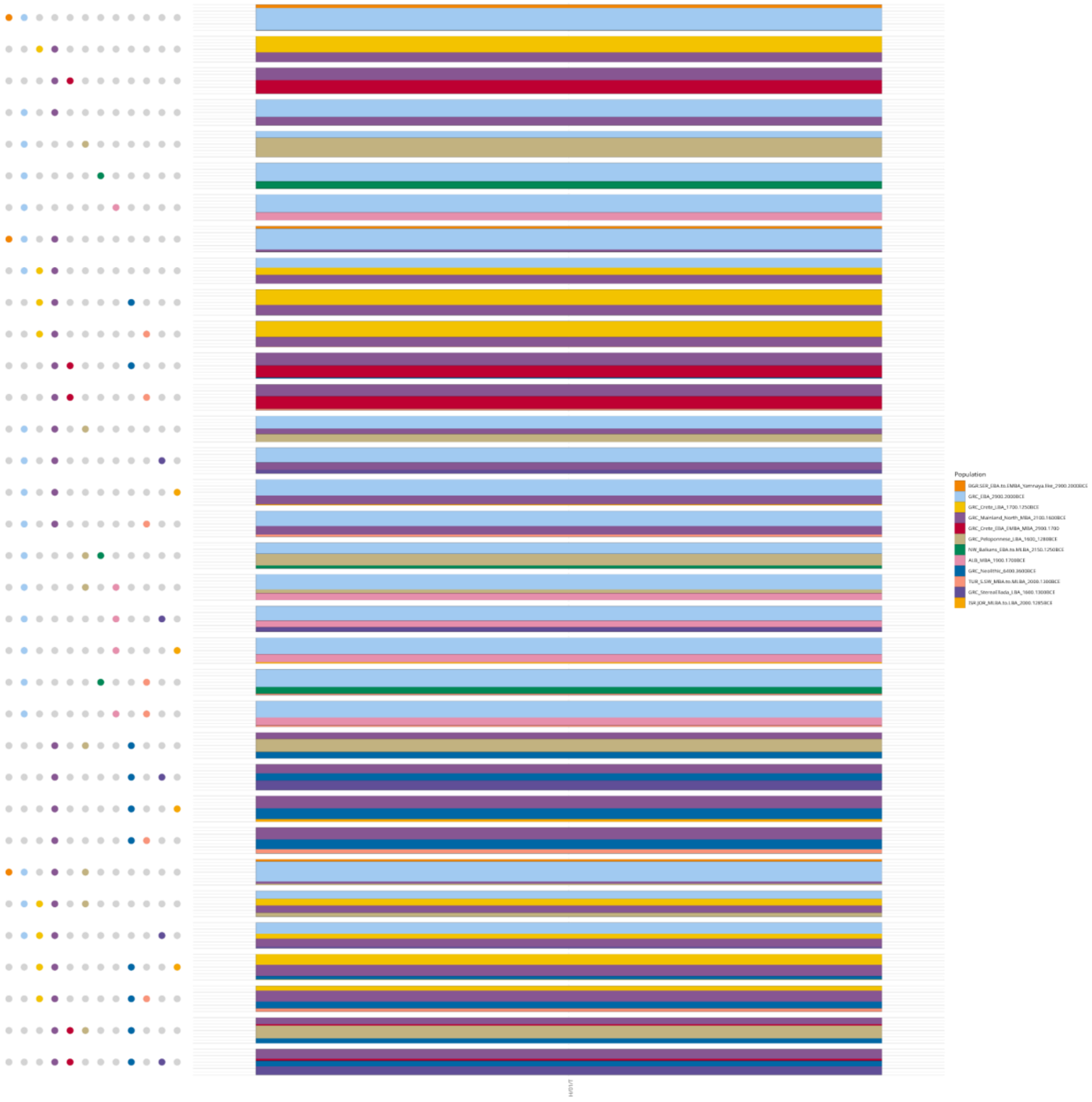
Population
GRC_EBA_2900.2000BCE
GRC_Crete_LBA_1700.1250BCE
GRC_Mainland_North_MBA_2100.1600BCE
ALB_MBA_1900.1700BCE
GRC_Neolithic_6400.3600BCE

**D.**

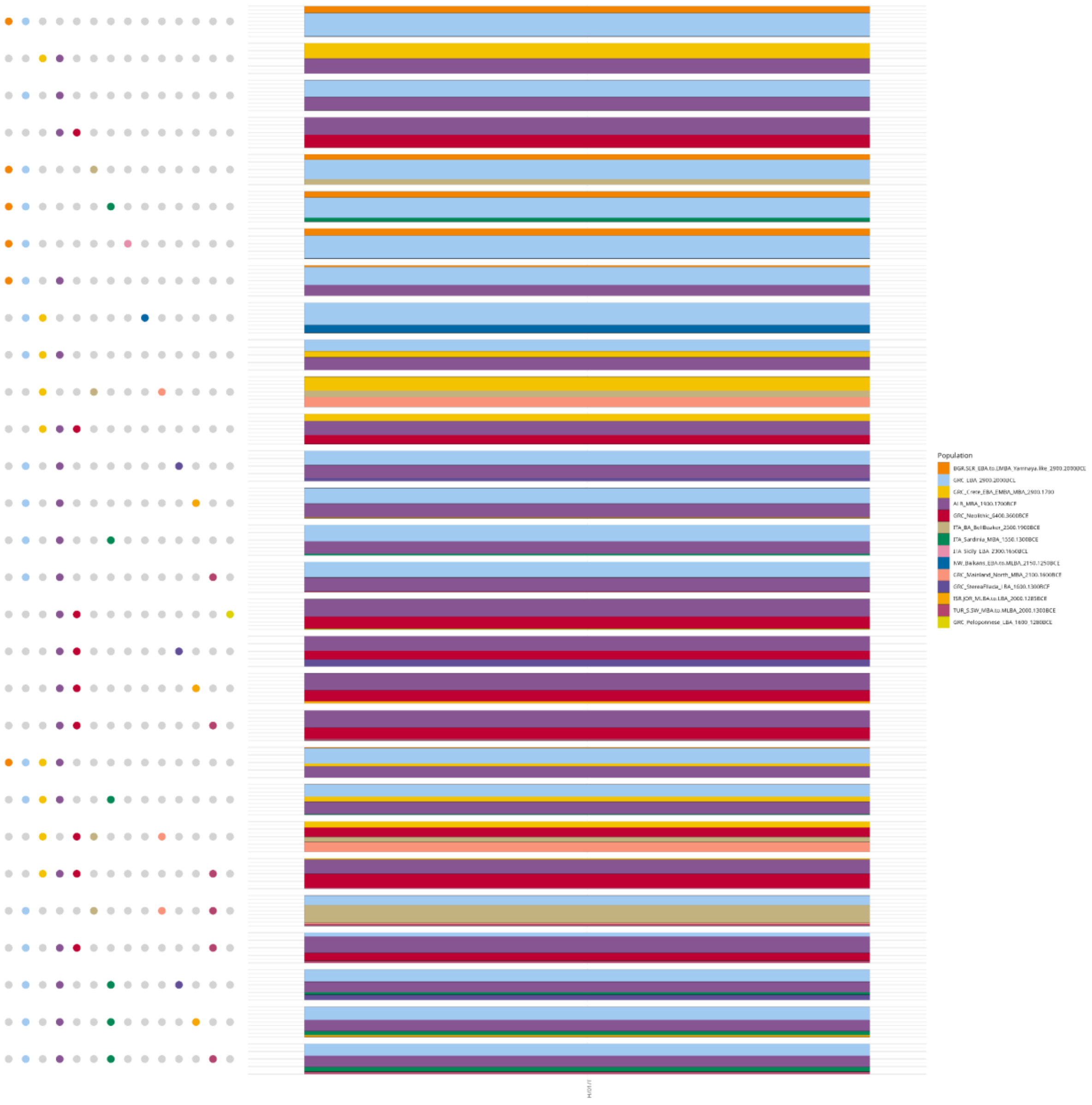

Population
GRC_Crete_EBA_EMBA_MBA_2900.1700
GRC_Neolithic_6400.3600BCE
ITA_Sardinia_MBA_1550.1300BCE
NW_Balkans_EBA.to.MLBA_2150.1250BCE
GRC_Mainland_North_MBA_2100.1600BCE
TUR_S.SW_MBA.to.MLBA_2000.1300BCE

**E.**

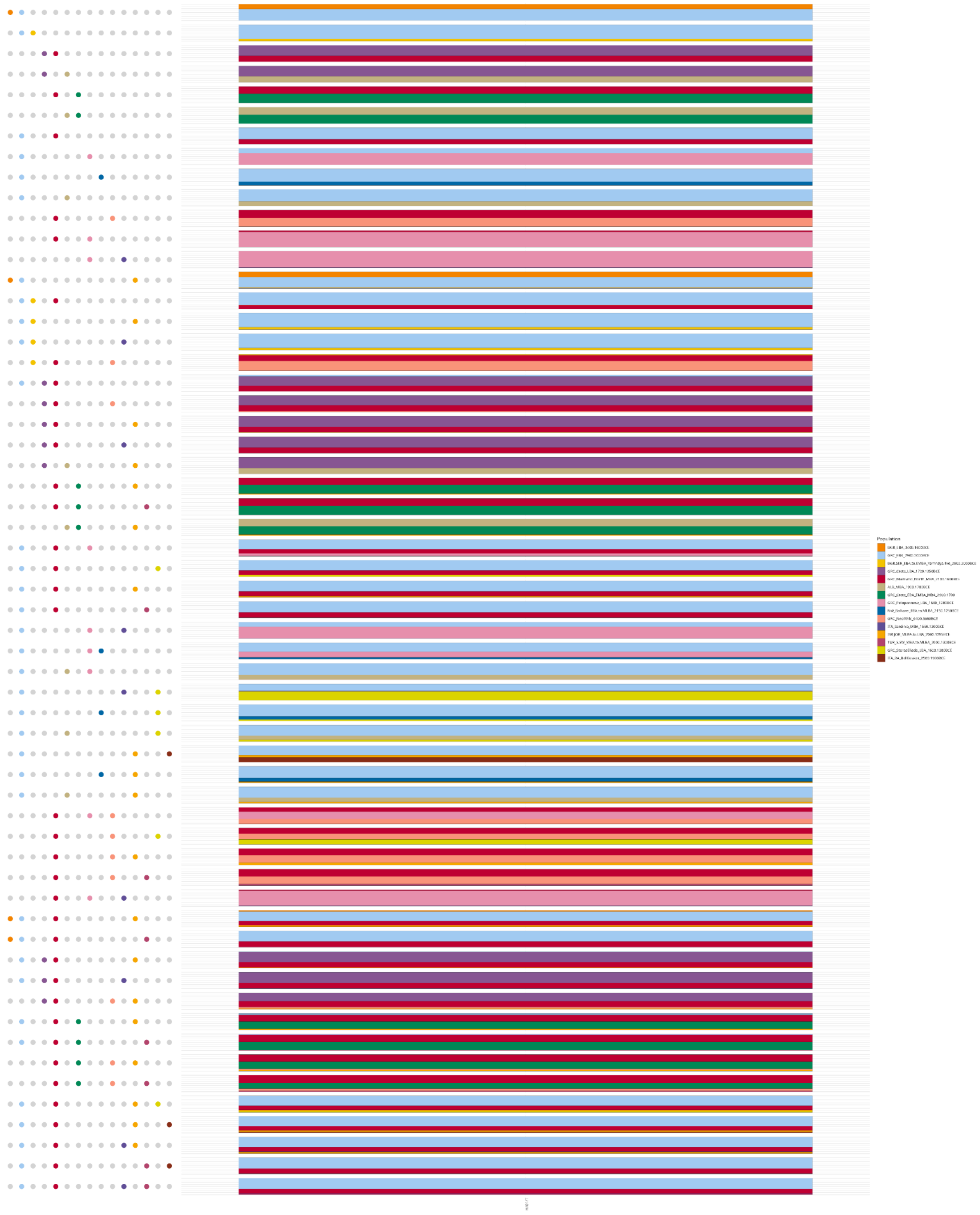
Population
BGR_EBA_3400.1600BCE
GRC_EBA_2900.2000BCE
BGR.SER_EBA.to.EMBA_Yamnaya.like_2900.2000BCE
GRC_Crete_LBA_1700.1250BCE
GRC_Mainland_North_MBA_2100.1600BCE
ALB_MBA_1900.1700BCE
GRC_Crete_EBA_EMBA_MBA_2900.1700
GRC_Peloponnese_LBA_1600_1200BCE
NW_Balkans_EBA.to.MLBA_2150.1250BCE
GRC_Neolithic_6400.3600BCE
ITA_Sardinia_MBA_1550.1300BCE
TUR_S.SW_MBA.to.MLBA_2000.1300BCE
GRC_StereaEllada_LBA_1600.1300BCE
ITA_BA_BellBeaker_2500.1000BCE

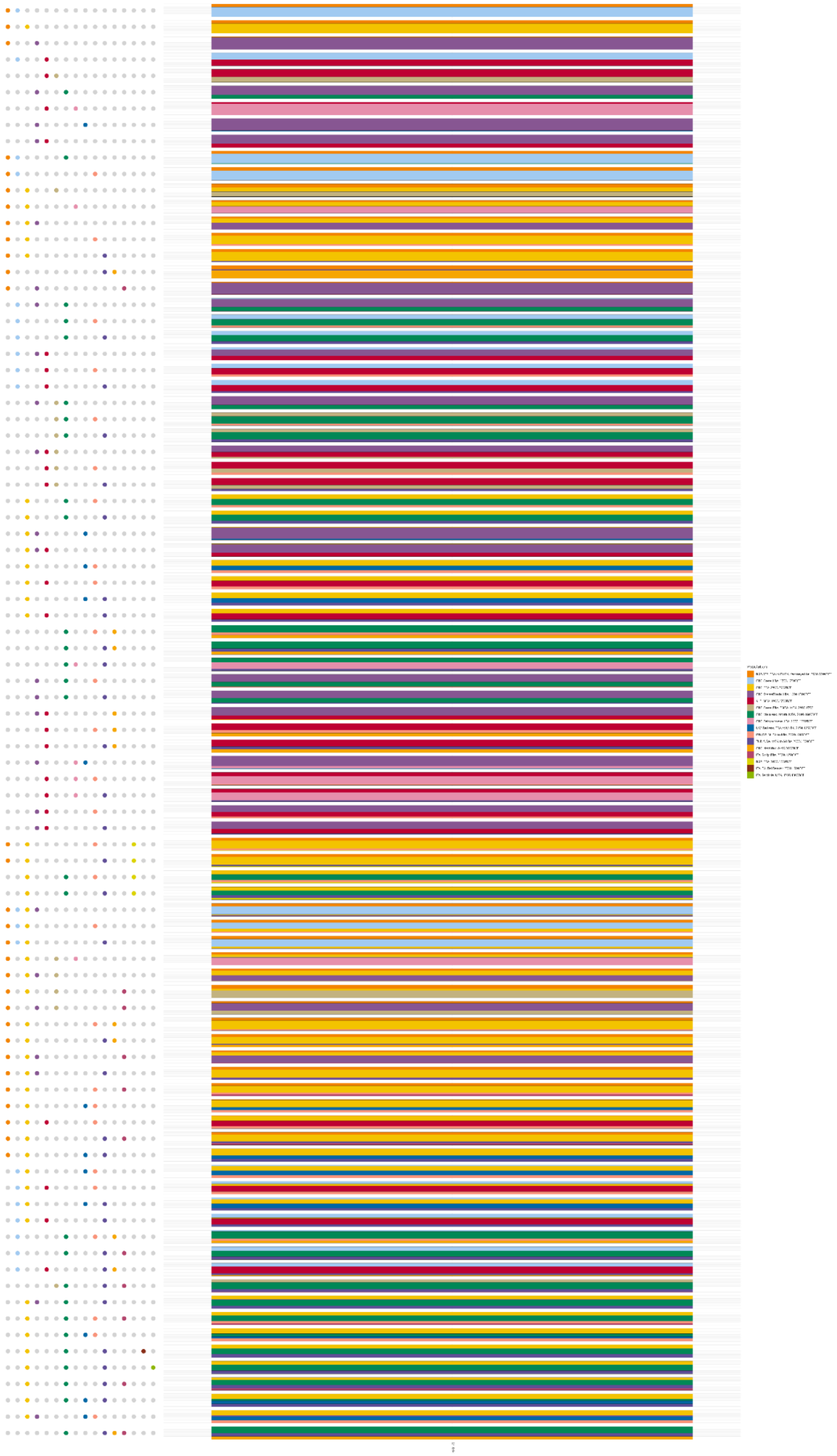
Population

**G.**

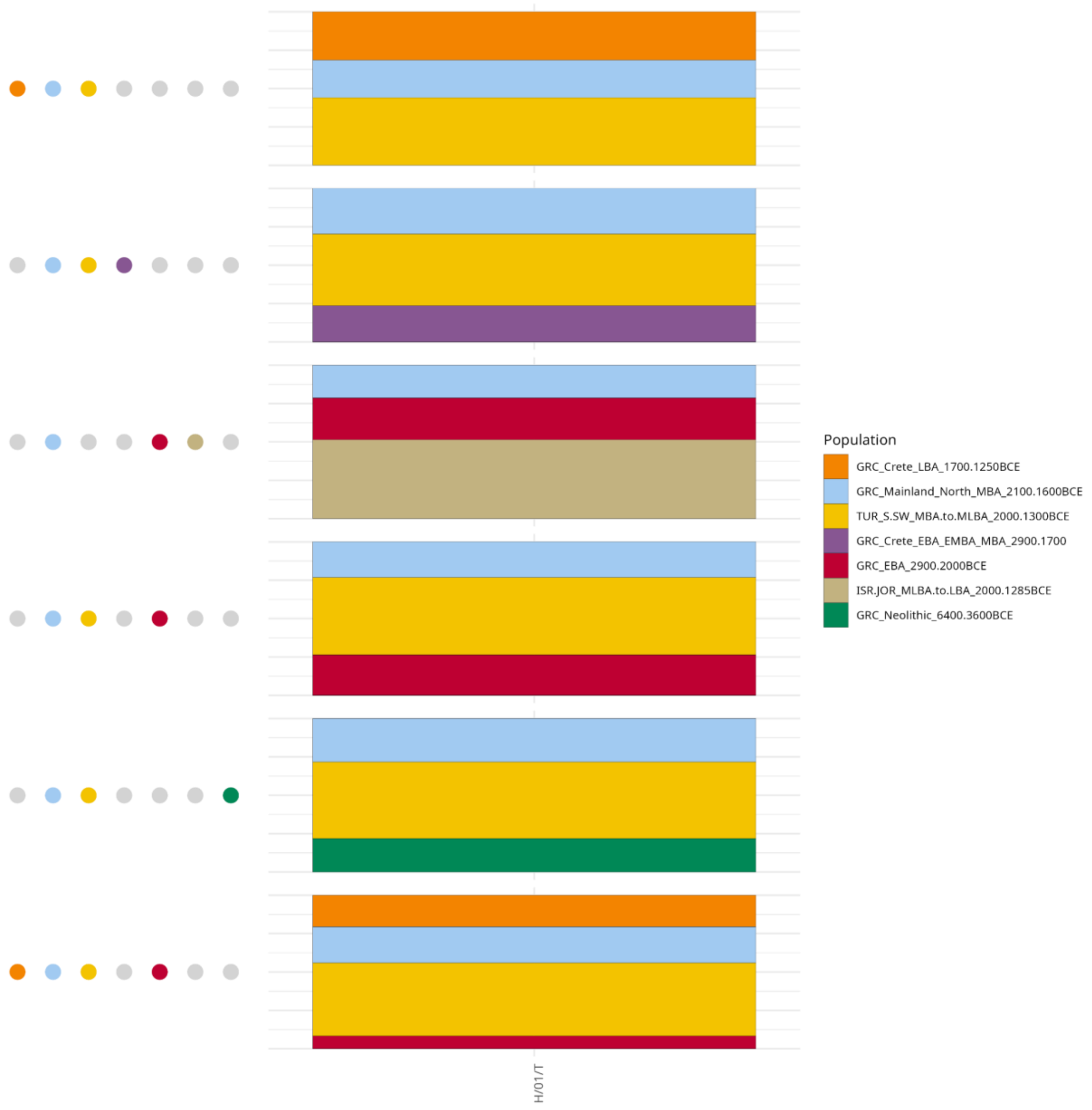


**Supplementary Figure S39.** Scatterplot of qpAdm analysis results at the population level using the “More_proximate” sample-set as putative sources, using the parameterization scheme “*adjust_pseudohaploid: TRUE - maxmmiss: 0.10 - afprod: TRUE*”. Only the feasible models with a p-value > 0.05 (accepted models) are depicted with the points representing the populations involved, and the colors indicating their admixture proportions. The models were sorted based on the number of involved sources (1-4). The tested target population is **A.** LBA Ammotopos **B.** Archaic Amvrakia, **C.** Classical Amvrakia **D.** Hellenistic Amvrakia **E.** Archaic Tenea **F.** Hellenistic Tenea **G.** Roman Tenea. The analyses for Archaic Amvrakia did not produce any feasible model. Archaic Amvrakia did not yield any feasible model.

**A.**

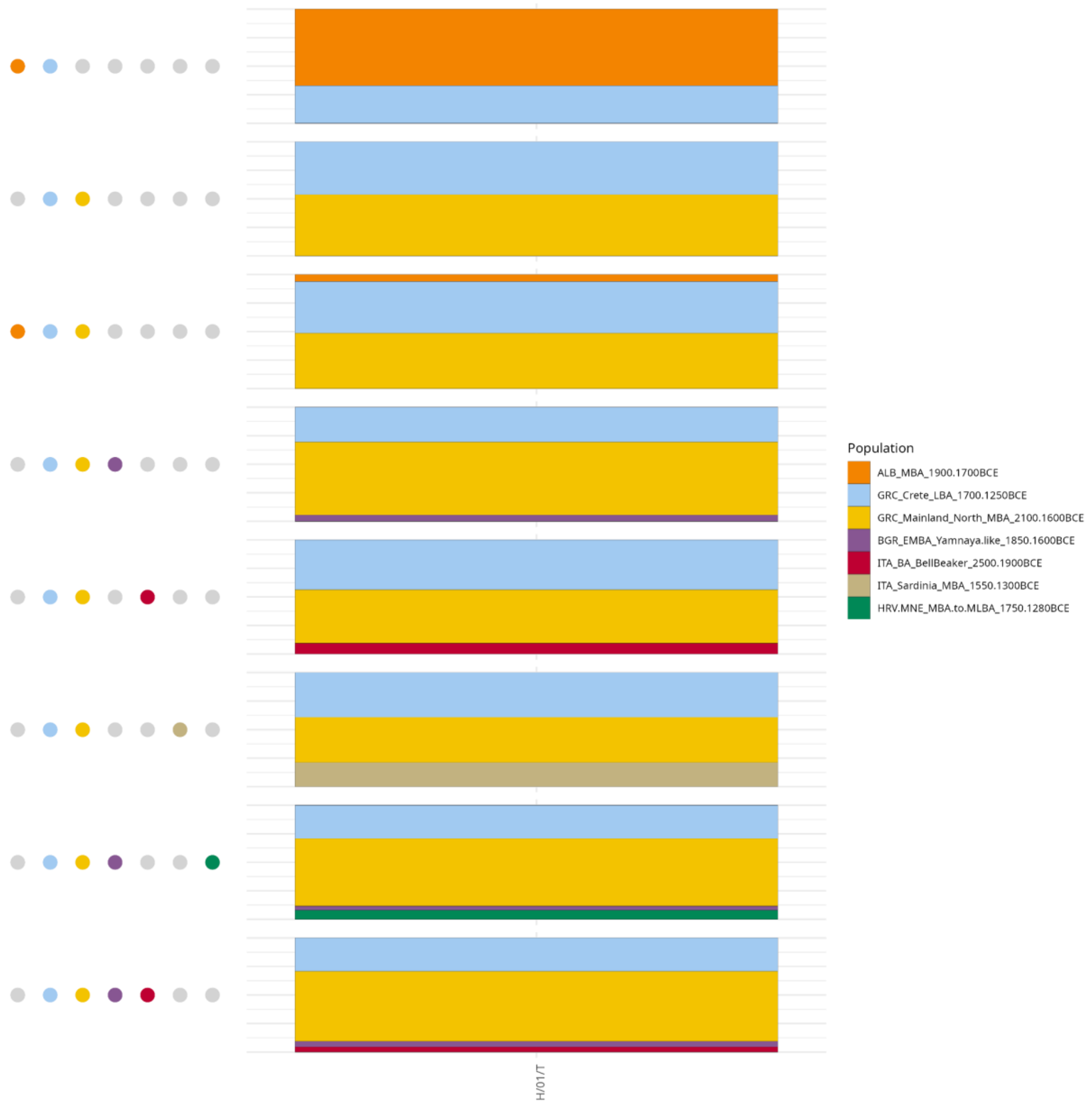

Population
ALB_MBA_1900.1700BCE
GRC_Crete_LBA_1700.1250BCE
GRC_Mainland_North_MBA_2100.1600BCE
BGR_EMBA_Yamnaya.like_1850.1600BCE
ITA_BA_BellBeaker_2500.1900BCE
ITA_Sardinia_MBA_1550.1300BCE
HRV.MNE_MBA.to.MLBA_1750.1280BCE
H/01/T

**B.**

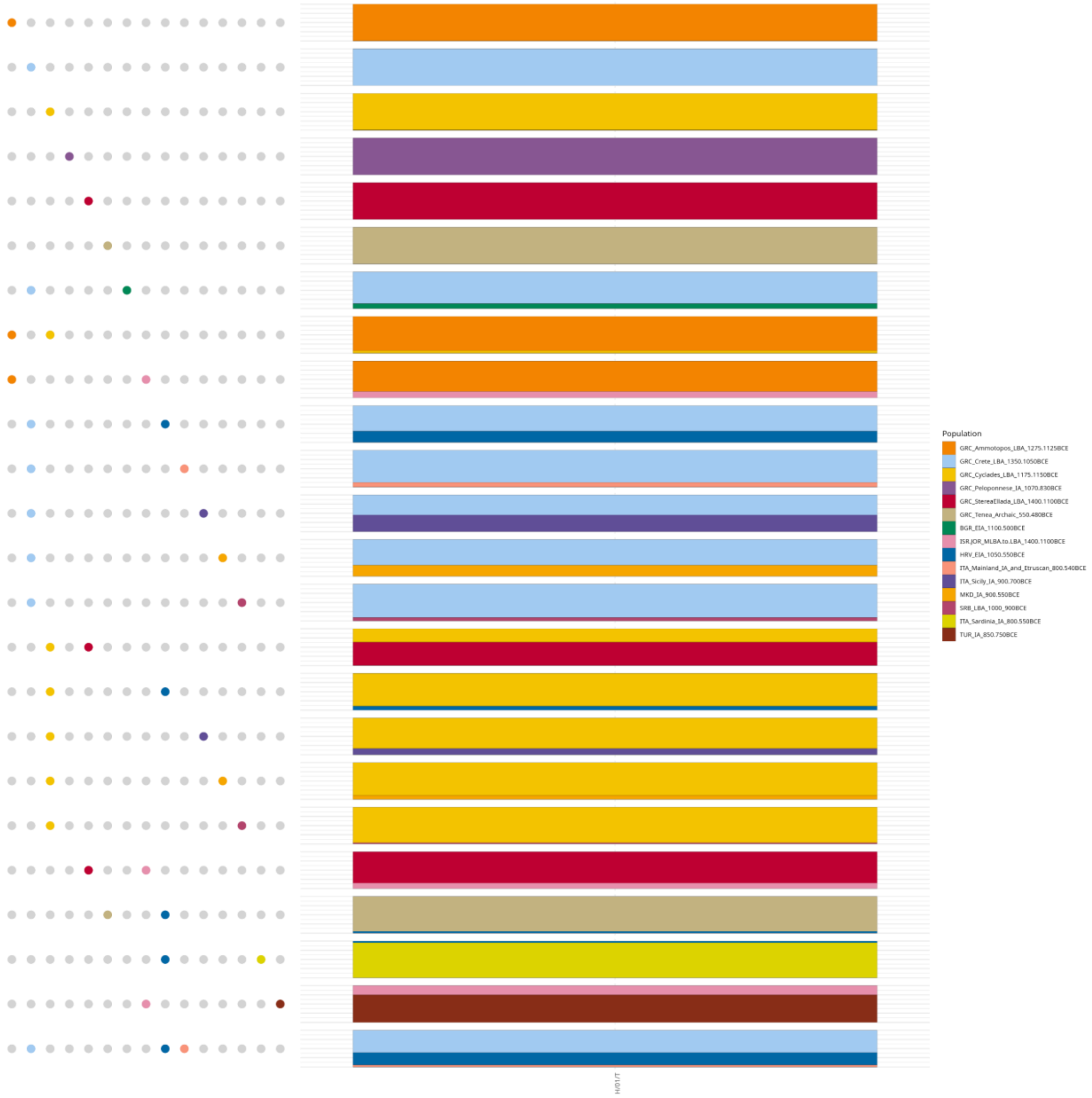

Population
GRC_Ammotopos_LBA_1275.1125BCE
GRC_Crete_LBA_1350.1050BCE
GRC_Cyclades_LBA_1175.1150BCE
GRC_Peloponnese_IA_1070.830BCE
GRC_StereaEllada_LBA_1400.1100BCE
GRC_Tenea_Archaic_550.480BCE
BGR_EIA_1100.500BCE
ISR.JOR_MLBA.to.LBA_1400.1100BCE
HRV_EIA_1050.550BCE
ITA_Mainland_IA_and_Etruscan_800.540BCE
ITA_Sicily_IA_900.700BCE
MKD_IA_900.550BCE
SRB_LBA_1000_900BCE
ITA_Sardinia_IA_800.550BCE
TUR_IA_850.750BCE
H/01/T

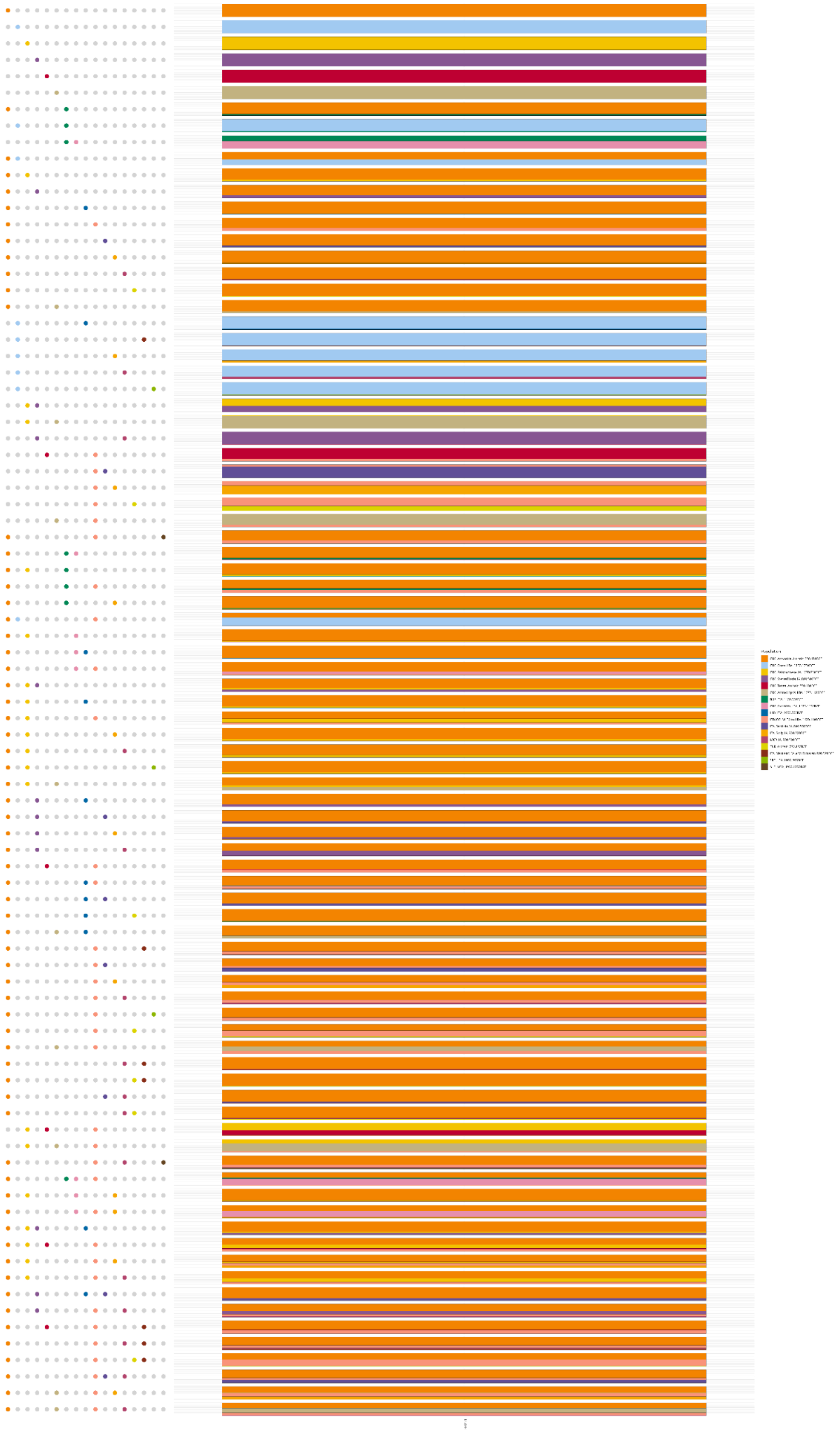


1774 **C.**

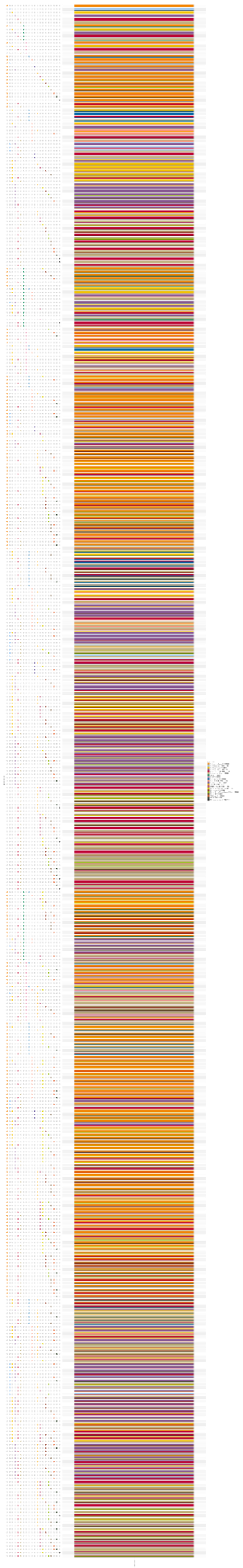

1775 **D.**

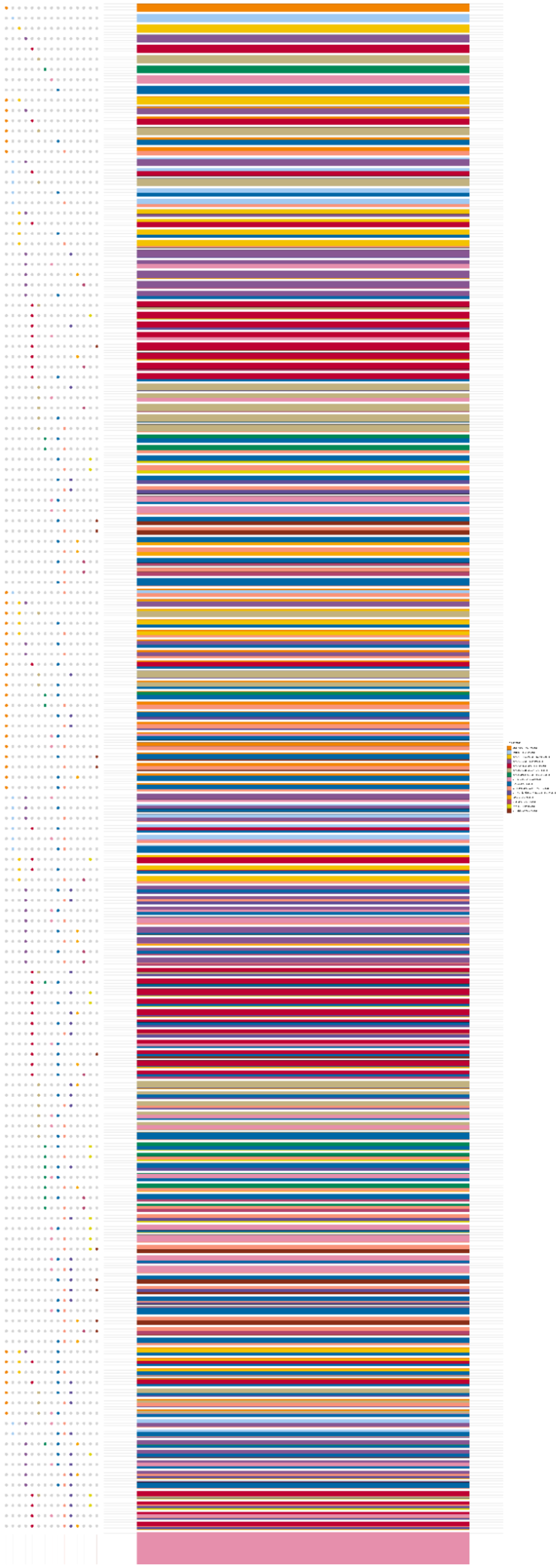

1776

**E.**

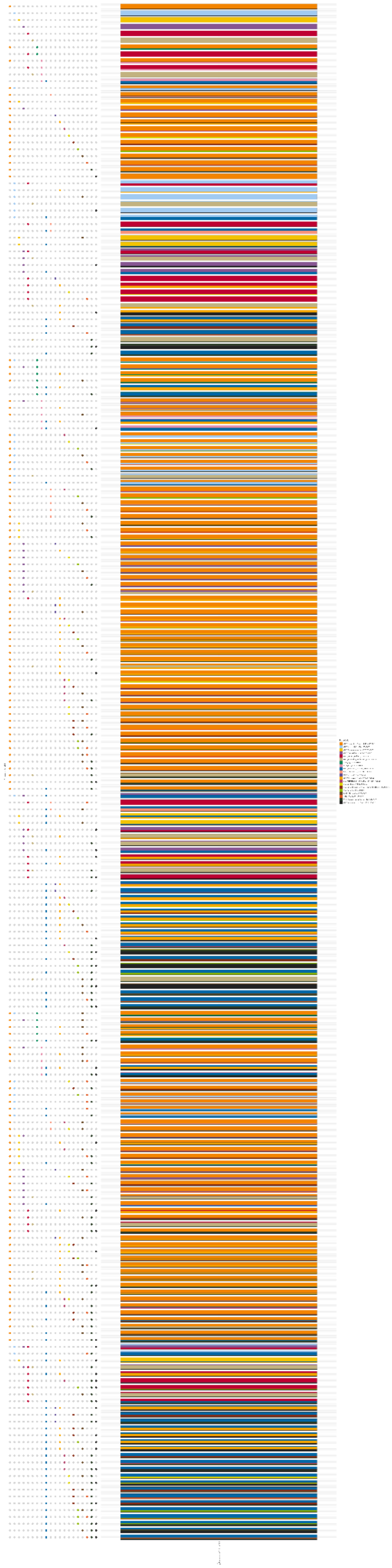

1777 **F.**

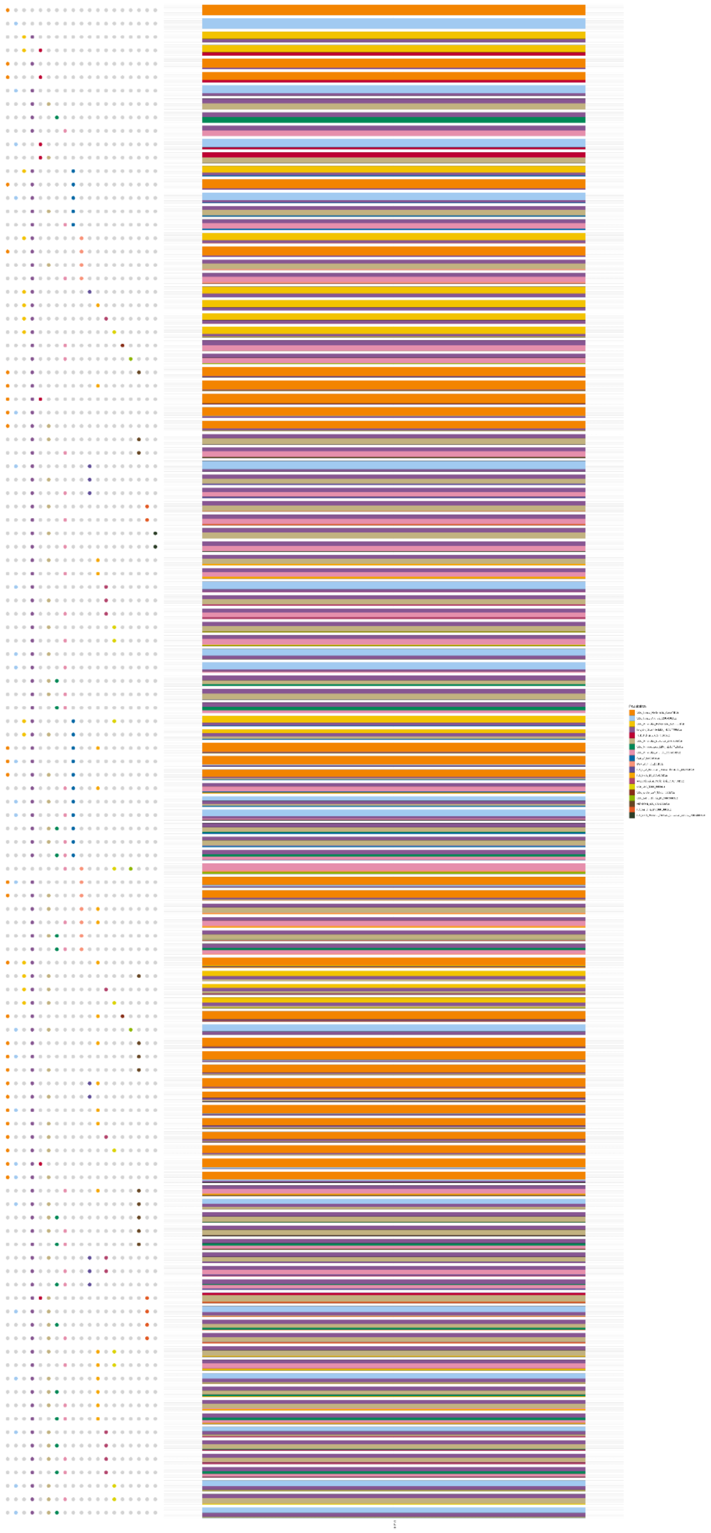

**G.**

**Supplementary Figure S40.** Scatterplot of qpAdm analyses results at the population level using the “Most_proximate” sample-set as putative sources, using the parameterization scheme “*adjust_pseudohaploid: TRUE - maxmmiss: 0.10 - afprod: TRUE*”. Only the feasible models with a p-value > 0.05 (accepted models) are depicted with the points representing the populations involved and the colors indicating their admixture proportions. The models were sorted based on the number of involved sources (1-4). **A.** LBA Ammotopos **B.** Archaic Amvrakia, **C.** Classical Amvrakia **D.** Hellenistic Amvrakia **E.** Archaic Tenea **F.** Hellenistic Tenea **G.** Roman Tenea.

The f3 analyses were performed using the R interface [142] of ADMIXTOOLS2 v.2.0.0, again deploying, as separate targets, the focal populations (Ammotopos, Amvrakia, Tenea) during different time periods (LBA, Archaic, Classical, Hellenistic, Roman, etc). For Admixture f3 analyses we used as potential sources of each target all possible putative pairs with the “Most-proximate” sources, in order to investigate if each one of the targets is the product of admixture (of equal genetic contribution, more or less) between the members of a given pair. Note that, rejection of the test, does not indicate absence of admixture. The Outgroup f3 analyses, on the other hand, were performed in order to estimate the genetic distance between a given target and a given set of other populations (members of the “Most-proximate” sources, as well as contemporary and subsequent populations of each target, i.e., for Classical Amvrakia we estimated the genetic distance to Archaic Amvrakia, but also to Hellenistic Amvrakia, Roman Tenea etc; see **Additional file 4** for details). As an outgroup population we used, in separate runs, the modern African Yoruba population and the modern East Asian Han population. Additionally, within-population genetic similarity levels were estimated by calculating the pair-wise Outgroup f3 values within each population and within a given period (**Figure 4B**). In calculations including outgroups, a higher f3 value indicates that the target is more similar to other tested population(s). The script is provided at https://doi.org/10.5281/zenodo.10848927 and includes the following R packages: Hmisc v.5.1-2 [143] and stringr. The Admixture f3 and the Outgroup f3 results are plotted in **Supplementary Figure S41** and **Supplementary Figure S42**, respectively, whereas all the Admixture and Outgroup f3 outputs, are provided in https://doi.org/10.5281/zenodo.10848927. All f3 calculations were performed using f2-block computations (*f2_from_geno*; https://uqrmaie1.github.io/admixtools/reference/f2_from_geno.html) and f2-blocks were calculated distinctly for all population triplets. Parameter *maxmiss* was set to 0.1 (in order to be consistent with qpAdm parameters) and *adjust_pseudohaploid* was set to TRUE as pou data are pseudohaploidized. In order to determine f3 values for the newly sequenced genomes, each individual was singled out and f3 calculations were repeated for each individual being the only representative of the population (**Supplementary Figure S42**, gray dots). Hence, individual estimates were computed for all samples. In the cases of close genetic relatedness (1st and 2nd degrees), the population estimate (**Supplementary Figure S42**, red dots) was computed by only retaining the genome with higher coverage.

No negative f3 value was observed in any of the pairwise Admixture f3 tests, indicating that we cannot statistically infer the presence of admixture between any given triplet of target and source pair, albeit the existence of admixture cannot be ruled out (see also simulation results above).

Regarding the Outgroup f3 tests, we observed similar results using either Yoruba or Han as the outgroup population, with the differences mostly being concentrated in the more distant populations in relation to each target. Overall, in most cases, the target population had the highest genetic similarity with populations from the geographic area of present-day Greece, indicating a general genetic continuity.

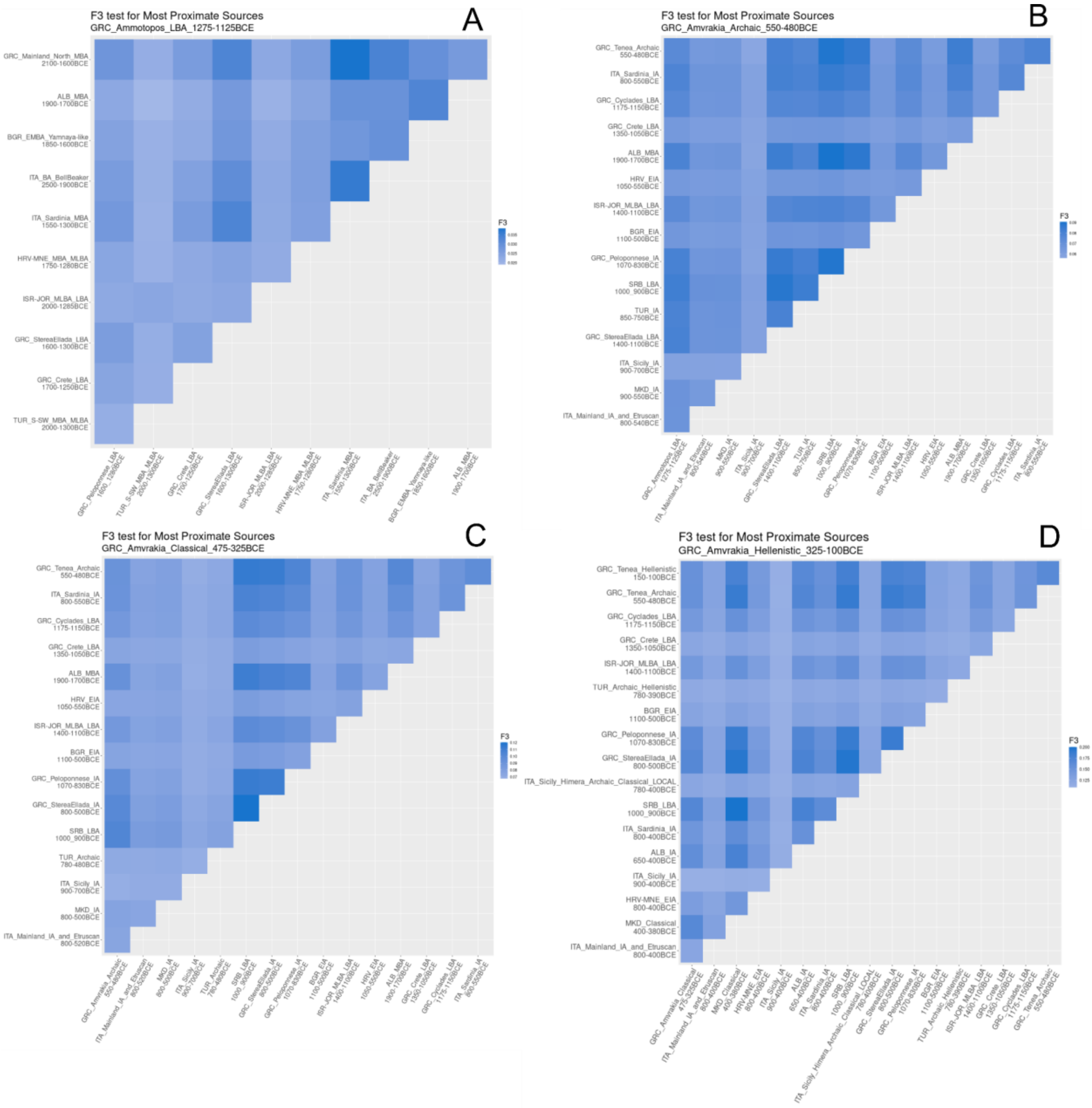

A
F3 test for Most Proximate Sources
GRC_Ammotopos_LBA_1275-1125BCE
GRC_Mainland_North_MBA 2100-1600BCE
ALB_MBA 1900-1700BCE
BGR_EMBA_Yamnaya-like 1850-1600BCE
ITA_BA_BellBeaker 2500-1900BCE
ITA_Sardinia_MBA 1550-1300BCE
HRV-MNE_MBA_MLBA 1750-1280BCE
ISR-JOR_MLBA_LBA 2000-1285BCE
GRC_StereaEllada_LBA 1600-1300BCE
GRC_Crete_LBA 1700-1250BCE
TUR_S-SW_MBA_MLBA 2000-1300BCE
F3
B
F3 test for Most Proximate Sources
GRC_Amvrakia_Archaic_550-480BCE
GRC_Tenea_Archaic 550-480BCE
ITA_Sardinia_IA 800-550BCE
GRC_Cyclades_LBA 1175-1150BCE
GRC_Crete_LBA 1350-1050BCE
ALB_MBA 1900-1700BCE
HRV_EIA 1050-550BCE
ISR-JOR_MLBA_LBA 1400-1100BCE
BGR_EIA 1100-500BCE
GRC_Peloponnese_IA 1070-830BCE
SRB_LBA 1000_900BCE
TUR_IA 850-750BCE
GRC_StereaEllada_LBA 1400-1100BCE
ITA_Sicily_IA 900-700BCE
MKD_IA 900-550BCE
ITA_Mainland_IA_and_Etruscan 800-540BCE
F3
C
F3 test for Most Proximate Sources
GRC_Amvrakia_Classical_475-325BCE
GRC_Tenea_Archaic 550-480BCE
ITA_Sardinia_IA 800-550BCE
GRC_Cyclades_LBA 1175-1150BCE
GRC_Crete_LBA 1350-1050BCE
ALB_MBA 1900-1700BCE
HRV_EIA 1050-550BCE
ISR-JOR_MLBA_LBA 1400-1100BCE
BGR_EIA 1100-500BCE
GRC_Peloponnese_IA 1070-830BCE
GRC_StereaEllada_IA 800-500BCE
SRB_LBA 1000_900BCE
TUR_Archaic 780-480BCE
ITA_Sicily_IA 900-700BCE
MKD_IA 800-500BCE
ITA_Mainland_IA_and_Etruscan 800-520BCE
F3
D
F3 test for Most Proximate Sources
GRC_Amvrakia_Hellenistic_325-100BCE
GRC_Tenea_Hellenistic 150-100BCE
GRC_Tenea_Archaic 550-480BCE
GRC_Cyclades_LBA 1175-1150BCE
GRC_Crete_LBA 1350-1050BCE
ISR-JOR_MLBA_LBA 1400-1100BCE
TUR_Archaic_Hellenistic 780-390BCE
BGR_EIA 1100-500BCE
GRC_Peloponnese_IA 1070-830BCE
GRC_StereaEllada_IA 800-500BCE
ITA_Sicily_Himera_Archaic_Classical_LOCAL 780-400BCE
SRB_LBA 1000_900BCE
ITA_Sardinia_IA 800-400BCE
ALB_IA 650-400BCE
ITA_Sicily_IA 900-400BCE
HRV-MNE_EIA 800-400BCE
MKD_Classical 400-380BCE
ITA_Mainland_IA_and_Etruscan 800-400BCE
F3

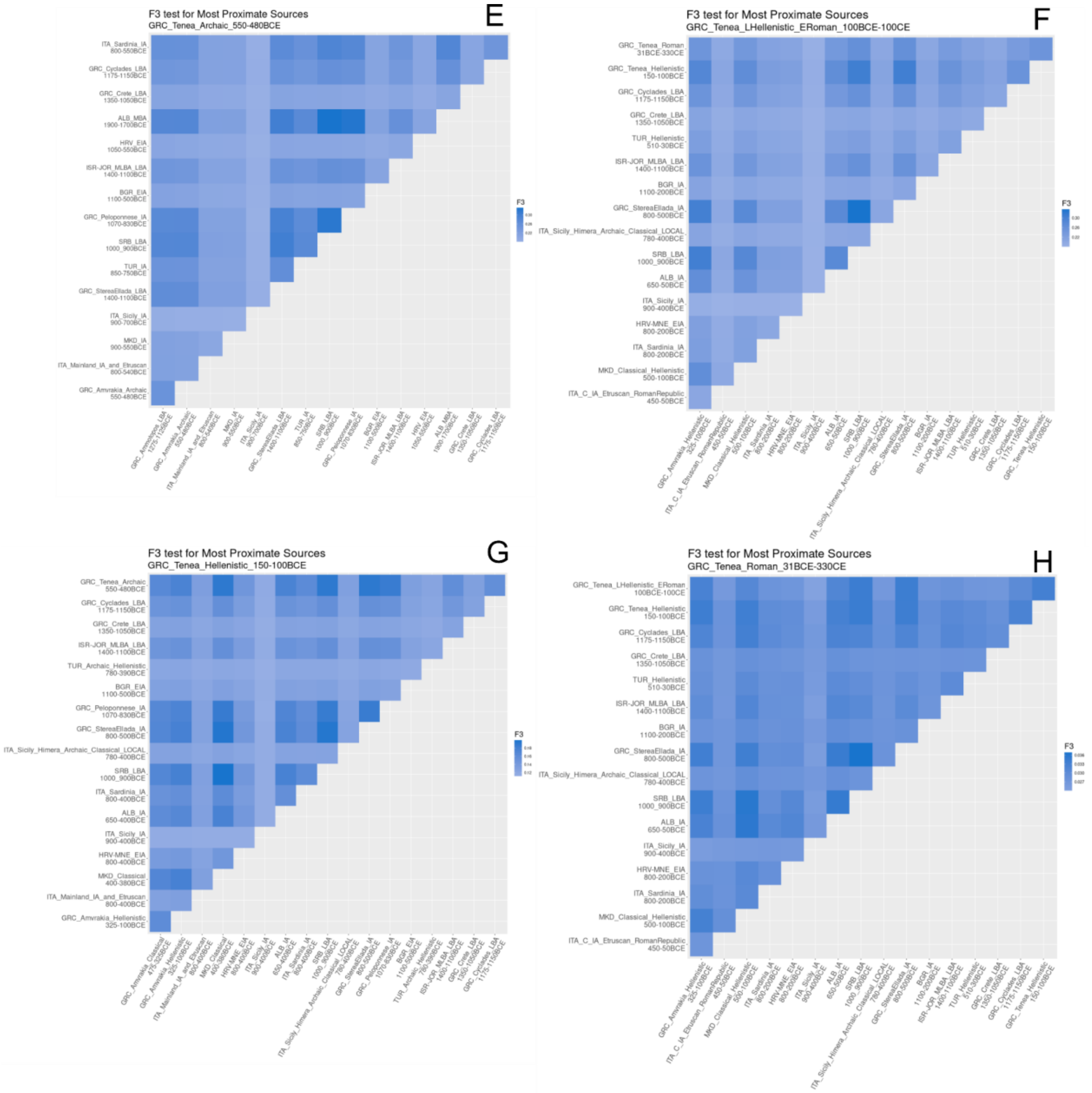


**Supplementary Figure S41.** Pairwise Admixture f3 plots using as potential sources of each target, all possible putative pairs with the "Most-proximate" sources, testing if each one of the targets might be the product of admixture between the members of a given pair (when f3 value is statistically significant negative). Targets are **A**) LBA Ammotopos, **B**) Archaic Amvrakia, **C**) Classical Amvrakia, **D**) Hellenistic Amvrakia, **E**) Archaic Tenea, **F**) Late Hellenistic - Early Roman Tenea, **G**) Hellenistic Tenea, and **H**) Roman Tenea. Abbreviations are given in **Figure 2**.

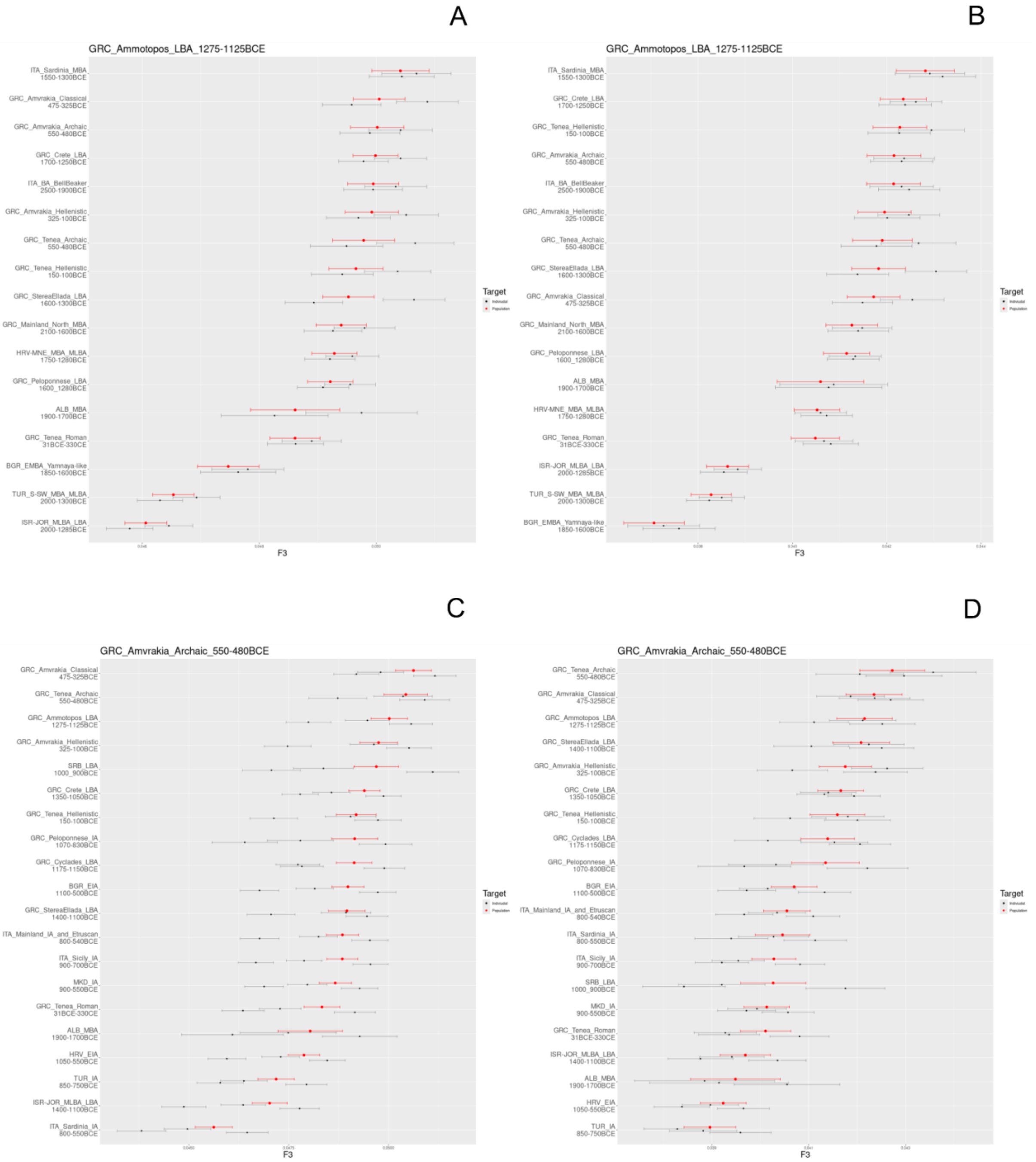
A
GRC_Ammotopos_LBA_1275-1125BCE
ITA_Sardinia_MBA 1550-1300BCE
GRC_Amvrakia_Classical 475-325BCE
GRC_Amvrakia_Archaic 550-480BCE
GRC_Crete_LBA 1700-1250BCE
ITA_BA_BellBeaker 2500-1900BCE
GRC_Amvrakia_Hellenistic 325-100BCE
GRC_Tenea_Archaic 550-480BCE
GRC_Tenea_Hellenistic 150-100BCE
GRC_StereaEllada_LBA 1600-1300BCE
GRC_Mainland_North_MBA 2100-1600BCE
HRV-MNE_MBA_MLBA 1750-1280BCE
GRC_Peloponnese_LBA 1600_1280BCE
ALB_MBA 1900-1700BCE
GRC_Tenea_Roman 31BCE-330CE
BGR_EMBA_Yamnaya-like 1850-1600BCE
TUR_S-SW_MBA_MLBA 2000-1300BCE
ISR-JOR_MLBA_LBA 2000-1285BCE
F3
B
GRC_Ammotopos_LBA_1275-1125BCE
ITA_Sardinia_MBA 1550-1300BCE
GRC_Crete_LBA 1700-1250BCE
GRC_Tenea_Hellenistic 150-100BCE
GRC_Amvrakia_Archaic 550-480BCE
ITA_BA_BellBeaker 2500-1900BCE
GRC_Amvrakia_Hellenistic 325-100BCE
GRC_Tenea_Archaic 550-480BCE
GRC_StereaEllada_LBA 1600-1300BCE
GRC_Amvrakia_Classical 475-325BCE
GRC_Mainland_North_MBA 2100-1600BCE
GRC_Peloponnese_LBA 1600_1280BCE
ALB_MBA 1900-1700BCE
HRV-MNE_MBA_MLBA 1750-1280BCE
GRC_Tenea_Roman 31BCE-330CE
ISR-JOR_MLBA_LBA 2000-1285BCE
TUR_S-SW_MBA_MLBA 2000-1300BCE
BGR_EMBA_Yamnaya-like 1850-1600BCE
F3
Target
C
GRC_Amvrakia_Archaic_550-480BCE
GRC_Amvrakia_Classical 475-325BCE
GRC_Tenea_Archaic 550-480BCE
GRC_Ammotopos_LBA 1275-1125BCE
GRC_Amvrakia_Hellenistic 325-100BCE
SRB_LBA 1000_900BCE
GRC_Crete_LBA 1350-1050BCE
GRC_Tenea_Hellenistic 150-100BCE
GRC_Peloponnese_IA 1070-830BCE
GRC_Cyclades_LBA 1175-1150BCE
BGR_EIA 1100-500BCE
GRC_StereaEllada_LBA 1400-1100BCE
ITA_Mainland_IA_and_Etruscan 800-540BCE
ITA_Sicily_IA 900-700BCE
MKD_IA 900-550BCE
GRC_Tenea_Roman 31BCE-330CE
ALB_MBA 1900-1700BCE
HRV_EIA 1050-550BCE
TUR_IA 850-750BCE
ISR-JOR_MLBA_LBA 1400-1100BCE
ITA_Sardinia_IA 800-550BCE
F3
Target
D
GRC_Amvrakia_Archaic_550-480BCE
GRC_Tenea_Archaic 550-480BCE
GRC_Amvrakia_Classical 475-325BCE
GRC_Ammotopos_LBA 1275-1125BCE
GRC_StereaEllada_LBA 1400-1100BCE
GRC_Amvrakia_Hellenistic 325-100BCE
GRC_Crete_LBA 1350-1050BCE
GRC_Tenea_Hellenistic 150-100BCE
GRC_Cyclades_LBA 1175-1150BCE
GRC_Peloponnese_IA 1070-830BCE
BGR_EIA 1100-500BCE
ITA_Mainland_IA_and_Etruscan 800-540BCE
ITA_Sardinia_IA 800-550BCE
ITA_Sicily_IA 900-700BCE
SRB_LBA 1000_900BCE
MKD_IA 900-550BCE
GRC_Tenea_Roman 31BCE-330CE
ISR-JOR_MLBA_LBA 1400-1100BCE
ALB_MBA 1900-1700BCE
HRV_EIA 1050-550BCE
TUR_IA 850-750BCE
F3
Target

E

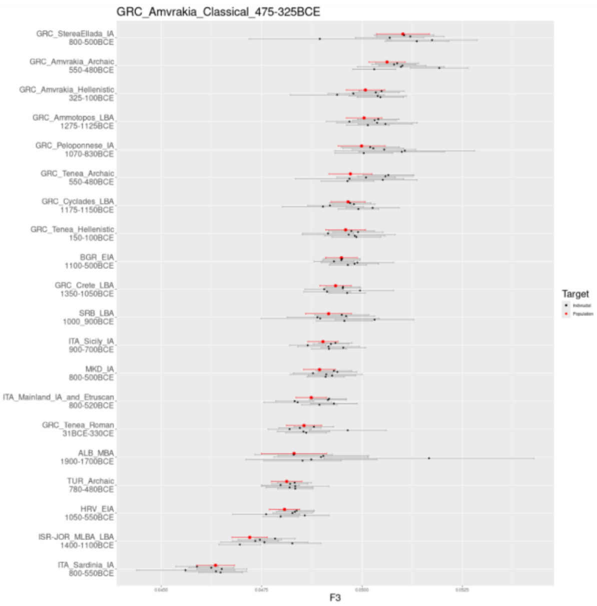
GRC_Amvrakia_Classical_475-325BCE
GRC_StereaEllada_IA 800-500BCE
GRC_Amvrakia_Archaic 550-480BCE
GRC_Amvrakia_Hellenistic 325-100BCE
GRC_Ammotopos_LBA 1275-1125BCE
GRC_Peloponnese_IA 1070-830BCE
GRC_Tenea_Archaic 550-480BCE
GRC_Cyclades_LBA 1175-1150BCE
GRC_Tenea_Hellenistic 150-100BCE
BGR_EIA 1100-500BCE
GRC_Crete_LBA 1350-1050BCE
SRB_LBA 1000_900BCE
ITA_Sicily_IA 900-700BCE
MKD_IA 800-500BCE
ITA_Mainland_IA_and_Etruscan 800-520BCE
GRC_Tenea_Roman 31BCE-330CE
ALB_MBA 1900-1700BCE
TUR_Archaic 780-480BCE
HRV_EIA 1050-550BCE
ISR-JOR_MLBA_LBA 1400-1100BCE
ITA_Sardinia_IA 800-550BCE
F3
Target

F

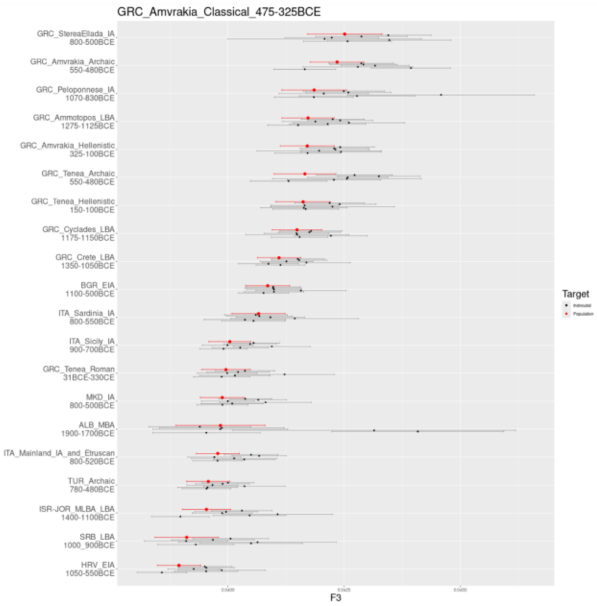
GRC_Amvrakia_Classical_475-325BCE
GRC_StereaEllada_IA 800-500BCE
GRC_Amvrakia_Archaic 550-480BCE
GRC_Peloponnese_IA 1070-830BCE
GRC_Ammotopos_LBA 1275-1125BCE
GRC_Amvrakia_Hellenistic 325-100BCE
GRC_Tenea_Archaic 550-480BCE
GRC_Tenea_Hellenistic 150-100BCE
GRC_Cyclades_LBA 1175-1150BCE
GRC_Crete_LBA 1350-1050BCE
BGR_EIA 1100-500BCE
ITA_Sardinia_IA 800-550BCE
ITA_Sicily_IA 900-700BCE
GRC_Tenea_Roman 31BCE-330CE
MKD_IA 800-500BCE
ALB_MBA 1900-1700BCE
ITA_Mainland_IA_and_Etruscan 800-520BCE
TUR_Archaic 780-480BCE
ISR-JOR_MLBA_LBA 1400-1100BCE
SRB_LBA 1000_900BCE
HRV_EIA 1050-550BCE
F3
Target

G

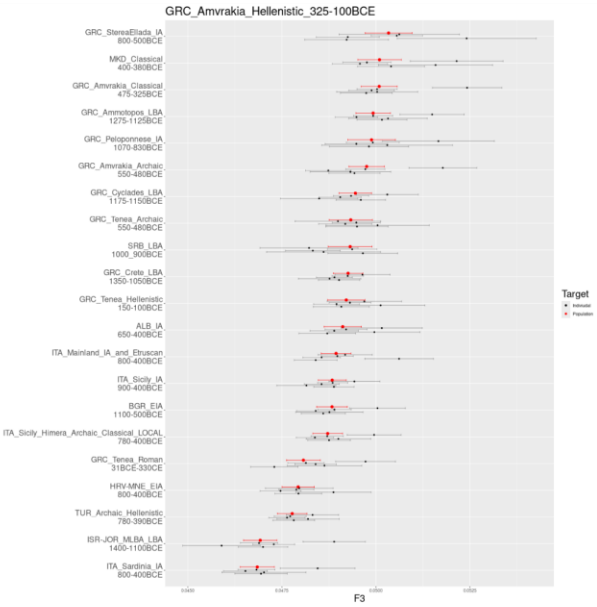
GRC_Amvrakia_Hellenistic_325-100BCE
GRC_StereaEllada_IA 800-500BCE
MKD_Classical 400-380BCE
GRC_Amvrakia_Classical 475-325BCE
GRC_Ammotopos_LBA 1275-1125BCE
GRC_Peloponnese_IA 1070-830BCE
GRC_Amvrakia_Archaic 550-480BCE
GRC_Cyclades_LBA 1175-1150BCE
GRC_Tenea_Archaic 550-480BCE
SRB_LBA 1000_900BCE
GRC_Crete_LBA 1350-1050BCE
GRC_Tenea_Hellenistic 150-100BCE
ALB_IA 650-400BCE
ITA_Mainland_IA_and_Etruscan 800-400BCE
ITA_Sicily_IA 900-400BCE
BGR_EIA 1100-500BCE
ITA_Sicily_Himera_Archaic_Classical_LOCAL 780-400BCE
GRC_Tenea_Roman 31BCE-330CE
HRV-MNE_EIA 800-400BCE
TUR_Archaic_Hellenistic 780-390BCE
ISR-JOR_MLBA_LBA 1400-1100BCE
ITA_Sardinia_IA 800-400BCE
F3
Target

H

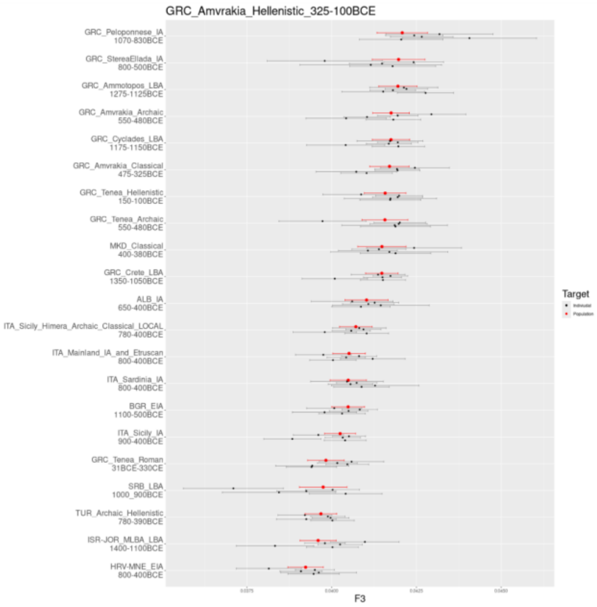
GRC_Amvrakia_Hellenistic_325-100BCE
GRC_Peloponnese_IA 1070-830BCE
GRC_StereaEllada_IA 800-500BCE
GRC_Ammotopos_LBA 1275-1125BCE
GRC_Amvrakia_Archaic 550-480BCE
GRC_Cyclades_LBA 1175-1150BCE
GRC_Amvrakia_Classical 475-325BCE
GRC_Tenea_Hellenistic 150-100BCE
GRC_Tenea_Archaic 550-480BCE
MKD_Classical 400-380BCE
GRC_Crete_LBA 1350-1050BCE
ALB_IA 650-400BCE
ITA_Sicily_Himera_Archaic_Classical_LOCAL 780-400BCE
ITA_Mainland_IA_and_Etruscan 800-400BCE
ITA_Sardinia_IA 800-400BCE
BGR_EIA 1100-500BCE
ITA_Sicily_IA 900-400BCE
GRC_Tenea_Roman 31BCE-330CE
SRB_LBA 1000_900BCE
TUR_Archaic_Hellenistic 780-390BCE
ISR-JOR_MLBA_LBA 1400-1100BCE
HRV-MNE_EIA 800-400BCE
F3
Target

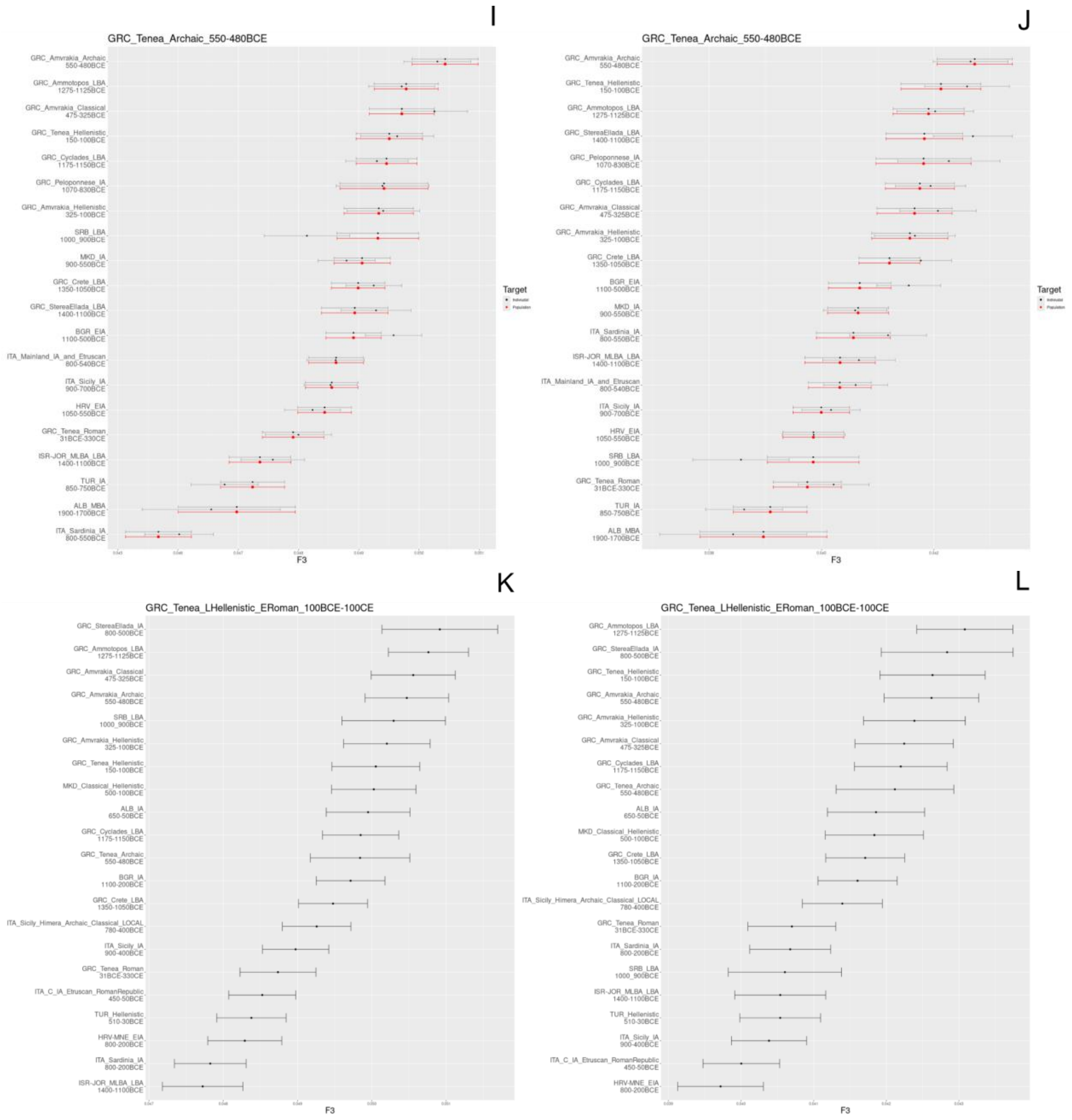
I
GRC_Tenea_Archaic_550-480BCE
GRC_Amvrakia_Archaic 550-480BCE
GRC_Ammotopos_LBA 1275-1125BCE
GRC_Amvrakia_Classical 475-325BCE
GRC_Tenea_Hellenistic 150-100BCE
GRC_Cyclades_LBA 1175-1150BCE
GRC_Peloponnese_IA 1070-830BCE
GRC_Amvrakia_Hellenistic 325-100BCE
SRB_LBA 1000_900BCE
MKD_IA 900-550BCE
GRC_Crete_LBA 1350-1050BCE
GRC_StereaEllada_LBA 1400-1100BCE
BGR_EIA 1100-500BCE
ITA_Mainland_IA_and_Etruscan 800-540BCE
ITA_Sicily_IA 900-700BCE
HRV_EIA 1050-550BCE
GRC_Tenea_Roman 31BCE-330CE
ISR-JOR_MLBA_LBA 1400-1100BCE
TUR_IA 850-750BCE
ALB_MBA 1900-1700BCE
ITA_Sardinia_IA 800-550BCE
F3
Target
J
GRC_Tenea_Archaic_550-480BCE
GRC_Amvrakia_Archaic 550-480BCE
GRC_Tenea_Hellenistic 150-100BCE
GRC_Ammotopos_LBA 1275-1125BCE
GRC_StereaEllada_LBA 1400-1100BCE
GRC_Peloponnese_IA 1070-830BCE
GRC_Cyclades_LBA 1175-1150BCE
GRC_Amvrakia_Classical 475-325BCE
GRC_Amvrakia_Hellenistic 325-100BCE
GRC_Crete_LBA 1350-1050BCE
BGR_EIA 1100-500BCE
MKD_IA 900-550BCE
ITA_Sardinia_IA 800-550BCE
ISR-JOR_MLBA_LBA 1400-1100BCE
ITA_Mainland_IA_and_Etruscan 800-540BCE
ITA_Sicily_IA 900-700BCE
HRV_EIA 1050-550BCE
SRB_LBA 1000_900BCE
GRC_Tenea_Roman 31BCE-330CE
TUR_IA 850-750BCE
ALB_MBA 1900-1700BCE
F3
Target
K
GRC_Tenea_LHellenistic_ERoman_100BCE-100CE
GRC_StereaEllada_IA 800-500BCE
GRC_Ammotopos_LBA 1275-1125BCE
GRC_Amvrakia_Classical 475-325BCE
GRC_Amvrakia_Archaic 550-480BCE
SRB_LBA 1000_900BCE
GRC_Amvrakia_Hellenistic 325-100BCE
GRC_Tenea_Hellenistic 150-100BCE
MKD_Classical_Hellenistic 500-100BCE
ALB_IA 650-50BCE
GRC_Cyclades_LBA 1175-1150BCE
GRC_Tenea_Archaic 550-480BCE
BGR_IA 1100-200BCE
GRC_Crete_LBA 1350-1050BCE
ITA_Sicily_Himera_Archaic_Classical_LOCAL 780-400BCE
ITA_Sicily_IA 900-400BCE
GRC_Tenea_Roman 31BCE-330CE
ITA_C_IA_Etruscan_RomanRepublic 450-50BCE
TUR_Hellenistic 510-30BCE
HRV-MNE_EIA 800-200BCE
ITA_Sardinia_IA 800-200BCE
ISR-JOR_MLBA_LBA 1400-1100BCE
F3
L
GRC_Tenea_LHellenistic_ERoman_100BCE-100CE
GRC_Ammotopos_LBA 1275-1125BCE
GRC_StereaEllada_IA 800-500BCE
GRC_Tenea_Hellenistic 150-100BCE
GRC_Amvrakia_Archaic 550-480BCE
GRC_Amvrakia_Hellenistic 325-100BCE
GRC_Amvrakia_Classical 475-325BCE
GRC_Cyclades_LBA 1175-1150BCE
GRC_Tenea_Archaic 550-480BCE
ALB_IA 650-50BCE
MKD_Classical_Hellenistic 500-100BCE
GRC_Crete_LBA 1350-1050BCE
BGR_IA 1100-200BCE
ITA_Sicily_Himera_Archaic_Classical_LOCAL 780-400BCE
GRC_Tenea_Roman 31BCE-330CE
ITA_Sardinia_IA 800-200BCE
SRB_LBA 1000_900BCE
ISR-JOR_MLBA_LBA 1400-1100BCE
TUR_Hellenistic 510-30BCE
ITA_Sicily_IA 900-400BCE
ITA_C_IA_Etruscan_RomanRepublic 450-50BCE
HRV-MNE_EIA 800-200BCE
F3

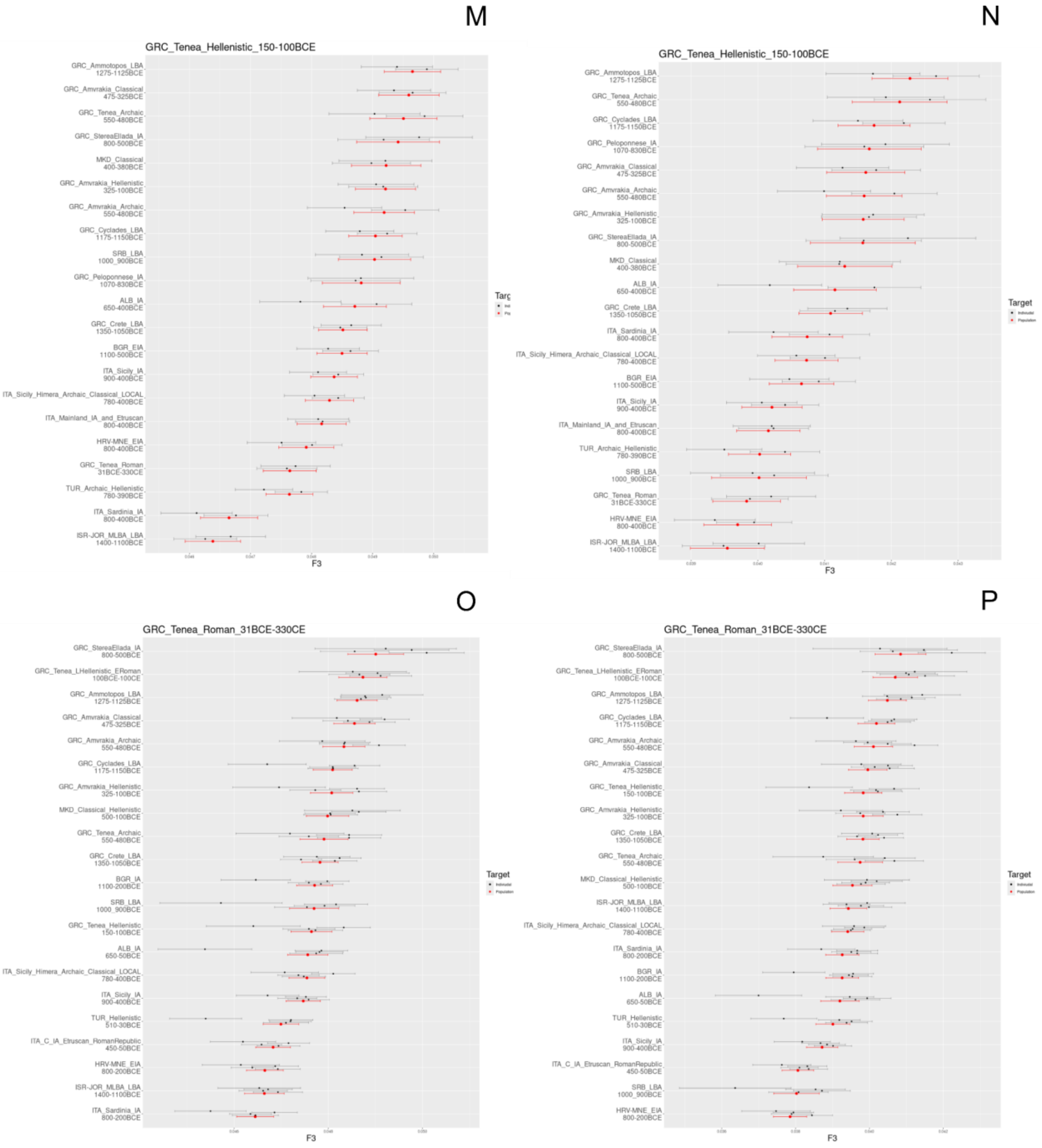


**Supplementary Figure S42.** Pairwise outgroup f3 tests between a given target and members of the “Most-proximate” sources set and contemporary and subsequent populations of each target, estimating the genetic distance between them (the higher the f3 value the lower the genetic distance). As outgroups we use the modern African Yoruba population (**left column plots**; **A**, **C**, **E**, **G**, **I**, **K**, **M**, **O**) and the modern Eastern Asia Han population (**right column plots**; **B**, **D**, **F**, **H**, **J**, **L**, **N**, **P**). Targets are **A-B**) LBA Ammotopos, **C-D**) Archaic Amvrakia, **E-F**) Classical Amvrakia, **G-H**) Hellenistic Amvrakia, **I-J**) Archaic Tenea, **K-L**) Late Hellenistic - Early Roman Tenea, **M-N**) Hellenistic Tenea, and **O-P**) Roman Tenea. Abbreviations are given in **Figure 2.**

#### 3.6 Phenotypes

***Angelos Souleles***

##### 3.6.1. Pigmentation

For hair, eye, and skin color prediction, we deployed the widely used HirisPlex-S [144,145] tool. We employed imputed data (see Section 3.5.5) for the 41 SNPs associated with HirisPlex-S, as recent studies have demonstrated that imputed data can reliably predict phenotypes, even for individuals with a coverage as low as 0.10-0.50× [146]. All newly generated genomes of the present study have >0.1× mean coverage depth. To minimize errors associated with reference-based imputation, we used BCFtools v.1.15 and its *filter* function to only retain genotypes from imputed SNPs if they had an INFO score of 0.50 or higher (`–e 'INFO/INFO<=0.5'`). For the remaining HIrisPlex-S SNPs with INFO score less than 0.50, we followed a similar approach as Marchi et al. [109]. We examined the BAM files directly using SAMtools v.1.15. We then created two HirisPlex-S input files for each individual: one by replacing missing genotypes with homozygous genotypes for the most abundant allele in the BAM file (*_main* in **Additional file 5**; **HIrisPlex-S_raw_results sheet**), and another one by replacing missing genotypes with heterozygous genotypes (*_secondary* in **Additional file 5**; **HIrisPlex-S_raw_results sheet**) to account for the uncertainty in heterozygosity observation associated with low-coverage aDNA data. However, for SNPs rs312262906 and rs201326893, we did not need to apply the aforementioned approach, because the presence of an alternative allele at each of the two SNPs sites predicts red hair with a probability of ~1.00 [145]. For both of these SNPs, we replaced missing genotypes with homozygous genotypes for the reference allele (0/0) for both main and secondary input files.

The color interpretations are detailed in **Additional file 5 (HIrisPlex-S_interpretations sheet)**. When both, the main, and the secondary probabilities exceeded 0.70, the predicted phenotype was accepted. In other cases, the phenotype with the highest and the second highest probability in the main run was accepted. This is based on the indication that the second most likely category can influence the main category phenotype [144,147]. Our approach reliably determined eye color, skin color, and hair shade for all individuals, with hair color results being available for 19 out of the 26 individuals. All individuals showed the highest probability for brown eyes. Most individuals likely had an intermediate skin tone, while three of them had a darker skin color (two from Classical Amvrakia and one from Roman Tenea). Similarly, most individuals likely had brown hair with a dark shade. Notably, despite our attempts to eliminate false positive results for red hair, the Late Hellenistic - Early Roman individual from Tenea still exhibited a high probability for this phenotype.

##### 3.6.2. Monogenic Phenotypes

Genotypes associated with metabolic traits (lactase persistence and sensitivity to fats) and human muscle strength and composition (muscle contraction type and muscle performance) were manually examined directly in the BAM files using SAMtools, as described above. The allele counts from the corresponding reads are provided in **Additional file 5** (**Monogenic_traits_counts sheet)**. For each genotype, the number of individuals covering the SNP varied (MCM6: n=14; FABP2: 15; ACTN3: 15; ACVR1B: 11). None of these individuals were found to be lactose tolerant, including the more recent ones. Seven individuals showed a moderately increased sensitivity to fats; however, four of them only had one single read at the specific genomic position, yielding these results inconclusive. Regarding muscle performance, eight individuals had an ACTN3 genotype associated with improved muscle performance as typically seen in sprinters (three of whom had only one read), whereas seven individuals likely had impaired muscle performance (three of these only had one read). Lastly, five individuals had higher muscle strength associated with the ACVR1B gene, with one individual only having one read. Additionally, we examined 29 SNPs associated with beta thalassemia (and malaria resistance), as it is the most common genetic disorder in modern Greece [148,149]. However, no alleles associated with beta thalassemia were found in any of the newly sequenced ancient Greek individuals.

#### 3.7 Microbial Metagenomics

***Nikolaos Psonis***

For the microbial metagenomics analyses, we used the FastQ files with the fully (non-truncated) collapsed reads as input that were produced after the residual adapter trimming step of the mapache pipeline (see **section 3.2.1** above). Hence, ancient microbial DNA screening analyses were performed at the FastQ level and not at the individual level. Due to its large size, one FastQ file (165_lys2_ex1_L1_fq_collapsed) had to be split into multiple ones of equal size (four in total), using fastqsplitter v.1.2.0 (https://github.com/LUMC/fastqsplitter) in order to be used in downstream metagenomic analyses. The taxonomic assignment (using a k-mer based approach) of each sequence was performed with v.1.0.4 KrakenUniq [150], a Lowest Common Ancestor (LCA) sequence alignment was performed with MALT v.0.61 [151], and authentication and validation of putative microbial species was conducted with the *MaltExtract* function of HOPS v.0.35 [152]. All the aforementioned software tools were used as implemented in the v.1.0.0 aMeta [153] pipeline by using the snakemake v.7.18.2 [58] workflow manager. As a reference database, we used the pre-built full NCBI non-redundant *nucleotide* (NT; https://www.ncbi.nlm.nih.gov/nucleotide/) database provided by aMeta. This database contains records until December 2020 and includes all microbial, vertebrate, non-vertebrate, and plant organisms.

No microbial DNA belonging to ancient systemic pathogens was detected in any of the samples examined (**Additional file 6;** metagenomic overview heatmap score >8). In dental samples, however, we did observe DNA traces belonging to ancient human oral bacteria (and/or are considered common dental pathogens causing diseases, such as gingivitis), including *Porphyromonas gingivalis, Tannerella forsythia, Streptococcus gordonii, Streptococcus anginosus, Streptococcus intermedius, Capnocytophaga sputigena, Eikenella corrodens, Neisseria elongata, Parvimonas micra, Streptococcus sanguinis, Tannerella* sp. oral taxon HOT-286, *Campylobacter showae, Gemella morbillorum, Neisseria mucosa, Aggregatibacter aphrophilus, Prevotella intermedia, Rothia dentocariosa, Streptococcus mutans, Aggregatibacter actinomycetemcomitans, Campylobacter rectus, Corynebacterium matruchotii, Fusobacterium nucleatum, Leptotrichia trevisanii*. Furthermore, the Human endogenous retrovirus K was detected (score of 8) in a Classical Amvrakia individual (Amv_Epi_Cl_1).

Moreover, authenticated DNA traces were obtained from environmental taxa, such as *Ralstonia solanacearum*, *Thermobispora bispora, Clostridium tetani, Streptosporangium roseum, Clostridium septicum, Candidatus Nitrososphaera gargensis, Alcaligenes faecalis, Proteus vulgaris, Acidipropionibacterium jensenii, Acinetobacter calcoaceticus, Advenella kashmirensis, Serratia rubidaea, Thermobispora bispora, Lysobacter gummosus, Frankia alni, Sanguibacter keddieii, Clostridium butyricum, Citrobacter freundii, Citrobacter braakii, Streptomyces malaysiensis*. Finally, there were cases that a human pathogen taxon was identified (e.g. *Spirometra erinaceieuropaei, Clostridium botulinum*), although in these cases they are considered false positives due to the increased genetic similarity with specific areas in the human genome, despite showing aDNA damage signatures (they largely differ, compared to the human genome, by transitions). In the latter group may be included *Paeniclostridium sordellii*, too, which was detected in Amv_Epi_Cl_5.

#### 3.8 Visualization

***Stefanos Papadantonakis, Angelos Souleles, Georgios Kousis Tsampazis, Angeliki Papadopoulou, Nikolaos Psonis***

The map of **Figure 1A** was created with R v.4.3.0 [154] using the packages rnaturalearth v.1.0.1.9000 [155], sf v.1.0-16 [156], proj4 v1.0-13 [157], ggplot2 [158] and ggrepel v.0.9.5 [159]. The mapache rulegraph (**Supplementary Figure S37)** was produced using snakemake. Plotting of ROHs (**Figure 5** and **Supplementary Figure S32)** and IBD (**Figure 3**) results was performed in Python v.3.8.1, using the seaborn v.0.12.1 [160] package. Plotting of PCA results (**Figure 2A**) was performed using R v.4.3.0 and the ggplot2 and ggmagnify v.0.4.1.9000 [161] packages. Plotting of Pandora results was performed online using Flourish (https://flourish.studio/). Plotting of ADMIXTURE (**Figure 2B** and **Supplementary Figures S34-S35**), qpAdm (**Figure 4** and **Supplementary Figures S36-S38**) and f3 (**Supplementary Figures S39-S40**) results was performed in R v.4.3.2 using the ggplot2 and ggtext v.0.1.2 [162] package. We merged multiple figures into a single one using Inkscape v.1.0.2-2 (https://inkscape.org).

### 4. Provenance, mobility and diet analysis using stable isotopes

***Argyro Nafplioti***

#### 4.1 Strontium isotope ratio analysis of bioarchaeological skeletal remains: principles

The strontium isotope ratio ($^{(87}$Sr/$^{86}$Sr) largely reflects local geology. As strontium isotopes in teeth are fixed in enamel biogenic apatite at the time of tooth formation and enamel undergoes little remodeling thereafter, the strontium isotope ratio values recorded reflect childhood provenance and provide evidence for geographical origins and potentially also for mobility [163–165]. Since the principles of $^{87}$Sr/$^{86}$Sr analysis in research of this kind are well documented [165–167] and have also been extensively discussed in earlier relevant work of one of the authors [e.g. 168,169,170], we only provide a summary in the following.

In nature, strontium occurs in the form of four stable isotopes, $^{87}$Sr (comprises c. 7.04% of total strontium), $^{88}$Sr (c. 82.53%), $^{86}$Sr (c. 9.87%), and $^{84}$Sr (c. 0.56%). The strontium isotope $^{87}$Sr is radiogenic and is the product of the radioactive decay of the rubidium isotope $^{87}$Rb, which has a half-life of approximately 47 billion years. All remaining three strontium isotopes are non-radiogenic [171]. Therefore, in any geology, the ratio of strontium isotope $^{87}$Sr to $^{86}$Sr depends on the relative abundance of rubidium and strontium at the time the rock crystallized and the age of the rocks [172]. Because rubidium is substantially more abundant in crustal materials than in the Earth’s mantle, old metamorphic rocks of crustal origin have higher $^{87}$Sr/$^{86}$Sr values (c. 0.715) than recent volcanic rocks (c. 0.704) [173]. Strontium isotope ratios in marine sedimentary rocks depend on the $^{87}$Sr/$^{86}$Sr value of seawater at the time they were formed and largely vary between 0.707 and 0.710 [174,175].

In essence, the $^{87}$Sr/$^{86}$Sr ratio largely reflects local geology, and passes from the bedrock into the soil, the groundwater, and the food chain. Thereby, $^{87}$Sr/$^{86}$Sr reaches the human skeletal tissues, where it substitutes for calcium in hydroxyapatite [171], largely from the food and water consumed with no fractionation related to biological processes [176,177]. Although other factors such as the proximity to marine environments and the $^{87}$Sr/$^{86}$Sr ratio in sea spray [178], atmospheric deposition [179] and in modern contexts fertilizers too [167,177], can also impact local $^{87}$Sr/$^{86}$Sr signatures, the latter largely reflect bedrock geology and mineral weathering. Thus, $^{87}$Sr/$^{86}$Sr signatures in human skeletal tissues match the geochemical profile of the catchment area of the individuals analyzed.

Because tooth enamel is a cell-free tissue that for most of the permanent dentition largely forms by the 8th year of life and does not remodel thereafter [164,180], $^{87}$Sr/$^{86}$Sr signatures from tooth enamel reflect early childhood diet and geographical origins. Conversely, bone and, to a lesser extent, also dentine, undergo continuous replacement of their mineral phase in the course of life. In addition, tooth enamel is denser, harder, and more inert than bone or dentine, and therefore more resistant to post-burial isotopic contamination than bone or dentine [163,181–184]. Thus, cortical bone $^{87}$Sr/$^{86}$Sr signatures more closely reflect the dietary intake of the last 10-20 years of life and human bone $^{87}$Sr/$^{86}$Sr values can be used to characterize local bioavailable $^{87}$Sr/$^{86}$Sr at one’s site of residence prior to death [170,185–189].

Acknowledging the possibility of recent immigrants among the tested individuals, samples from archaeological animal skeletal tissues offer a more reliable measure of the local bioavailable $^{87}Sr/^{86}Sr$ compared to bone signatures. They provide an average of the bioavailable $^{87}Sr/^{86}Sr$ signatures of the feeding territories that these animals occupied and are thereby widely accepted as an accurate measure for the local $^{87}Sr/^{86}Sr$ value range in soils, plants, animals, and waters [165,173]. In principle, if an individual was born and raised in the local area, the $^{87}Sr/^{86}Sr$ values measured from his/her tooth enamel should be similar to his/her bone $^{87}Sr/^{86}Sr$ and also to the local bioavailable $^{87}Sr/^{86}Sr$ signatures. They will also be in overall agreement with comparable data from local geological material(s) and archaeological animal skeletal tissues. Otherwise, if human tooth enamel $^{87}Sr/^{86}Sr$ signatures are found to be statistically significantly different from the local $^{87}Sr/^{86}Sr$, we may infer that the respective people spent their childhood at (a) location(s) that are geologically and isotopically different from their residence prior to death. For the reasons outlined above, in this paper in addition to information on the local geology we discuss $^{87}Sr/^{86}Sr$ data from archaeological animals and human $^{87}Sr/^{86}Sr$ tooth enamel signatures from Corinth and the region of Carinthia in order to track potential residential mobility using the $^{87}Sr/^{86}Sr$ methodology.

#### 4.2 Geological context of the study area

Epirus comprises most of the mainland of north-west Greece and largely falls within the Ionian and Gavrovo isotopic/tectonic zones, while its basement comprises nappes that represent rocks of several different environments, stacked up on top of each other during the Alpine compression [190]. The Gavrovo zone was a continental fragment for the early part of its history, where the Mesozoic shallow-water limestones were later almost completely covered by late Eocene flysch sediments [190]. In Epirus, the Gavrovo zone crops out in a narrow belt west of the Pindos zone, and its oldest rocks are limestone [190]. Further west, the Ionian zone, a deep-water trough, crops out throughout much of the western part of this region and largely consists of deep-water limestones. The site of Amvrakia in particular, is set on hard limestone, but there also exist outcrops of flysch, as well as alluvium and recent deposits in the immediate proximity to the site, at a distance below 5 km (**Supplementary Figure S43**).

The archaeological site of Amvrakia on the north-east coast of the Gulf of Arta is set on a narrow strip of limestone/s [190]. Less than 2 km further west and south of the site extend alluvial deposits, while for a small part to its east and further south these rocks are interrupted by outcrops of flysch. The site of Amvrakia and its immediate periphery are thus characterized by high geological variability.

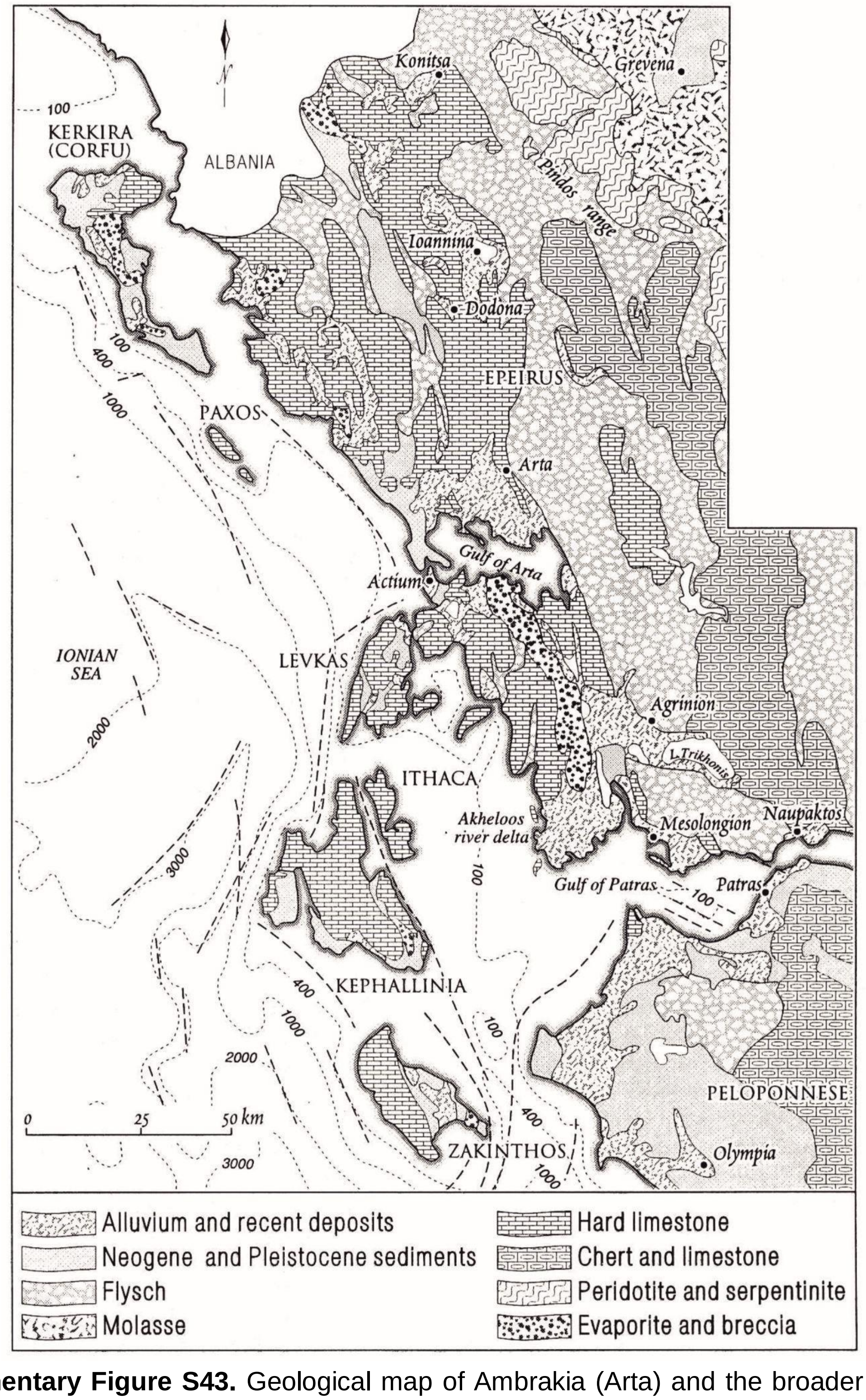


**Supplementary Figure S43.** Geological map of Ambrakia (Arta) and the broader region of Epirus. After [190].

### 4.3 Materials and Methods

#### 4.3.1 Samples

Strontium isotope ratios were measured from tooth enamel samples from 14 human burials of Amvrakia. With the exception of one burial, for which it was not possible to assign a relative date, eight of them were dated to the Archaic and the remaining six to the Classical period, respectively. Three of these burials were also analyzed for the corresponding $^{87}Sr/^{86}Sr$ signatures in tooth dentine samples. All human teeth sampled had previously been studied macroscopically. All human teeth were found attached to the associated maxillary/mandibular bone. There were seven M1s, five M2s, as well as one incisor and one canine. Relevant information is included in **Additional Table A2**.

#### 4.3.2 Sample preparation and analysis

**Strontium isotope ratio analysis**

The analytical protocol for $^{87}Sr/^{86}Sr$ analysis, including sample extraction and preparation prior to analysis, have been detailed in earlier publications [169,170]. Tooth enamel samples (> 20mg) were placed in an ultrasonic bath for a total of 30 minutes to remove surface contamination. The bath was interrupted every 10 minutes and the specimens were mechanically cleaned using distilled water. In order to remove diagenetic strontium, tooth enamel samples were leached in 2 ml of 5% acetic acid. All samples were rinsed in ultrapure water (four times) after the first hour of bathing in acetic acid and then placed back in fresh 5% and 2% acetic acid, respectively, and left overnight.

On the following day, samples were rinsed four times and dried in an oven (≤50 °C). All leachates were retained. Strontium columns were prepared by filling small Teflon columns up to the neck with cleaned Sr resin. The columns were cleaned with 3 ml $H_2O$ and 3 ml of SB 3M $HNO_3$. The matrix and everything except Sr and Rb was eluted with 2.5 ml of SB 3M $HNO_3$. Sr was collected by passing ultrapure water and dried down on a hotplate. The Sr fractions were loaded onto single tantalum filaments with Ta-activator and the $^{87}Sr/^{86}Sr$ values were measured to the sixth decimal digit with a ThermoFisher TRITON Plus Thermal Ionization Mass Spectrometer (7 Collectors). Preparation of the samples was carried out at the Department of Biology, University of Crete, while sample chemical analysis and measurement of the associated signatures were performed at the National Oceanography Centre in Southampton (NOCS).

### 4.4 Results and Discussion

Strontium isotope ratio ($^{87}Sr/^{86}Sr$) signatures from human tooth enamel from the 14 burials analyzed follow a broad distribution and range between 0.70808 and 0.70890. The results are largely consistent with consumption of regional livestock and agricultural products from the area of influence of the city of Amvrakia. In four of the examined cases, however, the human tooth enamel $^{87}Sr/^{86}Sr$ signatures (0.70859 to 0.70890) are similar to comparable data from human burials in Ancient Corinth (0.70848 to 0.70882) [191] and also to bioavailable data from the Corinthia region (0.70865 to 0.70869) [168,169], albeit there can be an overlap in Sr signatures between sites of similar geology. These data are thereby compatible with an origin from Corinth for the respective individuals and add support to the archaeological theory of the Corinthian colonization of ancient Amvrakia.

**Additional Table A2.** Strontium isotope ratio ($^{87}Sr/^{86}Sr$) signatures from human tooth enamel from the 14 burials analyzed in the present study and related burial metadata.

| isotope ID | Context | Element and tissue | Date |
|---|---|---|---|
| AN14 | LXXXVI B, AMV 30, ID164 | Tooth enamel, molar 1, upper right | Classical |
| AN187 | CCCLX, ID142 | Tooth enamel, molar 2, lower right | Archaic |
| AN15 | CCLXI, ID210 | Tooth enamel, molar 1, lower left | Archaic |
| AN13 | CCLXIV, ID102 | Tooth enamel, molar 1, lower right | Classical |
| AN186 | CXI, ID26 | Tooth enamel, molar 2, lower | Classical |
| AN11 | CXXVI 2, ID51 | Tooth enamel, molar 1, upper left | Archaic |
| AN189 | CXXVI, ID1 | Tooth enamel, molar 1, lower right | Classical |
| AN178 | T9, CCCLXXXVIII, ID13 | Tooth enamel, canine | Archaic |
| AN185 | T9, CCCXXXIX Burial 2, ID11 | Tooth enamel, molar 2, upper | Archaic |
| AN183 | T9, CCL Burial 2, ID171 | Tooth enamel, molar 2, upper | Classical |
| AN184 | T9, CCXCV, ID24 | Tooth enamel, incisor 1 | Archaic |
| AN182 | T9, CXXXIV | Tooth enamel, molar 2, upper | Classical |
| AN180 | T9, C | Tooth enamel, molar 1, upper | Archaic |
| AN181 | T9, CVII | Tooth enamel, molar 1, uower | Archaic |

### 5. Supplementary Information References


1. Dillon M, Dillion M, Garland L. Ancient Greece: Social and Historical Documents from Archaic Times to the Death of Alexander the Great. Routledge; 2010. Available from: https://play.google.com/store/books/details?id=ohYWBAAAQBAJ

2. Hansen MH, Nielsen TH. An Inventory of Archaic and Classical Poleis. OUP Oxford; 2004. Available from: https://play.google.com/store/books/details?id=9QZREAAAQBAJ

3. Graham AJ. Colony and Mother City in Ancient Greece. Manchester University Press; 1999. Available from: https://books.google.com/books/about/Colony_and_Mother_City_in_Ancient_Greece.html?hl=&id=z6XnAAAAIAAJ

4. Ridgway D. The First Western Greeks. CUP Archive; 1992. Available from: https://books.google.com/books/about/The_First_Western_Greeks.html?hl=&id=9F44AAAAIAAJ

5. Grammenos DV, Petropoulos EK. Ancient Greek Colonies in the Black Sea. Thessaloniki: Archaeological Institute of Northern Greece; 2003. Available from: https://books.google.com/books/about/Ancient_Greek_Colonies_in_the_Black_Sea.html?hl=&id=UtdoAAAAMAAJ

6. Tsetskhladze GR. Greek Colonisation: An Account of Greek Colonies and Other Settlements Overseas. Leiden, Boston, and Köln: Brill; 2008. Available from: https://books.google.com/books/about/Greek_Colonisation.html?hl=&id=PIgTAQAAIAAJ

7. Petropoulos EK. Problems in the history and archaeology of the Greek colonization of the Black Sea. In: Grammenos DV, Petropoulos EK, editors. Ancient Greek Colonies in the Black Sea. 2003 [cited 2024 Jul 18]. p. 17–92. Available from: https://www.academia.edu/32112108/ANCIENT_GREEK_COLONIES_IN_THE_BLACK_SEA_2_Grammenos_D_V_and_E_K_Petropoulos_eds_British_Archaeological_Reports_International_Series_1679_Oxford_2007

8. Petropoulos EK. Hellenic Colonization in Euxeinos Pontos: Penetration, Early Establishment, and the Problem of the “emporion” Revisited. British Archaeological Reports Oxford Limited; 2005. Available from: https://books.google.com/books/about/Hellenic_Colonization_in_Euxeinos_Pontos.html?hl=&id=0jBmAAAAMAAJ

9. van Dommelen P. Colonialism and Migration in the Ancient Mediterranean. Annu Rev Anthropol. 2012 [cited 2024 Apr 9];41:393–409. Available from: https://www.annualreviews.org/content/journals/10.1146/annurev-anthro-081309-145758

10. Malkin I. A Small Greek World: Networks in the Ancient Mediterranean. OUP USA; 2011. Available from: https://books.google.com/books/about/A_Small_Greek_World.html?hl=&id=CKQXm8sNgNkC

11. Osborne R. Greece in the Making 1200-479 BC. Routledge; 2009. Available from: https://books.google.com/books/about/Greece_in_the_Making_1200_479_BC.html?hl=&id=6WO7-Goh28IC

12. Malkin I. Foundations. A Companion to Archaic Greece. John Wiley & Sons, Ltd; 2009 [cited 2024 Jul 18]. p. 373–94. Available from:

https://onlinelibrary.wiley.com/doi/abs/10.1002/9781444308761.ch19

13. Hornblower S. Thucydides and “Chalkidic” Torone (IV.110.1). Oxford Journal of Archaeology. 1997 [cited 2024 Jul 18];16:177–86. Available from: https://onlinelibrary.wiley.com/doi/abs/10.1111/1468-0092.00033

14. Papadopoulos JK. Archaeology, Myth-History and the Tyranny of the Text: Chaldike, Torone and Thucydides. Oxford Journal of Archaeology. 1999 [cited 2024 Jul 18];18:377–94. Available from: https://onlinelibrary.wiley.com/doi/abs/10.1111/1468-0092.00091

15. Graham AJ. COMMERCIAL INTERCHANGES BETWEEN GREEKS AND NATIVES. Collected Papers on Greek Colonization. Brill; 2001 [cited 2024 Apr 17]. p. 45–56. Available from: https://brill.com/display/book/9789004351066/B9789004351066-s004.xml

16. Boardman J. The Greeks Overseas: Their Early Colonies and Trade. Thames and Hudson; 1999. Available from: https://play.google.com/store/books/details?id=EqHAQgAACAAJ

17. Reitsema LJ, Kyle B, Vassallo S. Food traditions and colonial interactions in the ancient Mediterranean: Stable isotope evidence from the Greek Sicilian colony Himera. Journal of Anthropological Archaeology. 2020;57:101144. Available from: https://www.sciencedirect.com/science/article/pii/S0278416519301734

18. Kaponis A. (In Greek) The Corinthian colonies around the Amvrakiko gulf from their foundation to the time of Philip II. 2020 [cited 2024 Jul 18]; Available from: https://pergamos.lib.uoa.gr/uoa/dl/object/2897438

19. Keenleyside A. Dental pathology and diet at Apollonia, a Greek colony on the Black Sea. International Journal of Osteoarchaeology. 2008 [cited 2024 Jul 18];18:262–79. Available from: https://onlinelibrary.wiley.com/doi/abs/10.1002/oa.934

20. Hammond N. The classical age of Greece. Harper & Row Publishers, Inc. USA; 1975. Available from: https://cir.nii.ac.jp/crid/1130000797935601664

21. Graham AJ. Patterns in Early Greek Colonisation. J Hell Stud. 1971 [cited 2024 Apr 19];91:35–47. Available from: https://www.cambridge.org/core/journals/journal-of-hellenic-studies/article/patterns-in-early-greek-colonisation/C5ECC7999BD671BDB82FC553D4DE3AB6

22. Strabo. Geography, Volume III: Books 6-7. Translated by Horace Leonard Jones. Loeb Classical Library 182. Cambridge. Harvard University Press; 1924. Available from: https://play.google.com/store/books/details?id=n4RiAAAAMAAJ

23. Wilkes J. GREEKS AND ILLYRIANS IN THE GREEK-LANGUAGE INSCRIPTIONS FROM EPIDAMNUS-DYRRHACIUM AND FROM APOLLONIA IN ILYRIA-PROCEEDINGS OF THE INTERNATIONAL ROUNDTABLE (CLERMONT-FERRAND, OCTOBER 19-21, 1989)-FRENCH-CABANES, P. SOC PROMOTION HELLENIC STUD 31-34 GORDON SQ, LONDON, UNITED KINGDOM WC1H OPP; 1995.

24. White ME. Greek Colonization. J Econ Hist. 1961;21:443–54. Available from: http://www.jstor.org/stable/2114410

25. Zhestokanov SM. The Corinth-Corcyra conflict of the seventh century BC. Saalburg Jahrb. 2020;26:15–23. Available from: http://saa.uaic.ro/the-corinth-corcyra-conflict-of-the-seventh-century-bc/

26. Papadopoulou V. RES GESTAE. The work of the Ephorate of Antiquities of Arta during

the years 2014 – 2020, Arta 2020 / RES GESTAE. 2020 [cited 2024 Jul 18]; Available from: https://www.academia.edu/51173824/RES_GESTAE_The_work_of_the_Ephorate_of_Antiquities_of_Arta_during_the_years_2014_2020_Arta_2020_RES_GESTAE_%CE%A4%CE%B1_%CE%A0%CE%B5%CF%80%CF%81%CE%B1%CE%B3%CE%BC%CE%AD%CE%BD%CE%B1_%CF%84%CE%B7%CF%82_%CE%95%CF%86%CE%BF%CF%81%CE%B5%CE%AF%CE%B1%CF%82_%CE%91%CF%81%CF%87%CE%B1%CE%B9%CE%BF%CF%84%CE%AE%CF%84%CF%89%CE%BD_%CE%86%CF%81%CF%84%CE%B1%CF%82_%CE%BA%CE%B1%CF%84%CE%AC_%CF%84%CE%B1_%CE%AD%CF%84%CE%B7_2014_2020_%CE%86%CF%81%CF%84%CE%B1_2020

27. Aggeli A. (In Greek) The burial precincts of Amvrakia, in: K. Sporn (ed.), Griechische Grabbezirke klassischer Zeit, Athenaia 6, 2013. 2013 [cited 2024 Jul 18]; Available from: https://www.academia.edu/19964480/%CE%9F%CE%B9_%CF%84%CE%B1%CF%86%CE%B9%CE%BA%CE%BF%CE%AF_%CF%80%CE%B5%CF%81%CE%AF%CE%B2%CE%BF%CE%BB%CE%BF%CE%B9_%CF%84%CE%B7%CF%82_%CE%91%CE%BC%CE%B2%CF%81%CE%B1%CE%BA%CE%AF%CE%B1%CF%82_in_K_Sporn_ed_Griechische_Grabbezirke_klassischer_Zeit_Athenaia_6_2013

28. Tzouvara-Souli H. (In Greek) Amvrakia. Artis Musicological Association "O Skoufas"; 1992 [cited 2024 Jul 18]. Available from: https://books.google.com/books/about/%CE%91%CE%BC%CE%B2%CF%81%CE%B1%CE%BA%CE%AF%CE%B1.html?hl=el&id=bJxMAAAAYAAJ

29. Exemplare: Ambracie, une ville ancienne se reconstitue peu à peu par les recherches. [cited 2024 Aug 26]. Available from: https://zenon.dainst.org/Record/001019928

30. Robinson EW. The First Democracies: Early Popular Government Outside Athens. Stuttgart: F. Steiner; 1997 [cited 2024 Jul 18]. p. 80–2. Available from: https://books.google.com/books/about/The_First_Democracies.html?hl=el&id=T1kfcobFRSMC

31. Chrysos EK. Nicopolis I: Proceedings of the First International Symposium on Nicopolis (23-29 September 1984). Actia Nicopolis Foundation; 1987 [cited 2024 Jul 18]. Available from: https://books.google.com/books/about/Nicopolis_I.html?hl=el&id=-hHVmQEACAAJ

32. Kordosis M. Ancient and Early Byzantine Tenea. Dodone. 1997 [cited 2024 Jul 18];26:465–580. Available from: http://dx.doi.org/https://olympias.lib.uoi.gr/jspui/handle/123456789/6127 http://dx.doi.org/10.26268/heal.uoi.9267

33. Wiseman J. Corinth and Rome I: 228 B.C.—A.D. 267. Band 7/1 Halbband Politische Geschichte (Provinzen und Randvölker: Griechischer Balkanraum; Kleinasien). De Gruyter; 2016 [cited 2024 Aug 28]. p. 438–548. Available from: https://www.degruyter.com/document/doi/10.1515/9783110837612-009/html

34. Tenea Project, Elena K, Lagos C. Korka, E. & Lagos, C. (2019), Numismatic Grave Finds of the Tenea-Chiliomodi Excavation Project 2013-2017. The Journal of Archaeological Numismatics,Volume 9, pp 349-362. 2019 [cited 2024 Aug 1]; Available from: https://www.academia.edu/42952562/Korka_E_and_Lagos_C_2019_Numismatic_Grave_Finds_of_the_Tenea_Chiliomodi_Excavation_Project_2013_2017

35. Korka E, Evaggeloglou P, Panailidis P, Christidis I. Systematic excavation of Ancient Tenea: A journey from the past to the present with an eye to the future. The partnership between local community and archaeology. Mare Ponticum. 2022 [cited 2024 Jul 18];10:84. Available from: http://mareponticum.bscc.duth.gr/index_htm_files/korka-evangelopoulou-panailidis-christidis_10.pdf

36. Korka E, Evaggeloglou P. A cemetery excavation unearths Tenea’s past. pp. 101-113. Griechische Nekropolen Bibliopolis / Möhnesee. 2019 [cited 2024 Jul 8]; Available from: https://www.academia.edu/43805124/Korka_E_and_Evaggeloglou_P_2019_A_cemetery_excavation_unearths_Teneas_past_pp_101_113

37. Papadopoulou B, Angeli A. (In Greek) Amvrakia. The city and its monuments. Arta: Ephorate of Antiquities of Arta; 2015 [cited 2024 Jul 18]. Available from: https://www.kardamitsa.gr/product/47869/ambrakia-i-poli-kai-ta-mnimeia-tis.html

38. Oikonomidis S, Papayiannis A, Tsonos A. The Emergence and the Architectural Development of the Tumulus Burial Custom in NW Greece (Epirus and the Ionian Islands) and Albania and its Connections to Settlement Organization. 2012 [cited 2024 Apr 23]; Available from: https://www.semanticscholar.org/paper/28eac8e9ce3e37f3d2f3fe70021225fb785cf1b3

39. Oikonomidis S, Papayiannis A, Tsonos A. (In Greek) The burial custom of tumulus along the Ionian and Adriatic Coast as a cultural and social phenomenon. AURA. 2018 [cited 2024 Jul 18]; Available from: https://www.academia.edu/36769273/S_Oikonomidis_A_Papayiannis_and_A_Tsonos_%CE%A4%CE%BF_%CF%84%CE%B1%CF%86%CE%B9%CE%BA%CF%8C_%CE%AD%CE%B8%CE%B9%CE%BC%CE%BF_%CF%84%CE%B7%CF%82_%CE%B1%CE%BD%CE%AD%CE%B3%CE%B5%CF%81%CF%83%CE%B7%CF%82_%CF%84%CF%8D%CE%BC%CE%B2%CE%BF%CF%85_%CE%BA%CE%B1%CF%84%CE%AC_%CE%BC%CE%AE%CE%BA%CE%BF%CF%82_%CF%84%CE%B7%CF%82_%CE%99%CE%BF%CE%BD%CE%AF%CE%B1%CF%82_%CE%BA%CE%B1%CE%B9_%CE%86%CE%B4%CF%81%CE%B9%CE%B1%CF%84%CE%B9%CE%BA%CE%AE%CF%82_%CE%B1%CE%BA%CF%84%CE%AE%CF%82_%CF%89%CF%82_%CF%80%CE%BF%CE%BB%CE%B9%CF%84%CE%B9%CF%83%CF%84%CE%B9%CE%BA%CF%8C_%CE%BA%CE%B1%CE%B9_%CE%BA%CE%BF%CE%B9%CE%BD%CF%89%CE%BD%CE%B9%CE%BA%CF%8C_%CF%86%CE%B1%CE%B9%CE%BD%CF%8C%CE%BC%CE%B5%CE%BD%CE%BF_The_burial_custom_of_tumulus_along_the_Ionian_and_Adriatic_Coast_as_a_cultural_and_social_phenomenon_

40. Angeli A. (In Greek) The cemeteries of Amvrakia during the Archaic and Classical times. National Documentation Centre (EKT); 2021. Available from: https://www.didaktorika.gr/eadd/handle/10442/49590?locale=en

41. Zafiropoulos NS. (In Greek) Excavation of Amvrakia tombs under Nikolaos S. Zafiropoulou. 1962 [cited 2024 Aug 6]; Available from: http://hdl.handle.net/11615/11552

42. Kirkoy T. (In Greek) Hellenistic pottery from the eastern cemetery of ancient Amvrakia (Merkovitis and Kokkinelli plot), 8th Scientific Meeting on Hellenistic Ceramics, Ioannina 5-9 May 2009, Athens 2014. 2009 [cited 2024 Jul 18]; Available from: https://www.academia.edu/12089457/%CE%95%CE%BB%CE%BB%CE%B7%CE%BD%CE%B9%CF%83%CF%84%CE%B9%CE%BA%CE%AE_%CE%BA%CE%B5%CF%81%CE%B1%CE%BC%CE%B9%CE%BA%CE%AE_%CE%B1%CF%80%CF%8C_%CF%84%CE%BF_%CE%B1%CE%BD%CE%B1%CF%84%CE%BF%CE%BB%CE%B9%CE%BA%CF%8C_%CE%BD%CE%B5%CE%BA%CF%81%CE%BF%CF%84%CE%B1%CF%86%CE%B5%CE%AF%CE%BF_%CF%84%CE%B7%CF%82_%CE%B1%CF%81%CF%87%CE%B1%CE%AF%CE%B1%CF%82_%CE%91%CE%BC%CE%B2%CF%81%CE%B1%CE%BA%CE%AF%CE%B1%CF%82_%CE%BF%CE%B9%CE%BA%CF%8C%CF%80%CE%B5%CE%B4%CE%BF_%CE%9C%CE%B5%CF%81%CE%BA%CE%BF%CE%B2%CE%AF%CF%84%CE%B7_%CE%BA%CE%B1%CE%B9_%CE%9A%CE%BF%CE%BA%CE%BA%CE%B9%CE%BD%CE%AD%CE%BB%CE%BB%CE%B7_%CE%97_%CE%95%CF%80%CE%B9%CF%83%CF%84%CE%B7%CE%BC%CE%BF%CE%BD%CE%B9%CE%BA%CE%AE_%CE%A3%CF%85%CE%BD%CE%AC%CE%BD%CF%84%CE%B7%CF%83%CE

%B7_%CE%B3%CE%B9%CE%B1_%CF%84%CE%B7%CE%BD_%CE%95%CE%BB%CE%BB%CE%B7%CE%BD%CE%B9%CF%83%CF%84%CE%B9%CE%BA%CE%AE_%CE%9A%CE%B5%CF%81%CE%B1%CE%BC%CE%B9%CE%BA%CE%AE_%CE%99%CF%89%CE%AC%CE%BD%CE%BD%CE%B9%CE%BD%CE%B1_5_9_%CE%9C%CE%B1%E1%BF%92%CE%BF%CF%85_2009_%CE%91%CE%B8%CE%AE%CE%BD%CE%B1_2014

43. Katsadima I. (In Greek) Epitaphic columns from Ambrakia. University of Ioannina. Faculty of Philosophy. Department of History and Archaeology; 2003 [cited 2024 Jul 18]. Available from: http://hdl.handle.net/10442/hedi/17630

44. Vasileiou E. Neolithic and Bronze Age Epirus revisited, Archaeology in Greece 2019–2020 (2019-2020). Archaeological Reports. 2020;66:67–81. Available from: https://www.jstor.org/stable/27098049

45. Baladima A., Raptopoulos S., Kptsokostas I.. (In Greek) Chronicles A.D. 71, 2016 (Arta). Chronicles. 2016 [cited 2024 Jul 18];71. Available from: https://www.academia.edu/95794876/%CE%91%CE%9D%CE%94%CE%A1%CE%9F%CE%9C%CE%91%CE%A7%CE%97_%CE%9C%CE%A0%CE%91%CE%9B%CE%91%CE%94%CE%97%CE%9C%CE%91_%CE%A3%CE%A9%CE%A4%CE%97%CE%A1%CE%99%CE%9F%CE%A3_%CE%A1%CE%91%CE%A0%CE%A4%CE%9F%CE%A0%CE%9F%CE%A5%CE%9B%CE%9F%CE%A3_%CE%99%CE%A9%CE%91%CE%9D%CE%9D%CE%97%CE%A3_%CE%9A%CE%A9%CE%A4%CE%A3%CE%9F%CE%9A%CE%A9%CE%A3%CE%A4%CE%91%CE%A3_%CE%A7%CF%81%CE%BF%CE%BD%CE%B9%CE%BA%CE%AC_%CE%91_%CE%94_71_2016_%CE%86%CF%81%CF%84%CE%B1_

46. Vokotopoulou I. (In Greek) Vitsa: the cemeteries of a Molossian count. Archaeological Resources and Expropriations Fund; 1986 [cited 2024 Jul 18]. Available from: https://books.google.com/books/about/%CE%92%CE%AF%CF%84%CF%83%CE%B1.html?hl=el&id=x0_6zQEACAAJ

47. Korka E, Agelarakis A. (In Greek) New evidence on the early written sarcophagus of Faneromenes Chiliomodios. The archaeological project in peloponnese (AEPEL1), Proceedings of the Tripoli International Conference (November 7-11, 2012). 2018;627–34. Available from: https://www.researchgate.net/publication/329759926_Nea_stoicheia_peri_tes_porines_graptes_sarkophagou_Phaneromenes_Chiliomodiou_Korka_E_kai_Agelarakes_A_TO_ARCHAIOLOGIKO_ERGO_STEN_PELOPONNESO_AEPEL1_Praktika_tou_Diethnous_Synedriou_Tripole_7-11_Noembr/citation/download

48. Korka E. (In Greek) The written early sarcophagus of Faneromenis Chiliomodius of Corinth. The Corinthia and the Northeast Peloponnese Topography and History from Prehistoric Times until the end of Antiquity. 2013 [cited 2024 Aug 1]. p. 305–11. Available from: https://www.aegeussociety.org/new_book/the-corinthia-and-the-northeast-peloponnese-topography-and-history-from-prehistoric-times-until-the-end-of-antiquity/

49. Tenea Project, Evaggeloglou P (vivi), Elena K. Korka, E. & Evaggeloglou, P. (2020) (In Greek) Ancient Tenea. Systematic archaeological research in Chiliomodi, Corinth, 2013-2017. AEPEL(2). 2020 [cited 2024 Aug 1]; Available from: https://www.academia.edu/45328998/%CE%9A%CF%8C%CF%81%CE%BA%CE%B1_%CE%95_and_%CE%95%CF%85%CE%B1%CE%B3%CE%B3%CE%AD%CE%BB%CE%BF%CE%B3%CE%BB%CE%BF%CF%85_%CE%A0_2020_%CE%91%CF%81%CF%87%CE%B1%CE%AF%CE%B1_%CE%A4%CE%B5%CE%BD%CE%AD%CE%B1_%CE%A3%CF%85%CF%83%CF%84%CE%B7%CE%BC%CE%B1%CF%84%CE%B9%CE%BA%CE%AE_%CE%B1%CF%81%CF%87%CE%B1%CE%B9%CE%BF%CE%BB%CE%BF%CE%B3%CE%B9%CE%BA%CE%AE_%CE%AD%CF%81%CE%B5%CF%85%CE%BD%CE%B1_

%CF%83%CF%84%CE%BF_%CE%A7%CE%B9%CE%BB%CE%B9%CE%BF%CE%BC%CF%8C%CE%B4%CE%B9_%CE%9A%CE%BF%CF%81%CE%B9%CE%BD%CE%B8%CE%AF%CE%B1%CF%82_2013_2017

50. Harney É, Cheronet O, Fernandes DM, Sirak K, Mah M, Bernardos R, et al. A minimally destructive protocol for DNA extraction from ancient teeth. Genome Res. 2021;31:472–83. Available from: http://dx.doi.org/10.1101/gr.267534.120

51. Rohland N, Glocke I, Aximu-Petri A, Meyer M. Extraction of highly degraded DNA from ancient bones, teeth and sediments for high-throughput sequencing. Nat Protoc. 2018;13:2447–61. Available from: http://dx.doi.org/10.1038/s41596-018-0050-5

52. Orfanou E, Himmel M, Aron F, Haak W. Minimally-invasive sampling of pars petrosa (os temporale) for ancient DNA extraction. 2020 [cited 2023 Nov 27]; Available from: https://www.protocols.io/view/minimally-invasive-sampling-of-pars-petrosa-os-tem-bqd8ms9w.pdf

53. Allentoft ME, Sikora M, Sjögren K-G, Rasmussen S, Rasmussen M, Stenderup J, et al. Population genomics of Bronze Age Eurasia. Nature. 2015;522:167–72. Available from: http://dx.doi.org/10.1038/nature14507

54. Meyer M, Kircher M. Illumina sequencing library preparation for highly multiplexed target capture and sequencing. Cold Spring Harb Protoc. 2010;2010:db.prot5448. Available from: http://dx.doi.org/10.1101/pdb.prot5448

55. Dabney J, Knapp M, Glocke I, Gansauge M-T, Weihmann A, Nickel B, et al. Complete mitochondrial genome sequence of a Middle Pleistocene cave bear reconstructed from ultrashort DNA fragments. Proc Natl Acad Sci U S A. 2013;110:15758–63. Available from: http://dx.doi.org/10.1073/pnas.1314445110

56. Rohland N, Harney E, Mallick S, Nordenfelt S, Reich D. Partial uracil–DNA–glycosylase treatment for screening of ancient DNA. Philos Trans R Soc Lond B Biol Sci. 2015 [cited 2023 Nov 27];370. Available from: https://www.ncbi.nlm.nih.gov/pmc/articles/PMC4275898/

57. Neuenschwander S, Cruz Dávalos DI, Anchieri L, Sousa da Mota B, Bozzi D, Rubinacci S, et al. Mapache: a flexible pipeline to map ancient DNA. Bioinformatics. 2023 [cited 2023 May 31];39:btad028. Available from: https://academic.oup.com/bioinformatics/article-pdf/39/2/btad028/50436112/btad028.pdf

58. Mölder F, Jablonski KP, Letcher B, Hall MB, Tomkins-Tinch CH, Sochat V, et al. Sustainable data analysis with Snakemake. F1000Res. 2021;10:33. Available from: http://dx.doi.org/10.12688/f1000research.29032.2

59. Schubert M, Lindgreen S, Orlando L. AdapterRemoval v2: rapid adapter trimming, identification, and read merging. BMC Res Notes. 2016;9:88. Available from: http://dx.doi.org/10.1186/s13104-016-1900-2

60. Prüfer K, Stenzel U, Hofreiter M, Pääbo S, Kelso J, Green RE. Computational challenges in the analysis of ancient DNA. Genome Biol. 2010 [cited 2023 Jul 5];11:R47. Available from: https://www.ncbi.nlm.nih.gov/pmc/articles/PMC2898072/

61. A global reference for human genetic variation. Nature. 2015 [cited 2023 Jul 12];526:68–74. Available from: https://www.nature.com/articles/nature15393

62. Li H, Durbin R. Fast and accurate short read alignment with Burrows–Wheeler transform. Bioinformatics. 2009 [cited 2023 May 31];25:1754–60. Available from:

https://academic.oup.com/bioinformatics/article-pdf/25/14/1754/48994219/bioinformatics_25_14_1754.pdf

63. Schubert M, Ginolhac A, Lindgreen S, Thompson JF, Al-Rasheid KA, Willerslev E, et al. Improving ancient DNA read mapping against modern reference genomes. BMC Genomics. 2012 [cited 2023 May 31];13. Available from: https://pubmed.ncbi.nlm.nih.gov/22574660/

64. Oliva A, Tobler R, Cooper A, Llamas B, Souilmi Y. Systematic benchmark of ancient DNA read mapping. Brief Bioinform. 2021 [cited 2023 May 31];22:bbab076. Available from: https://academic.oup.com/bib/article-pdf/22/5/bbab076/40260467/bbab076.pdf

65. Li H, Handsaker B, Wysoker A, Fennell T, Ruan J, Homer N, et al. The Sequence Alignment/Map format and SAMtools. Bioinformatics. 2009 [cited 2023 May 31];25:2078–9. Available from: https://academic.oup.com/bioinformatics/article-pdf/25/16/2078/48994296/bioinformatics_25_16_2078.pdf

66. Jun G, Wing MK, Abecasis GR, Kang HM. An efficient and scalable analysis framework for variant extraction and refinement from population-scale DNA sequence data. Genome Res. 2015;25:918–25. Available from: http://dx.doi.org/10.1101/gr.176552.114

67. Malaspinas A-S, Tange O, Moreno-Mayar JV, Rasmussen M, DeGiorgio M, Wang Y, et al. bammds: a tool for assessing the ancestry of low-depth whole-genome data using multidimensional scaling (MDS). Bioinformatics. 2014 [cited 2023 Jun 1];30:2962–4. Available from: https://academic.oup.com/bioinformatics/article-pdf/30/20/2962/48929902/bioinformatics_30_20_2962.pdf

68. McKenna A, Hanna M, Banks E, Sivachenko A, Cibulskis K, Kernytsky A, et al. The Genome Analysis Toolkit: a MapReduce framework for analyzing next-generation DNA sequencing data. Genome Res. 2010;20:1297–303. Available from: http://dx.doi.org/10.1101/gr.107524.110

69. Okonechnikov K, Conesa A, García-Alcalde F. Qualimap 2: advanced multi-sample quality control for high-throughput sequencing data. Bioinformatics. 2016;32:292–4. Available from: http://dx.doi.org/10.1093/bioinformatics/btv566

70. Mittnik A, Wang C-C, Svoboda J, Krause J. A Molecular Approach to the Sexing of the Triple Burial at the Upper Paleolithic Site of Dolní Věstonice. PLoS One. 2016 [cited 2023 Jun 23];11:e0163019. Available from: https://journals.plos.org/plosone/article/file?id=10.1371/journal.pone.0163019&type=printable

71. Skoglund P, Storå J, Götherström A, Jakobsson M. Accurate sex identification of ancient human remains using DNA shotgun sequencing. J Archaeol Sci. 2013 [cited 2023 Jun 23];40:4477–82. Available from: http://dx.doi.org/10.1016/j.jas.2013.07.004

72. Fu Q, Mittnik A, Johnson PLF, Bos K, Lari M, Bollongino R, et al. A revised timescale for human evolution based on ancient mitochondrial genomes. Curr Biol. 2013;23:553–9. Available from: http://dx.doi.org/10.1016/j.cub.2013.02.044

73. Green RE, Malaspinas A-S, Krause J, Briggs AW, Johnson PLF, Uhler C, et al. A complete Neandertal mitochondrial genome sequence determined by high-throughput sequencing. Cell. 2008;134:416–26. Available from: http://dx.doi.org/10.1016/j.cell.2008.06.021

74. Korneliussen TS, Albrechtsen A, Nielsen R. ANGSD: Analysis of Next Generation Sequencing Data. BMC Bioinformatics. 2014;15:356. Available from: http://dx.doi.org/10.1186/s12859-014-0356-4

75. Katoh K, Standley DM. MAFFT Multiple Sequence Alignment Software Version 7: Improvements in Performance and Usability. Mol Biol Evol. 2013 [cited 2023 Jun 1];30:772. Available from: https://www.ncbi.nlm.nih.gov/pmc/articles/PMC3603318/

76. Renaud G, Slon V, Duggan AT, Kelso J. Schmutzi: estimation of contamination and endogenous mitochondrial consensus calling for ancient DNA. Genome Biol. 2015;16:224. Available from: http://dx.doi.org/10.1186/s13059-015-0776-0

77. Schönherr S, Weissensteiner H, Kronenberg F, Forer L. Haplogrep 3 - an interactive haplogroup classification and analysis platform. Nucleic Acids Res. 2023;51:W263–8. Available from: http://dx.doi.org/10.1093/nar/gkad284

78. Rubin JD, Vogel NA, Gopalakrishnan S, Sackett PW, Renaud G. HaploCart: Human mtDNA haplogroup classification using a pangenomic reference graph. PLoS Comput Biol. 2023;19:e1011148. Available from: http://dx.doi.org/10.1371/journal.pcbi.1011148

79. Mallick S, Reich D. The Allen Ancient DNA Resource (AADR): A curated compendium of ancient human genomes. Harvard Dataverse; 2023 [cited 2024 Jul 29]. Available from: https://dataverse.harvard.edu/dataset.xhtml?persistentId=doi:10.7910/DVN/FFIDCW

80. Ralf A, Montiel González D, Zhong K, Kayser M. Yleaf: Software for Human Y-Chromosomal Haplogroup Inference from Next-Generation Sequencing Data. Mol Biol Evol. 2018 [cited 2023 Jun 22];35:1291–4. Available from: https://academic.oup.com/mbe/article-pdf/35/5/1291/24704771/msy032.pdf

81. David Poznik G. Identifying Y-chromosome haplogroups in arbitrarily large samples of sequenced or genotyped men. bioRxiv. 2016 [cited 2023 Jun 22]. p. 088716. Available from: https://www.biorxiv.org/content/10.1101/088716v1.abstract

82. Danecek P, Bonfield JK, Liddle J, Marshall J, Ohan V, Pollard MO, et al. Twelve years of SAMtools and BCFtools. Gigascience. 2021;10. Available from: http://dx.doi.org/10.1093/gigascience/giab008

83. Mallick S, Micco A, Mah M, Ringbauer H, Lazaridis I, Olalde I, et al. The Allen Ancient DNA Resource (AADR): A curated compendium of ancient human genomes. bioRxiv. 2023; Available from: http://dx.doi.org/10.1101/2023.04.06.535797

84. Koptekin D, Yüncü E, Rodríguez-Varela R, Altınışık NE, Psonis N, Kashuba N, et al. Spatial and temporal heterogeneity in human mobility patterns in Holocene Southwest Asia and the East Mediterranean. Curr Biol. 2023;33:41–57.e15. Available from: http://dx.doi.org/10.1016/j.cub.2022.11.034

85. Patterson N, Price AL, Reich D. Population Structure and Eigenanalysis. PLoS Genet. 2006 [cited 2023 Jun 23];2:e190. Available from: https://journals.plos.org/plosgenetics/article/file?id=10.1371/journal.pgen.0020190&type=printable

86. Kuhn JMM, Jakobsson M, Günther T. Estimating genetic kin relationships in prehistoric populations. PLoS One. 2018 [cited 2023 Jun 22];13:e0195491. Available from: https://journals.plos.org/plosone/article/file?id=10.1371/journal.pone.0195491&type=printable

87. Popli D, Peyrégne S, Peter BM. KIN: a method to infer relatedness from low-coverage ancient DNA. Genome Biol. 2023 [cited 2023 Jun 22];24:1–22. Available from: https://genomebiology.biomedcentral.com/articles/10.1186/s13059-023-02847-7

88. Psonis N, Vassou D, Nafplioti A, Tabakaki E, Pavlidis P, Stamatakis A, et al. Identification of the 18 World War II executed citizens of Adele, Rethymnon, Crete using an ancient DNA approach and low coverage genomes. Forensic Sci Int Genet. 2024;71:103060. Available from: http://dx.doi.org/10.1016/j.fsigen.2024.103060

89. Purcell S, Neale B, Todd-Brown K, Thomas L, Ferreira MAR, Bender D, et al. PLINK: a tool set for whole-genome association and population-based linkage analyses. Am J Hum Genet. 2007;81:559–75. Available from: http://dx.doi.org/10.1086/519795

90. Ringbauer H, Novembre J, Steinrücken M. Parental relatedness through time revealed by runs of homozygosity in ancient DNA. Nat Commun. 2021;12:5425. Available from: http://dx.doi.org/10.1038/s41467-021-25289-w

91. Rivollat M, Rohrlach AB, Ringbauer H, Childebayeva A, Mendisco F, Barquera R, et al. Extensive pedigrees reveal the social organization of a Neolithic community. Nature. 2023;620:600–6. Available from: http://dx.doi.org/10.1038/s41586-023-06350-8

92. Gretzinger J, Schmitt F, Mötsch A, Carlhoff S, Lamnidis TC, Huang Y, et al. Evidence for dynastic succession among early Celtic elites in Central Europe. Nat Hum Behav. 2024; Available from: http://dx.doi.org/10.1038/s41562-024-01888-7

93. Rubinacci S, Ribeiro DM, Hofmeister RJ, Delaneau O. Efficient phasing and imputation of low-coverage sequencing data using large reference panels. Nat Genet. 2021;53:120–6. Available from: http://dx.doi.org/10.1038/s41588-020-00756-0

94. Allentoft ME, Sikora M, Refoyo-Martínez A, Irving-Pease EK, Fischer A, Barrie W, et al. Population genomics of post-glacial western Eurasia. Nature. 2024;625:301–11. Available from: http://dx.doi.org/10.1038/s41586-023-06865-0

95. Link V, Kousathanas A, Veeramah K, Sell C, Scheu A, Wegmann D. ATLAS: Analysis Tools for Low-depth and Ancient Samples. bioRxiv. 2017 [cited 2023 May 20]. p. 105346. Available from: https://www.biorxiv.org/content/10.1101/105346v2.abstract

96. International HapMap Consortium, Frazer KA, Ballinger DG, Cox DR, Hinds DA, Stuve LL, et al. A second generation human haplotype map of over 3.1 million SNPs. Nature. 2007;449:851–61. Available from: http://dx.doi.org/10.1038/nature06258

97. Ringbauer H, Huang Y, Akbari A, Mallick S, Olalde I, Patterson N, et al. Accurate detection of identity-by-descent segments in human ancient DNA. Nat Genet. 2024;56:143–51. Available from: http://dx.doi.org/10.1038/s41588-023-01582-w

98. Agranat-Tamir L, Waldman S, Martin MAS, Gokhman D, Mishol N, Eshel T, et al. The Genomic History of the Bronze Age Southern Levant. Cell. 2020;181:1146–57.e11. Available from: http://dx.doi.org/10.1016/j.cell.2020.04.024

99. Fu Q, Posth C, Hajdinjak M, Petr M, Mallick S, Fernandes D, et al. The genetic history of Ice Age Europe. Nature. 2016;534:200–5. Available from: http://dx.doi.org/10.1038/nature17993

100. Clemente F, Unterländer M, Dolgova O, Amorim CEG, Coroado-Santos F, Neuenschwander S, et al. The genomic history of the Aegean palatial civilizations. Cell. 2021;184:2565–86.e21. Available from: http://dx.doi.org/10.1016/j.cell.2021.03.039

101. Freilich S, Ringbauer H, Los D, Novak M, Pavičić DT, Schiffels S, et al. Reconstructing genetic histories and social organisation in Neolithic and Bronze Age Croatia. Sci Rep. 2021;11:16729. Available from: http://dx.doi.org/10.1038/s41598-021-94932-9

102. Feldman M, Fernández-Domínguez E, Reynolds L, Baird D, Pearson J, Hershkovitz I, et al. Late Pleistocene human genome suggests a local origin for the first farmers of central Anatolia. Nat Commun. 2019;10:1218. Available from: http://dx.doi.org/10.1038/s41467-019-09209-7

103. De Angelis F, Romboni M, Veltre V, Catalano P, Martínez-Labarga C, Gazzaniga V, et al. First Glimpse into the Genomic Characterization of People from the Imperial Roman Community of Casal Bertone (Rome, First–Third Centuries AD). Genes. 2022 [cited 2024 Mar 22];13:136. Available from: https://www.mdpi.com/2073-4425/13/1/136

104. Fernandes DM, Mittnik A, Olalde I, Lazaridis I, Cheronet O, Rohland N, et al. The spread of steppe and Iranian-related ancestry in the islands of the western Mediterranean. Nat Ecol Evol. 2020;4:334–45. Available from: http://dx.doi.org/10.1038/s41559-020-1102-0

105. Hofmanová Z, Kreutzer S, Hellenthal G, Sell C, Diekmann Y, Díez-Del-Molino D, et al. Early farmers from across Europe directly descended from Neolithic Aegeans. Proc Natl Acad Sci U S A. 2016;113:6886–91. Available from: http://dx.doi.org/10.1073/pnas.1523951113

106. Jones ER, Gonzalez-Fortes G, Connell S, Siska V, Eriksson A, Martiniano R, et al. Upper Palaeolithic genomes reveal deep roots of modern Eurasians. Nat Commun. 2015;6:8912. Available from: http://dx.doi.org/10.1038/ncomms9912

107. Lazaridis I, Nadel D, Rollefson G, Merrett DC, Rohland N, Mallick S, et al. Genomic insights into the origin of farming in the ancient Near East. Nature. 2016;536:419–24. Available from: http://dx.doi.org/10.1038/nature19310

108. Lazaridis I, Mittnik A, Patterson N, Mallick S, Rohland N, Pfrengle S, et al. Genetic origins of the Minoans and Mycenaeans. Nature. 2017;548:214–8. Available from: http://dx.doi.org/10.1038/nature23310

109. Marchi N, Winkelbach L, Schulz I, Brami M, Hofmanová Z, Blöcher J, et al. The genomic origins of the world’s first farmers. Cell. 2022;185:1842–59.e18. Available from: http://dx.doi.org/10.1016/j.cell.2022.04.008

110. Marcus JH, Posth C, Ringbauer H, Lai L, Skeates R, Sidore C, et al. Genetic history from the Middle Neolithic to present on the Mediterranean island of Sardinia. Nat Commun. 2020;11:939. Available from: http://dx.doi.org/10.1038/s41467-020-14523-6

111. Mathieson I, Lazaridis I, Rohland N, Mallick S, Patterson N, Roodenberg SA, et al. Genome-wide patterns of selection in 230 ancient Eurasians. Nature. 2015;528:499–503. Available from: http://dx.doi.org/10.1038/nature16152

112. Mathieson I, Alpaslan-Roodenberg S, Posth C, Szécsényi-Nagy A, Rohland N, Mallick S, et al. The genomic history of southeastern Europe. Nature. 2018;555:197–203. Available from: http://dx.doi.org/10.1038/nature25778

113. Moots HM, Antonio M, Sawyer S, Spence JP, Oberreiter V, Weiß CL, et al. A genetic history of continuity and mobility in the Iron Age central Mediterranean. Nat Ecol Evol. 2023;7:1515–24. Available from: http://dx.doi.org/10.1038/s41559-023-02143-4

114. Narasimhan VM, Patterson N, Moorjani P, Rohland N, Bernardos R, Mallick S, et al. The formation of human populations in South and Central Asia. Science. 2019;365. Available from: http://dx.doi.org/10.1126/science.aat7487

115. Olalde I, Allentoft ME, Sánchez-Quinto F, Santpere G, Chiang CWK, DeGiorgio M, et al. Derived immune and ancestral pigmentation alleles in a 7,000-year-old Mesolithic European.

Nature. 2014;507:225–8. Available from: http://dx.doi.org/10.1038/nature12960

116. Olalde I, Brace S, Allentoft ME, Armit I, Kristiansen K, Booth T, et al. Erratum: The Beaker phenomenon and the genomic transformation of northwest Europe. Nature. 2018;555:543. Available from: http://dx.doi.org/10.1038/nature26164

117. Patterson N, Isakov M, Booth T, Büster L, Fischer C-E, Olalde I, et al. Large-scale migration into Britain during the Middle to Late Bronze Age. Nature. 2022;601:588–94. Available from: http://dx.doi.org/10.1038/s41586-021-04287-4

118. Posth C, Zaro V, Spyrou MA, Vai S, Gnecchi-Ruscone GA, Modi A, et al. The origin and legacy of the Etruscans through a 2000-year archeogenomic time transect. Sci Adv. 2021;7:eabi7673. Available from: http://dx.doi.org/10.1126/sciadv.abi7673

119. Raghavan M, Skoglund P, Graf KE, Metspalu M, Albrechtsen A, Moltke I, et al. Upper Palaeolithic Siberian genome reveals dual ancestry of Native Americans. Nature. 2014;505:87–91. Available from: http://dx.doi.org/10.1038/nature12736

120. Skourtanioti E, Erdal YS, Frangipane M, Balossi Restelli F, Yener KA, Pinnock F, et al. Genomic History of Neolithic to Bronze Age Anatolia, Northern Levant, and Southern Caucasus. Cell. 2020;181:1158–75.e28. Available from: http://dx.doi.org/10.1016/j.cell.2020.04.044

121. van de Loosdrecht M, Bouzouggar A, Humphrey L, Posth C, Barton N, Aximu-Petri A, et al. Pleistocene North African genomes link Near Eastern and sub-Saharan African human populations. Science. 2018;360:548–52. Available from: http://dx.doi.org/10.1126/science.aar8380

122. van den Brink ECM, Beeri R, Kirzner D, Bron E, Cohen-Weinberger A, Kamaisky E, et al. A Late Bronze Age II clay coffin from Tel Shaddud in the Central Jezreel Valley, Israel: context and historical implications. Levantina. 2017;49:105–35. Available from: https://doi.org/10.1080/00758914.2017.1368204

123. Wang C-C, Reinhold S, Kalmykov A, Wissgott A, Brandt G, Jeong C, et al. Ancient human genome-wide data from a 3000-year interval in the Caucasus corresponds with eco-geographic regions. Nat Commun. 2019;10:590. Available from: http://dx.doi.org/10.1038/s41467-018-08220-8

124. Yu H, van de Loosdrecht MS, Mannino MA, Talamo S, Rohrlach AB, Childebayeva A, et al. Genomic and dietary discontinuities during the Mesolithic and Neolithic in Sicily. iScience. 2022;25:104244. Available from: http://dx.doi.org/10.1016/j.isci.2022.104244

125. Ingman T, Eisenmann S, Skourtanioti E, Akar M, Ilgner J, Gnecchi Ruscone GA, et al. Human mobility at Tell Atchana (Alalakh), Hatay, Turkey during the 2nd millennium BC: Integration of isotopic and genomic evidence. PLoS One. 2021;16:e0241883. Available from: http://dx.doi.org/10.1371/journal.pone.0241883

126. Skourtanioti E, Ringbauer H, Gnecchi Ruscone GA, Bianco RA, Burri M, Freund C, et al. Ancient DNA reveals admixture history and endogamy in the prehistoric Aegean. Nat Ecol Evol. 2023;7:290–303. Available from: http://dx.doi.org/10.1038/s41559-022-01952-3

127. Reitsema LJ, Mittnik A, Kyle B, Catalano G, Fabbri PF, Kazmi ACS, et al. The diverse genetic origins of a Classical period Greek army. Proc Natl Acad Sci U S A. 2022;119:e2205272119. Available from: http://dx.doi.org/10.1073/pnas.2205272119

128. Haag J, Jordan AI, Stamatakis A. Pandora: A Tool to Estimate Dimensionality Reduction

Stability of Genotype Data. bioRxiv. 2024 [cited 2024 Aug 27]. p. 2024.03.14.584962. Available from: https://www.biorxiv.org/content/10.1101/2024.03.14.584962v1

129. Alexander DH, Novembre J, Lange K. Fast model-based estimation of ancestry in unrelated individuals. Genome Res. 2009;19:1655–64. Available from: http://dx.doi.org/10.1101/gr.094052.109

130. Chang CC, Chow CC, Tellier LC, Vattikuti S, Purcell SM, Lee JJ. Second-generation PLINK: rising to the challenge of larger and richer datasets. Gigascience. 2015;4:7. Available from: http://dx.doi.org/10.1186/s13742-015-0047-8

131. Davis TL. Command Line Optional and Positional Argument Parser [R package argparse version 2.2.3]. 2024 [cited 2024 Jul 29]; Available from: https://CRAN.R-project.org/package=argparse

132. Analytics R, Weston S. Foreach Parallel Adaptor for "parallel" [R package doMC version 1.3.8]. 2022 [cited 2024 Jul 29]; Available from: https://CRAN.R-project.org/package=doMC

133. Microsoft, Weston. Provides Foreach Looping Construct [R package foreach version 1.5.2]. 2022 [cited 2024 Jul 29]; Available from: https://CRAN.R-project.org/package=foreach

134. van den Brand T. Hacks for "ggplot2" [R package ggh4x version 0.2.8]. 2024 [cited 2024 Jul 29]; Available from: https://CRAN.R-project.org/package=ggh4x

135. Arnold JB. Extra Themes, Scales and Geoms for "ggplot2" [R package ggthemes version 5.0.0]. 2023 [cited 2024 Jul 29]; Available from: https://CRAN.R-project.org/package=ggthemes

136. Auguie B. Miscellaneous Functions for "Grid" Graphics [R package gridExtra version 2.3]. 2017 [cited 2024 Jul 29]; Available from: https://CRAN.R-project.org/package=gridExtra

137. Wickham H. Reshaping Data with the reshape Package. J Stat Softw. 2007 [cited 2024 Jul 8];21:1–20. Available from: http://www.jstatsoft.org/v21/i12/.

138. Wickham H. Simple, Consistent Wrappers for Common String Operations [R package stringr version 1.5.1]. 2023 [cited 2024 Jul 29]; Available from: https://CRAN.R-project.org/package=stringr

139. Harney É, Patterson N, Reich D, Wakeley J. Assessing the performance of qpAdm: a statistical tool for studying population admixture. Genetics. 2021;217. Available from: http://dx.doi.org/10.1093/genetics/iyaa045

140. Maier R, Flegontov P, Flegontova O, Işıldak U, Changmai P, Reich D. On the limits of fitting complex models of population history to f-statistics. Elife. 2023;12. Available from: http://dx.doi.org/10.7554/eLife.85492

141. Lazaridis I, Alpaslan-Roodenberg S, Acar A, Açıkkol A, Agelarakis A, Aghikyan L, et al. The genetic history of the Southern Arc: A bridge between West Asia and Europe. Science. 2022;377:eabm4247. Available from: http://dx.doi.org/10.1126/science.abm4247

142. Maier R, Patterson N. admixtools: Inferring demographic history from genetic data. 2024. Available from: https://uqrmaie1.github.io/admixtools/

143. Harrell FE Jr. Harrell Miscellaneous [R package Hmisc version 5.1-3]. 2024 [cited 2024 Aug 9]; Available from: https://CRAN.R-project.org/package=Hmisc

144. Chaitanya L, Breslin K, Zuñiga S, Wirken L, Pośpiech E, Kukla-Bartoszek M, et al. The

HIrisPlex-S system for eye, hair and skin colour prediction from DNA: Introduction and forensic developmental validation. Forensic Sci Int Genet. 2018;35:123–35. Available from: http://dx.doi.org/10.1016/j.fsigen.2018.04.004

145. Walsh S, Liu F, Wollstein A, Kovatsi L, Ralf A, Kosiniak-Kamysz A, et al. The HIrisPlex system for simultaneous prediction of hair and eye colour from DNA. Forensic Sci Int Genet. 2013;7:98–115. Available from: http://dx.doi.org/10.1016/j.fsigen.2012.07.005

146. Maróti Z, Nyerki E, Neparaczki E, Török T, Varga GI, Kalmár T. aHISplex: an imputation based method for eye, hair and skin colour prediction from low coverage ancient DNA. bioRxiv. 2023 [cited 2024 Jul 8]. p. 2023.11.02.565295. Available from: https://www.biorxiv.org/content/10.1101/2023.11.02.565295v1

147. Brace S, Diekmann Y, Booth TJ, van Dorp L, Faltyskova Z, Rohland N, et al. Ancient genomes indicate population replacement in Early Neolithic Britain. Nat Ecol Evol. 2019;3:765–71. Available from: http://dx.doi.org/10.1038/s41559-019-0871-9

148. Boussiou M, Karababa P, Sinopoulou K, Tsaftaridis P, Plata E, Loutradi-Anagnostou A. The molecular heterogeneity of β-thalassemia in Greece. Blood Cells Mol Dis. 2008;40:317–9. Available from: https://www.sciencedirect.com/science/article/pii/S1079979607002549

149. Georgiou I, Makis A, Chaidos A, Bouba I, Hatzi E, Kranas V, et al. Distribution and frequency of beta-thalassemia mutations in northwestern and central Greece. Eur J Haematol. 2003;70:75–8. Available from: http://dx.doi.org/10.1034/j.1600-0609.2003.00017.x

150. Breitwieser FP, Baker DN, Salzberg SL. KrakenUniq: confident and fast metagenomics classification using unique k-mer counts. Genome Biol. 2018;19:198. Available from: http://dx.doi.org/10.1186/s13059-018-1568-0

151. Herbig A, Maixner F, Bos KI, Zink A, Krause J, Huson DH. MALT: Fast alignment and analysis of metagenomic DNA sequence data applied to the Tyrolean Iceman. bioRxiv. 2016 [cited 2024 Jul 2]. p. 050559. Available from: https://www.biorxiv.org/content/10.1101/050559v1

152. Hübler R, Key FM, Warinner C, Bos KI, Krause J, Herbig A. HOPS: automated detection and authentication of pathogen DNA in archaeological remains. Genome Biol. 2019;20:280. Available from: http://dx.doi.org/10.1186/s13059-019-1903-0

153. Pochon Z, Bergfeldt N, Kırdök E, Vicente M, Naidoo T, van der Valk T, et al. aMeta: an accurate and memory-efficient ancient metagenomic profiling workflow. Genome Biol. 2023;24:242. Available from: http://dx.doi.org/10.1186/s13059-023-03083-9

154. R Core Team. R: A Language and Environment for Statistical Computing. R Foundation for Statistical Computing, Vienna, Austria. 2023;

155. Massicotte P, South A. World Map Data from Natural Earth [R package rnaturalearth version 1.0.1]. 2023 [cited 2024 Aug 6]; Available from: https://CRAN.R-project.org/package=rnaturalearth

156. Pebesma E. Simple Features for R [R package sf version 1.0-16]. 2024 [cited 2024 Aug 6]; Available from: https://CRAN.R-project.org/package=sf

157. Urbanek S. proj4: A simple interface to the PROJ.4 cartographic projections library. [cited 2024 Aug 6]; Available from: https://CRAN.R-project.org/package=proj4

158. Wickham H. ggplot2. Springer International Publishing; [cited 2024 Jul 5]. Available from:

https://link.springer.com/book/10.1007/978-3-319-24277-4

159. Slowikowski K. Automatically Position Non-Overlapping Text Labels with “ggplot2” [R package ggrepel version 0.9.5]. 2024 [cited 2024 Aug 7]; Available from: https://CRAN.R-project.org/package=ggrepel

160. Waskom M. seaborn: statistical data visualization. J Open Source Softw. 2021;6:3021. Available from: https://joss.theoj.org/papers/10.21105/joss.03021

161. Hugh-Jones D. ggmagnify: Create a Magnified Inset of Part of a “Ggplot” Object. https://github.com/hughjonesd/ggmagnify. 2024;

162. Wilke CO, Wiernik BM. Improved Text Rendering Support for “ggplot2” [R package ggtext version 0.1.2]. 2022 [cited 2024 Sep 10]; Available from: https://CRAN.R-project.org/package=ggtext

163. Hoppe KA, Koch PL, Furutani TT. Assessing the preservation of biogenic strontium in fossil bones and tooth enamel. Int J Osteoarchaeol. 2003;13:20–8. Available from: https://onlinelibrary.wiley.com/doi/10.1002/oa.663

164. Hillson S. Dental Anthropology. Cambridge University Press; 1996. Available from: https://play.google.com/store/books/details?id=WIcgAwAAQBAJ

165. Price TD, Burton JH, Bentley RA. The Characterization of Biologically Available Strontium Isotope Ratios for the Study of Prehistoric Migration. Archaeometry. 2002 [cited 2024 Aug 22];44:117–35. Available from: https://onlinelibrary.wiley.com/doi/abs/10.1111/1475-4754.00047

166. Evans J, Montgomery J, Wildman G. Isotope domain mapping of 87Sr/86Sr biosphere variation on the Isle of Skye, Scotland. J Geol Soc London. 2009 [cited 2024 Aug 22];166:617–31. Available from: http://nora.nerc.ac.uk/id/eprint/7960/

167. Alexander Bentley R. Strontium Isotopes from the Earth to the Archaeological Skeleton: A Review. Journal of Archaeological Method and Theory. 2006 [cited 2024 Aug 22];13:135–87. Available from: https://link.springer.com/article/10.1007/s10816-006-9009-x

168. Nafplioti A. Moving Forward: Strontium Isotope Mobility Research in the Aegean. Mediterranean Archaeology and Archaeometry. 2021 [cited 2024 Aug 22];21:165–79. Available from: https://www.academia.edu/76198598/%CE%9Coving_Forward_Strontium_Isotope_Mobility_Research_in_the_Aegean

169. Nafplioti A. Tracing population mobility in the Aegean using isotope geochemistry: a first map of local biologically available 87Sr/86Sr signatures. J Archaeol Sci. 2011 [cited 2024 Aug 22];38:1560–70. Available from: http://dx.doi.org/10.1016/j.jas.2011.02.021

170. Nafplioti A. “Mycenaean” political domination of Knossos following the Late Minoan IB destructions on Crete: negative evidence from strontium isotope ratio analysis (87Sr/86Sr). J Archaeol Sci. 2008 [cited 2024 Aug 22];35:2307–17. Available from: http://dx.doi.org/10.1016/j.jas.2008.03.006

171. Faure G. Principles of Isotope Geology. New York: John Wile and Sons; 1986. Available from: https://books.google.com/books/about/Principles_of_Isotope_Geology.html?hl=&id=xlfwAAAAMAAJ

172. Graeme R, Christopher JH. A geochemical traverse across the North Chilean Andes: evidence for crust generation from the mantle wedge. Earth Planet Sci Lett. 1989 [cited 2024 Aug 22];91:271–85. Available from: http://dx.doi.org/10.1016/0012-821X(89)90003-4

173. Wright LE. Identifying immigrants to Tikal, Guatemala: Defining local variability in strontium isotope ratios of human tooth enamel. J Archaeol Sci. 2005 [cited 2024 Aug 22];32:555–66. Available from: http://dx.doi.org/10.1016/j.jas.2004.11.011

174. Palmer MR, Elderfield H. Sr isotope composition of sea water over the past 75 Myr. Nature. 1985 [cited 2024 Aug 22];314:526–8. Available from: https://www.nature.com/articles/314526a0

175. Elderfield H. Strontium isotope stratigraphy. Palaeogeogr Palaeoclimatol Palaeoecol. 1986 [cited 2024 Aug 22];57:71–90. Available from: http://dx.doi.org/10.1016/0031-0182(86)90007-6

176. Blum JD, Taliaferro EH, Weisse MT, Holmes RT. Changes in Sr/Ca, Ba/Ca and 87Sr/86Sr ratios between trophic levels in two forest ecosystems in the northeastern U.S.A. Biogeochemistry. 2000 [cited 2024 Aug 22];49:87–101. Available from: https://link.springer.com/article/10.1023/A:1006390707989

177. Graustein WC. 87Sr/86Sr Ratios Measure the Sources and Flow of Strontium in Terrestrial Ecosystems. Stable Isotopes in Ecological Research. 1989 [cited 2024 Aug 22];491–512. Available from: https://link.springer.com/chapter/10.1007/978-1-4612-3498-2_28

178. Veizer J. Strontium Isotopes in Seawater through Time. Annu Rev Earth Planet Sci. 1989 [cited 2024 Aug 29]; Available from: https://doi.org/10.1146/ANNUREV.EA.17.050189.001041

179. Miller EK, Panek JA, Friedland AJ, Kadlecek J, Mohnen VA. Atmospheric deposition to a high-elevation forest at Whiteface Mountain, New York, USA. Tellus B Chem Phys Meteorol. 1993 [cited 2024 Aug 29];45:209–27. Available from: https://onlinelibrary.wiley.com/doi/abs/10.1034/j.1600-0889.1993.t01-2-00001.x

180. Ubelaker DH. Human Skeletal Remains: Excavation, Analysis, Interpretation. Washington: Taraxacum Press; 1989. Available from: https://books.google.com/books/about/Human_Skeletal_Remains.html?hl=&id=5bfczgEACAAJ

181. Chiaradia M, Gallay A, Todt W. Different contamination styles of prehistoric human teeth at a Swiss necropolis (Sion, Valais) inferred from lead and strontium isotopes. Appl Geochem. 2003 [cited 2024 Aug 22];18:353–70. Available from: http://dx.doi.org/10.1016/S0883-2927(02)00072-0

182. Lee-Thorp JA, Sponheimer M. Three case studies used to reassess the reliability of fossil bone and enamel isotope signals for paleodietary studies. Journal of Anthropological Archaeology. 2003 [cited 2024 Aug 22];22:208–16. Available from: http://dx.doi.org/10.1016/S0278-4165(03)00035-7

183. Kohn MJ, Schoeninger MJ, Barker WW. Altered states: Effects of diagenesis on fossil tooth chemistry. GeCoA. 1999 [cited 2024 Aug 22];63:2737–47. Available from: https://ui.adsabs.harvard.edu/abs/1999GeCoA..63.2737K/abstract

184. Price TD, Schoeninger MJ, Armelagos GJ. Bone chemistry and past behavior: an overview. JHumE. 1985 [cited 2024 Aug 22];14:419–47. Available from: https://ui.adsabs.harvard.edu/abs/1985JHumE..14..419P/abstract

185. Hedges REM, Clement JG, Thomas CDL, O'Connell TC. Collagen turnover in the adult femoral mid-shaft: Modeled from anthropogenic radiocarbon tracer measurements. Am J Phys Anthropol. 2007 [cited 2024 Aug 22];133:808–16. Available from: https://onlinelibrary.wiley.com/doi/abs/10.1002/ajpa.20598

186. Tafuri MA, Alexander Bentley R, Manzi G, di Lernia S. Mobility and kinship in the prehistoric Sahara: Strontium isotope analysis of Holocene human skeletons from the Acacus Mts. (southwestern Libya). Journal of Anthropological Archaeology. 2006 [cited 2024 Aug 22];3:390–402. Available from: https://www.infona.pl//resource/bwmeta1.element.elsevier-feb8936c-ce90-3401-b94d-dfc7249a1351

187. Manolagas SC. Birth and Death of Bone Cells: Basic Regulatory Mechanisms and Implications for the Pathogenesis and Treatment of Osteoporosis*. Endocr Rev. 2000 [cited 2024 Aug 22];21:115–37. Available from: https://academic.oup.com/edrv/article-pdf/21/2/115/8859550/edrv0115.pdf

188. Ezzo J, Price T. Migration, Regional Reorganization, and Spatial Group Composition at Grasshopper Pueblo, Arizona. J Archaeol Sci. 2002 [cited 2024 Aug 22]; Available from: https://doi.org/10.1006/JASC.2001.0745

189. Price TD, Johnson CM, Ezzo JA, Ericson J, Burton JH. Residential Mobility in the Prehistoric Southwest United States: A Preliminary Study using Strontium Isotope Analysis. JArSc. 1994 [cited 2024 Aug 22];21:315–30. Available from: https://ui.adsabs.harvard.edu/abs/1994JArSc..21..315P/abstract

190. Higgins MD, Higgins RA. A Geological Companion to Greece and the Aegean. Cornell University Press; 1996. Available from: https://books.google.com/books/about/A_Geological_Companion_to_Greece_and_the.html?hl=&id=Q-0jraoNqvUC

191. Nafplioti A. Population movement, biological and cultural interactions in the EBA Aegean: The case of Manika. Unpublished report submitted to the American School of Classical Studies at Athens (ASCSA) for research carried out through a J.L. Angel Fellowship in Human Skeletal Studies. 2008 [cited 2024 Sep 5]. Available from: https://www.ascsa.edu.gr/research/wiener-laboratory/research/research-archive

2784
